# Maternal lineages from 10-11^th^ century commoner cemeteries of the Carpathian Basin

**DOI:** 10.1101/2021.01.26.428268

**Authors:** Kitti Maár, Gergely I. B. Varga, Bence Kovács, Oszkár Schütz, Zoltán Maróti, Tibor Kalmár, Emil Nyerki, István Nagy, Dóra Latinovics, Balázs Tihanyi, Antónia Marcsik, György Pálfi, Zsolt Bernert, Zsolt Gallina, Sándor Varga, László Költő, István Raskó, Tibor Török, Endre Neparáczki

## Abstract

**Background:** Nomadic groups of conquering Hungarians played a predominant role in Hungarian prehistory, but genetic data are available only from the immigrant elite strata. Most of the 10-11th century remains in the Carpathian Basin belong to common people, whose origin and relation to the immigrant elite has been widely debated.

**Methods:** Mitogenome sequences were determined from 202 individuals with Next Generation Sequencing combined with hybridization capture. Median Joining networks were used for phylogenetic analysis. The commoner population was compared to 87 ancient Eurasian populations with sequence based (Fst) and haplogroup based population genetic methods.

**Results:** Haplogroup composition of the commoner population markedly differs from the elite and in contrast to the elite, commoners cluster with European populations. Besides, detectable sub-haplogroup sharing indicates admixture between the elite and commoners.

**Conclusions:** Majority of the 10-11^th^ century commoners most likely represent local populations of the Carpathian Basin, which admixed with the eastern immigrant groups including conquering Hungarians.

## 1. Introduction

Hungarian history was profoundly determined by the conquering Hungarians (succinctly Conquerors), which arrived to the Carpathian Basin from the Eastern European steppe at the end of the 9^th^ century AD as an alliance of seven tribes. Leaders of the alliance, Álmos and his son Árpád founded a steppe state upon the ashes of the Avar Khaganate [1,2], and their descendants later established the Hungarian Kingdom. The archeological legacy of the Conquerors is well defined, especially the 10^th^ century small cemeteries of the military leader strata whose grave finds included precious metal jewels, costume ornaments as well as decorated horse riding- and weapon-related grave goods. Most of the larger cemeteries attributed to the common people are dated somewhat later, to the 10-12^th^ centuries. People in these so called village cemeteries were buried with simpler jewels and grave goods, with sporadic appearance of weapons or harness accessories. There is a general agreement that elite graves with typical grave goods represent first or second generation immigrant Conquerors, but affiliation of people in the village cemeteries is far less clear. The several hypothesis presumed among others Slavic ethnicity, immigrant commoner strata of the conquering Hungarians, and local inhabitants from the previous Avar period (see Appendix A for details). Origin of the commoner strata considerably determines the historical interpretation of the conquer and subsequent events in the Carpathian Basin and genetic data may contribute to clarify this issue.

Hitherto most genetic studies were focused on the elite graves, as these promised an answer for the origin of the immigrant groups. In [3] HVR sequences of 76 individuals were investigated, picked from 23 cemeteries mainly representing 10^th^ century elite graves, from which they identified 23% east Eurasian and 77% west Eurasian maternal lineages. Another study [4], aimed at characterizing the population of entire elite cemeteries, and sequenced 102 mitogenomes of which 30% had Central-Inner Asian maternal ancestry, while most of the remaining lineages originated from western Eurasia. Y-chromosome studies [5] found that male lineages had similar phylogeographic composition to female ones, thus all studies had congruent results, inferring that the Conqueror elite population was assembled from an admixture of Asian and European groups on the Pontic steppe.

This raises the question whether the commoners were genetically similar to the elite, if so they could be one and the same population, or the poorer strata had different origin? This question had been addressed in the first HVR-based study [6], in which 27 selected graves from 15 cemeteries were grouped according to grave goods and the population with “classical” grave goods were found to contain higher proportion of east Eurasian Hgs than the group with poor archaeological remains. However, this conclusion was based on small sample size and low resolution HVR study and a systematic characterization of the commoner population with representative dataset has not been performed yet.

We set out to implement a comprehensive study in this matter and for this end selected 8 cemeteries archaeologically evaluated as belonging to the 10-11th century commoners, from which we determined altogether 202 whole mitogenome sequences. Phylogenetic analysis was performed to illuminate the origin of each maternal sub-lineages of the studied remains. We compared the mitochondrial haplogroup composition of the commoner and elite population to find out their genetic relations and applied different population genetic methods to elucidate the relationship of the commoners with other ancient Eurasian populations. For this reason, we also built a comprehensive database of ancient Eurasian populations including all available published mitogenome data.

## 2. Materials and Methods

### 2.1 Archeological background

In contrast to the small 10^th^ century cemeteries with characteristic grave goods [7] representing the conquering Hungarian elite (ConqE) archaeologists classify large 10-11^th^ century cemeteries containing poor grave goods with sporadic appearance of ConqE findings (see Appendix A for details) as belonging to the Hungarian commoners (ConqC). We collected petrous bones or where it was unavailable teeth from 220 human remains from 9 archeological sites (Figure 1), associated with Hungarian commoners. We made an effort to carry out representative sampling, thus graves were selected from each part of the cemeteries, including males and females from burials both with and without grave goods.

**Figure 1.**
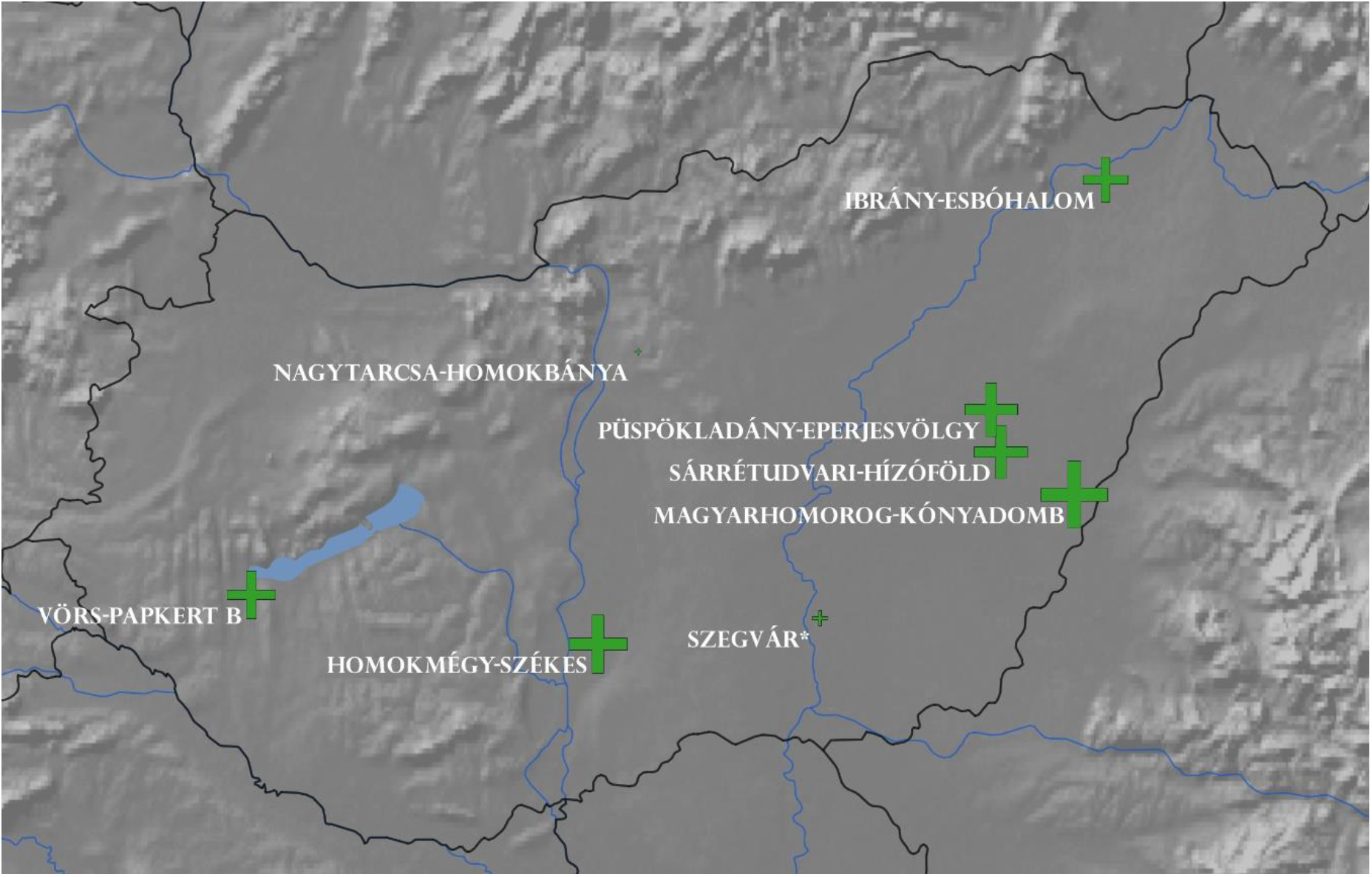
Locations of graveyards of the Hungarian commoners (ConqC) under study. Sizes of the crosses are proportional to the number of examined remains. Asterisk (*) denotes that two nearby cemeteries, Szegvár-Oromdűlő and Szegvár-Szőlőkalja were studied from Szegvár. The map was generated using QGIS 3.12.0 [14]

The largest 10th century commoner cemetery with 262 graves was excavated in Sárrétudvari-Hízóföld [8], with high proportion of graves containing archery equipment and stirrups. We selected 32 new samples from this site, of which 31 mitogenomes could be determined. Further 8 sequences were added from our previous study [4,9].

Another large commoner cemetery with 637 graves is located in the nearby Püspökladány-Eperjesvölgy [8]. This cemetery was used from the 10th to the early 12th centuries, therefore contains a “pagan”and “Christian” part. We sampled altogether 36 remains from both part of the graveyard and could obtain 31 mitogenomes.

The Ibrány-Esbó-halom commoner cemetery with 269 graves was dated to the 10-11th century [10], thus also stretches into the Christian era. We analyzed 32 remains from this site resulting in 26 mitogenomes.

We studied 36 remains from the Homokmégy-Székes cemetery excavated at the Duna-Tisza Interfluve [11] with 206 graves, which was referred by the archaeologist as “typical cemetery of conquer period commoners”, and obtained 34 mitogenomes.

Among the studied cemeteries Magyarhomorog-Kónyadomb [12] is an exceptional case, as archaeologically it can be divided into two parts: A small part with 17 individuals was used by the 10th century Conqueror elite, while the larger part with 523 graves by 11-12th century commoners raising the question of potential continuity. We sequenced 14 samples from the elite part and processed 27 samples from the commoner part resulting in 25 mitogenomes.

From the Transdanubia region, we included the Vörs-Papkert-B cemetery [13], which was in use between the 8-11th centuries and accordingly the 716 excavated burials are mostly from the late Avar and Carolingian periods, but 33 people can be dated to the time of the Hungarian conquest. The uninterrupted usage of this graveyard raises the possibility that it might represent the same population in the subsequent periods, thus we sampled graves from each period, namely 9 from the Avar (8 mitogenomes), 11 from Carolingian, and 10 from the Hungarian conquer period (9 mitogenomes).

Finally, we complemented our sample set with few individuals from the Nagytarcsa-Homokbánya (n=4), Szegvár-Oromdűlő (n=4) and Szegvár-Szőlőkalja (n=5) commoner cemeteries. All of the 13 samples came from poor burials or from graves devoid of archaeological grave goods. For detailed description of the sites and samples see Appendix A and Table S1.

### 2.2 Library preparation, sequencing and Hg assignment

All pre-PCR laboratory procedures leading to Next-Generation Sequencing (NGS) were conducted in the common ancient DNA laboratory of the Department of Archaeogenetics of the Institute of Hungarian Research and Department of Genetics, University of Szeged, Hungary. Details of the ancient DNA purification, library preparation, hybridization capture, sequencing and sequence analysis method are given in [9]. All libraries were made from partial uracil-DNA-glycosylase (UDG) treated DNA extracts. We estimated the endogenous human DNA content of each library with low coverage shotgun sequencing (Table S2a) then mitogenomes from samples with similar proportions of human DNA content were pooled and enriched together. Captured and amplified libraries were purified on MinElute columns. Quantity and quality measurements were performed with Qubit fluorometric quantification system and TapeStation automated electrophoresis system (Agilent). Further 13 mitogenome sequences were determined from whole genome sequencing as indicated in Table S2.

Ancient DNA damage patterns were assessed using MapDamage 2.0 [15] and read quality scores were modified with the rescale option to account for post-mortem damage. Mitochondrial genome contamination was estimated using Schmutzi algorithm [16] (Table S2b). Mitochondrial haplogroup (Hg) determination was performed using HaploGrep v2.1.25 [17] (Table S3a). Biological sex of the individuals was identified according to [18] based on X/Y ratio of reads from shotgun sequencing.

The raw nucleotide sequence data of the 202 samples were deposited to the European Nucleotide Archive (http://www.ebi.ac.uk/ena) under accession number: PRJEB40566.

### 2.3 Assembling an ancient Eurasian mitogenome database

For the phylogenetic and population genetic analyses we built a database containing 4191 published ancient Eurasian mitogenomes (Table S4). Sequences were downloaded from the NCBI and European Nucleotide Archive databases and where it was necessary mitogenome sequences were sorted out from whole genomes. This database was then augmented with the 202 new mitogenomes from this study. We ordered the published samples into 88 populations based on time period, archeological site and context, as well as classification of published genome data. In cases when populations were underrepresented due to low sample size, we grouped samples from related cultures like Alans and Saltovo-Mayaki, Medieval samples from Italy, Germany and England, Iberian Chalcolithic and Bronze Age, Chalcolithic samples from Iran and Turan, early and late Sarmatians etc. (Table S4).

### 2.4 Phylogenetic and population genetic study

A subset of the published sequences was of poor quality, we excluded sequences with >5% missing data from the phylogenetic and Fst analysis, and used 3844 fasta files of ancient and 11682 modern sequences for building median joining (MJ) networks as described in [4]. Phylogeographic origin of samples were assessed from the geographic origin of the nearest Hgs. We distinguished four regions; east Eurasia, west Eurasia, Eurasia and Caucasus-Middle East (Figure S1).

For population genetic analysis we merged all 169 ConqC data to a single population (Tables S3c and S4) excluding members of the elite Magyarhomorog cemetery, as well as Avar and Caroling samples from the Vörs-Papkert cemetery (excluded samples are color labelled in Table S2b). These were supplemented with 13 commoner mitogenomes published previously [4]. The merged ConqC population was compared to the 88 ancient Eurasian groups from the newly assembled mitogenome database, including the previously published military elite strata of the Conquerors [4,19,20], which was supplemented with the Magyarhomorog elite graveyard data from present study (Tables S3c and S4).

Three independent methods were applied to measure the genetic distances of ConqC from other ancient populations. In the first analysis we reduced the Hg assignments of all samples to major Hgs as listed in Figure 4, which decreased population data to 34 dimensions, what is sufficient to display main correlations. Then major Hg frequencies were calculated and Principal Component Analysis (PCA) was conducted employing the function prcomp in R 3.6.3. [21]. We also compared the major Hg frequencies of the ConqC and ConqE groups separately.

In a second approach traditional sequence based method was implemented calculating pair-wise population differentiation values (Fst) with Arlequin 3.5.2.2 [22] from entire mtDNA genomes, as described in [4]. Multidimensional scaling (MDS) was applied on the matrix of linearized Slatkin Fst values [23] and the values were visualized in the two-dimensional space using the cmdscale function implemented in R 3.6.3 [21].

In a third approach Shared Haplogroup Distance (SHD) values were measured between the populations according to our previous study [4], which calculates the frequency of identical terminal sub-Hgs between populations as these testify shared ancestry or past admixture.

## 3. Results

### 3.1. Sequencing results and assigned Haplogroups

We collected altogether 220 samples from the listed sites, but could not obtain DNA from 5 samples, another 10 samples were excluded from the analysis due to low mitogenome sequence coverage and 3 further samples due to high contamination. Using the NGS method combined with target enrichment, we determined 189 ancient mitogenome sequences and further 13 were determined from whole genome sequencing, thus we report 202 new mitogenomes in this paper (Table S3a). We obtained 4.2-3068x mitogenome coverage, average coverage was 231x. Schmutzi estimated negligible contamination for most of the 202 samples, 7 samples were indicated to carry significant (15-21%) contamination, nonetheless Schmutzi could determine the endogenous sequence unambiguously for these samples due to high coverage, enabling correct Hg assignment. For details of sequencing data see Table S2. On the ground of haplogroup determination by HaploGrep 2.0, the 202 samples belong to 154 sub-Hg-s and 187 different haplotypes (Table S3a).

### 3.2 Kinship analysis

We examined possible kinship relation between and within cemeteries. We detected 10 pairs of identical mitochondrial haplotypes within cemeteries and 4 pairs between cemeteries (Table S3b), which indicate potential direct maternal relationship of these individuals.

### 3.3. Phylogenetic analysis

As some of the mitochondrial subclades may have specific geographical distribution [24,25] we elucidated the phylogenetic relations of each mitogenome sequences using M-J Networks as shown in Figure S1. The closest sequence matches pointed at a well-defined geographical region in most cases, which is indicated next to the phylogenetic trees and is summarized on Figure 2.

**Figure 2.**
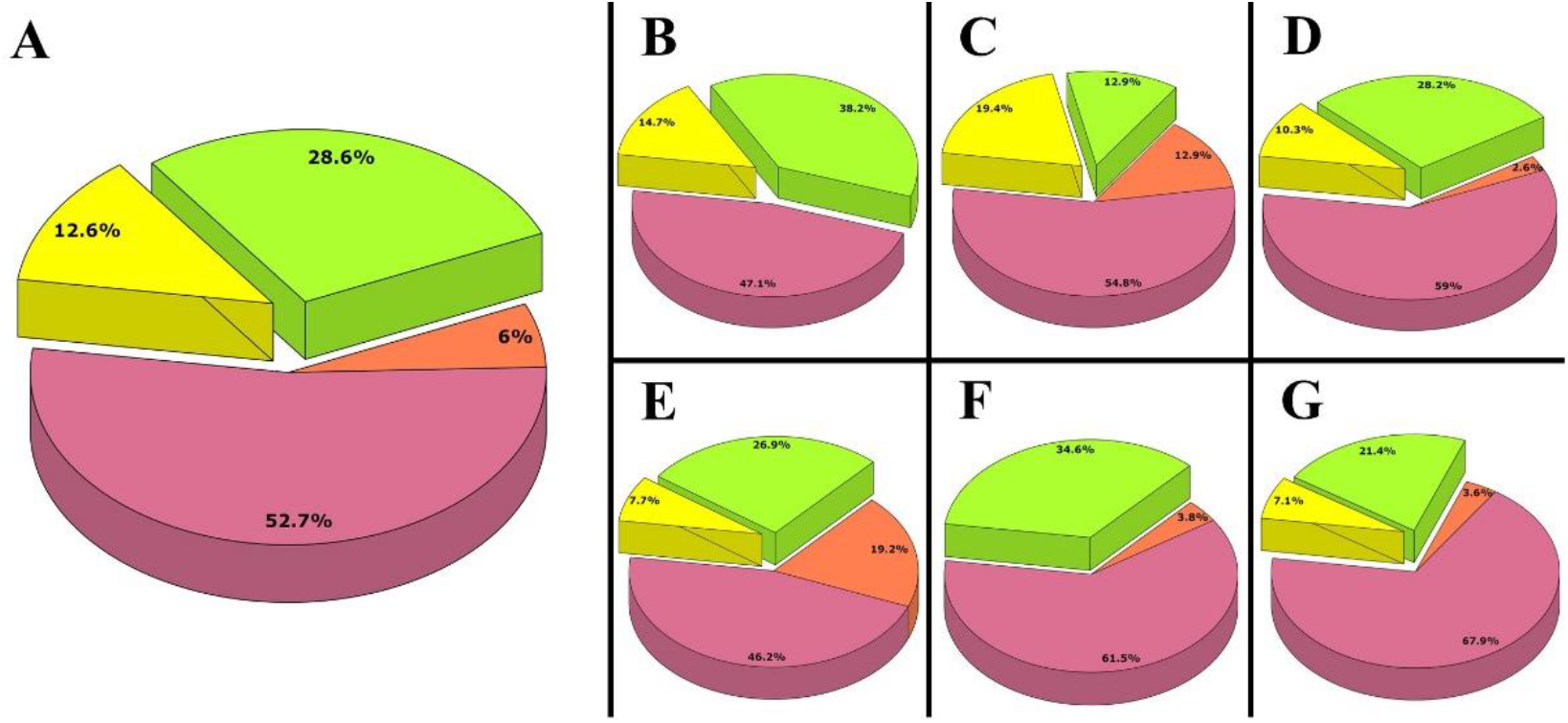
Phylogeographic origin of the ConqC maternal lineages from different cemeteries. Data are summarized from Figure S1 and from previous study [4]. West Eurasian Hgs are marked with pink, east Eurasian Hgs with yellow, Eurasian Hgs with green, Caucasus-Middle East Hgs with brown. (A) Distribution of merged data of 182 Hungarian commoner samples from all cemeteries. (B-G) Phylogeographic distribution of the maternal lineages from individual cemeteries: (B) Homokmégy-Székes (n=34); (C) Püspökladány-Eperjesvölgy (n=31); (D) Sárrétudvari-Hízóföld (n=39); (E) Ibrány-Esbóhalom (n=26); (F) Magyarhomorog-Kónyadomb (n=26, samples just from the commoner part); (G) Vörs-Papkert-B (n=28, including all samples from this cemetery).

Phylogenetic trees revealed that out of the 182 commoner maternal lineages 23 were unequivocally derived from east Eurasia, 107 from west Eurasia, while 52 are widespread throughout Eurasia. Out of the European lineages 11 have primarily Caucasus-Middle East distribution (Figure 2A).

### 3.4 Haplogroup composition of individual cemeteries

The 34 investigated samples from Homokmégy-Székes belonged to 30 Hgs (Table S3a), 47.1% of the lineages were of European and 14.7% of east Eurasian origin, while 38.2% showed general Eurasian distribution (Figure 2B).

From the Püspökladány-Eperjesvölgy cemetery 31 remains were analyzed. The maternal lineages were classified into 28 Hgs (Table S3a), and they showed 54.8% European, 19.4% east Eurasian, 12.9% Eurasian ancestry while 12.9% had Caucasus-Middle East affinity (Figure 2C).

The newly reported mitogenomes of 31 individuals from Sárrétudvari-Hízóföld belonged to 26 Hgs (Table S3a). In a previous study the mitochondrial lineage of 8 individuals from this cemetery were determined [4]. Merging these data 59% of the lineages had European, 10.3%, east Eurasian 28.2% Eurasian and 2.6% Caucasian-Middle East maternal ancestry (Figure 2D).

The Ibrány-Esbóhalom cemetery was represented by 26 samples falling to 26 different Hgs (Table S3a). 46.2% of the maternal lineages originated from Europe, 7.7% from east Eurasia and 19.2% from Caucasus-Middle East regions, while 26.9% of the lineages had Eurasian distribution (Figure 2E).

We sequenced 14 samples from the elite part of Magyarhomorog-Kónyadomb, and their Hg composition was very similar to those of previously studied elite cemeteries [4]; 35,7% of the lineages were of east Eurasian, of 42,9% European and 21,4% of Eurasian origin (Table S1) and their Hg content was also overlapping with the Karos and Kenézlő dataset supporting the archaeological evaluation, thus we included these data to the elite dataset (Table S3c). From the 11-12th century commoner part of Magyarhomorog we sequenced 25 samples which belonged into 22 mitochondrial Hgs (Table S3a), this was supplemented with one published sample from this site [4]. From the 26 samples 61.5% had west Eurasian, 34.6% Eurasian and 3.8% Caucasus-Middle East affinity (Figure 2F), thus genetic data also corroborated that the large graveyard represent a separable commoner population.

The cemetery of Vörs-Papkert is another special case, as it was used for centuries by successive populations of Avars, Carolingians and Conquerors. Evaluating the entire 28 sample set from this cemetery together (Figure 2G) it showed very similar overall picture to other commoner cemeteries, with 25 Hgs of which 67.9% had European, 7.1% east Eurasian, 21.4% Eurasian and 3.6% Caucasus-Middle-East affinity, suggesting moderate Asian impact from immigrant Avar and Conqueror groups. By all means for the population genetic analysis we removed Avar and Carolingian samples from this dataset.

We also investigated a few individuals from other commoner cemeteries, namely 4 samples from Nagytarcsa-Homokbánya, 4 from Szegvár-Oromdűlő and 5 from Szegvár-Szőlőkalja resulting in two east Eurasian lineages besides the European ones (Table S3a).

We acknowledge that the average of 30 samples per site may poorly represent the individual cemeteries, but the total number of 182 commoner remains (Table S3c) can be regarded considerably representative for population genetic analysis.

### 3.5. Population genetic analysis

First, we compared the major Hg distribution of the conqueror period elite and commoner populations (Figure 3). The heterogeneity of major-Hg distribution of ConqE is comparable to that of ConqC (22 and 19 main Hgs respectively), however the Hg composition of the two groups shows considerable differences. The ratio of east Eurasian major-Hgs in the commoners is 7.69% contrary to the 19.64% of the elite, besides the elite contains a broad spectrum of these; A, B, C, D, F, G, and Y, while only C and D occur with notable frequencies in the commoners, with single appearance of B.

**Figure 3.**
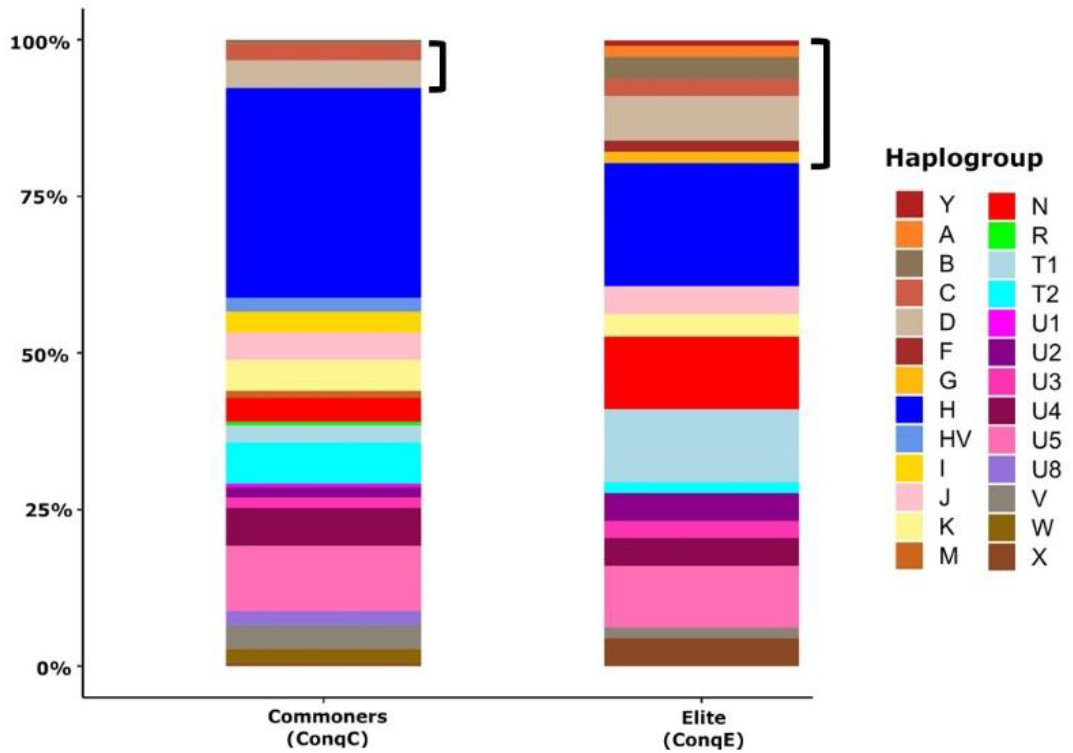
Comparison of major Hg distributions from ancient Hungarian populations. Major Hg distribution of Commoner samples (n=182) from this study is distinct from Conqueror Elite samples (n=112) taken from previous studies [4,19,20] and including elite data from Magyarhomorog of present study. Brackets mark East Eurasian Hgs.

West Eurasian Hgs of ConqC and ConqE also show notable differences: Hgs HV, I, M, R, U1, U8 and W occur with moderate frequencies in commoners, while these are completely absent from the elite. Three Hgs N, T1 and X, typically widespread both in east and west Eurasia, show much higher ratio in the elite than in commoners: N 11.61% in elite, 3.85% in commoners; T1 11.61% in elite, 2.75% in commoners; X 4.46% in elite and 0.55% in commoners. The opposite is true for Hgs H and T2; among commoners H is the most prevalent Hg with 33.52% frequency, while in the elite group its proportion is significantly lower 19.64%; T2 has 6.59% proportion in the commoners and 1.79% in the elite.

As the Hg composition of the studied commoner samples markedly differs from that of the elite, we measured ConqC genetic distances from ConqE as well as from 87 published ancient Eurasian populations (Table S5). PCA obtained from major Hg frequencies of 88 populations (Figure 4) highlights the distance between ConqE and ConqC. The ConqC clustered in the eastern side of the European aggregation, with closest genetic affinity to Baltic Bronze Age, Baltic Iron Age, Baltic Medieval, Bell Baker Germany and Great Britain Bronze Age populations and is not far away from the Steppe Early-Middle Bronze Age (Steppe_EMBA). In contrast the Conqueror Elite is located between ancient European and Asian populations and its closest clusters are Sarmatian Iron Age, Tien Shan Iron Age, Karasuk late Bronze Age and the two groups suggested to be in connection with the Conquerors [20]: the Cis-Ural Medieval and Uyelgi trans-Ural Medieval.

**Figure 4.**
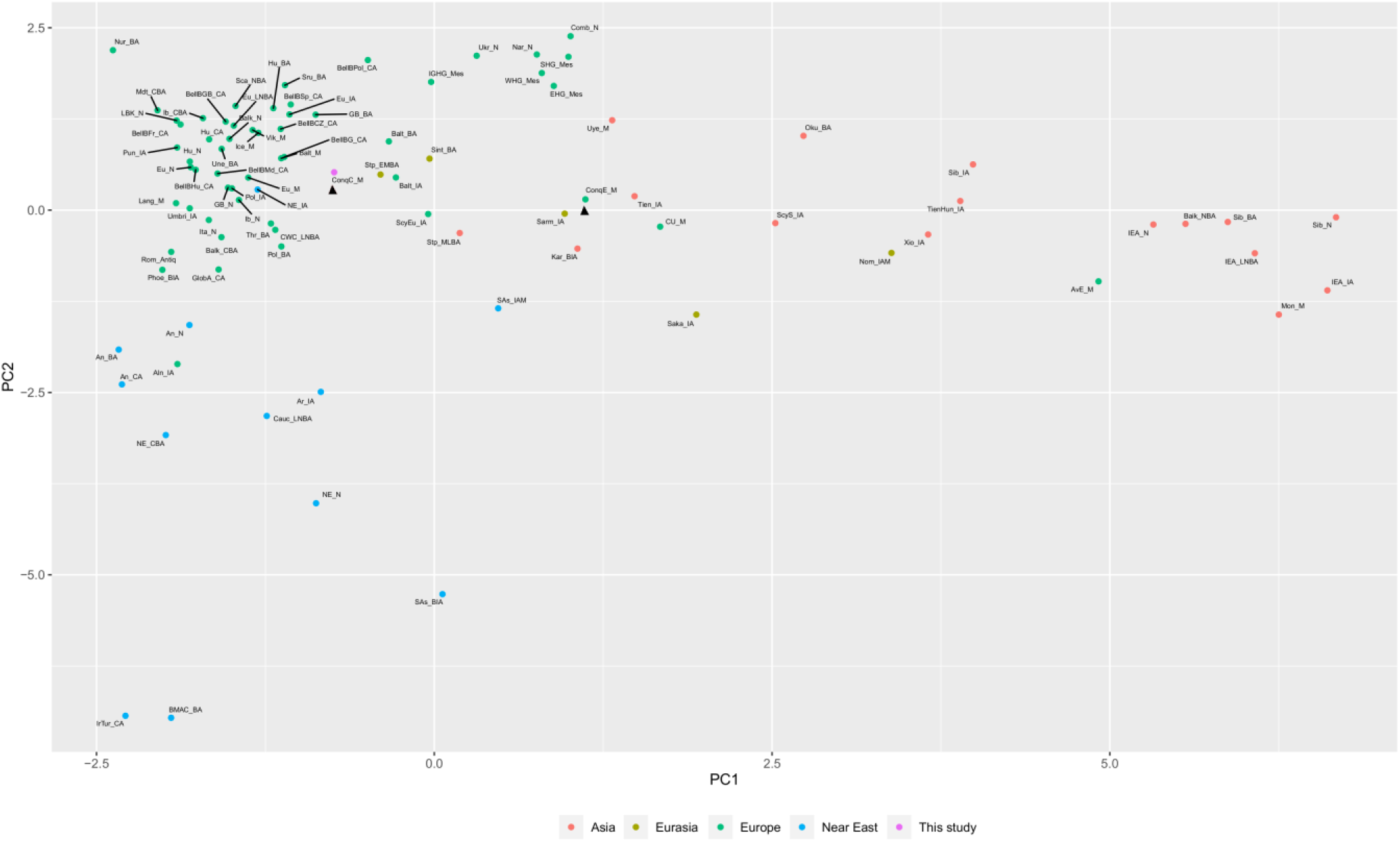
PCA plot of the major mtDNA haplogroup distribution (distinguishing Hgs A, B, C, D, F, G, H, HV, I, J, K, L, M, N, N1a, N1b, R, T, T1, T2, U, U1, U2, U3, U4, U5, U6, U7, U8, V, W, X, Y, Z) of 88 Eurasian populations. Abbreviations of population names are given in Table S4. Color shading denote Asian (salmon), Eurasian (olive), European (green), Near East (blue) populations, and the commoner group of this study (violet). ConqC and ConqE are highlighted with arrowheads. PC1 separates European populations to the left, Asian populations to the right side. PC2 separates Anatolian-Caucasus groups to the bottom, hunter-gatherers to top.

In order to further reveal the genetic relationships of ConqC with other ancient groups we draw an MDS plot (Figure. 5) from linearized Slatkin Fst values (Table S5a). Fst distances confirmed that ConqC is nearest to ancient European and Near Eastern populations; in the Pairwise FST matrix the closest groups are the European Medieval (0.0098), Anatolia Bronze Age (0.00991), Iceland Medieval (0.01433), pre-Roman (Umbri) Iron Age from Italy (0.01691) and Roman Antiquity (0.01701) followed by other European Bronze Age, Neolithic and Chalcolithic groups and accordingly these are located close on the MDS plot.

**Figure 5.**
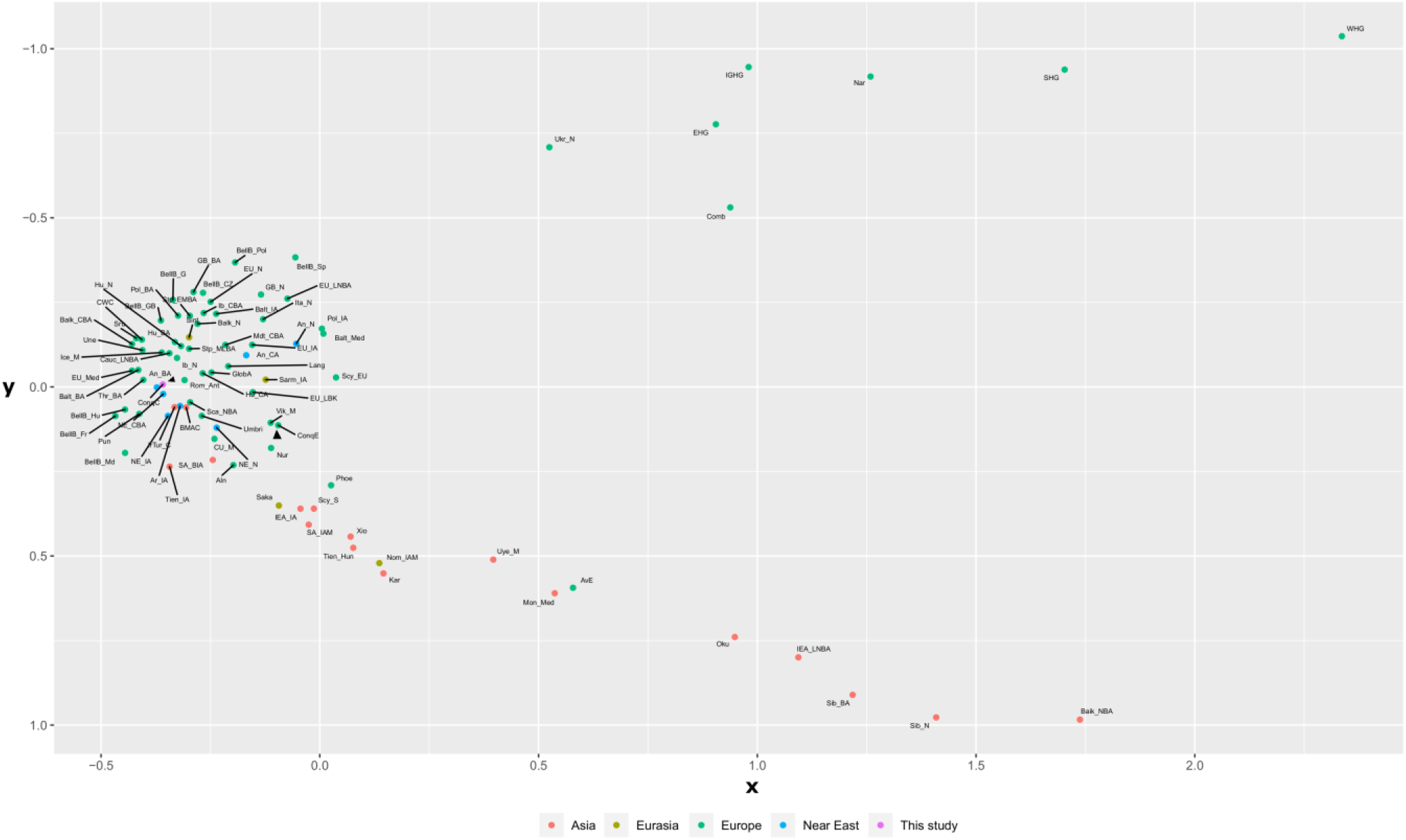
MDS plot from linearized Slatkin Fst values of Table S5a. Abbreviations of population names are given in Table S4. European populations are sequestered to the left, Asian populations to the bottom right, hunter-gatherers to the top right. Color shading denote Asian (salmon), Eurasian (olive), European (green), Near East (blue) populations, and the commoner group of this study (violet). ConqC and ConqE are highlighted with arrowheads.

The novel SHD population genetic method gave similar results, but also revealed new information (Table S5b). ConqE has the smallest SHD distance from ConqC followed by European populations from Neolithic to Medieval periods. It is also notable that SHD and Fst distances of Steppe_EMBA are comparable to that of European groups and European Scythian and Scytho-Siberian populations have noteworthy SHD distance as well, indicating that ConqC significantly shared sub-Hgs with these Eurasian steppe populations.

## 4. Discussion

In this paper an attempt was made to provide a genetic description of the common people of the Carpathian Basin who lived in the 10-11^th^ centuries, during the period of the Hungarian conquer. Of the 202 determined mitogenomes 169 belonged to commoners, while 14 samples from the Magyarhomorog cemetery were revealed to represent a small elite graveyard, not related to the adjacent commoner remains. Thus, beside the commoner samples we described a new small elite cemetery with 14 mitogenomes out of the 17 remains. The high frequency of N1a1a1a1a and T1a1 as well as the occurence of N1a1a1a1 and D4 in this small cemetery finds its best parallels in the Karos and Kenézlő elite graveyards [4]. From the Vörs-Papkert-B cemetery 11 Carolingian and 8 Avar Age samples were investigated next to the 9 ConqC ones. Hg H dominated this graveyard, as 16 out of the 28 remains carried Hg H irrespective of time period. A single D4e4 Hg was detected among the studied ConqC and a single A16 among the Avar period samples as weak signs of Asian impact. Thus, the Vörs-Papkert-B data rather imply a continuous population between the 8-11th centuries with mild influences from eastern immigrants.

The 6 ConqC graveyards with meaningful sample size showed a rather similar overall picture with an average of 12.6% east Eurasian Hgs almost confined to C and D, inferring a similar overall east Eurasian impact throughout cis-Danubia. The presence of not negligible proportion of east Eurasian Hgs in the ConqC population is a clear sign of admixture with eastern immigrants, presumably with Avars and/or Conquerors. This effect distinguishes ConqC from contemporary European populations as well as from modern Hungarians in which east Eurasian Hgs are neglectable.

Though the 10-11^th^ century genetic makeup of the Carpathian Basin certainly showed more local differences than in later periods, what is also reflected in Figure 2, but our result indicate signs of genetic consolidation. Next to comparable mitogenome composition of distant cemeteries, we identified five pairs of potential relatives in various combinations between commoner cemeteries (Table S3b), a potential sign of exogamy.

The overall Hg composition of the commoner population proved to be significantly dissimilar from the elite in respect to both east and West Eurasian Hgs, indicating that these two groups had highly different origin. Population genetic analysis definitely clustered ConqC primarily with European and Near Eastern populations, well separating them from the elite, suggesting that people with local European origin dominated the ConqC population. On the other hand, ConqC had smallest SHD distance from ConqE (Table S5b) suggesting that in spite of their dissimilarity, out of the studied ancient populations ConqE shared highest proportion of identical terminal Hgs with ConqC, which can be best explained by admixture between these groups. It is especially telling that the most frequent ConqE Hgs N1a1a1a1, its derivative N1a1a1a1a and T1a1 were present in numerous commoner cemeteries (Table S3c). The east Eurasian N1a1a1a1 ConqE marker most likely originated from the Afanasievo or Sintashta-Tagar cultures, while despite its general Eurasian range, Mongolian Chemurchek-Uyuk-Deer stone-khirigsuur [26] origin of T1a1 in ConqE is very plausible. SHD data (Table S5b) imply that the Steppe_EMBA affinity of ConqC, observed on Figure. 4, was probably also a consequence of ConqE admixture.

As the SHD value perfectly represent the common gene pool, the SHD distance of 0.85 indicates 14% common gene pool between ConqE and ConqC. Of course, this value cannot be directly interpreted as headcount proportion of immigrants and local people, as the studied ConqE population very likely also contains a yet unidentified admixture fraction from ConqC, furthermore both could have acquired common elements from other unknown populations. Reciprocal gene flow from ConqC to ConqE is indicated by their shared Hgs; H7, K1c1, T2b and V7a (Table S3c) which are absent from east Eurasia but had been present in the Carpathian Basin from the Neolithic-Bronze Age. On the whole above data imply that the proportion of immigrant Conquerors did not exceed 14% of the local population, but the more realistic value must be far smaller.

The contemporary local population descended from previous cultures of the Carpathian Basin, and it has indeed been shown that large number of people survived to the 10^th^ century from the previous Avar period [27,28]. The Avars also brought along a package of east Eurasian Hgs, and a significant fraction of these in ConqC, not shared with ConqE, are potential candidates of Avar heritage.

For more accurate conclusions future investigations are necessary, including high-resolution genome analysis of commoner and elite cemeteries, furthermore genome data from the pre-Avar, Avar, and later Árpádian populations would provide a more complete picture about the exact contribution of subsequent nomadic migrations to the demographic history of the Carpathian Basin.

## Supplementary Materials

Appendix A: Brief archeological background of the Hungarian commoner cemeteries.Figure S1: MJ Networs, Table S1: Summary and archaeological details of studied samples, Table S2a: Shotgun sequence data, Table S2b: Mitogenome sequence data, Table S3a: SNP positions of the mitogenomes, Table S3b: List of identical haplotypes. Table S3c: List of ConqC and ConqE individuals and shared haplogroups Table S4: Ancient mitogenome database, Table S5a: Pairwise Fst matrix of ancient populations. Table S5b: SHD distance matrix of ancient populations.

## Author Contributions

Conceptualization, T.T. and E.N.; data curation, K.M., Z.M., E.Ny. and E.N.; formal analysis, K.M., G.I.B.V., Z.M. and E.N.; funding acquisition, T.T. and E.N.; investigation, K.M., B.K., D.L., O.S., B.T., G.I.B.V., I.N. and E.N.; methodology, T.T., Z.M. and E.N.; project administration, K.M. and E.N.; resources, B.T., Zs.B., A.M., Zs.G., S.V., L.K. and Gy.P.; software, Z.M.; supervision, T.T. and E.N.; visualization, K.M. and E.N.; writing – original draft preparation, K.M. and E.N.; writing – review & editing: Z.M., I.R., T.K., G.I.B.V and T.T. All authors have read and agreed to the published version of the manuscript.

## Supporting information

Tablse S1

## Acknowledgments

We are grateful to our archaeologist colleagues, and especially László Kovács, Eszter Istvánovits, Gábor Lőrinczy, Ibolya M. Nepper and József Szentpéteri for their help. We are also grateful to Miklós Kásler and Gábor Horváth-Lugossy for their encouragement and Szabolcs Tóth for his administrative work.

