## Supplementary material for "Maternal lineages from 10-11^th^ century commoner cemeteries of the Carpathian Basin": Figure S1

### S1 Figure: Median-Joining Networks (1-98) for mtDNA sequences of the 202 studied samples

Phylogenetic trees (1–98), made with Median-Joining Network, from mtDNA sequences of the 202 archaic samples.

Phylogenetic trees are arranged in alphabetic order according to haplogroups. The 154 sub-haplogroups are depicted on 98 Networks. Samples falling into the same sub-haplogroup with the studied sample are encircled. The smallest colored circles represent one individual; circle sizes are proportional to the number of individuals with identical sequences. (When large number of sequences with few phylogenetically informative SNP-s are aligned, the algorithm may force the most similar but not identical sequences into the same large circle.) Green circles identify studied  samples, red circles represent modern samples, and violet circles correspond to ancient samples.

Number of crosslines between neighboring circles denotes mutation distances. Length of connecting lines is irrelevant, as they were modified in order to fit page. Genebank accesion number and origin of samples closest to the studied conquerors are listed next to the circles.

We summarized the probable origin of the samples’ Hg lineage in colored framed text.

Eastern Eurasia

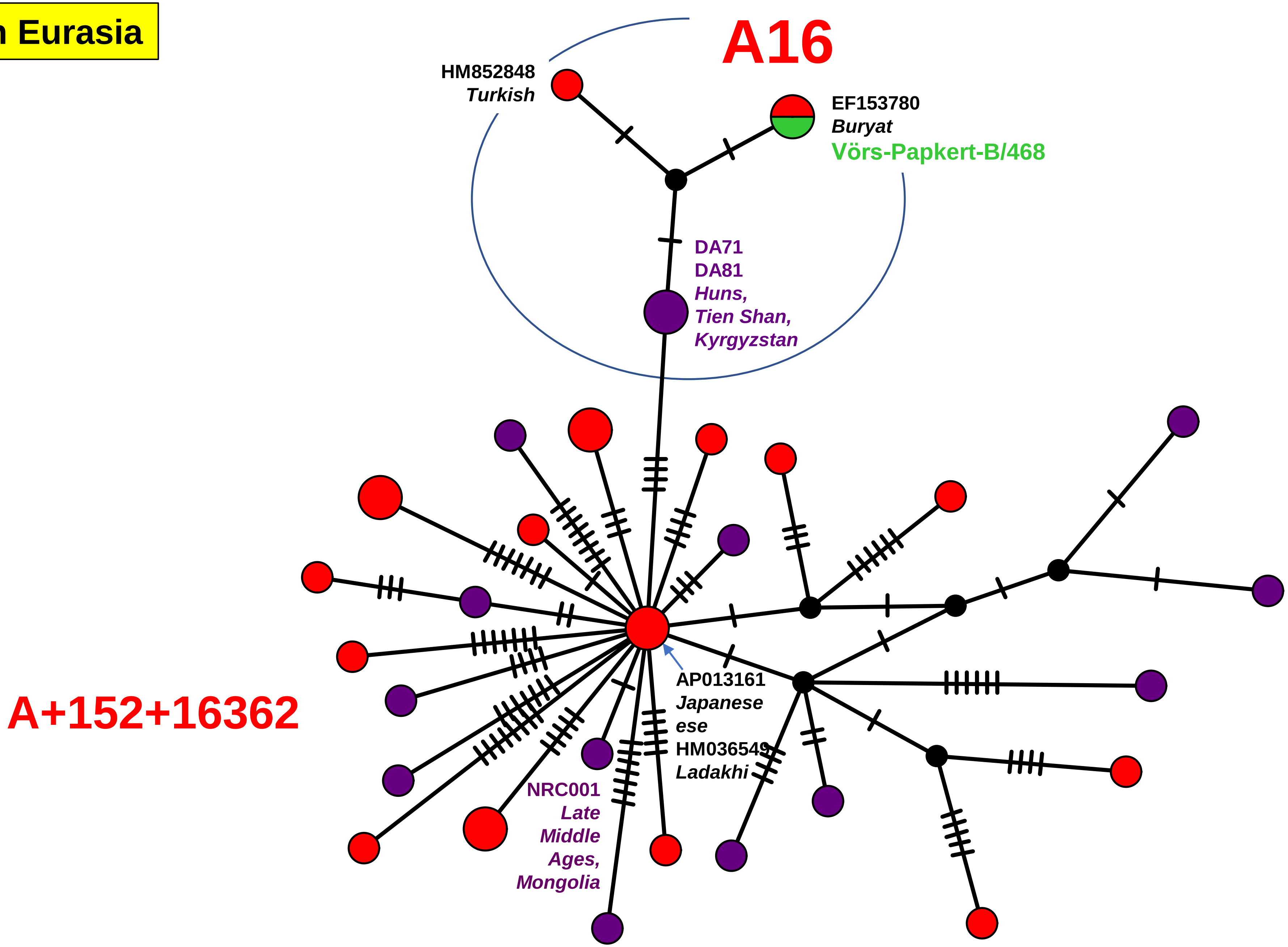


Network 1

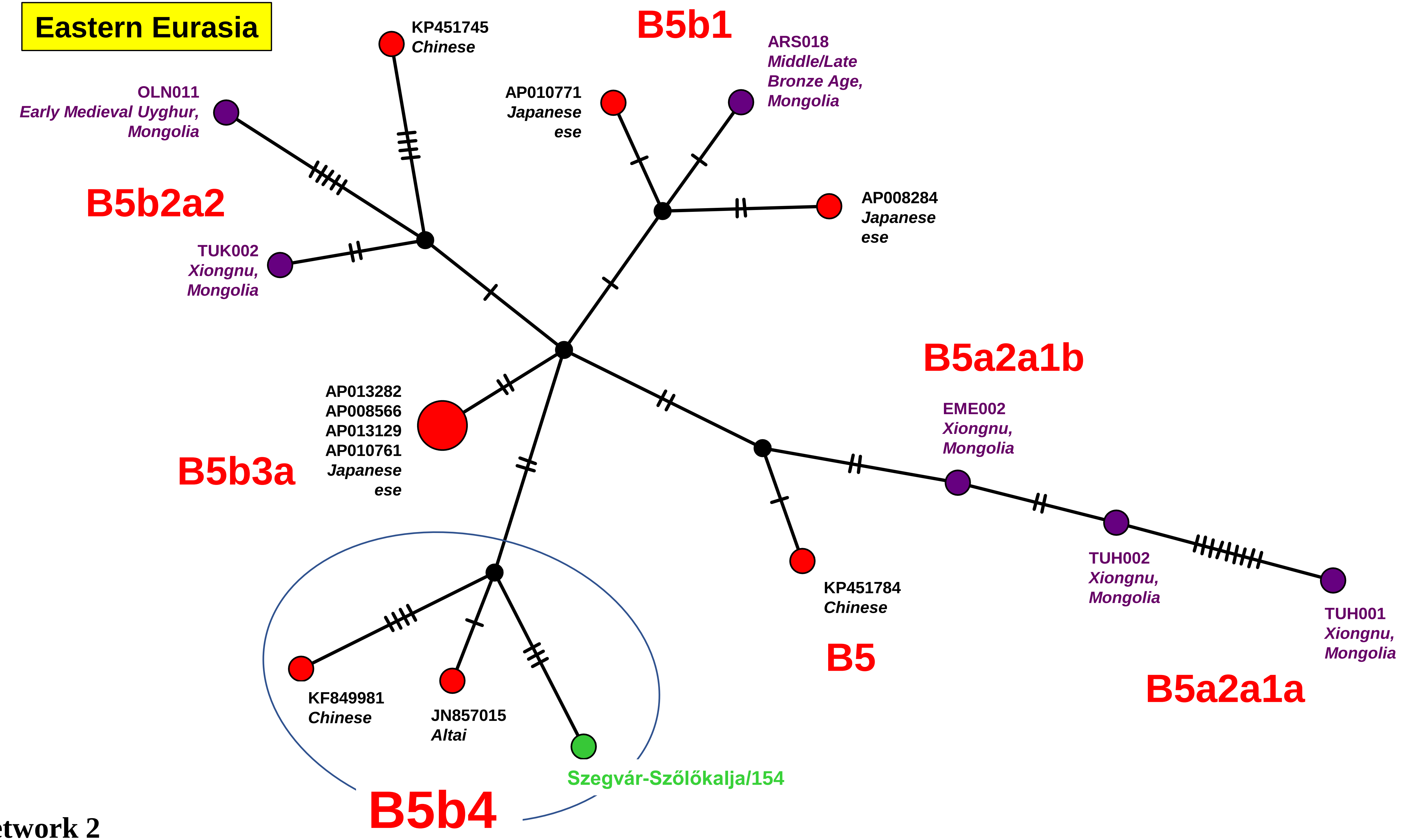
Eastern Eurasia
KP451745
Chinese
B5b1
ARS018
Middle/Late
Bronze Age,
Mongolia
OLN011
Early Medieval Uyghur,
Mongolia
AP010771
Japanese
ese
B5b2a2
AP008284
Japanese
ese
TUK002
Xiongnu,
Mongolia
B5a2a1b
AP013282
AP008566
AP013129
AP010761
Japanese
ese
EME002
Xiongnu,
Mongolia
B5b3a
TUH002
Xiongnu,
Mongolia
KP451784
Chinese
TUH001
Xiongnu,
Mongolia
B5
B5a2a1a
KF849981
Chinese
JN857015
Altai
Szegvár-Szőlőkalja/154
B5b4

Network 2

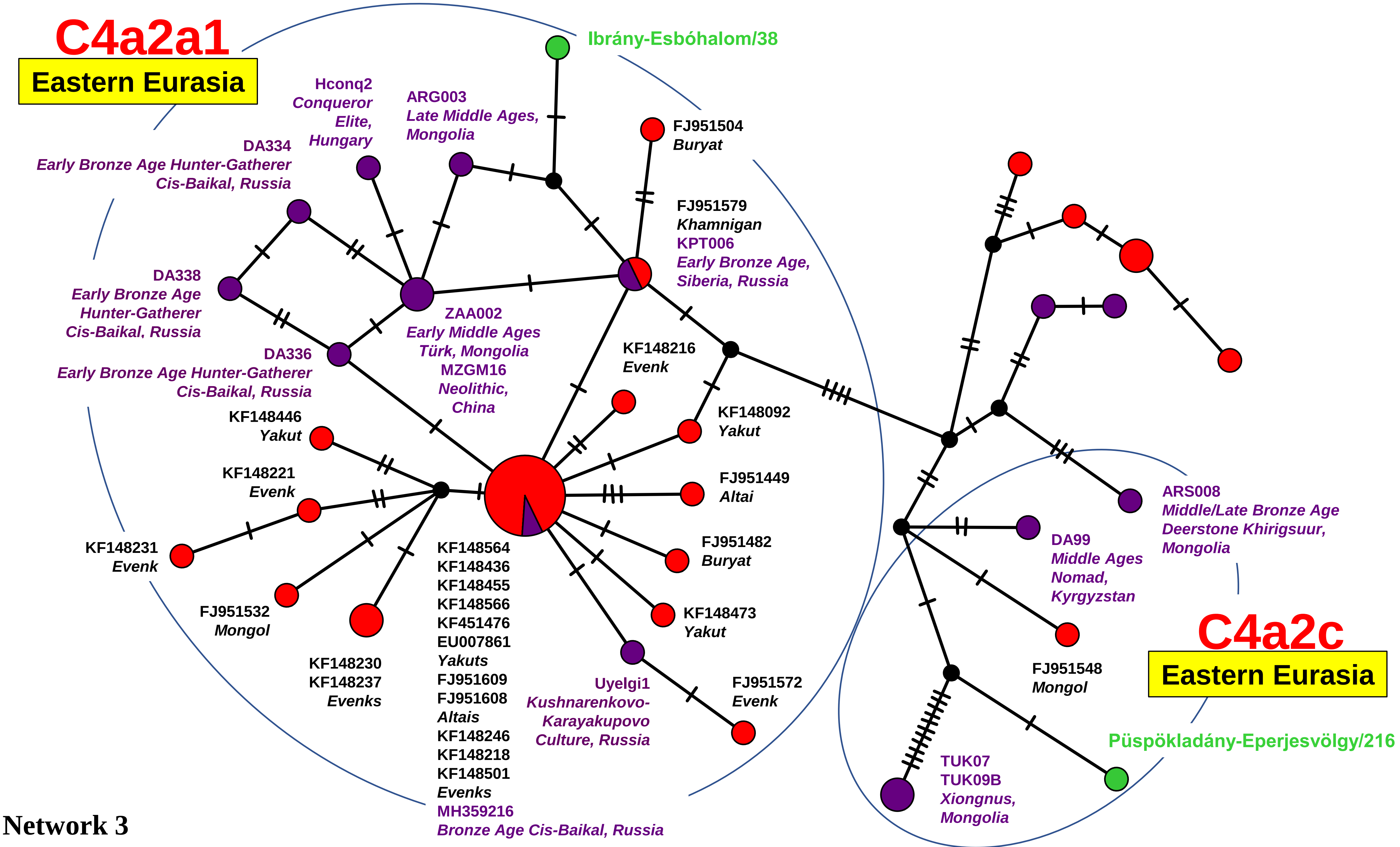

C4a2a1
Eastern Eurasia
Ibrány-Esbóhalom/38
Hconq2
Conqueror
Elite,
Hungary
ARG003
Late Middle Ages,
Mongolia
FJ951504
Buryat
DA334
Early Bronze Age Hunter-Gatherer
Cis-Baikal, Russia
FJ951579
Khamnigan
KPT006
Early Bronze Age,
Siberia, Russia
DA338
Early Bronze Age
Hunter-Gatherer
Cis-Baikal, Russia
ZAA002
Early Middle Ages
Türk, Mongolia
MZGM16
Neolithic,
China
DA336
Early Bronze Age Hunter-Gatherer
Cis-Baikal, Russia
KF148216
Evenk
KF148446
Yakut
KF148092
Yakut
KF148221
Evenk
FJ951449
Altai
KF148231
Evenk
FJ951482
Buryat
FJ951532
Mongol
KF148564
KF148436
KF148455
KF148566
KF451476
EU007861
Yakuts
FJ951609
FJ951608
Altais
KF148246
KF148218
KF148501
Evenks
MH359216
Bronze Age Cis-Baikal, Russia
KF148230
KF148237
Evenks
KF148473
Yakut
Uyelgi1
Kushnarenkovo-
Karayakupovo
Culture, Russia
FJ951572
Evenk
ARS008
Middle/Late Bronze Age
Deerstone Khirigsuur,
Mongolia
DA99
Middle Ages
Nomad,
Kyrgyzstan
C4a2c
Eastern Eurasia
FJ951548
Mongol
Püspökladány-Eperjesvölgy/216
TUK07
TUK09B
Xiongnus,
Mongolia


Network 3

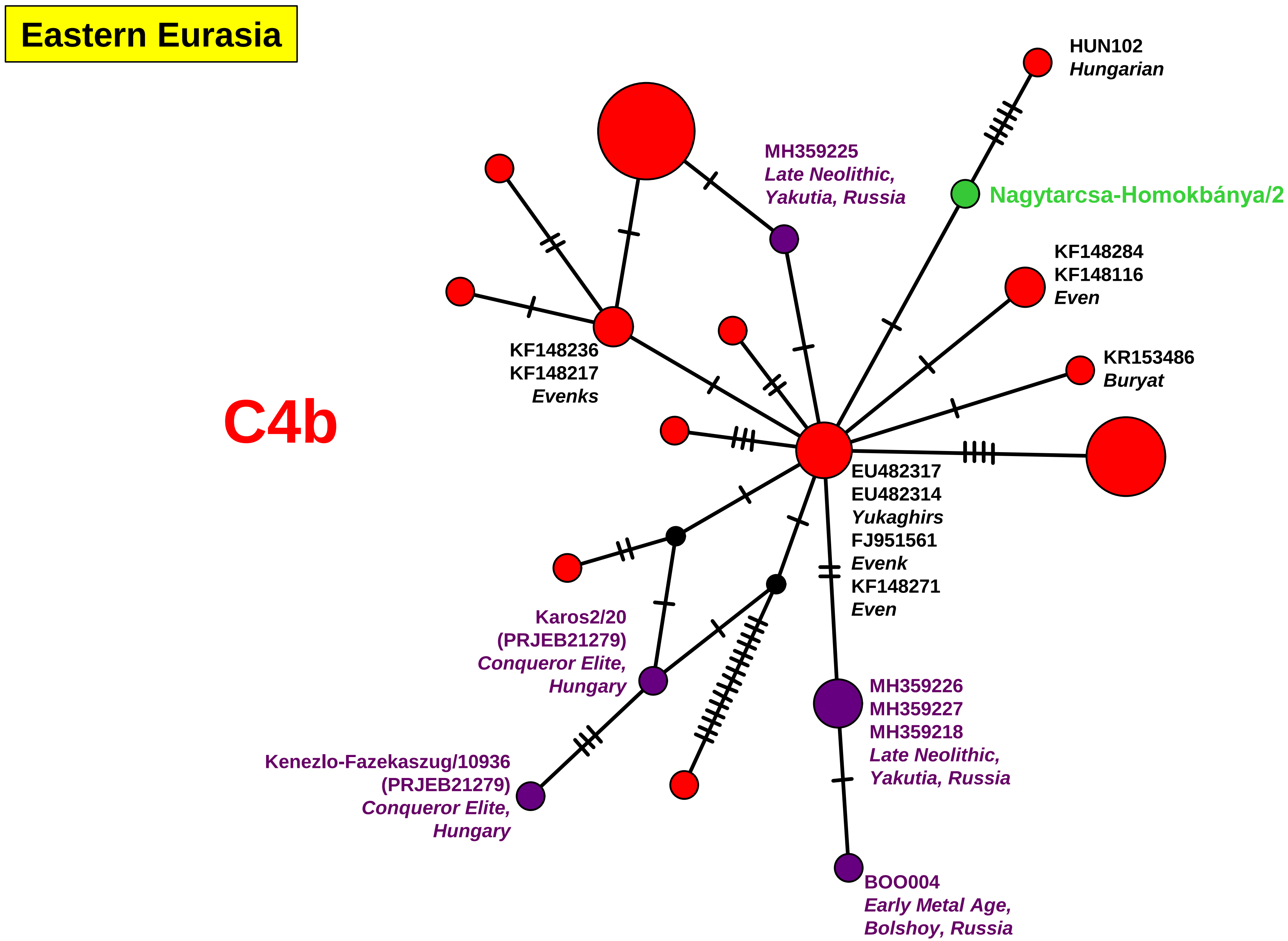
Eastern Eurasia
C4b
HUN102
Hungarian
MH359225
Late Neolithic,
Yakutia, Russia
Nagytarcsa-Homokbánya/2
KF148284
KF148116
Even
KR153486
Buryat
KF148236
KF148217
Evenks
EU482317
EU482314
Yukaghirs
FJ951561
Evenk
KF148271
Even
Karos2/20
(PRJEB21279)
Conqueror Elite,
Hungary
MH359226
MH359227
MH359218
Late Neolithic,
Yakutia, Russia
Kenezlo-Fazekaszug/10936
(PRJEB21279)
Conqueror Elite,
Hungary
BOO004
Early Metal Age,
Bolshoy, Russia

Network 4

Eastern Eurasia

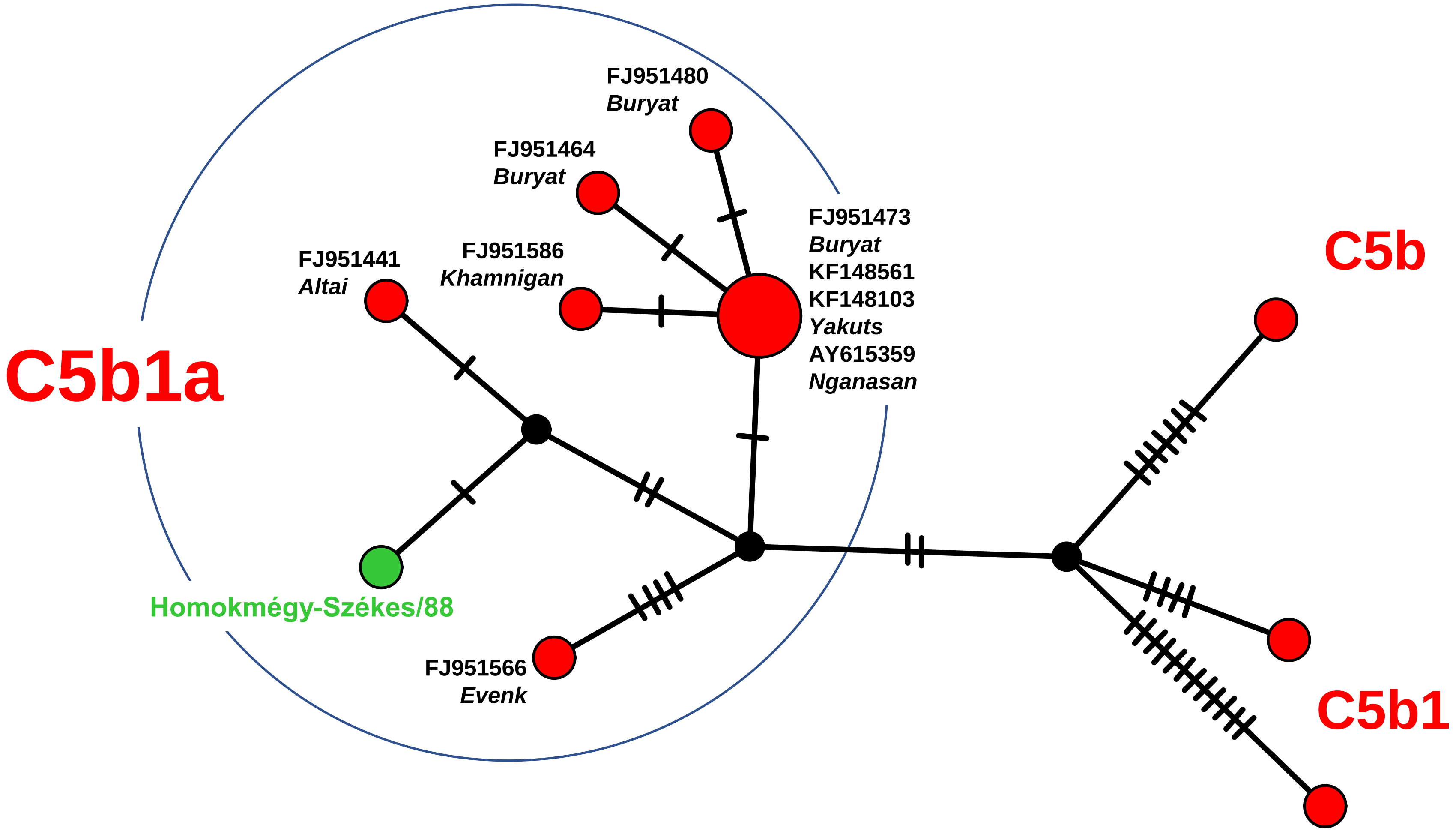


Network 5

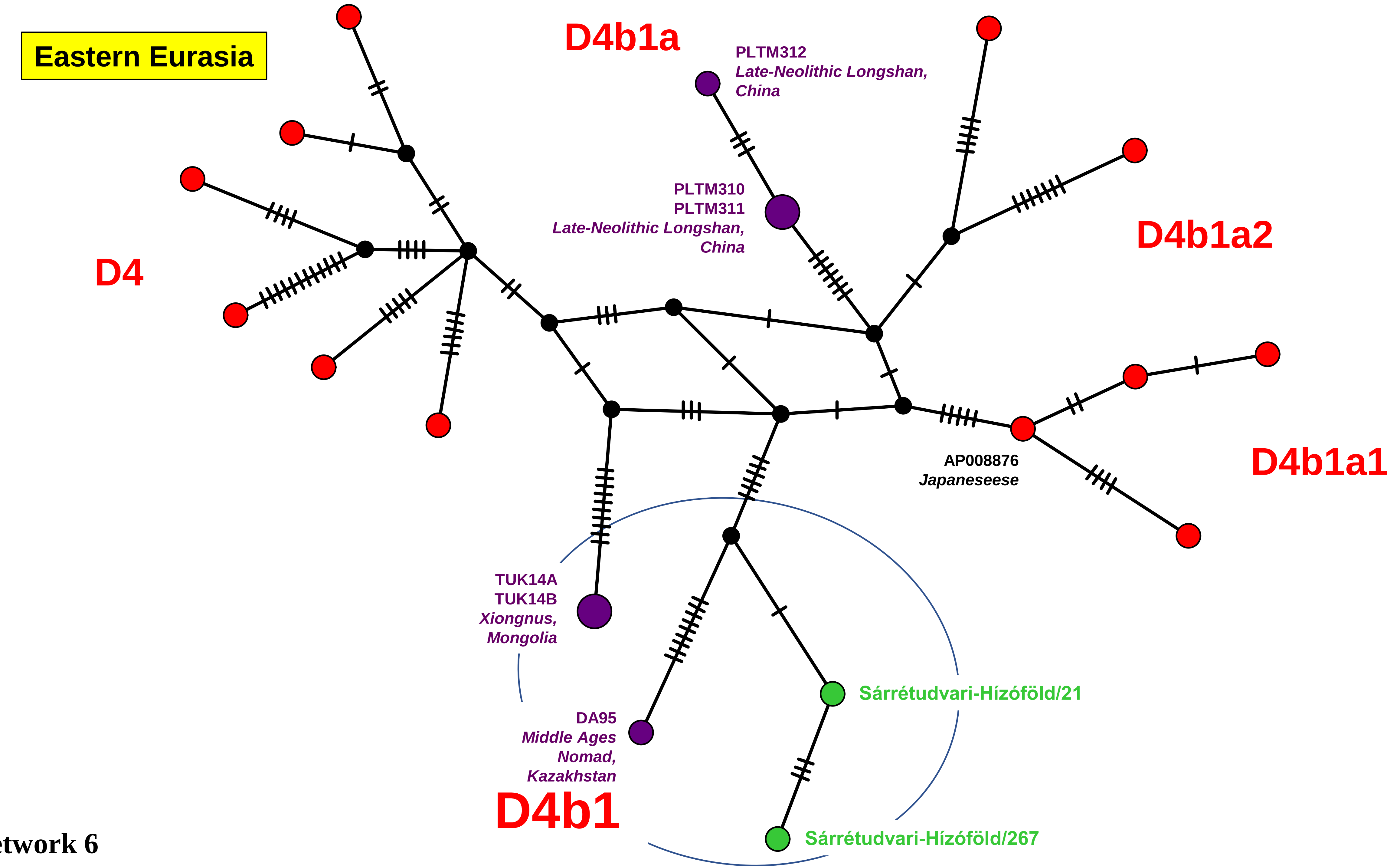

Eastern Eurasia
D4b1a
PLTM312
Late-Neolithic Longshan, China
PLTM310
PLTM311
Late-Neolithic Longshan, China
D4b1a2
D4
D4b1a1
AP008876
Japaneseese
TUK14A
TUK14B
Xiongnus, Mongolia
Sárrétudvari-Hízóföld/21
DA95
Middle Ages Nomad, Kazakhstan
D4b1
Sárrétudvari-Hízóföld/267


Network 6

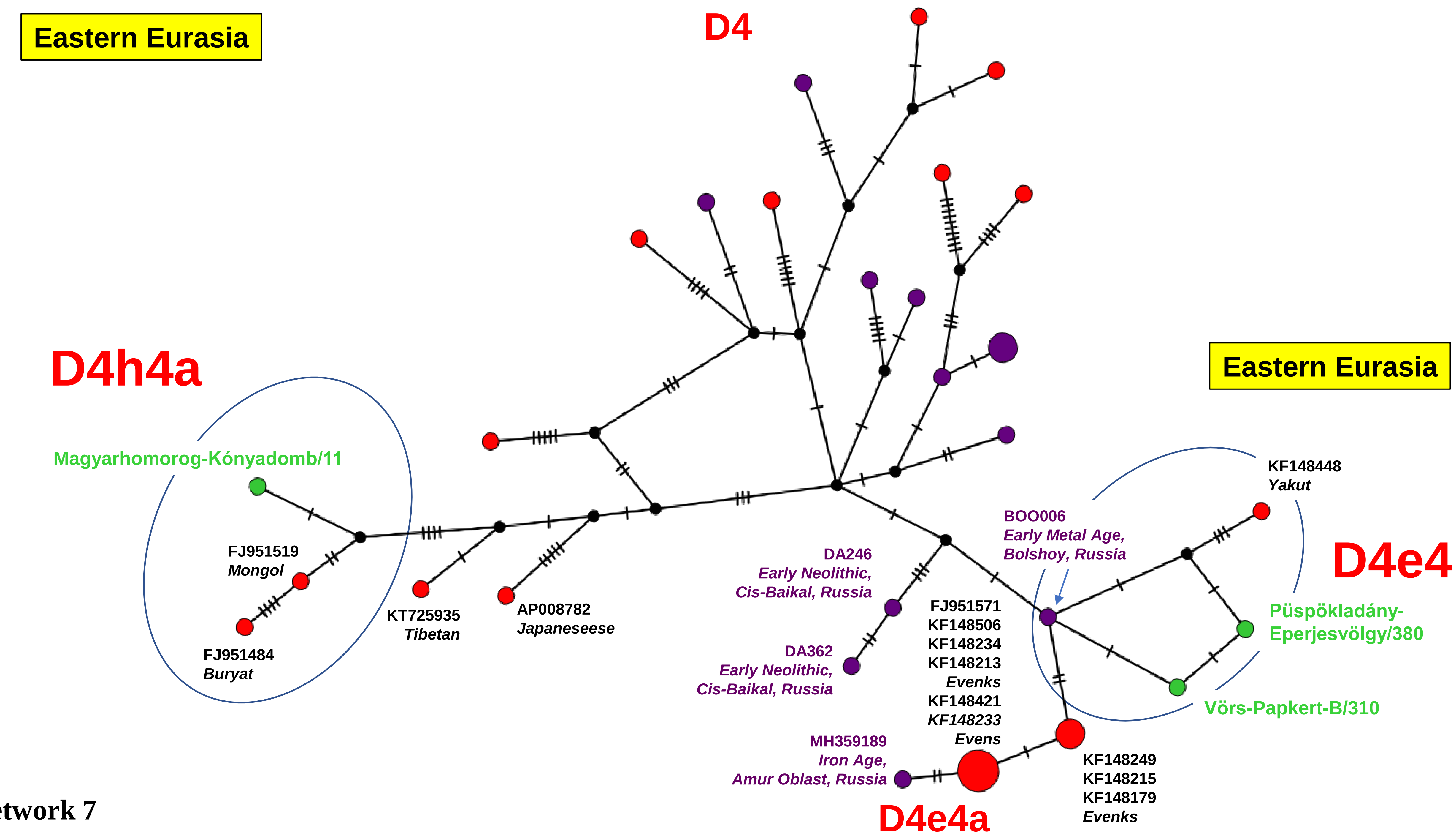
Eastern Eurasia
D4
D4h4a
Magyarhomorog-Kónyadomb/11
FJ951519
Mongol
FJ951484
Buryat
KT725935
Tibetan
AP008782
Japaneseese
Eastern Eurasia
KF148448
Yakut
BOO006
Early Metal Age,
Bolshoy, Russia
D4e4
DA246
Early Neolithic,
Cis-Baikal, Russia
Püspökladány-
Eperjesvölgy/380
FJ951571
KF148506
KF148234
KF148213
Evenks
KF148421
KF148233
Evens
DA362
Early Neolithic,
Cis-Baikal, Russia
Vörs-Papkert-B/310
MH359189
Iron Age,
Amur Oblast, Russia
KF148249
KF148215
KF148179
Evenks
D4e4a

Network 7

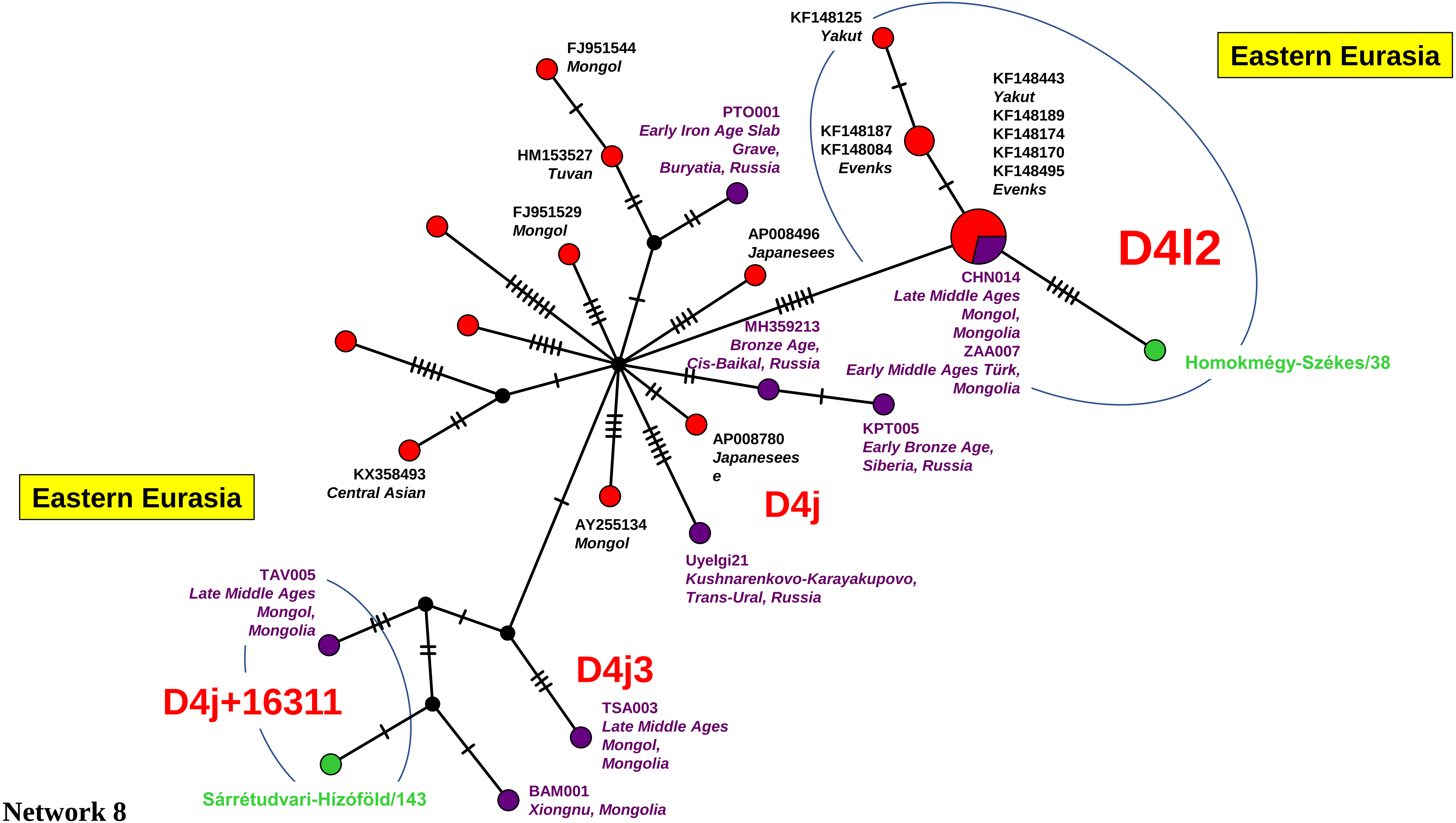
Eastern Eurasia
Eastern Eurasia
KF148125
Yakut
FJ951544
Mongol
KF148443
Yakut
KF148189
KF148174
KF148170
KF148495
Evenks
PTO001
Early Iron Age Slab
Grave,
Buryatia, Russia
KF148187
KF148084
Evenks
HM153527
Tuvan
FJ951529
Mongol
AP008496
Japanesees
D4l2
CHN014
Late Middle Ages
Mongol,
Mongolia
ZAA007
Early Middle Ages Türk,
Mongolia
MH359213
Bronze Age,
Cis-Baikal, Russia
Homokmégy-Székes/38
KPT005
Early Bronze Age,
Siberia, Russia
AP008780
Japanesees
e
KX358493
Central Asian
D4j
AY255134
Mongol
Uyelgi21
Kushnarenkovo-Karayakupovo,
Trans-Ural, Russia
TAV005
Late Middle Ages
Mongol,
Mongolia
D4j3
D4j+16311
TSA003
Late Middle Ages
Mongol,
Mongolia
BAM001
Xiongnu, Mongolia
Sárrétudvari-Hízóföld/143

Network 8

Eastern Eurasia

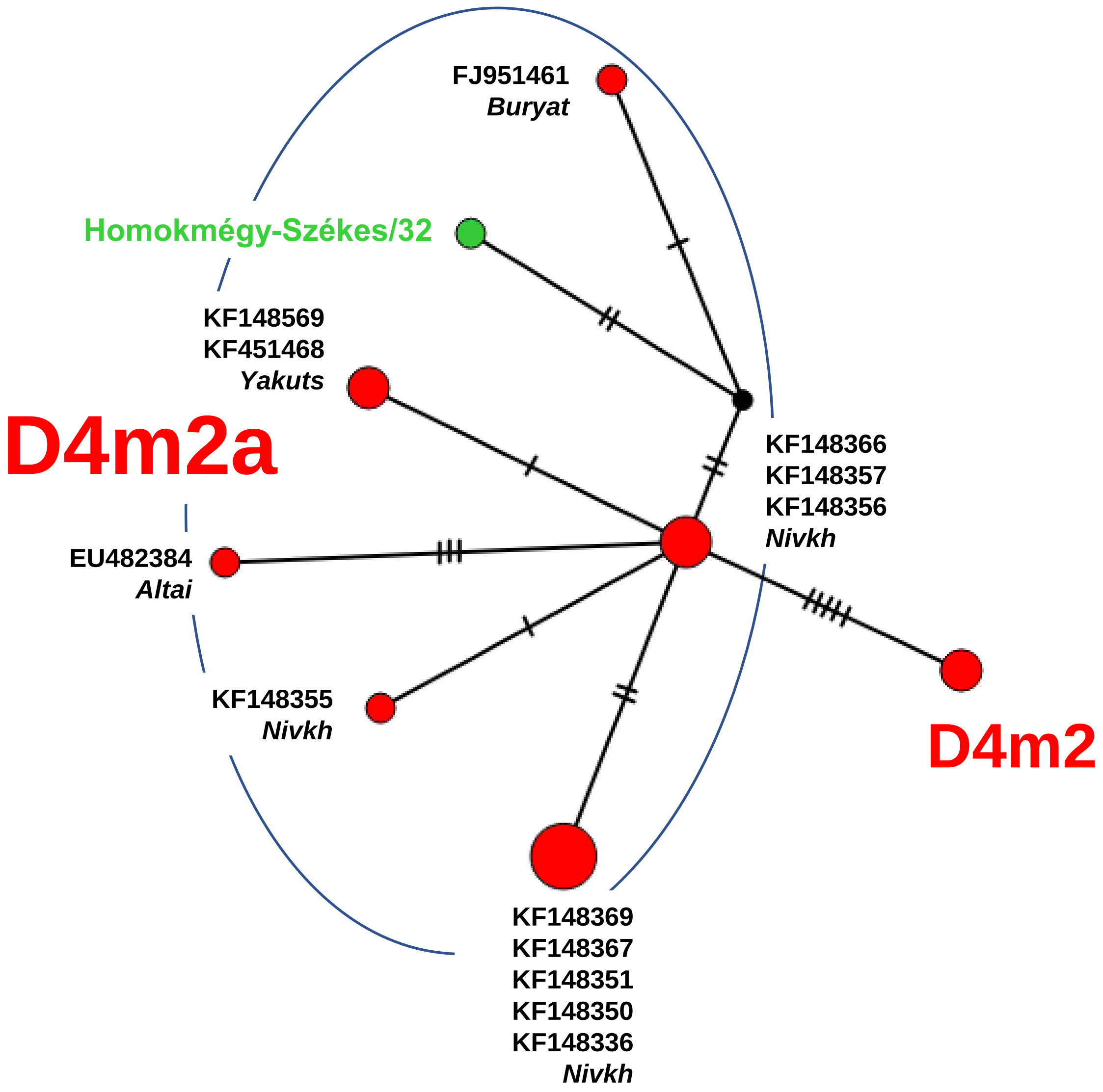


Network 9

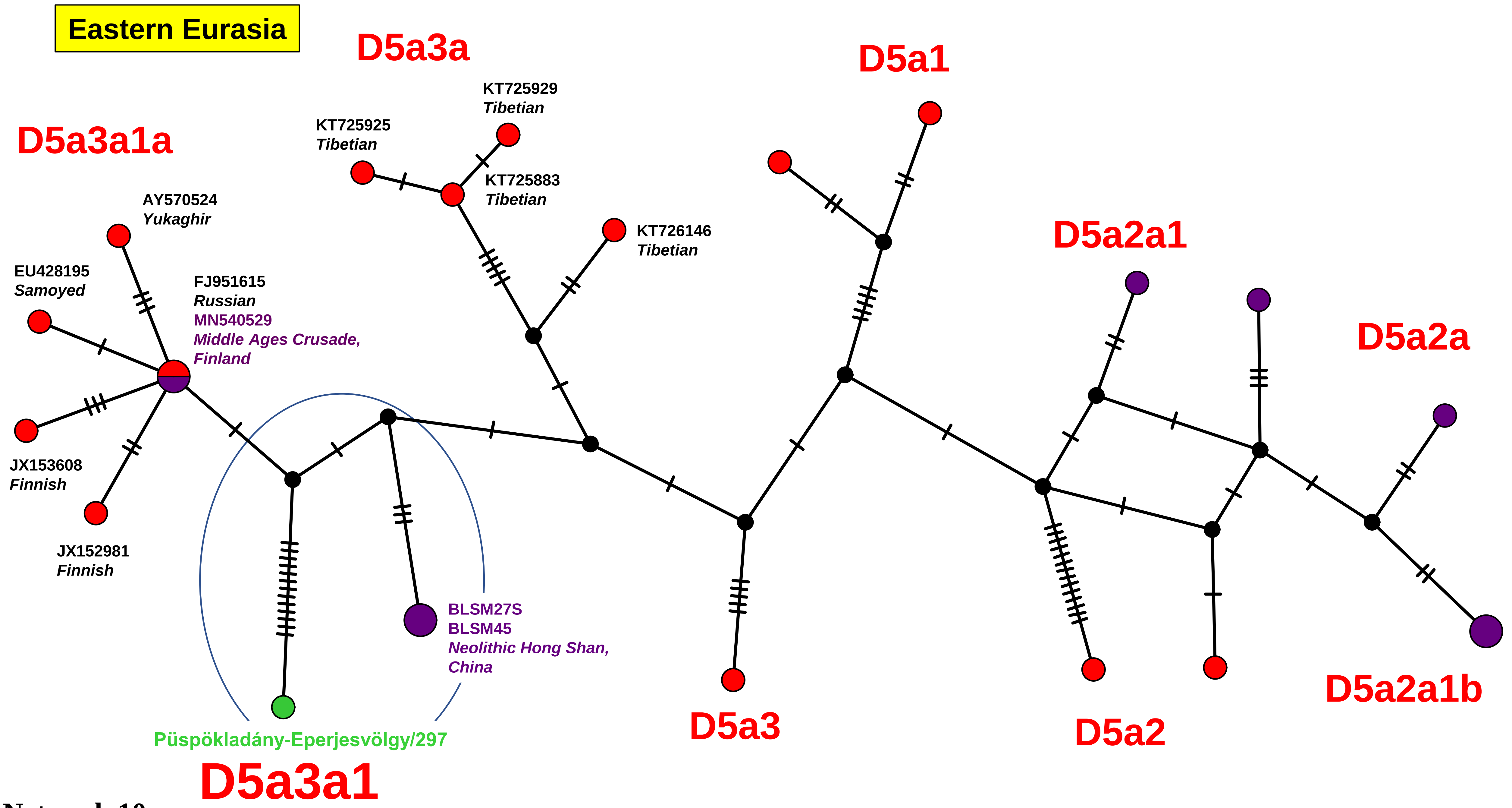
Eastern Eurasia
D5a3a
D5a1
D5a3a1a
KT725929
Tibetian
KT725925
Tibetian
KT725883
Tibetian
KT726146
Tibetian
AY570524
Yukaghir
D5a2a1
EU428195
Samoyed
FJ951615
Russian
MN540529
Middle Ages Crusade,
Finland
D5a2a
JX153608
Finnish
JX152981
Finnish
BLSM27S
BLSM45
Neolithic Hong Shan,
China
D5a2a1b
D5a3
D5a2
Püspökladány-Eperjesvölgy/297
D5a3a1

Network 10

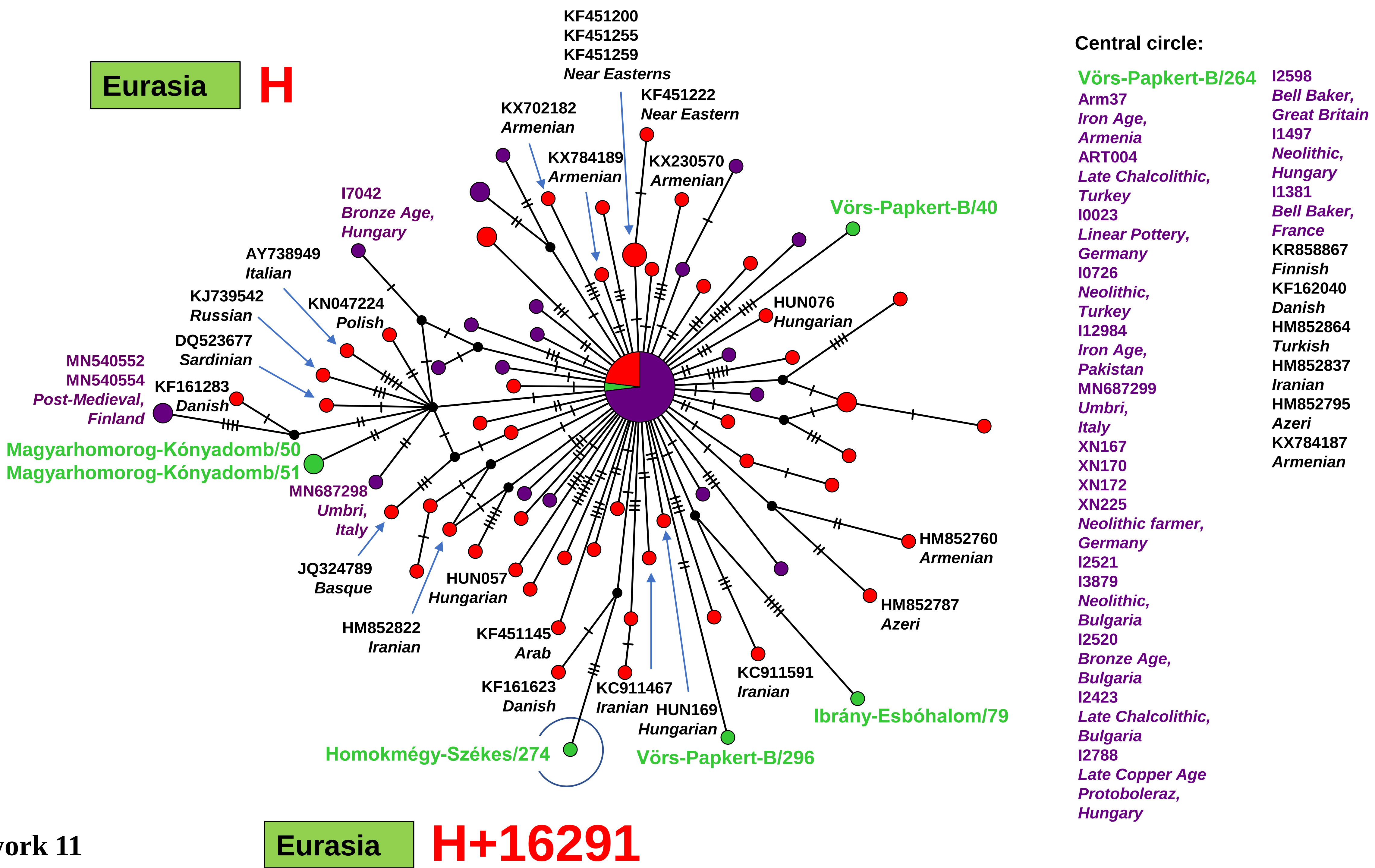
Eurasia
H
KF451200
KF451255
KF451259
Near Easterns
KF451222
Near Eastern
KX702182
Armenian
KX784189
Armenian
KX230570
Armenian
I7042
Bronze Age,
Hungary
Vörs-Papkert-B/40
AY738949
Italian
KJ739542
Russian
KN047224
Polish
HUN076
Hungarian
DQ523677
Sardinian
MN540552
MN540554
Post-Medieval,
Finland
KF161283
Danish
Magyarhomorog-Kónyadomb/50
Magyarhomorog-Kónyadomb/51
MN687298
Umbri,
Italy
JQ324789
Basque
HUN057
Hungarian
HM852760
Armenian
HM852787
Azeri
HM852822
Iranian
KF451145
Arab
KF161623
Danish
KC911467
Iranian
HUN169
Hungarian
KC911591
Iranian
Ibrány-Esbóhalom/79
Homokmégy-Székes/274
Vörs-Papkert-B/296
Central circle:
Vörs-Papkert-B/264
Arm37
Iron Age,
Armenia
ART004
Late Chalcolithic,
Turkey
I0023
Linear Pottery,
Germany
I0726
Neolithic,
Turkey
I12984
Iron Age,
Pakistan
MN687299
Umbri,
Italy
XN167
XN170
XN172
XN225
Neolithic farmer,
Germany
I2521
I3879
Neolithic,
Bulgaria
I2520
Bronze Age,
Bulgaria
I2423
Late Chalcolithic,
Bulgaria
I2788
Late Copper Age
Protoboleraz,
Hungary
I2598
Bell Baker,
Great Britain
I1497
Neolithic,
Hungary
I1381
Bell Baker,
France
KR858867
Finnish
KF162040
Danish
HM852864
Turkish
HM852837
Iranian
HM852795
Azeri
KX784187
Armenian
Eurasia
H+16291

Network 11

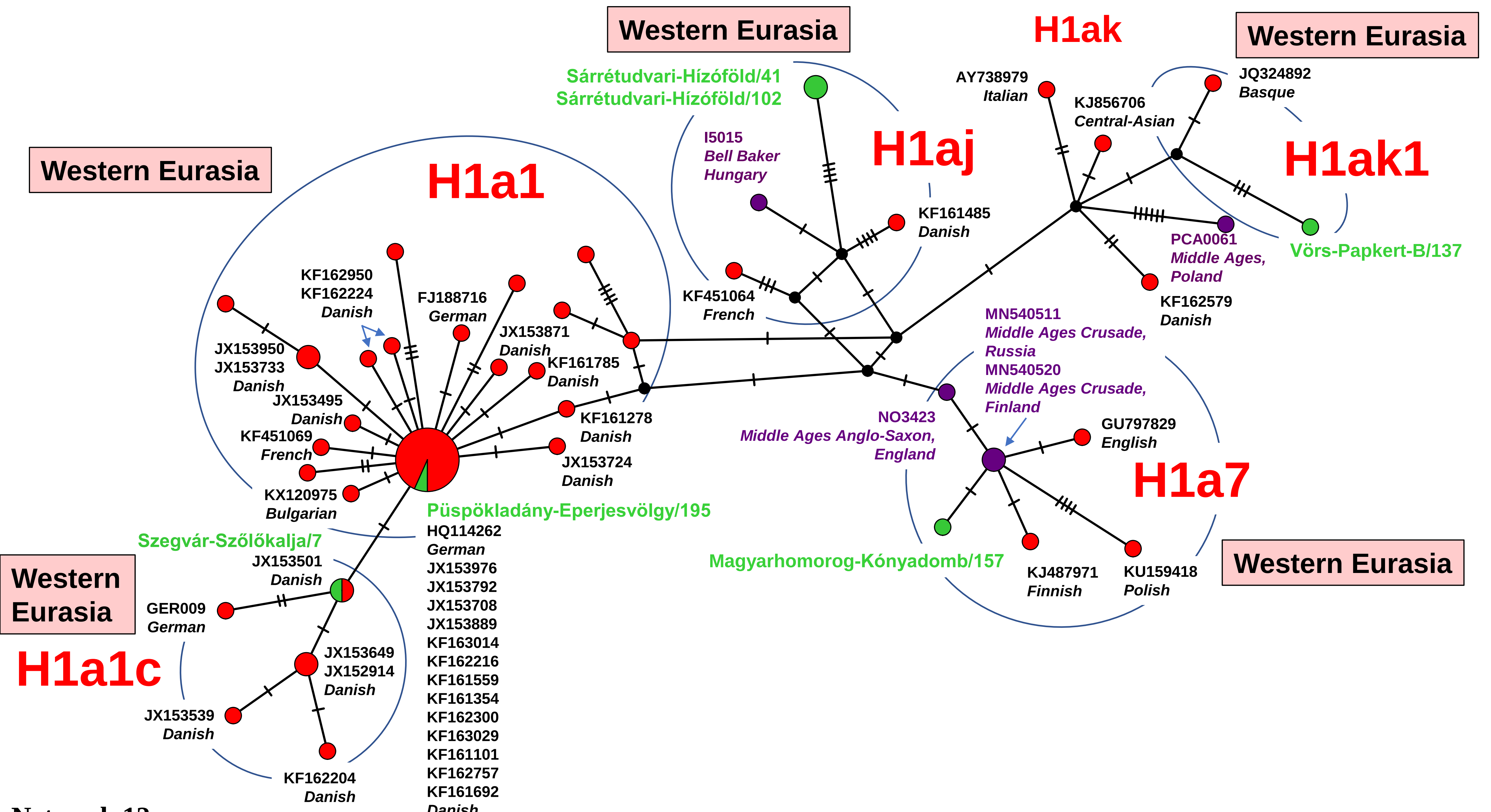
Western Eurasia
H1ak
Western Eurasia
Sárrétudvari-Hízóföld/41
Sárrétudvari-Hízóföld/102
AY738979
Italian
JQ324892
Basque
KJ856706
Central-Asian
I5015
Bell Baker
Hungary
H1aj
H1ak1
Western Eurasia
H1a1
KF161485
Danish
PCA0061
Middle Ages,
Poland
Vörs-Papkert-B/137
KF162950
KF162224
Danish
FJ188716
German
KF451064
French
KF162579
Danish
JX153871
Danish
MN540511
Middle Ages Crusade,
Russia
MN540520
Middle Ages Crusade,
Finland
JX153950
JX153733
Danish
KF161785
Danish
JX153495
Danish
KF161278
Danish
NO3423
Middle Ages Anglo-Saxon,
England
GU797829
English
KF451069
French
JX153724
Danish
H1a7
KX120975
Bulgarian
Püspökladány-Eperjesvölgy/195
HQ114262
German
JX153976
JX153792
JX153708
JX153889
KF163014
KF162216
KF161559
KF161354
KF162300
KF163029
KF161101
KF162757
KF161692
Danish
Szegvár-Szőlőkalja/7
JX153501
Danish
Western
Eurasia
GER009
German
Magyarhomorog-Kónyadomb/157
KJ487971
Finnish
KU159418
Polish
Western Eurasia
H1a1c
JX153649
JX152914
Danish
JX153539
Danish
KF162204
Danish
Network 12

Western Eurasia

# H1b

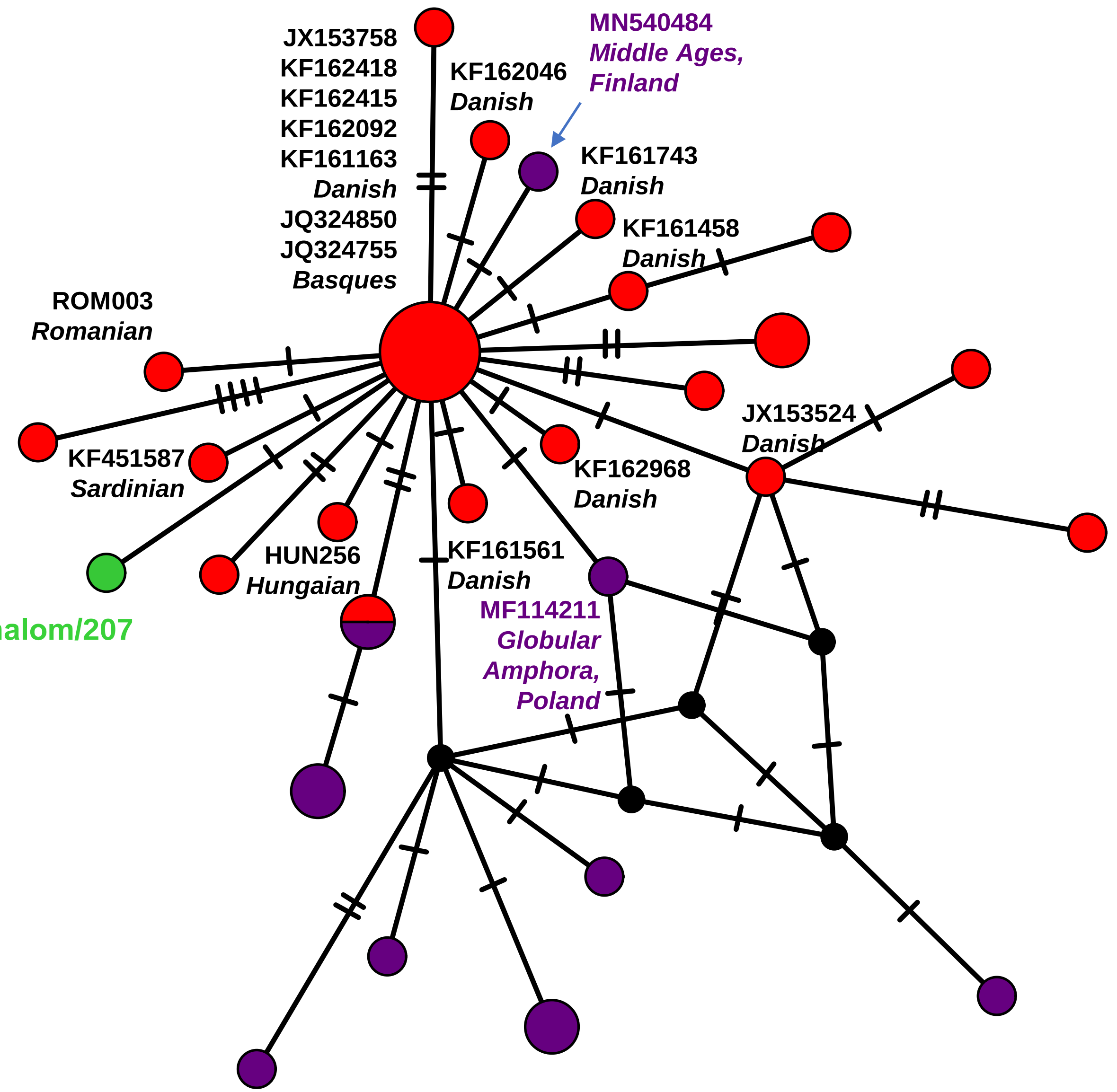


Network 13

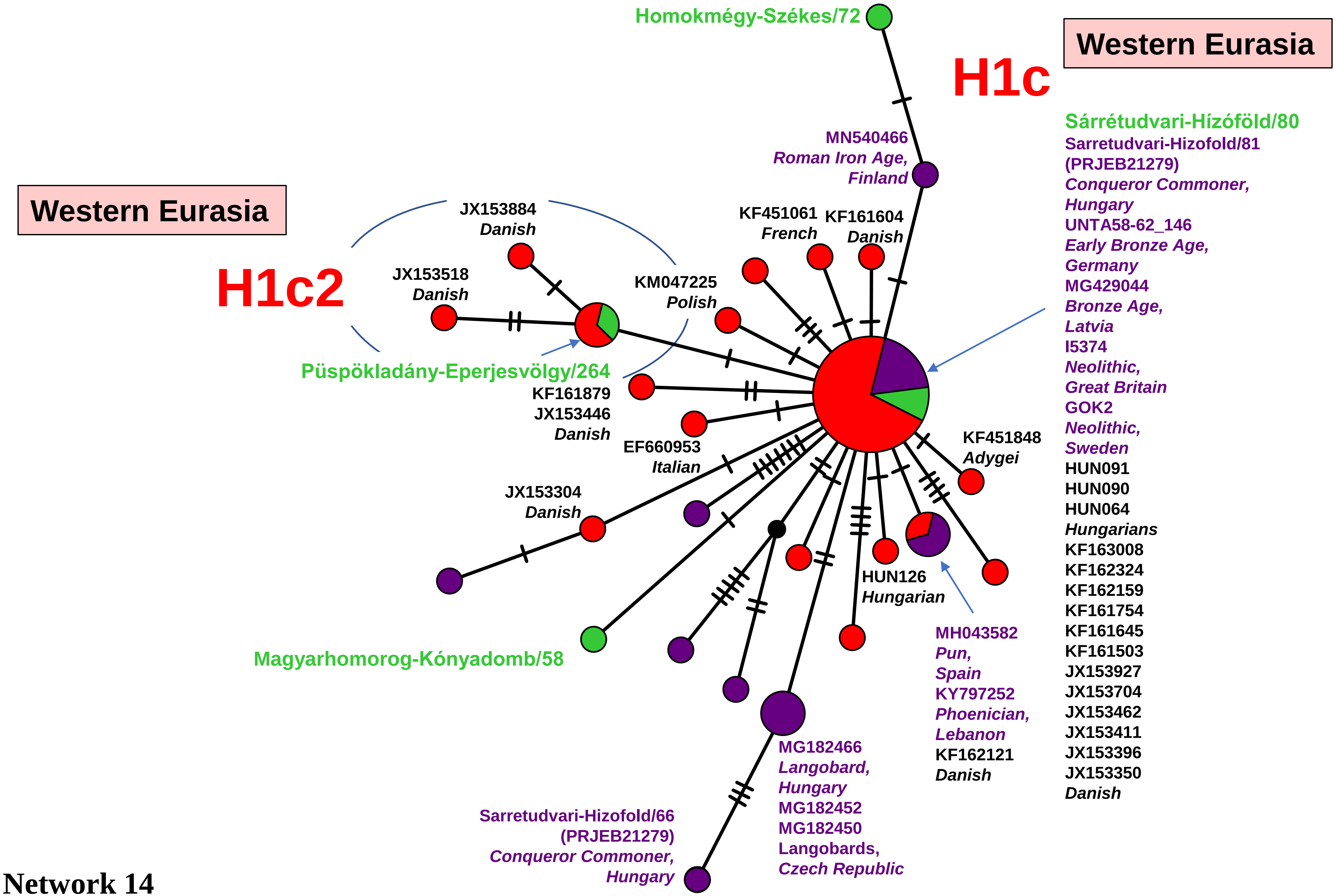
Western Eurasia
H1c
Homokmégy-Székes/72
Sárrétudvari-Hízóföld/80
Sarretudvari-Hizofold/81
(PRJEB21279)
Conqueror Commoner,
Hungary
UNTA58-62_146
Early Bronze Age,
Germany
MG429044
Bronze Age,
Latvia
I5374
Neolithic,
Great Britain
GOK2
Neolithic,
Sweden
HUN091
HUN090
HUN064
Hungarians
KF163008
KF162324
KF162159
KF161754
KF161645
KF161503
JX153927
JX153704
JX153462
JX153411
JX153396
JX153350
Danish
MN540466
Roman Iron Age,
Finland
Western Eurasia
H1c2
JX153884
Danish
JX153518
Danish
KM047225
Polish
KF451061
French
KF161604
Danish
Püspökladány-Eperjesvölgy/264
KF161879
JX153446
Danish
EF660953
Italian
KF451848
Adygei
JX153304
Danish
HUN126
Hungarian
MH043582
Pun,
Spain
KY797252
Phoenician,
Lebanon
KF162121
Danish
Magyarhomorog-Kónyadomb/58
MG182466
Langobard,
Hungary
MG182452
MG182450
Langobards,
Czech Republic
Sarretudvari-Hizofold/66
(PRJEB21279)
Conqueror Commoner,
Hungary

Network 14

**Western Eurasia**

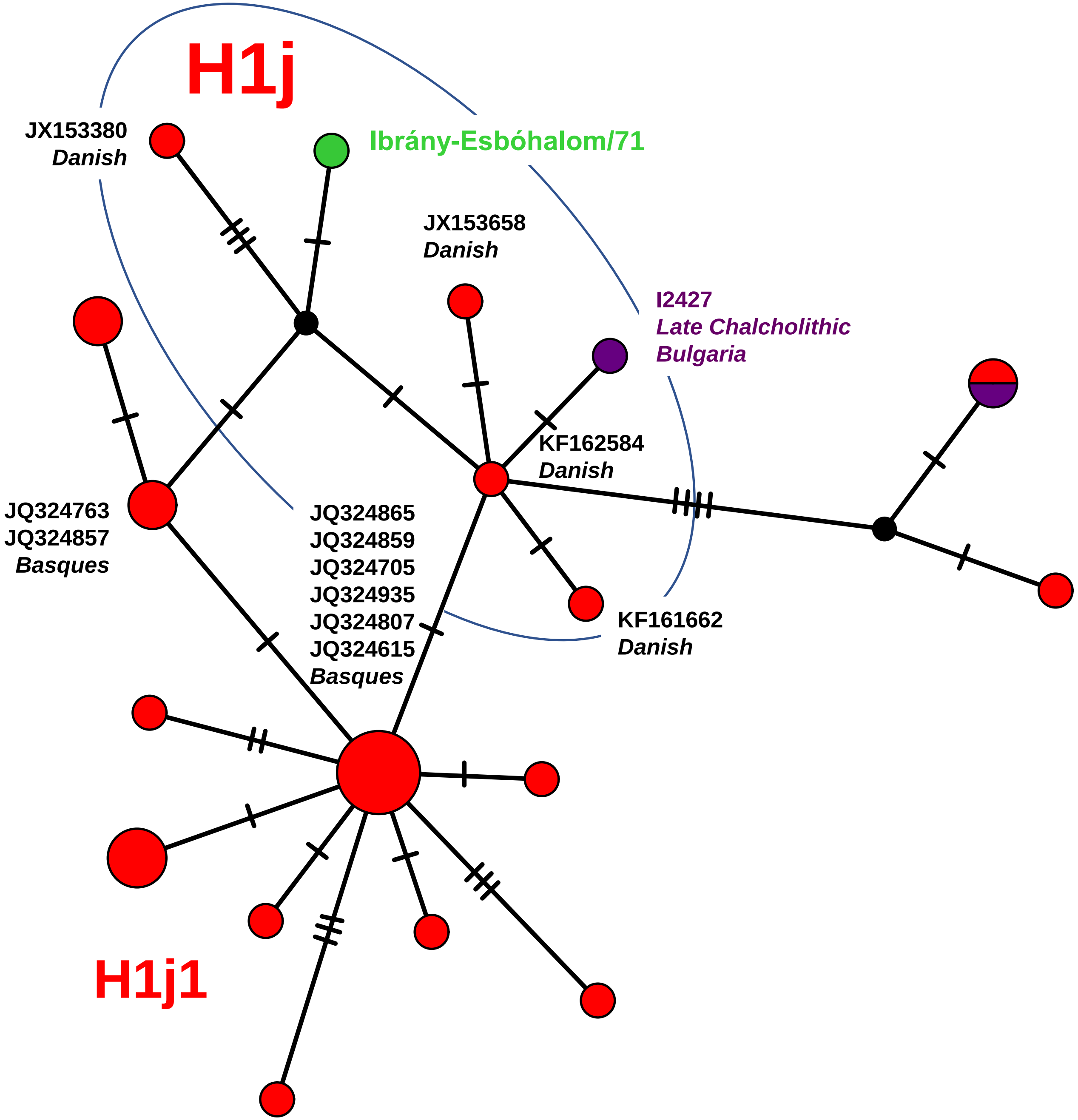


Network 15

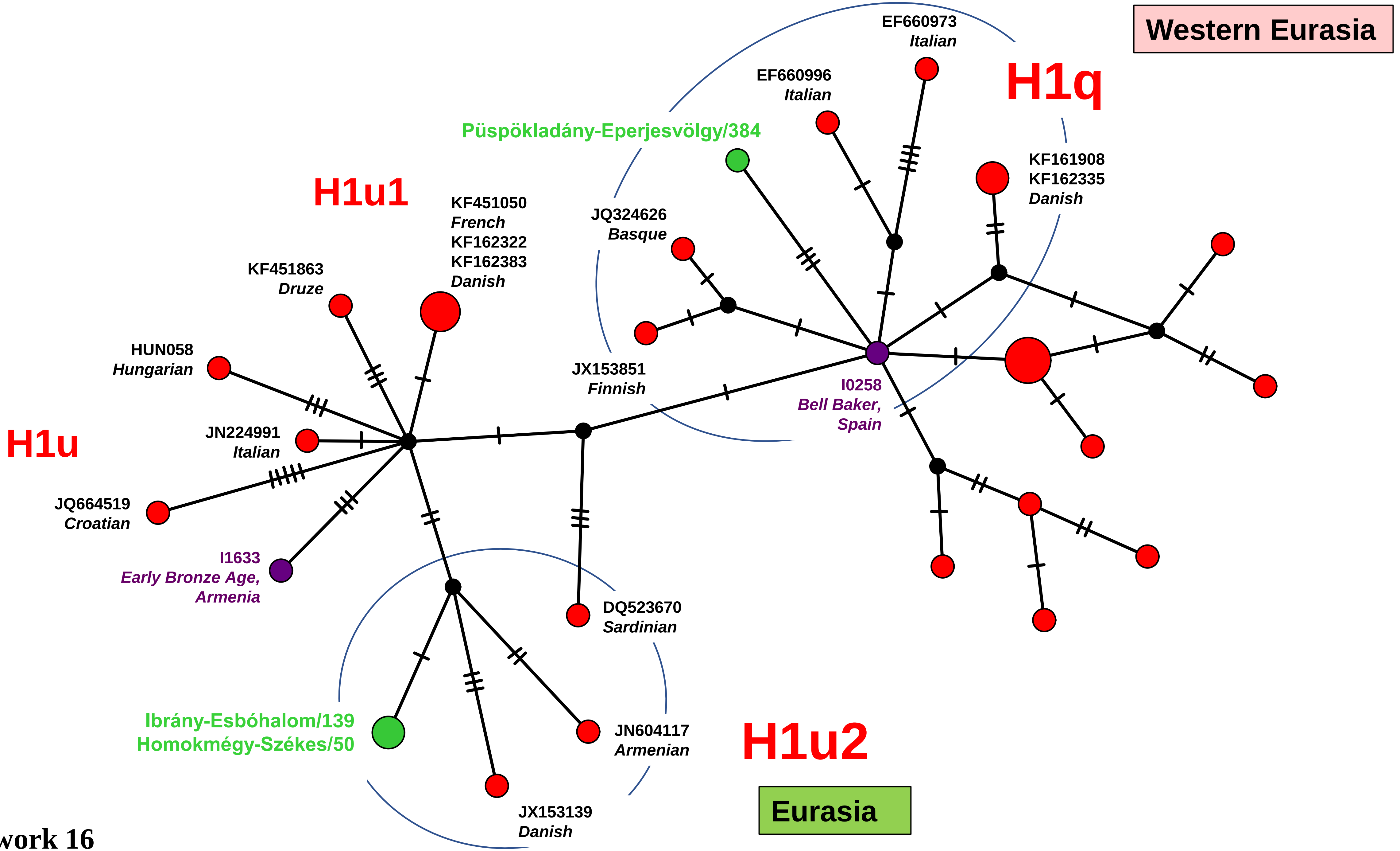
Western Eurasia
H1q
EF660973
Italian
EF660996
Italian
KF161908
KF162335
Danish
Püspökladány-Eperjesvölgy/384
H1u1
KF451050
French
KF162322
KF162383
Danish
JQ324626
Basque
KF451863
Druze
HUN058
Hungarian
JX153851
Finnish
I0258
Bell Baker,
Spain
H1u
JN224991
Italian
JQ664519
Croatian
I1633
Early Bronze Age,
Armenia
DQ523670
Sardinian
Ibrány-Esbóhalom/139
Homokmégy-Székes/50
JN604117
Armenian
H1u2
JX153139
Danish
Eurasia

Network 16

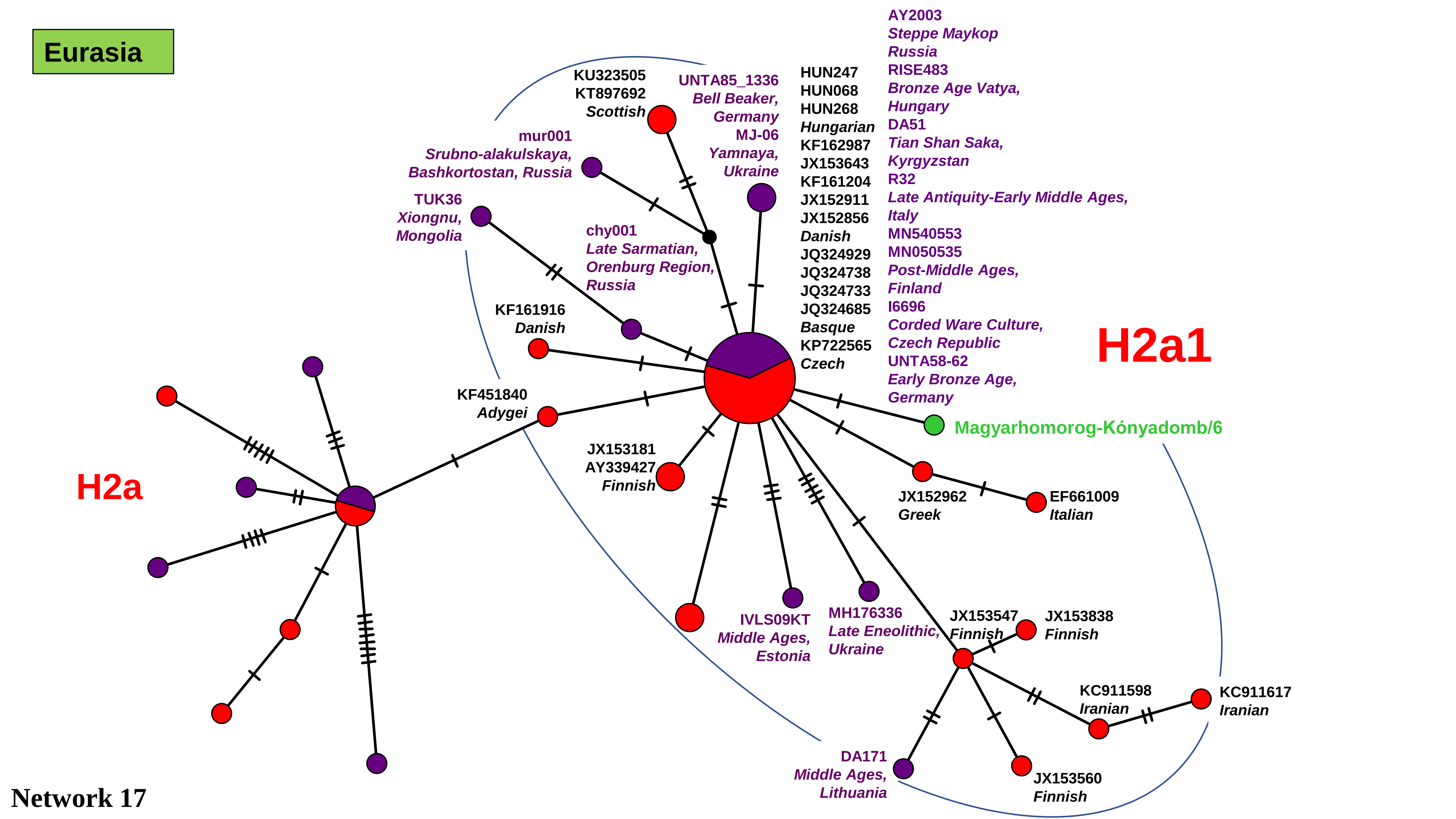
Eurasia
H2a
H2a1
KU323505
KT897692
Scottish
UNTA85_1336
Bell Beaker,
Germany
MJ-06
Yamnaya,
Ukraine
HUN247
HUN068
HUN268
Hungarian
KF162987
JX153643
KF161204
JX152911
JX152856
Danish
JQ324929
JQ324738
JQ324733
JQ324685
Basque
KP722565
Czech
AY2003
Steppe Maykop
Russia
RISE483
Bronze Age Vatya,
Hungary
DA51
Tian Shan Saka,
Kyrgyzstan
R32
Late Antiquity-Early Middle Ages,
Italy
MN540553
MN050535
Post-Middle Ages,
Finland
I6696
Corded Ware Culture,
Czech Republic
UNTA58-62
Early Bronze Age,
Germany
mur001
Srubno-alakulskaya,
Bashkortostan, Russia
TUK36
Xiongnu,
Mongolia
chy001
Late Sarmatian,
Orenburg Region,
Russia
KF161916
Danish
KF451840
Adygei
Magyarhomorog-Kónyadomb/6
JX153181
AY339427
Finnish
JX152962
Greek
EF661009
Italian
IVLS09KT
Middle Ages,
Estonia
MH176336
Late Eneolithic,
Ukraine
JX153547
Finnish
JX153838
Finnish
KC911598
Iranian
KC911617
Iranian
DA171
Middle Ages,
Lithuania
JX153560
Finnish

Network 17

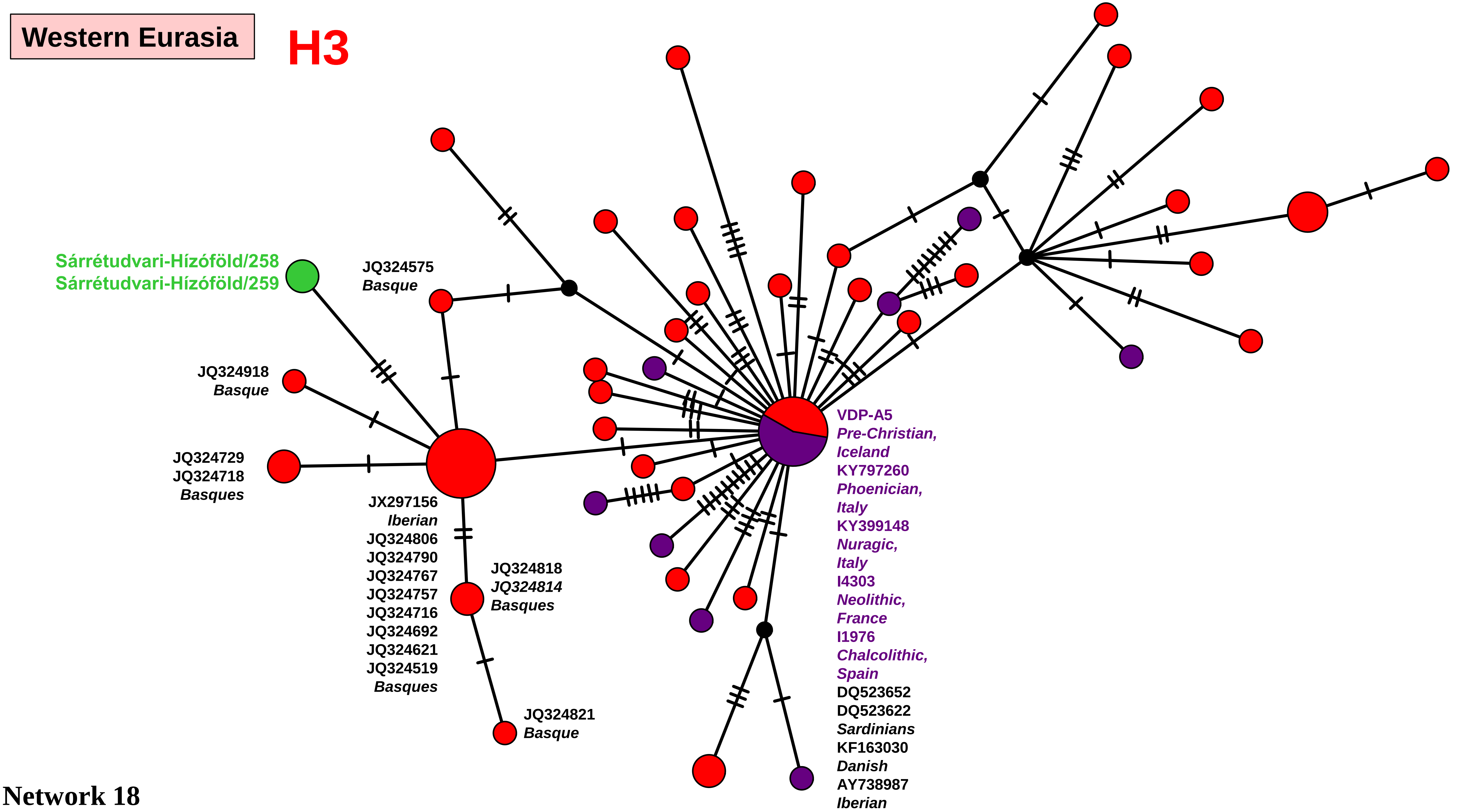

Western Eurasia
H3
Sárrétudvari-Hízóföld/258
Sárrétudvari-Hízóföld/259
JQ324575
Basque
JQ324918
Basque
JQ324729
JQ324718
Basques
JX297156
Iberian
JQ324806
JQ324790
JQ324767
JQ324757
JQ324716
JQ324692
JQ324621
JQ324519
Basques
JQ324818
JQ324814
Basques
JQ324821
Basque
VDP-A5
Pre-Christian,
Iceland
KY797260
Phoenician,
Italy
KY399148
Nuragic,
Italy
I4303
Neolithic,
France
I1976
Chalcolithic,
Spain
DQ523652
DQ523622
Sardinians
KF163030
Danish
AY738987
Iberian

Network 18

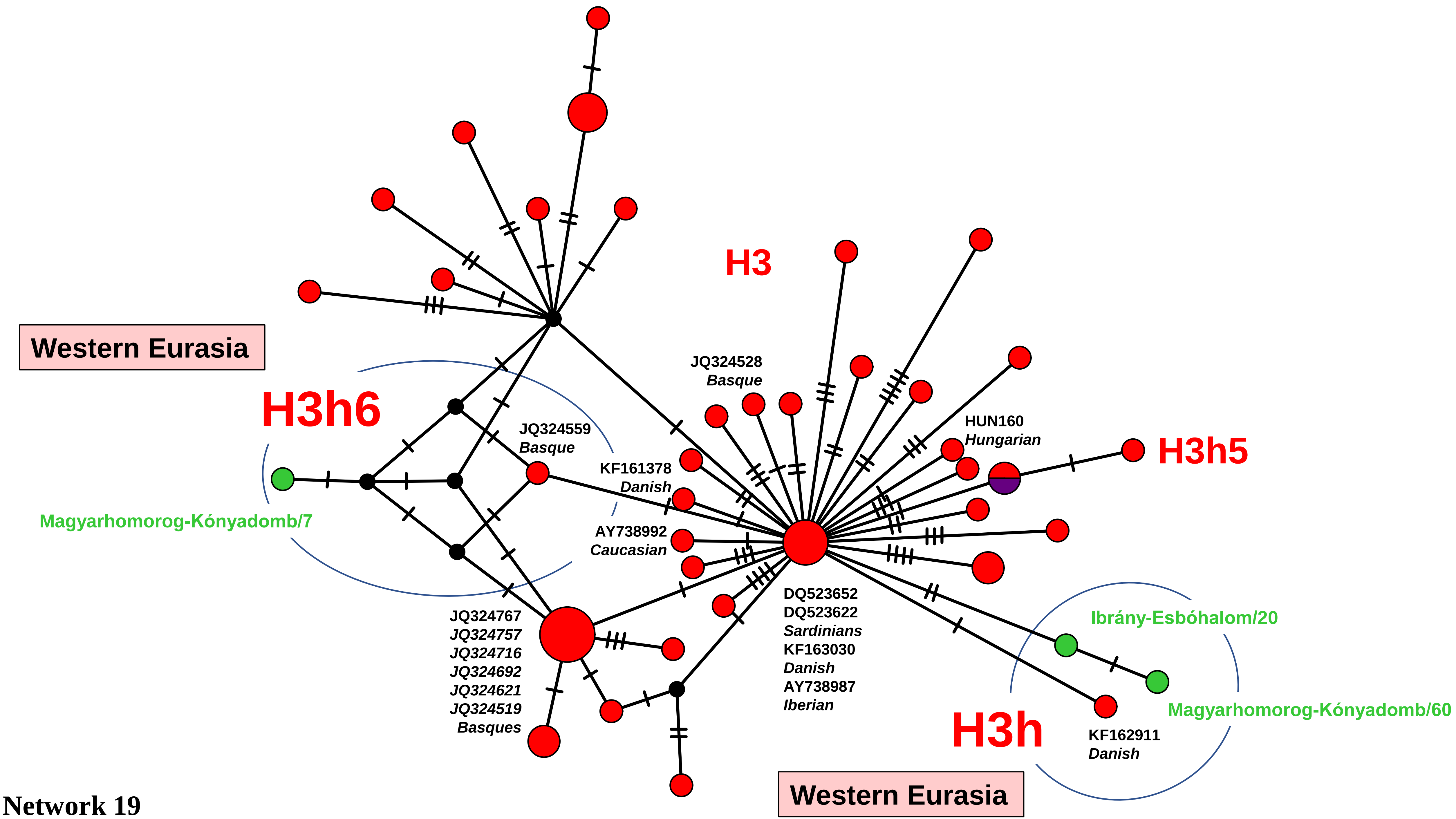
H3
Western Eurasia
H3h6
JQ324528
Basque
JQ324559
Basque
HUN160
Hungarian
H3h5
KF161378
Danish
Magyarhomorog-Kónyadomb/7
AY738992
Caucasian
DQ523652
DQ523622
Sardinians
KF163030
Danish
AY738987
Iberian
Ibrány-Esbóhalom/20
JQ324767
JQ324757
JQ324716
JQ324692
JQ324621
JQ324519
Basques
Magyarhomorog-Kónyadomb/60
H3h
KF162911
Danish
Western Eurasia

Network 19

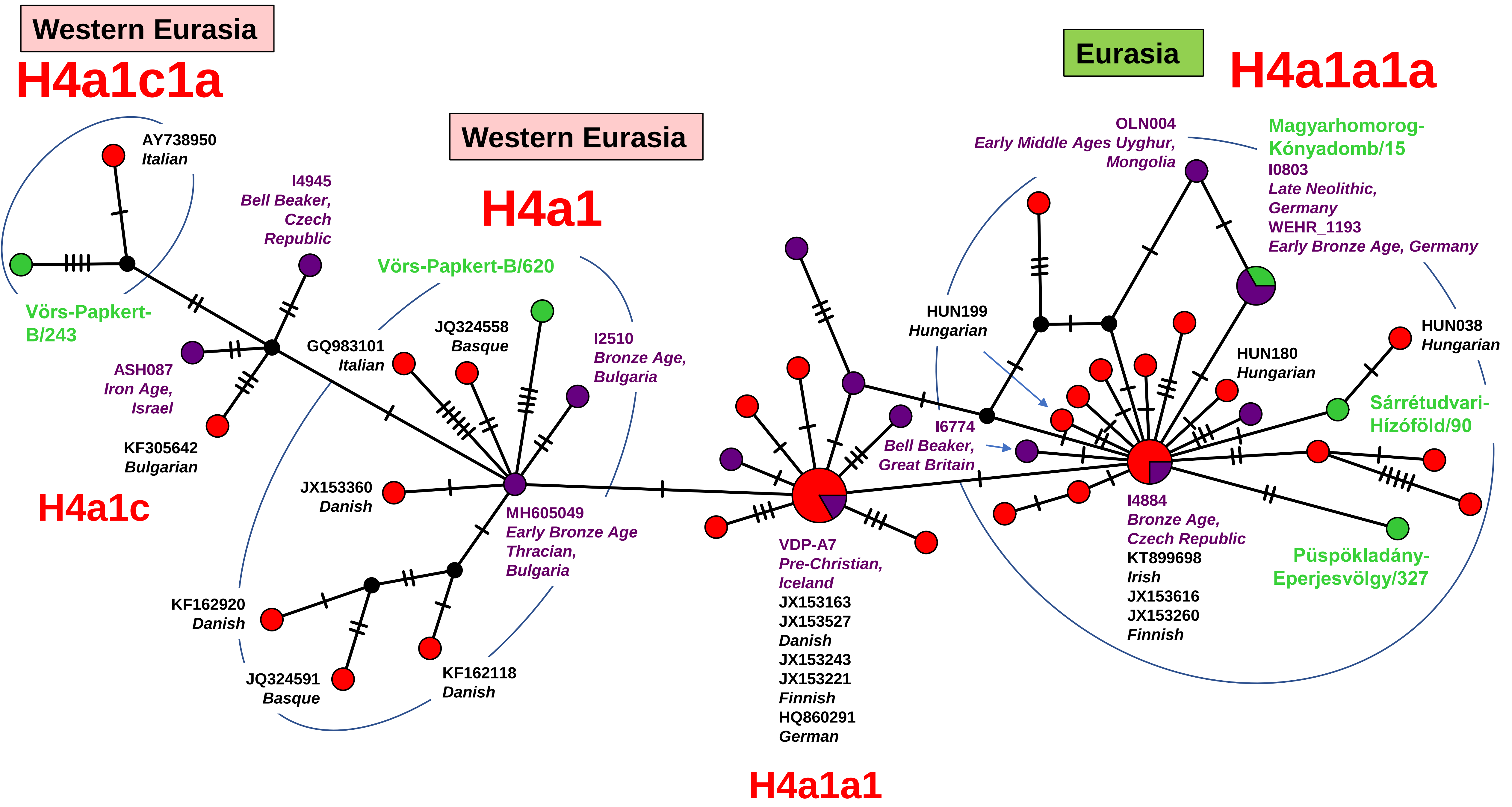

Western Eurasia
H4a1c1a
Western Eurasia
H4a1
Eurasia
H4a1a1a
AY738950
Italian
I4945
Bell Beaker,
Czech
Republic
Vörs-Papkert-B/620
Vörs-Papkert-
B/243
ASH087
Iron Age,
Israel
KF305642
Bulgarian
H4a1c
GQ983101
Italian
JQ324558
Basque
I2510
Bronze Age,
Bulgaria
JX153360
Danish
MH605049
Early Bronze Age
Thracian,
Bulgaria
KF162920
Danish
JQ324591
Basque
KF162118
Danish
OLN004
Early Middle Ages Uyghur,
Mongolia
Magyarhomorog-
Kónyadomb/15
I0803
Late Neolithic,
Germany
WEHR_1193
Early Bronze Age, Germany
HUN199
Hungarian
HUN180
Hungarian
HUN038
Hungarian
I6774
Bell Beaker,
Great Britain
Sárrétudvari-
Hízóföld/90
VDP-A7
Pre-Christian,
Iceland
JX153163
JX153527
Danish
JX153243
JX153221
Finnish
HQ860291
German
I4884
Bronze Age,
Czech Republic
KT899698
Irish
JX153616
JX153260
Finnish
Püspökladány-
Eperjesvölgy/327
H4a1a1


Network 20

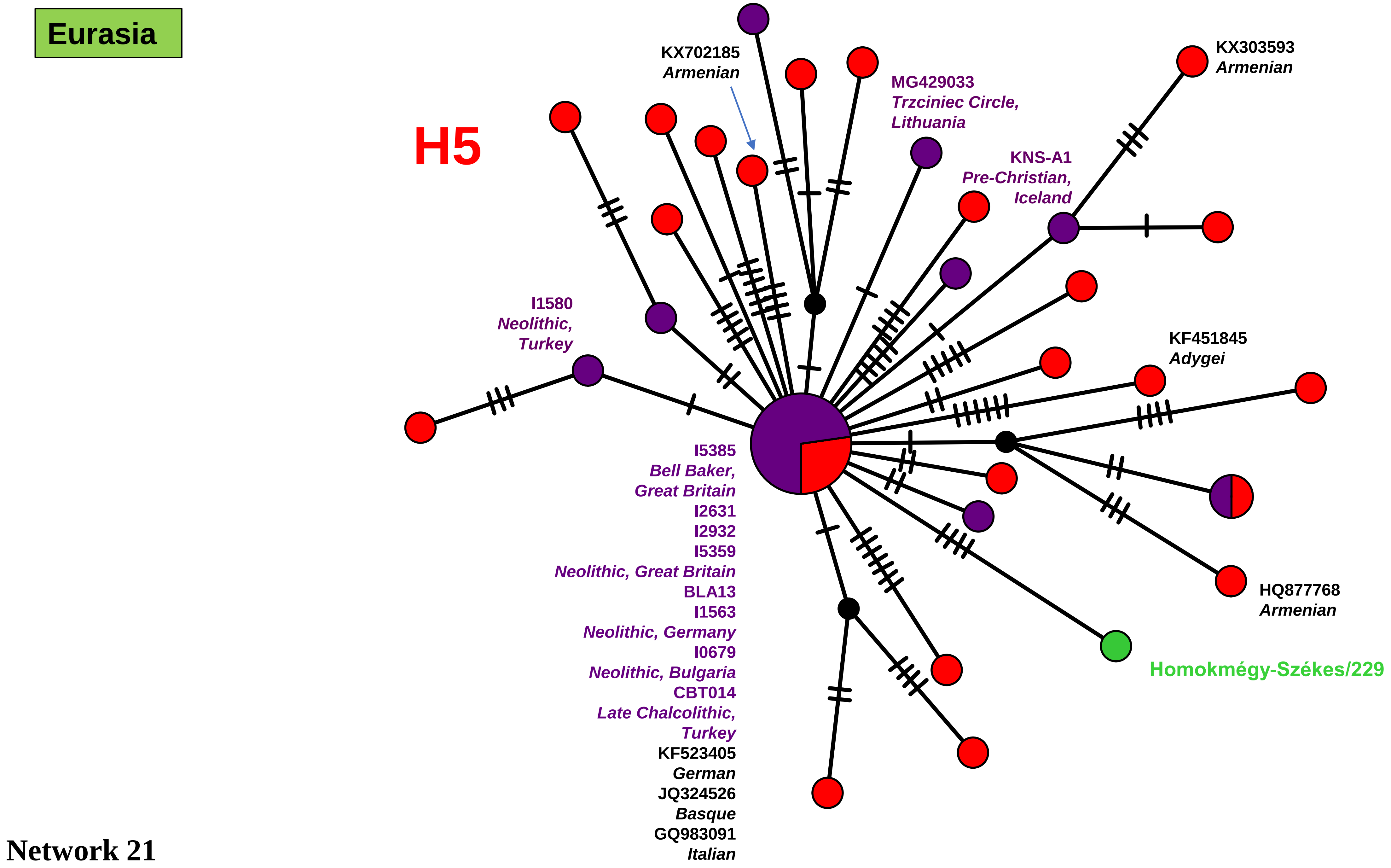
Eurasia
H5
KX702185
Armenian
MG429033
Trzciniec Circle,
Lithuania
KX303593
Armenian
KNS-A1
Pre-Christian,
Iceland
I1580
Neolithic,
Turkey
KF451845
Adygei
I5385
Bell Baker,
Great Britain
I2631
I2932
I5359
Neolithic, Great Britain
BLA13
I1563
Neolithic, Germany
I0679
Neolithic, Bulgaria
CBT014
Late Chalcolithic,
Turkey
KF523405
German
JQ324526
Basque
GQ983091
Italian
HQ877768
Armenian
Homokmégy-Székes/229
Network 21

Western Eurasia

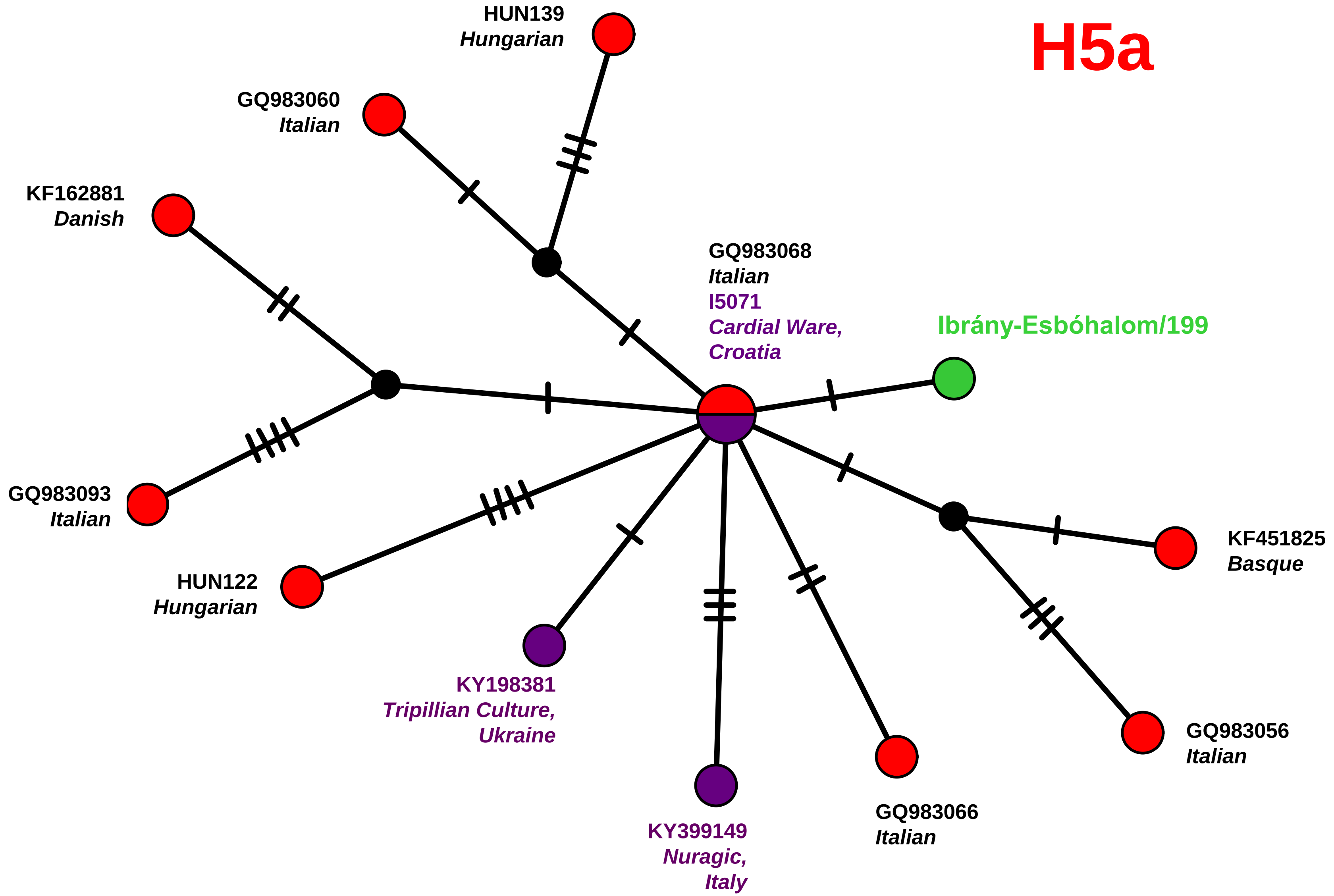


Network 22

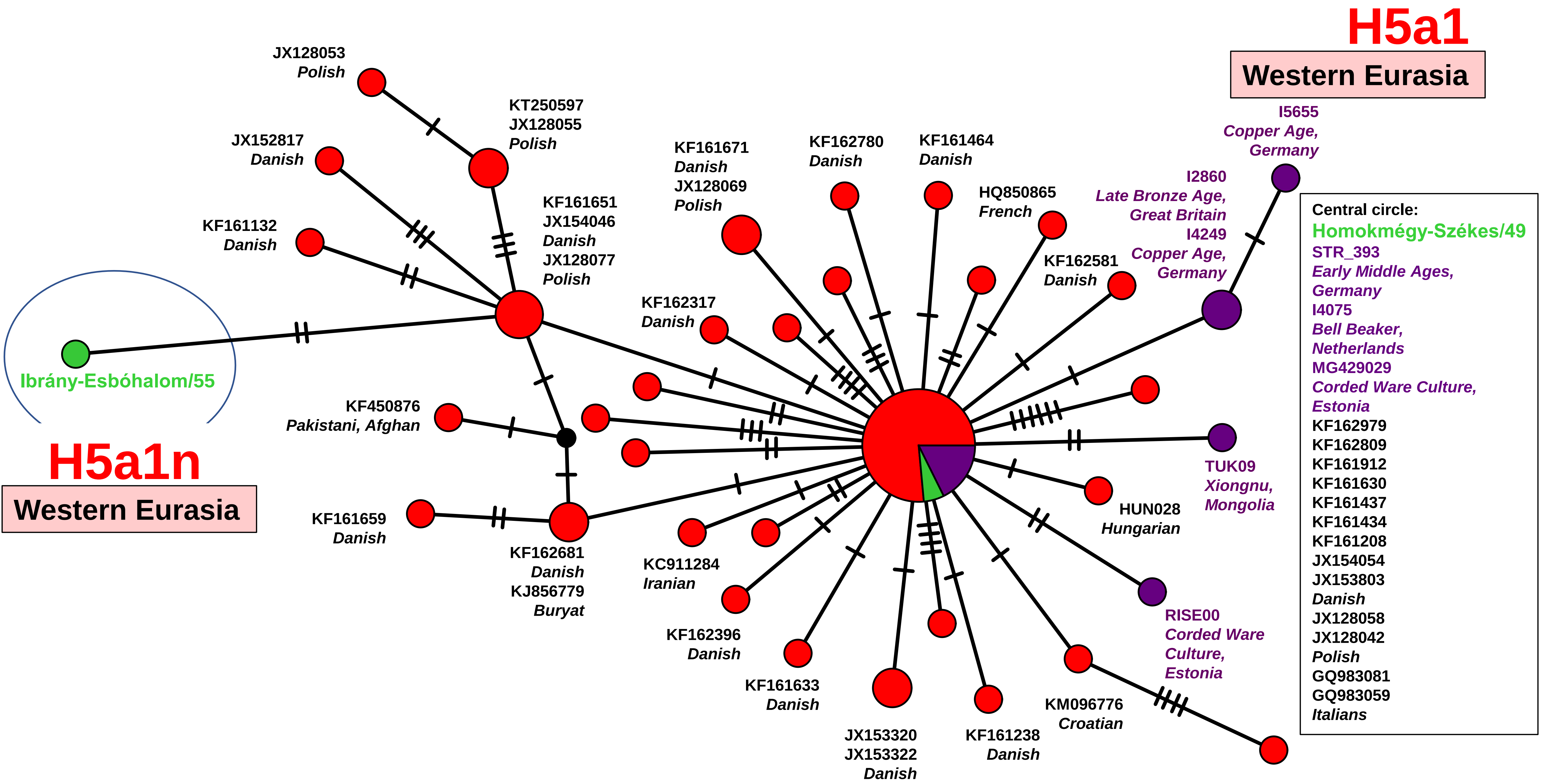

H5a1
Western Eurasia
I5655
Copper Age,
Germany
I2860
Late Bronze Age,
Great Britain
I4249
Copper Age,
Germany
Central circle:
Homokmégy-Székes/49
STR_393
Early Middle Ages,
Germany
I4075
Bell Beaker,
Netherlands
MG429029
Corded Ware Culture,
Estonia
KF162979
KF162809
KF161912
KF161630
KF161437
KF161434
KF161208
JX154054
JX153803
Danish
JX128058
JX128042
Polish
GQ983081
GQ983059
Italians
JX128053
Polish
KT250597
JX128055
Polish
JX152817
Danish
KF161132
Danish
KF161651
JX154046
Danish
JX128077
Polish
KF161671
Danish
JX128069
Polish
KF162780
Danish
KF161464
Danish
HQ850865
French
KF162581
Danish
KF162317
Danish
Ibrány-Esbóhalom/55
H5a1n
Western Eurasia
KF450876
Pakistani, Afghan
KF161659
Danish
KF162681
Danish
KJ856779
Buryat
KC911284
Iranian
KF162396
Danish
KF161633
Danish
JX153320
JX153322
Danish
KF161238
Danish
KM096776
Croatian
RISE00
Corded Ware
Culture,
Estonia
HUN028
Hungarian
TUK09
Xiongnu,
Mongolia


Network 23

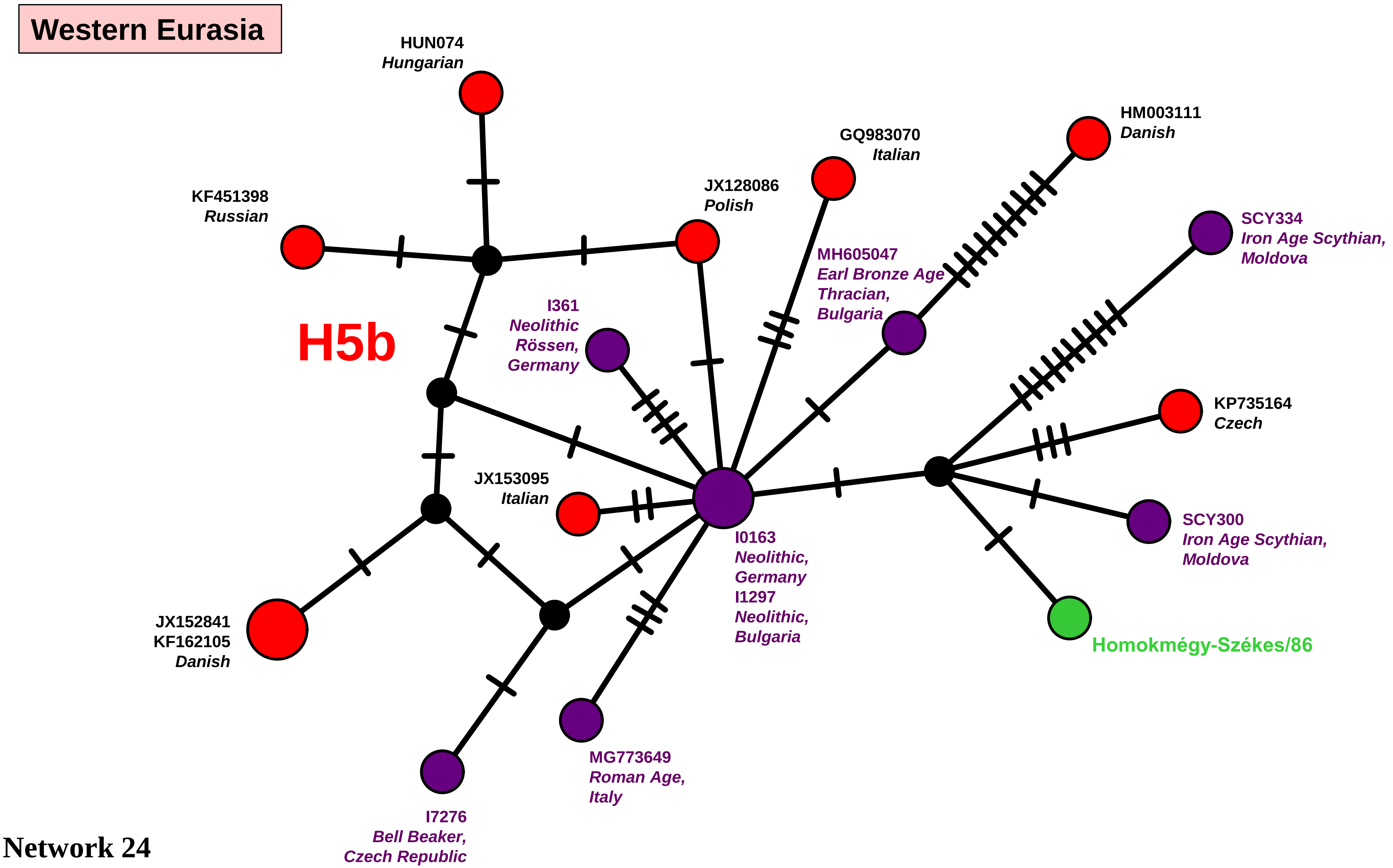
Western Eurasia
H5b
HUN074
Hungarian
KF451398
Russian
JX128086
Polish
GQ983070
Italian
HM003111
Danish
SCY334
Iron Age Scythian,
Moldova
MH605047
Earl Bronze Age
Thracian,
Bulgaria
I361
Neolithic
Rössen,
Germany
KP735164
Czech
JX153095
Italian
SCY300
Iron Age Scythian,
Moldova
I0163
Neolithic,
Germany
I1297
Neolithic,
Bulgaria
JX152841
KF162105
Danish
Homokmégy-Székes/86
MG773649
Roman Age,
Italy
I7276
Bell Beaker,
Czech Republic

Network 24

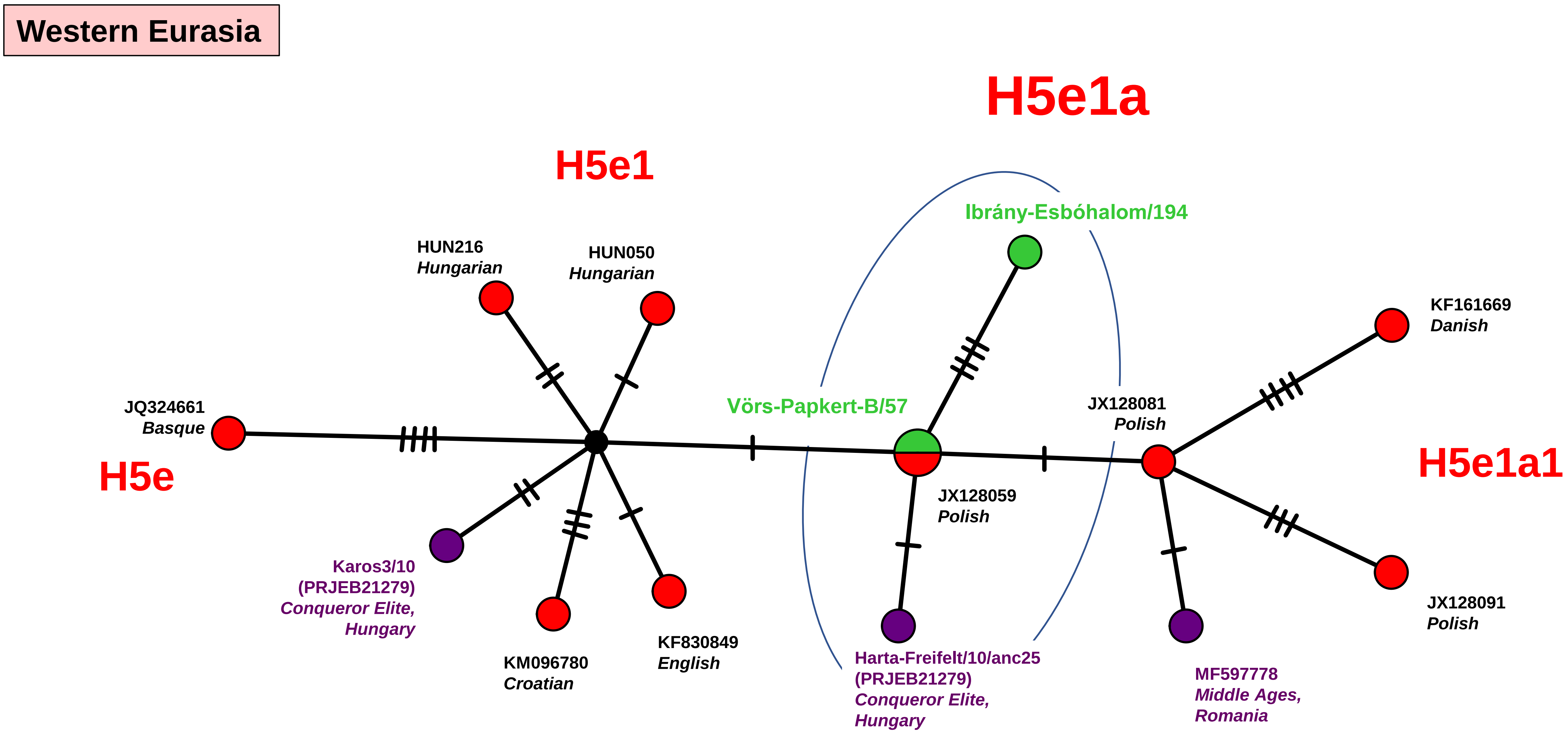
Western Eurasia
H5e1a
H5e1
Ibrány-Esbóhalom/194
HUN216
Hungarian
HUN050
Hungarian
KF161669
Danish
JQ324661
Basque
Vörs-Papkert-B/57
JX128081
Polish
H5e1a1
H5e
JX128059
Polish
Karos3/10
(PRJEB21279)
Conqueror Elite,
Hungary
JX128091
Polish
KF830849
English
KM096780
Croatian
Harta-Freifelt/10/anc25
(PRJEB21279)
Conqueror Elite,
Hungary
MF597778
Middle Ages,
Romania

Network 25

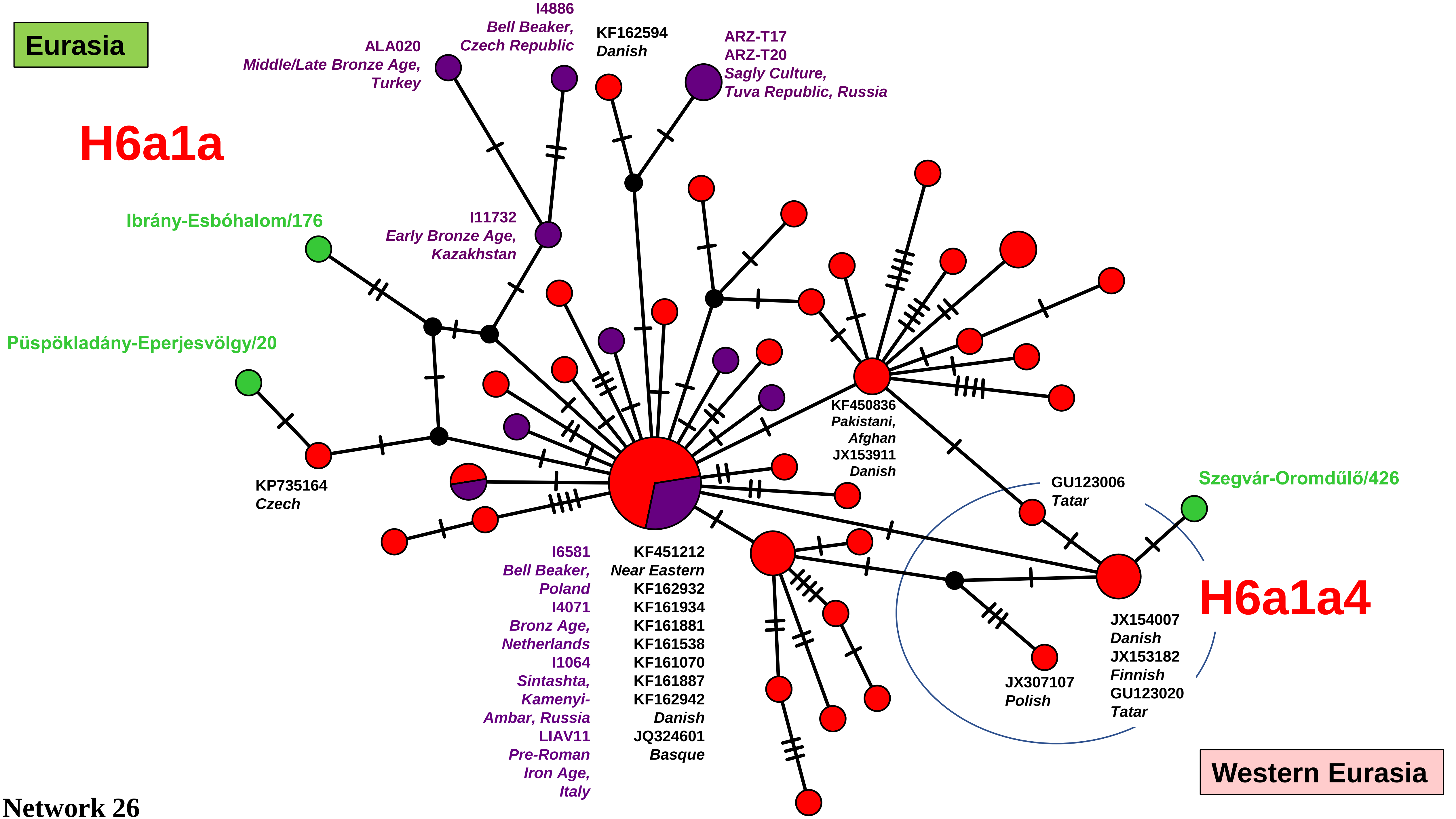

Eurasia
H6a1a
I4886
Bell Beaker,
Czech Republic
ALA020
Middle/Late Bronze Age,
Turkey
KF162594
Danish
ARZ-T17
ARZ-T20
Sagly Culture,
Tuva Republic, Russia
Ibrány-Esbóhalom/176
I11732
Early Bronze Age,
Kazakhstan
Püspökladány-Eperjesvölgy/20
KP735164
Czech
KF450836
Pakistani,
Afghan
JX153911
Danish
GU123006
Tatar
Szegvár-Oromdűlő/426
H6a1a4
I6581
Bell Beaker,
Poland
I4071
Bronz Age,
Netherlands
I1064
Sintashta,
Kamenyi-
Ambar, Russia
LIAV11
Pre-Roman
Iron Age,
Italy
KF451212
Near Eastern
KF162932
KF161934
KF161881
KF161538
KF161070
KF161887
KF162942
Danish
JQ324601
Basque
JX154007
Danish
JX153182
Finnish
GU123020
Tatar
JX307107
Polish
Western Eurasia


Network 26

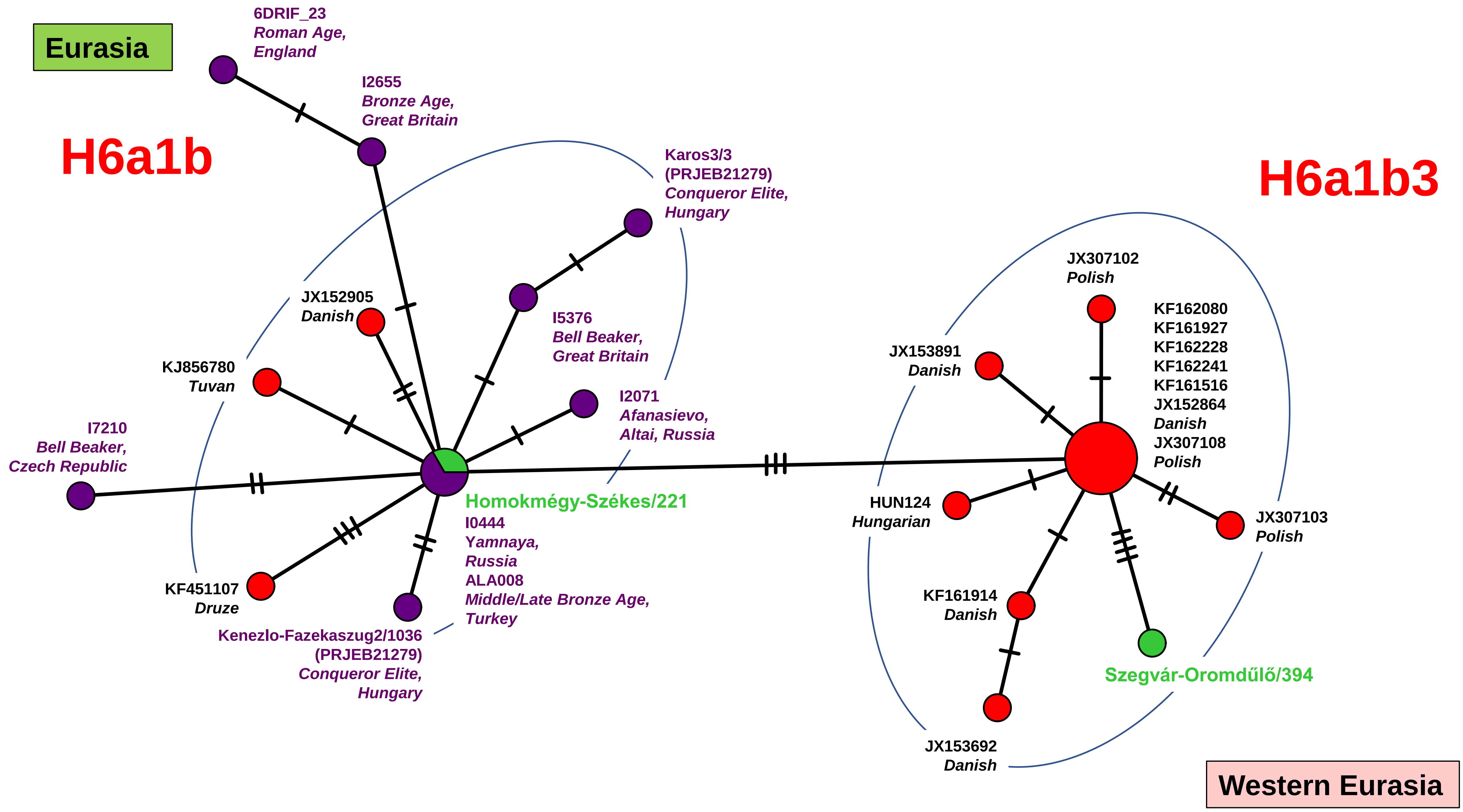

Eurasia
H6a1b
H6a1b3
6DRIF_23
Roman Age,
England
I2655
Bronze Age,
Great Britain
Karos3/3
(PRJEB21279)
Conqueror Elite,
Hungary
I5376
Bell Beaker,
Great Britain
I2071
Afanasievo,
Altai, Russia
JX152905
Danish
KJ856780
Tuvan
I7210
Bell Beaker,
Czech Republic
KF451107
Druze
Kenezlo-Fazekaszug2/1036
(PRJEB21279)
Conqueror Elite,
Hungary
Homokmégy-Székes/221
I0444
Yamnaya,
Russia
ALA008
Middle/Late Bronze Age,
Turkey
JX307102
Polish
KF162080
KF161927
KF162228
KF162241
KF161516
JX152864
Danish
JX307108
Polish
JX153891
Danish
HUN124
Hungarian
JX307103
Polish
KF161914
Danish
Szegvár-Oromdűlő/394
JX153692
Danish
Western Eurasia

Network 27

**Eastern Eurasia**

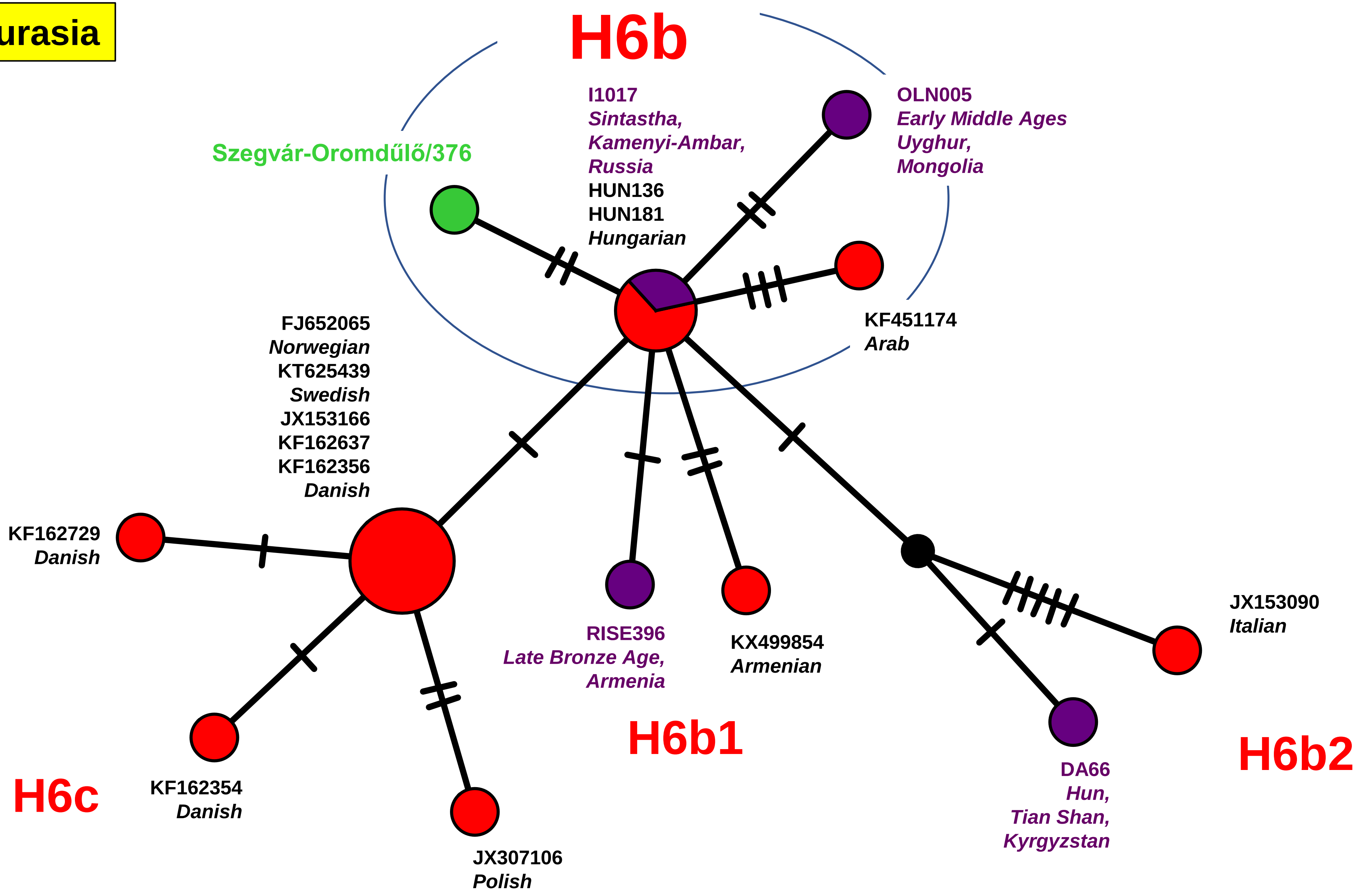


Network 28

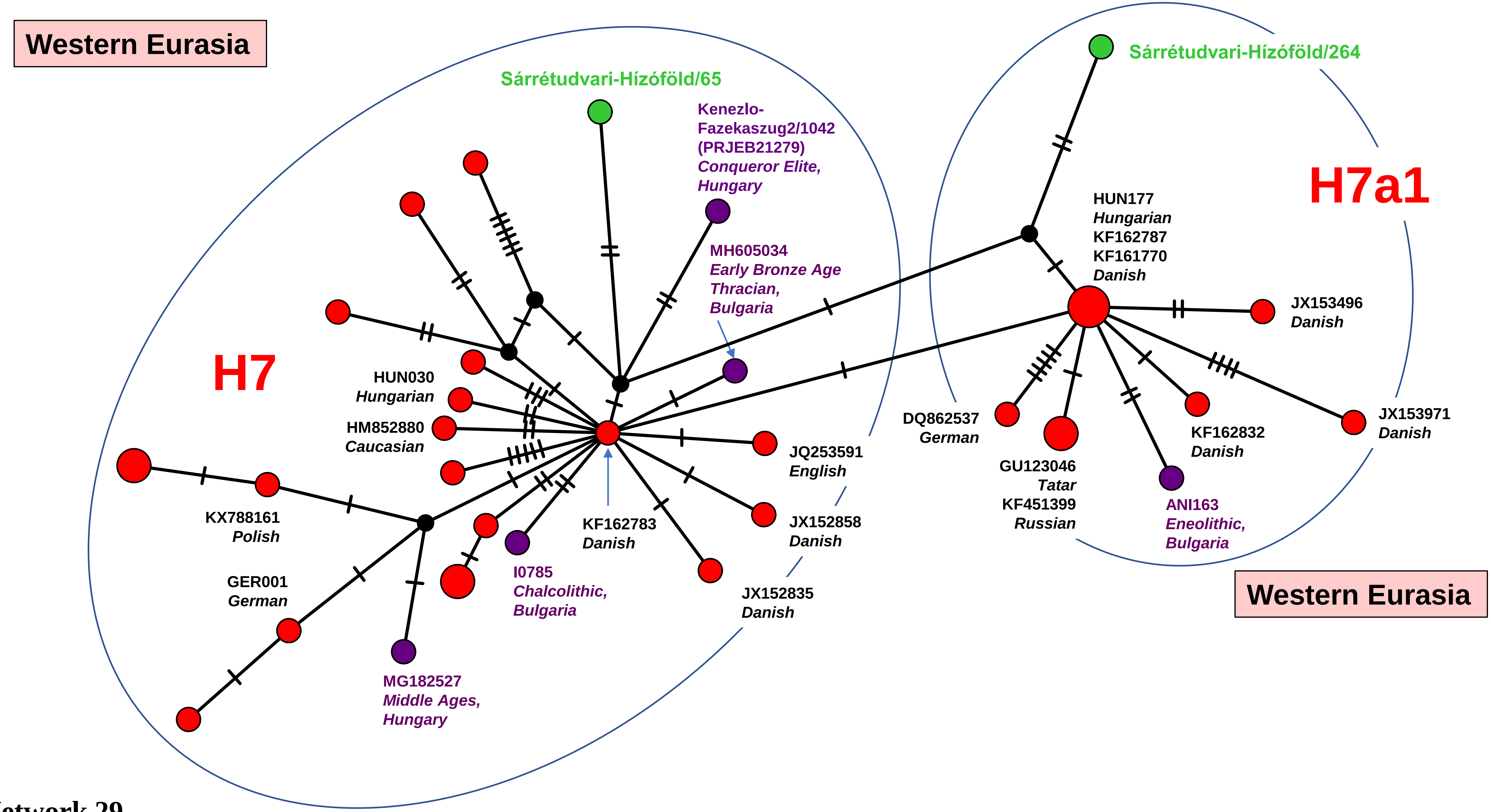
Western Eurasia
Sárrétudvari-Hízóföld/65
Kenezlo-
Fazekaszug2/1042
(PRJEB21279)
Conqueror Elite,
Hungary
MH605034
Early Bronze Age
Thracian,
Bulgaria
H7
HUN030
Hungarian
HM852880
Caucasian
KX788161
Polish
GER001
German
MG182527
Middle Ages,
Hungary
I0785
Chalcolithic,
Bulgaria
KF162783
Danish
JQ253591
English
JX152858
Danish
JX152835
Danish
Sárrétudvari-Hízóföld/264
H7a1
HUN177
Hungarian
KF162787
KF161770
Danish
JX153496
Danish
DQ862537
German
GU123046
Tatar
KF451399
Russian
ANI163
Eneolithic,
Bulgaria
KF162832
Danish
JX153971
Danish
Western Eurasia

Network 29

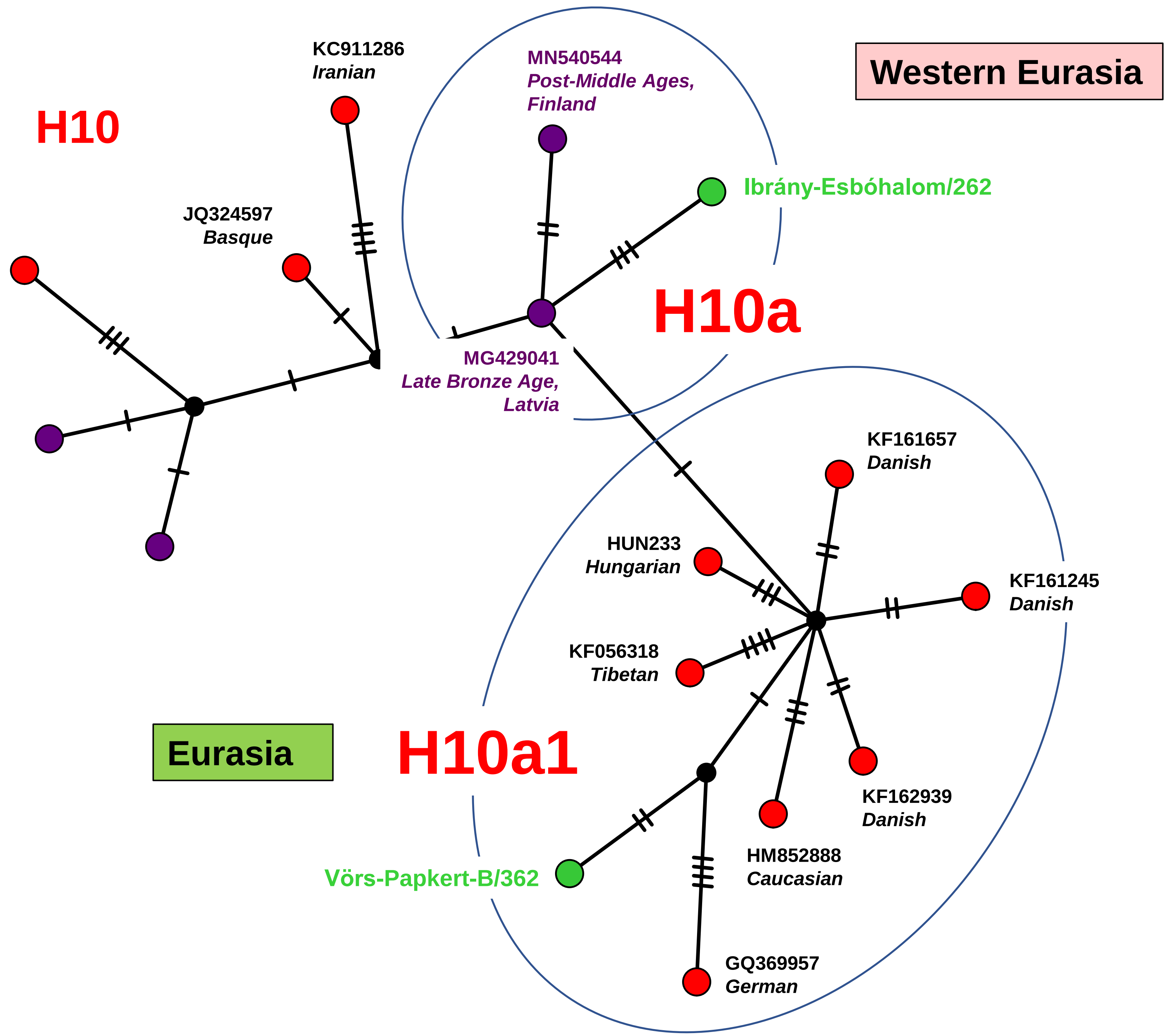

H10
KC911286
Iranian
JQ324597
Basque
MN540544
Post-Middle Ages,
Finland
Ibrány-Esbóhalom/262
H10a
MG429041
Late Bronze Age,
Latvia
Western Eurasia
KF161657
Danish
HUN233
Hungarian
KF161245
Danish
KF056318
Tibetan
Eurasia
H10a1
KF162939
Danish
HM852888
Caucasian
Vörs-Papkert-B/362
GQ369957
German


Network 30

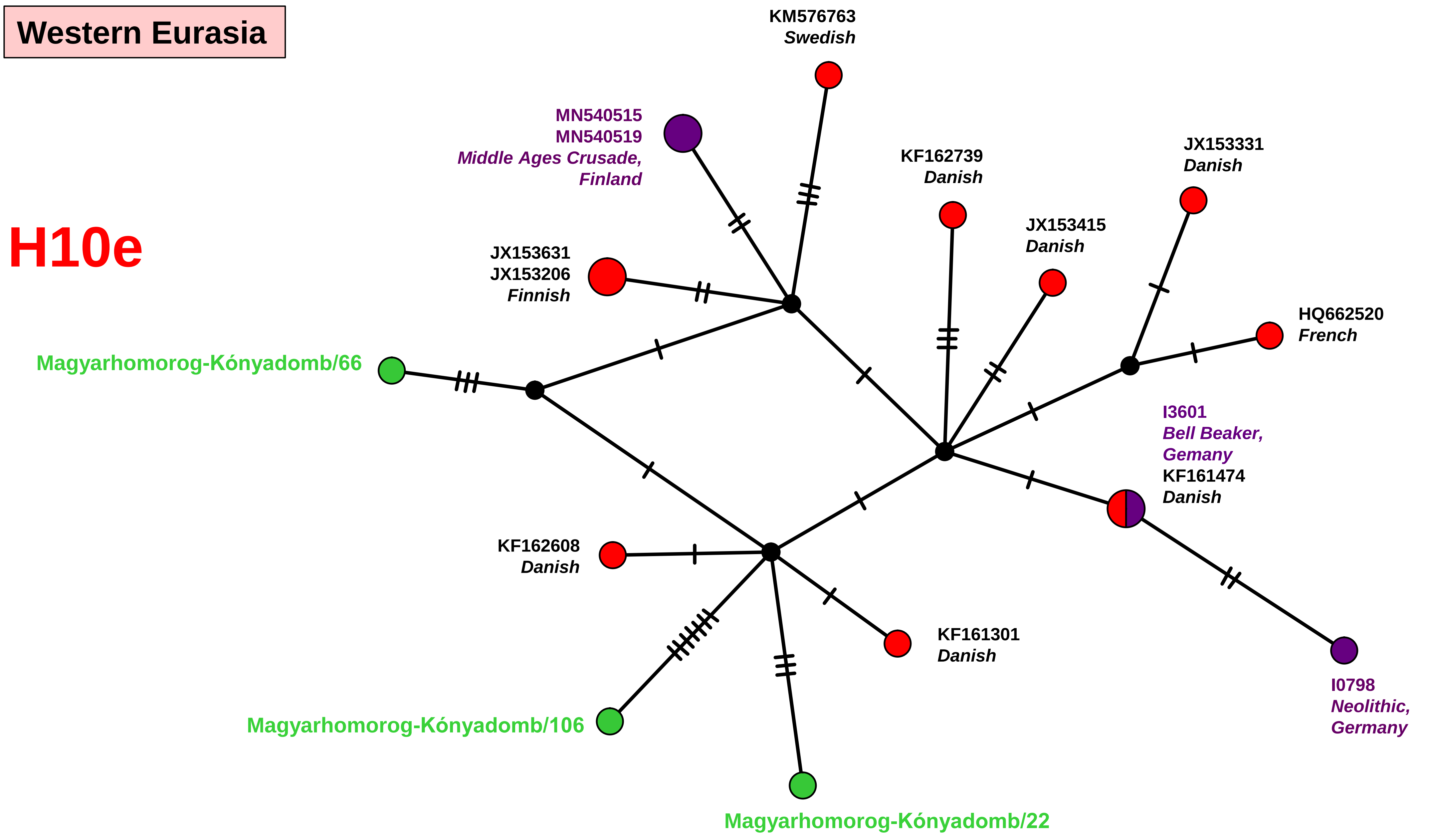
Western Eurasia
H10e
KM576763
Swedish
MN540515
MN540519
Middle Ages Crusade,
Finland
KF162739
Danish
JX153331
Danish
JX153415
Danish
JX153631
JX153206
Finnish
HQ662520
French
Magyarhomorog-Kónyadomb/66
I3601
Bell Beaker,
Gemany
KF161474
Danish
KF162608
Danish
KF161301
Danish
I0798
Neolithic,
Germany
Magyarhomorog-Kónyadomb/106
Magyarhomorog-Kónyadomb/22

Network 31

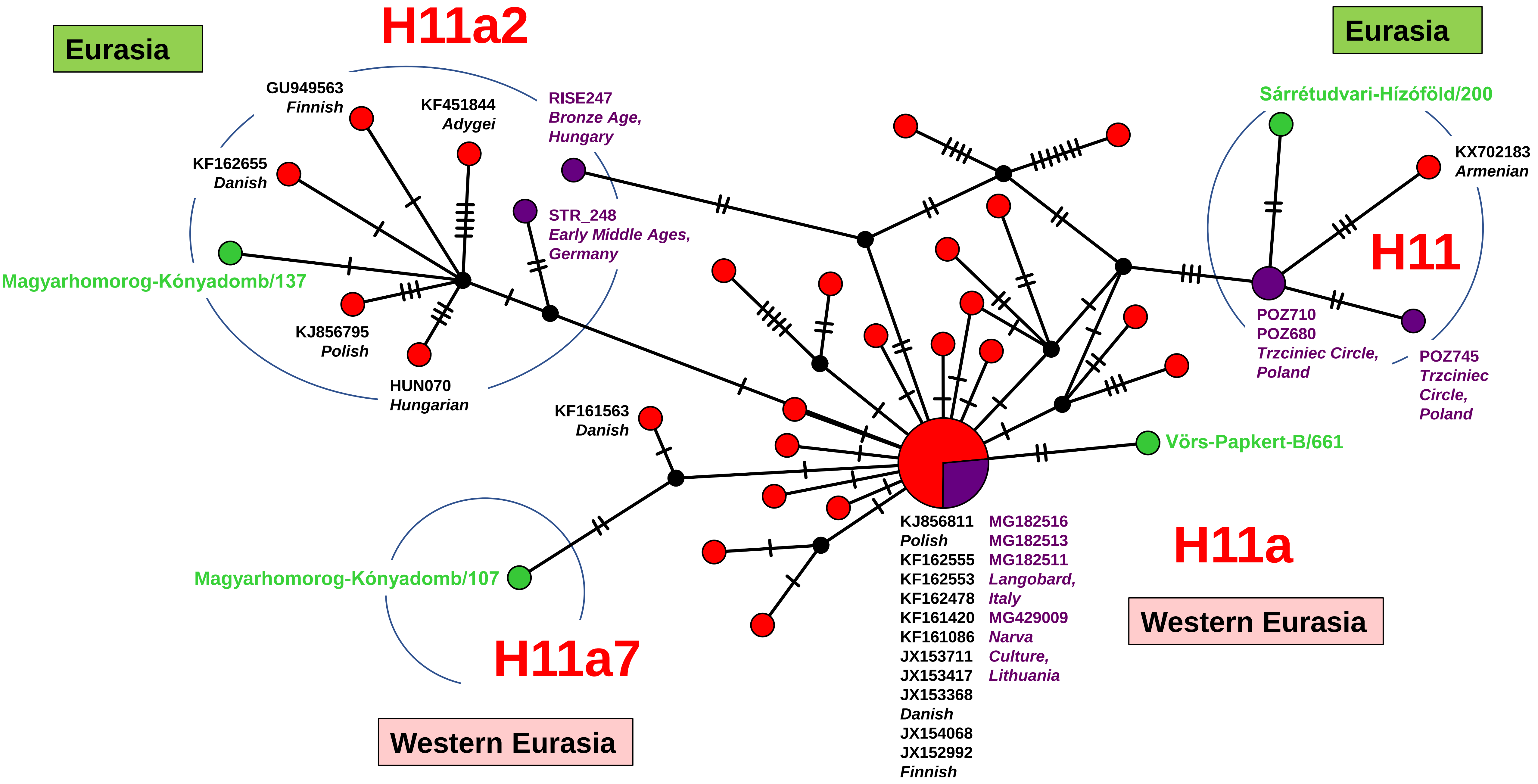

H11a2
Eurasia
GU949563
Finnish
KF451844
Adygei
RISE247
Bronze Age,
Hungary
KF162655
Danish
STR_248
Early Middle Ages,
Germany
Magyarhomorog-Kónyadomb/137
KJ856795
Polish
HUN070
Hungarian
KF161563
Danish
Magyarhomorog-Kónyadomb/107
H11a7
Western Eurasia
Eurasia
Sárrétudvari-Hízóföld/200
KX702183
Armenian
H11
POZ710
POZ680
Trzciniec Circle,
Poland
POZ745
Trzciniec
Circle,
Poland
Vörs-Papkert-B/661
H11a
Western Eurasia
KJ856811
Polish
KF162555
KF162553
KF162478
KF161420
KF161086
JX153711
JX153417
JX153368
Danish
JX154068
JX152992
Finnish
MG182516
MG182513
MG182511
Langobard,
Italy
MG429009
Narva
Culture,
Lithuania


Network 32

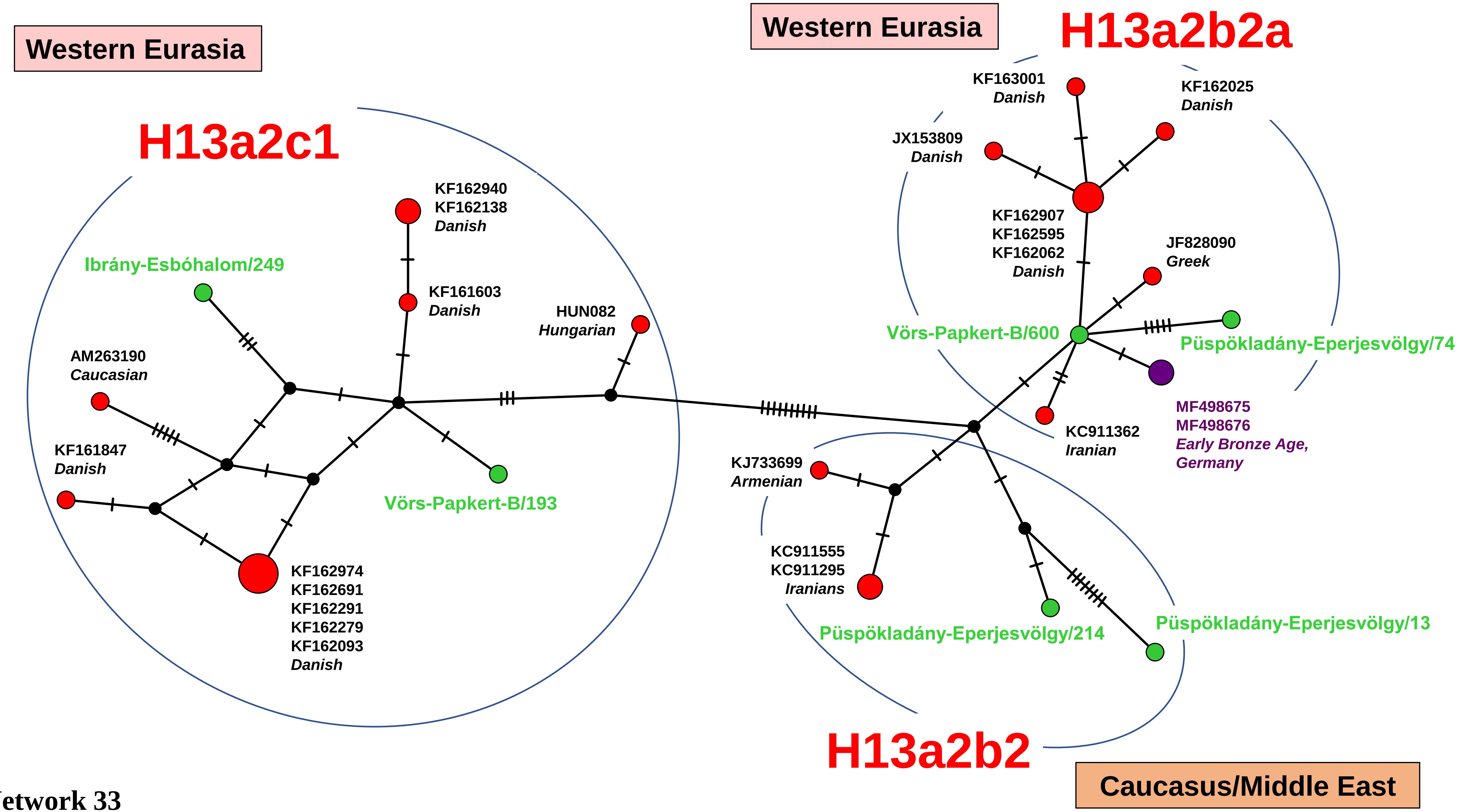
Western Eurasia
H13a2c1
KF162940
KF162138
Danish
Ibrány-Esbóhalom/249
KF161603
Danish
HUN082
Hungarian
AM263190
Caucasian
KF161847
Danish
Vörs-Papkert-B/193
KF162974
KF162691
KF162291
KF162279
KF162093
Danish
Western Eurasia
H13a2b2a
KF163001
Danish
KF162025
Danish
JX153809
Danish
KF162907
KF162595
KF162062
Danish
JF828090
Greek
Vörs-Papkert-B/600
Püspökladány-Eperjesvölgy/74
KC911362
Iranian
MF498675
MF498676
Early Bronze Age,
Germany
KJ733699
Armenian
KC911555
KC911295
Iranians
Püspökladány-Eperjesvölgy/214
Püspökladány-Eperjesvölgy/13
H13a2b2
Caucasus/Middle East

Network 33

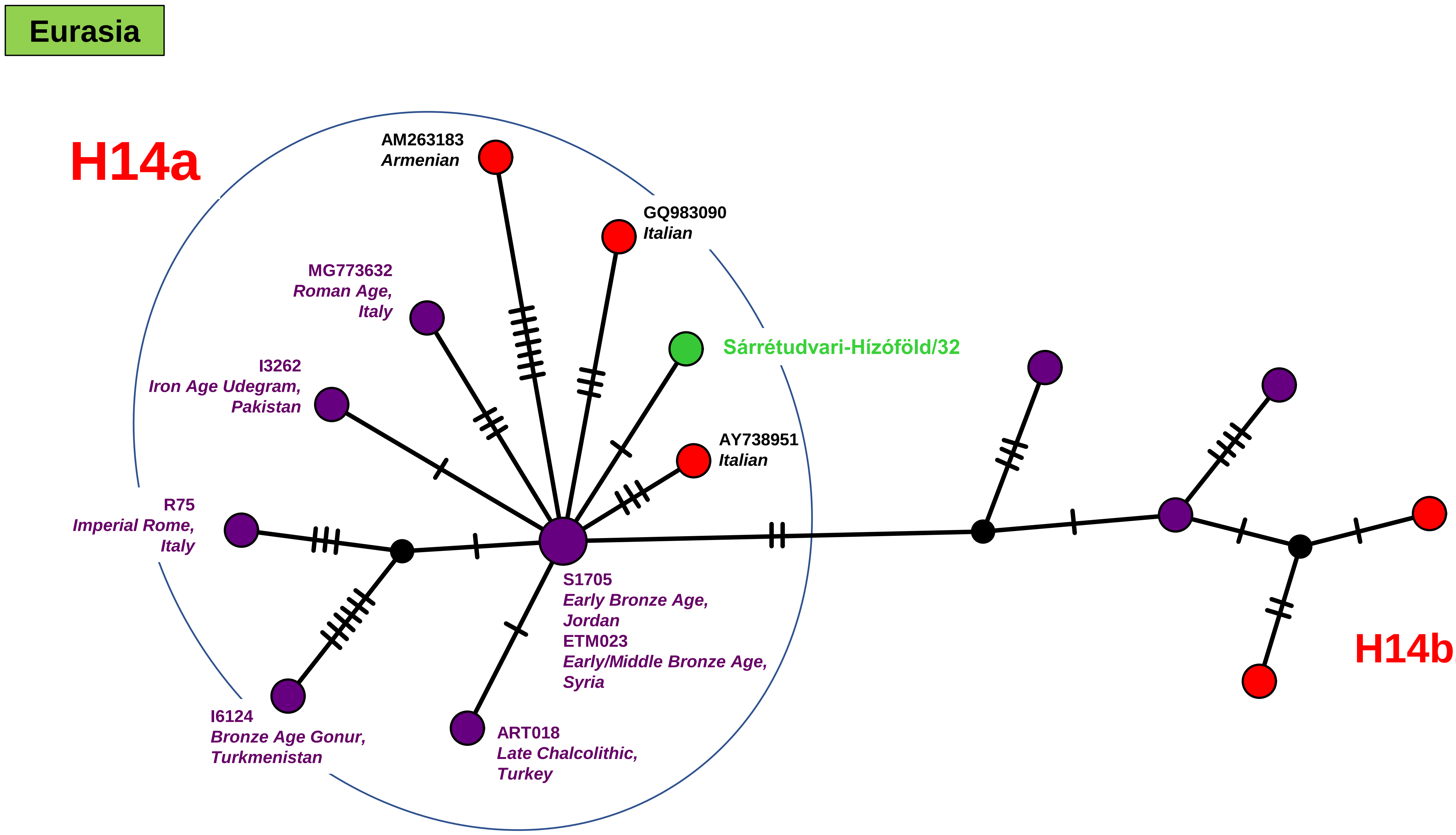
Eurasia
H14a
AM263183
Armenian
GQ983090
Italian
MG773632
Roman Age,
Italy
Sárrétudvari-Hízóföld/32
I3262
Iron Age Udegram,
Pakistan
AY738951
Italian
R75
Imperial Rome,
Italy
S1705
Early Bronze Age,
Jordan
ETM023
Early/Middle Bronze Age,
Syria
I6124
Bronze Age Gonur,
Turkmenistan
ART018
Late Chalcolithic,
Turkey
H14b2

Network 34

**Western Eurasia**

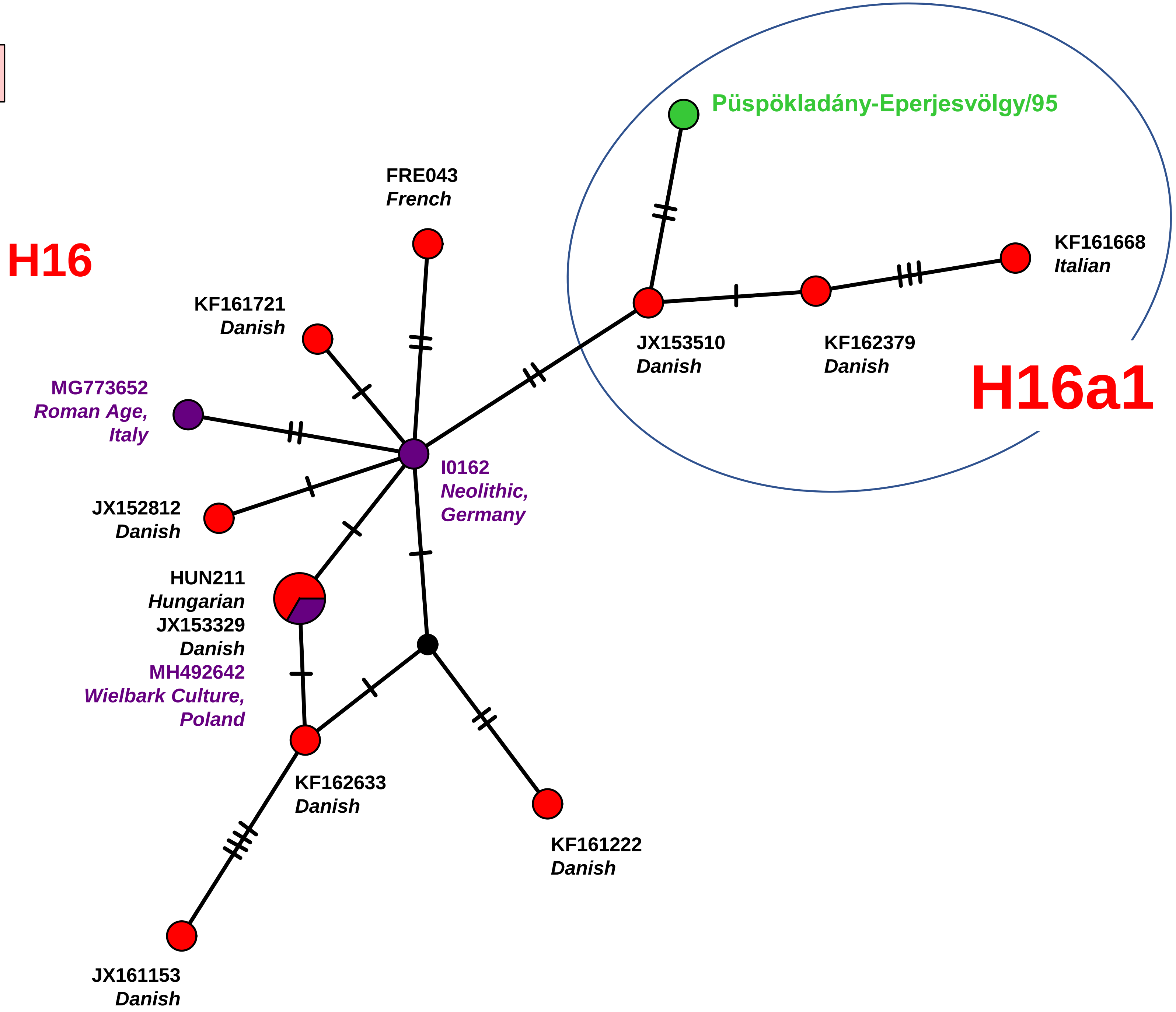


**Network 35**

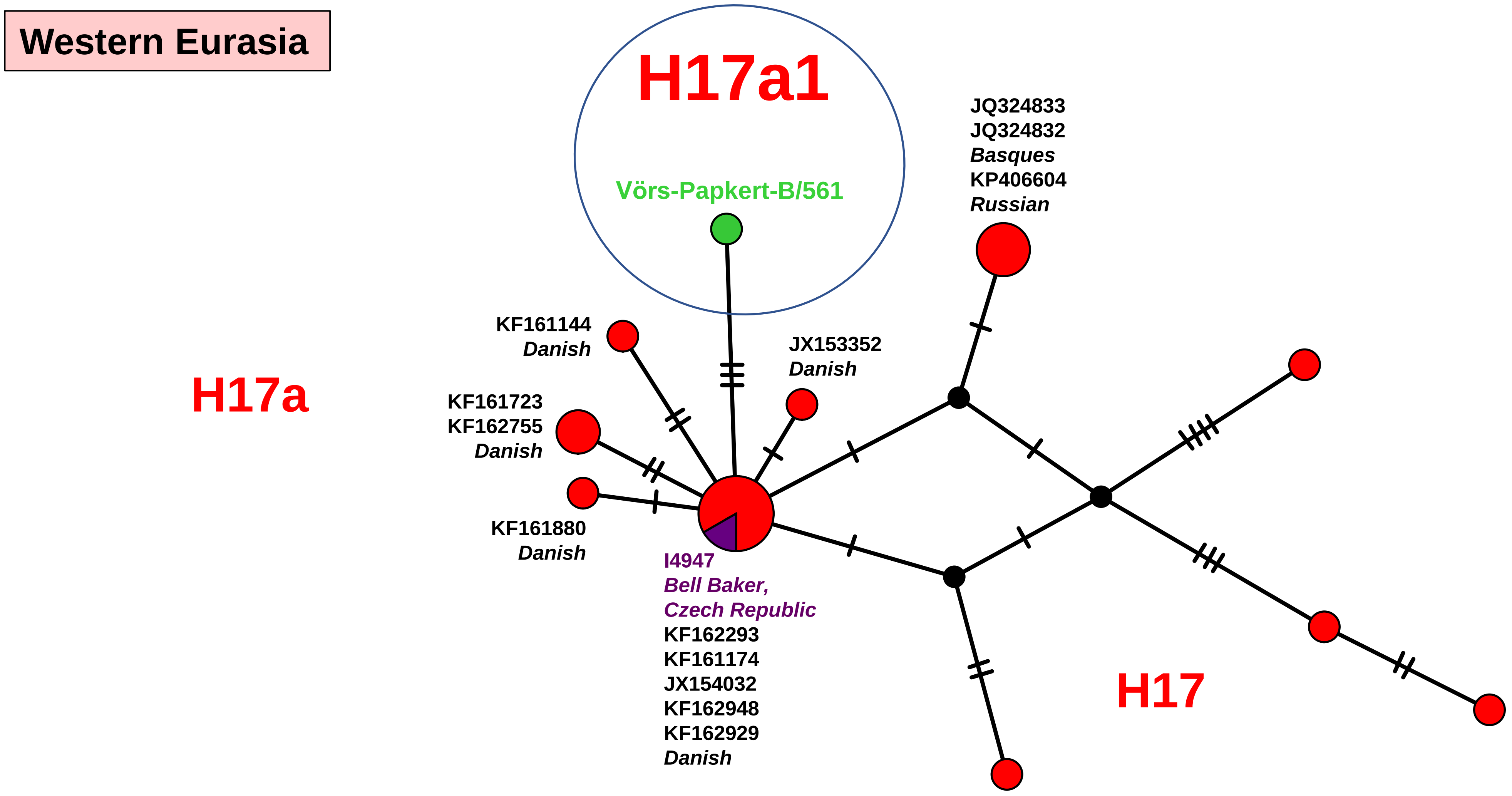
Western Eurasia
H17a1
Vörs-Papkert-B/561
JQ324833
JQ324832
Basques
KP406604
Russian
KF161144
Danish
JX153352
Danish
H17a
KF161723
KF162755
Danish
KF161880
Danish
I4947
Bell Baker,
Czech Republic
KF162293
KF161174
JX154032
KF162948
KF162929
Danish
H17

Network 36

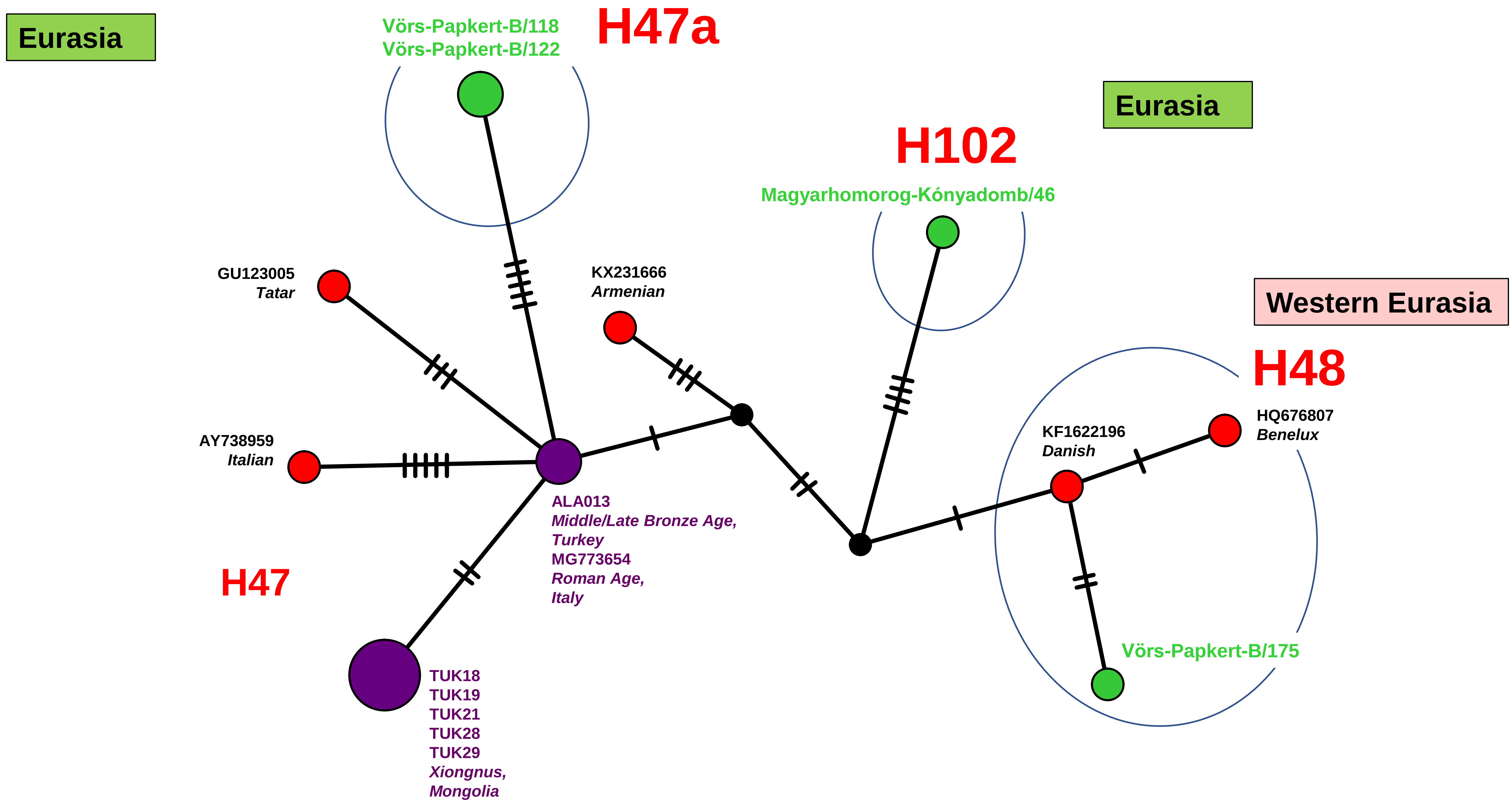
Eurasia
Vörs-Papkert-B/118
Vörs-Papkert-B/122
H47a
Eurasia
H102
Magyarhomorog-Kónyadomb/46
GU123005
Tatar
KX231666
Armenian
Western Eurasia
H48
HQ676807
Benelux
KF1622196
Danish
AY738959
Italian
ALA013
Middle/Late Bronze Age,
Turkey
MG773654
Roman Age,
Italy
H47
Vörs-Papkert-B/175
TUK18
TUK19
TUK21
TUK28
TUK29
Xiongnus,
Mongolia

Network 37

Eastern Eurasia

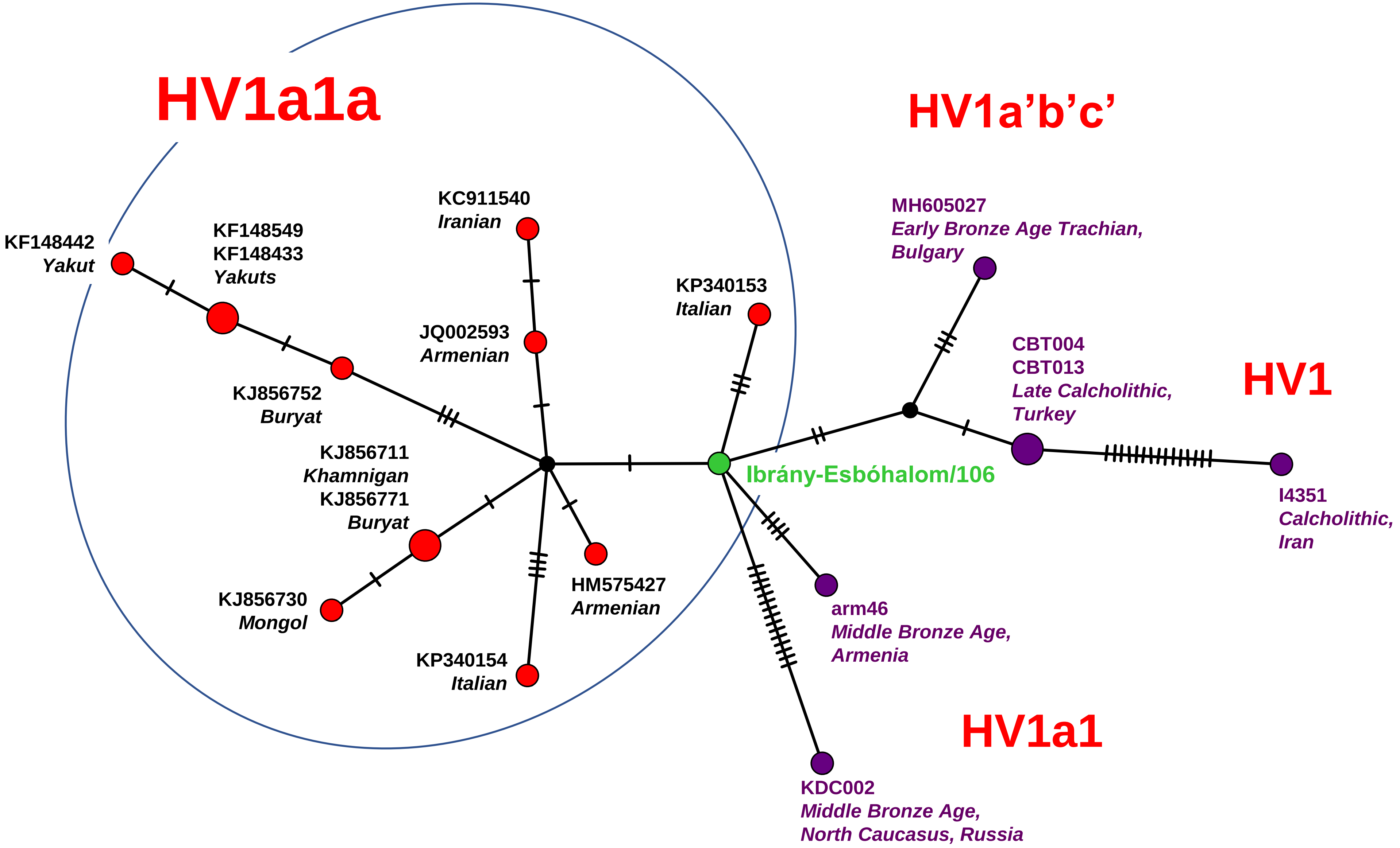


Network 38

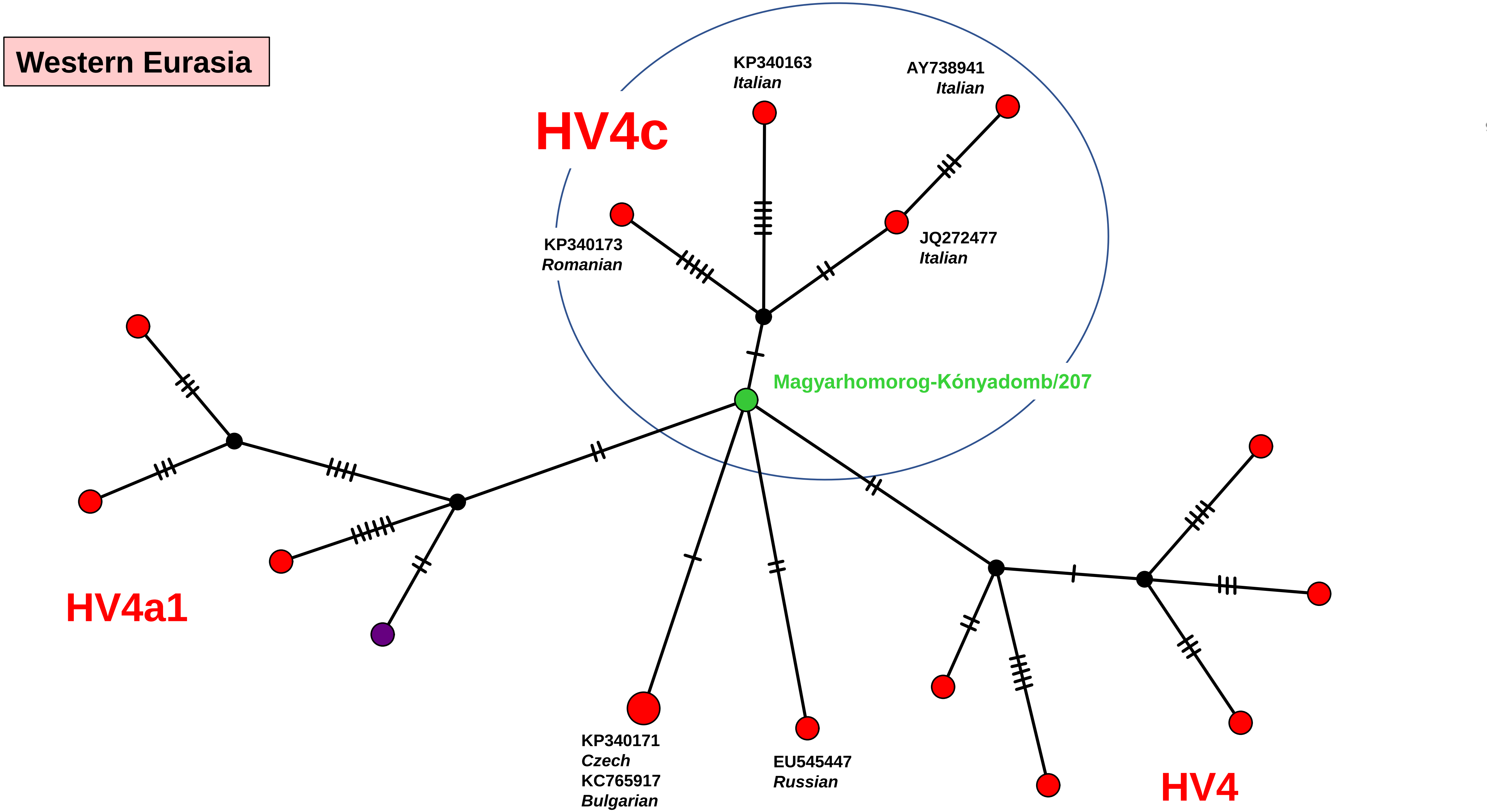

Western Eurasia
HV4c
KP340163
Italian
AY738941
Italian
KP340173
Romanian
JQ272477
Italian
Magyarhomorog-Kónyadomb/207
HV4a1
KP340171
Czech
KC765917
Bulgarian
EU545447
Russian
HV4


Network 39

### Western Eurasia

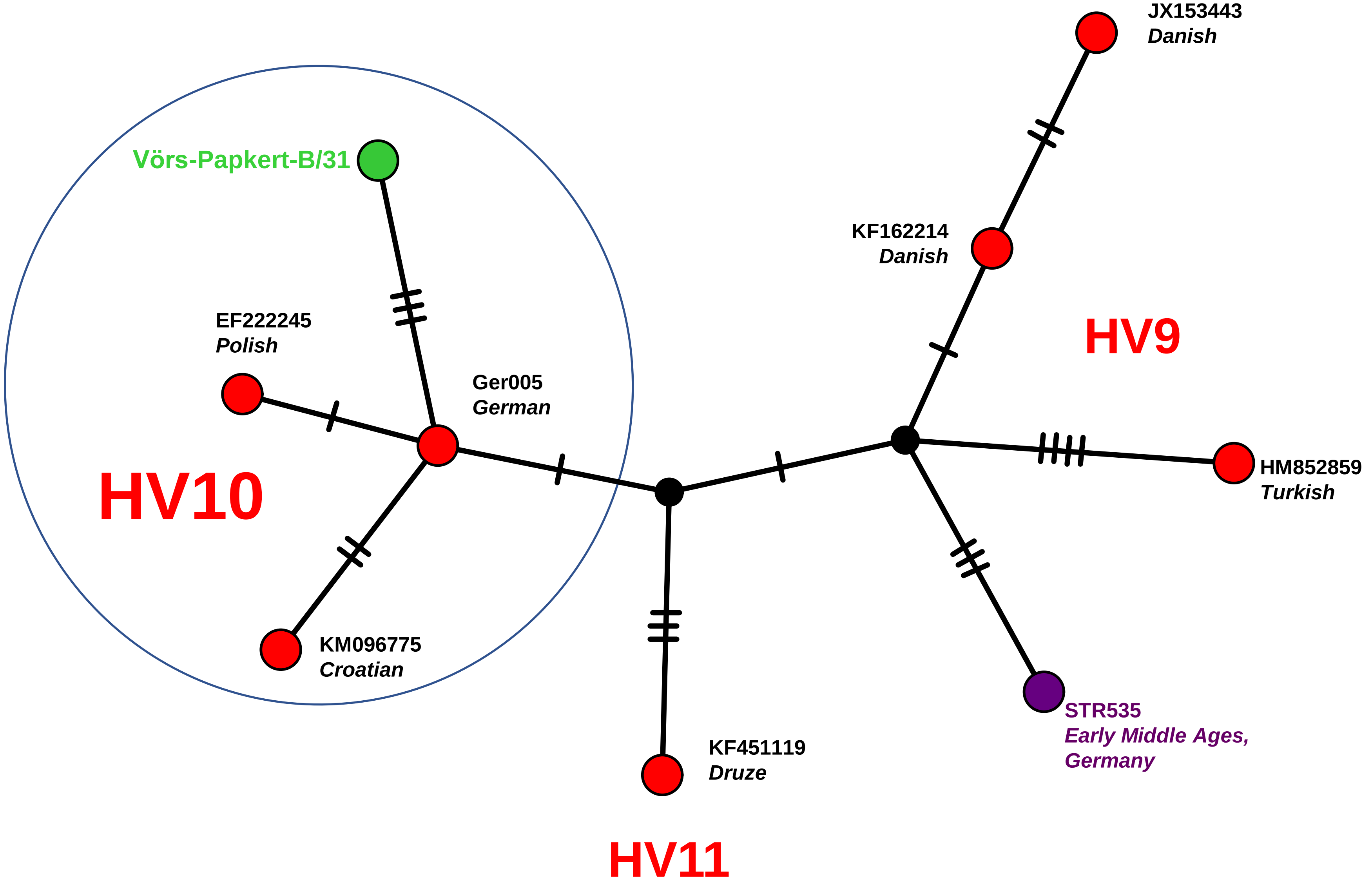


Network 40

**Western Eurasia**

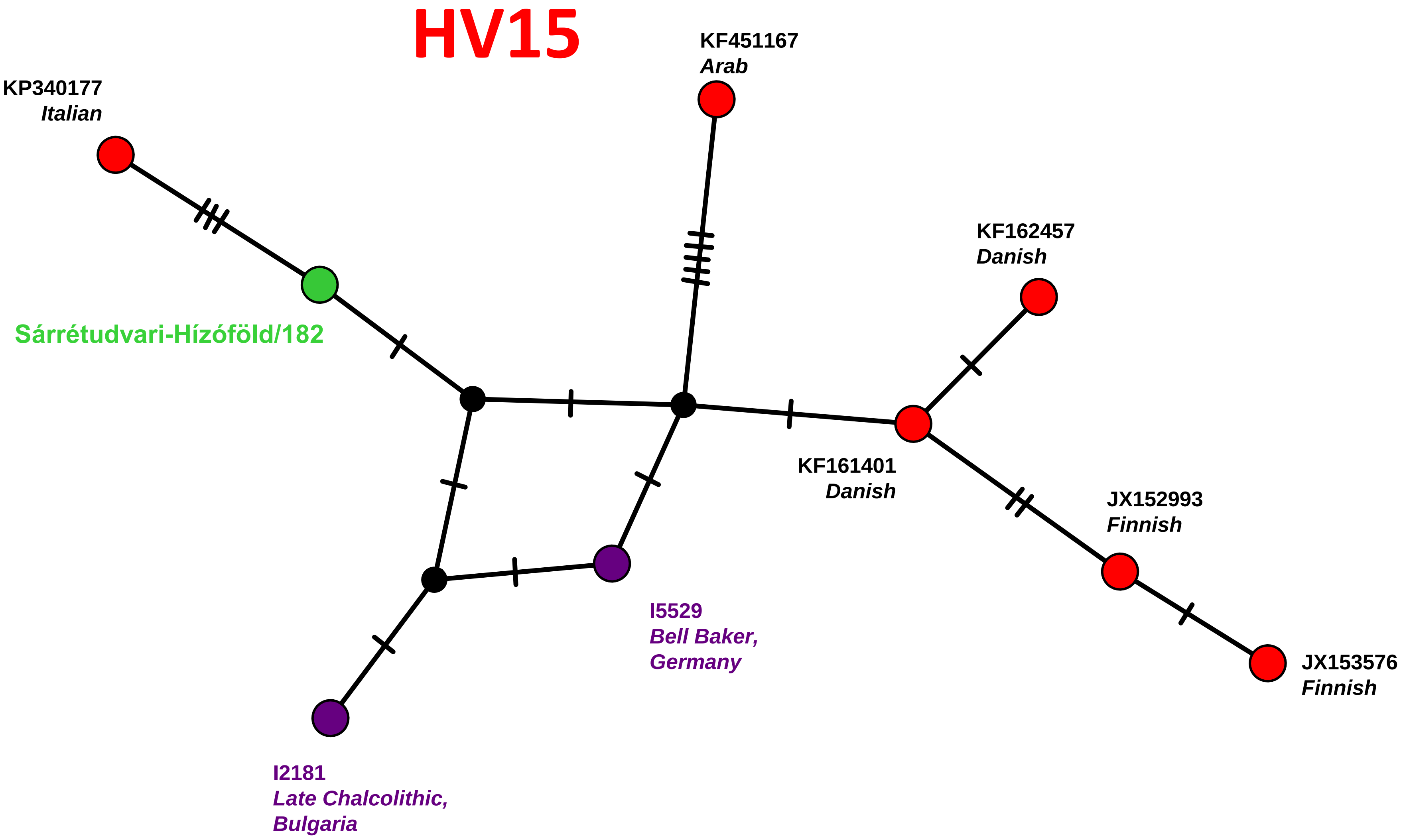


**Network 41**

Eurasia

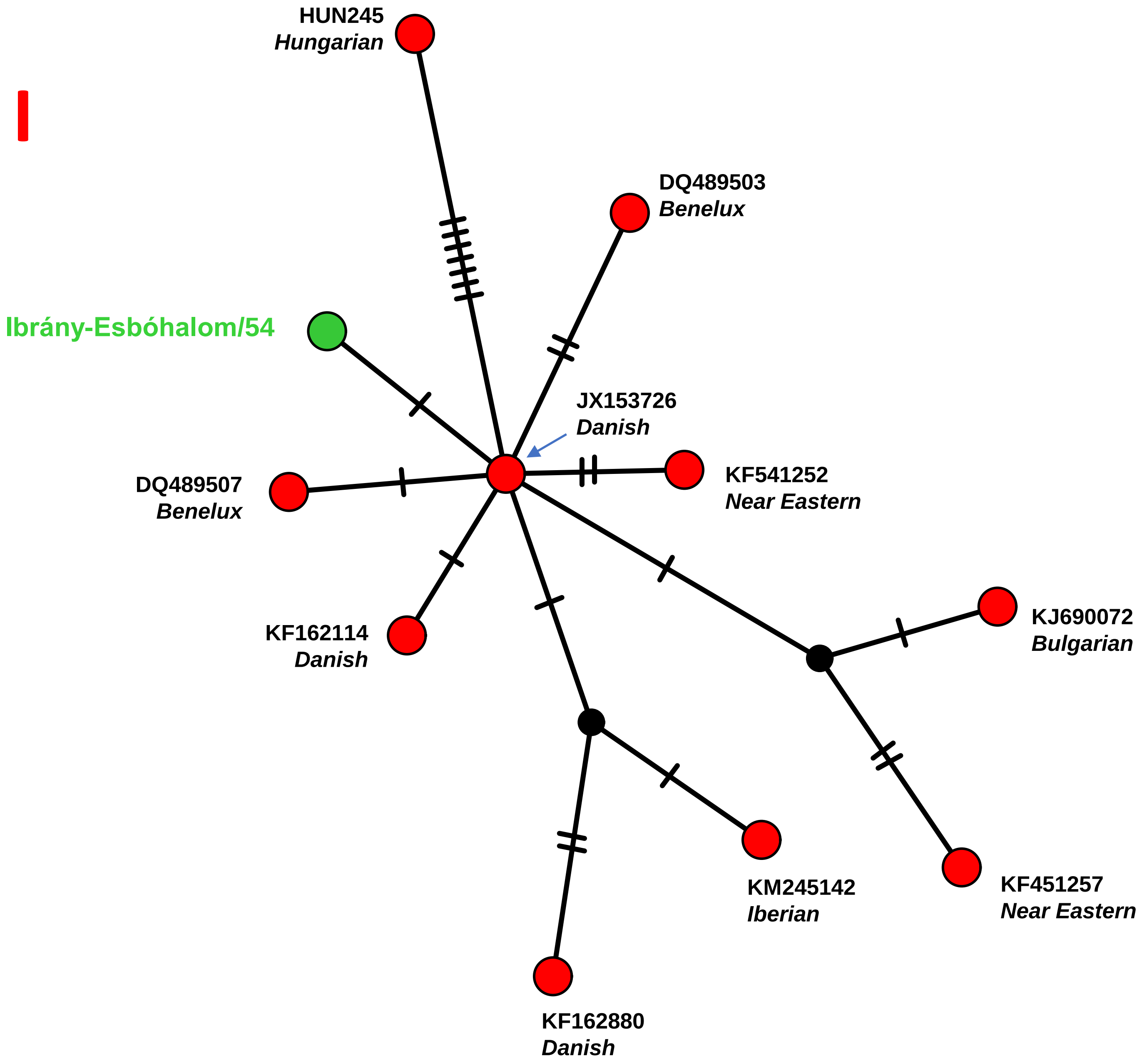


Network 42

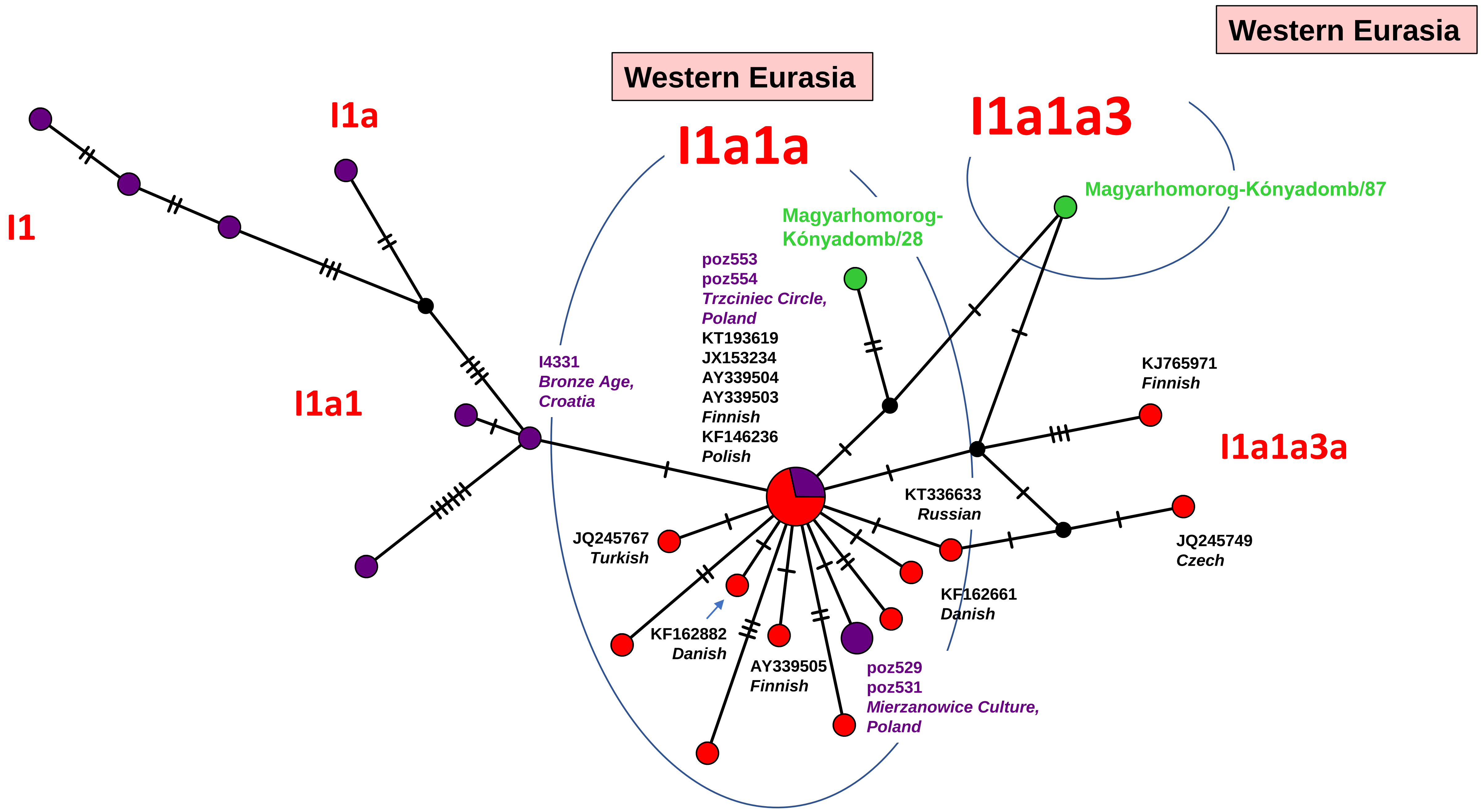
Western Eurasia
Western Eurasia
I1a
I1
I1a1
I1a1a
I1a1a3
I1a1a3a
Magyarhomorog-Kónyadomb/87
Magyarhomorog-
Kónyadomb/28
poz553
poz554
Trzciniec Circle,
Poland
KT193619
JX153234
AY339504
AY339503
Finnish
KF146236
Polish
I4331
Bronze Age,
Croatia
KJ765971
Finnish
KT336633
Russian
JQ245767
Turkish
KF162882
Danish
AY339505
Finnish
poz529
poz531
Mierzanowice Culture,
Poland
KF162661
Danish
JQ245749
Czech

Network 43

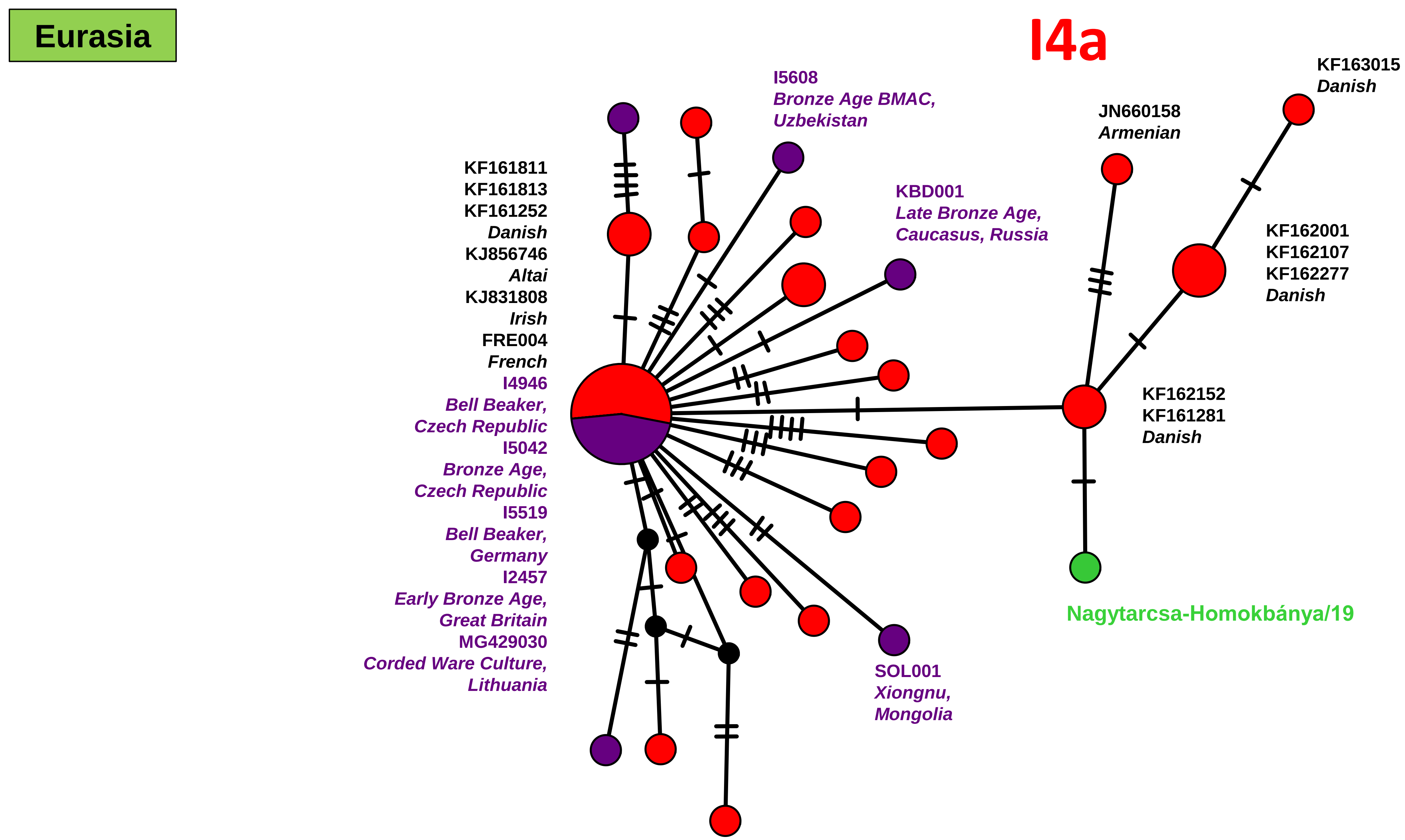
Eurasia
I4a
I5608
Bronze Age BMAC,
Uzbekistan
KF163015
Danish
JN660158
Armenian
KBD001
Late Bronze Age,
Caucasus, Russia
KF161811
KF161813
KF161252
Danish
KJ856746
Altai
KJ831808
Irish
FRE004
French
I4946
Bell Beaker,
Czech Republic
I5042
Bronze Age,
Czech Republic
I5519
Bell Beaker,
Germany
I2457
Early Bronze Age,
Great Britain
MG429030
Corded Ware Culture,
Lithuania
KF162001
KF162107
KF162277
Danish
KF162152
KF161281
Danish
Nagytarcsa-Homokbánya/19
SOL001
Xiongnu,
Mongolia

Network 44

### Western Eurasia

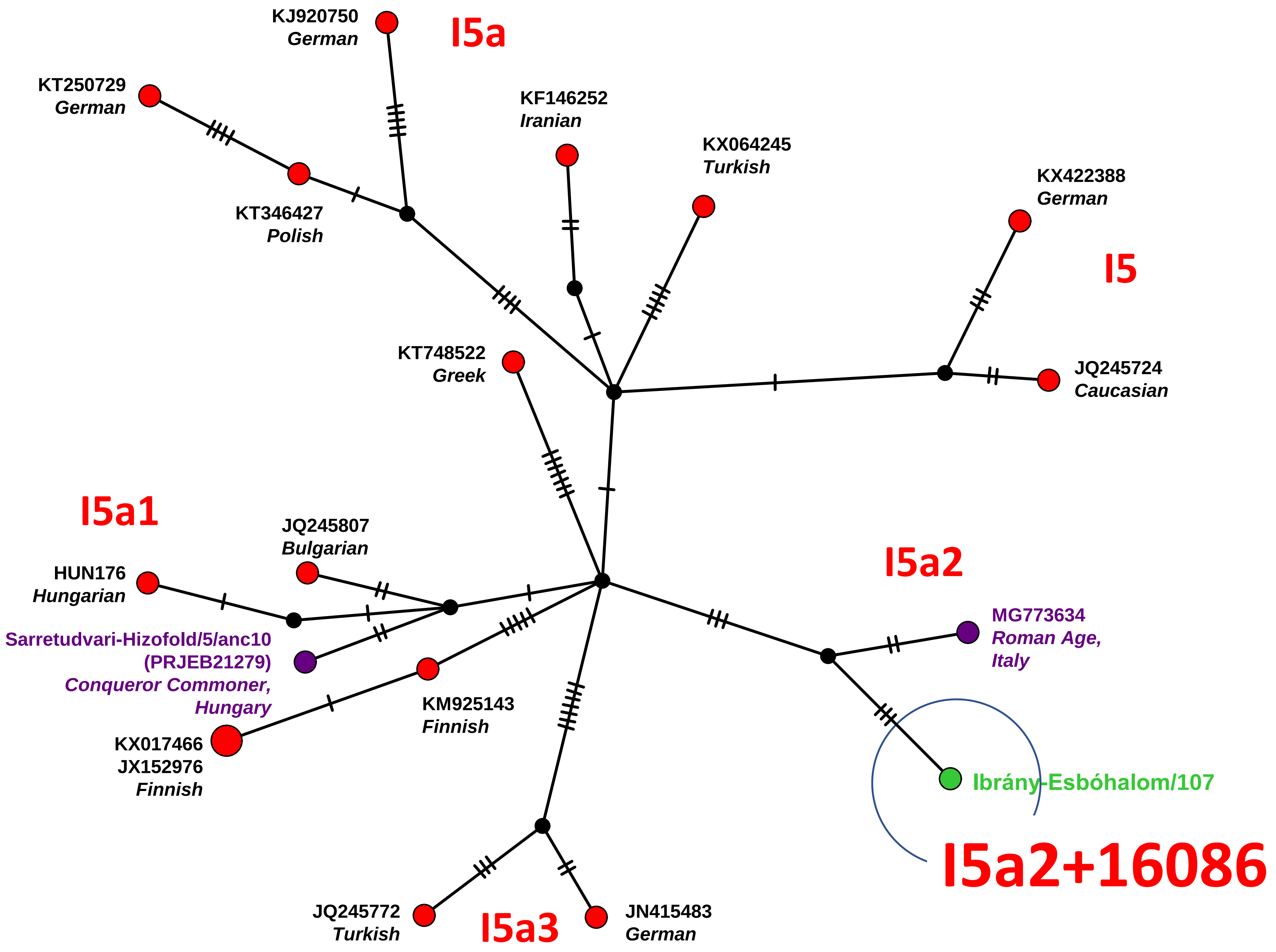


Network 45

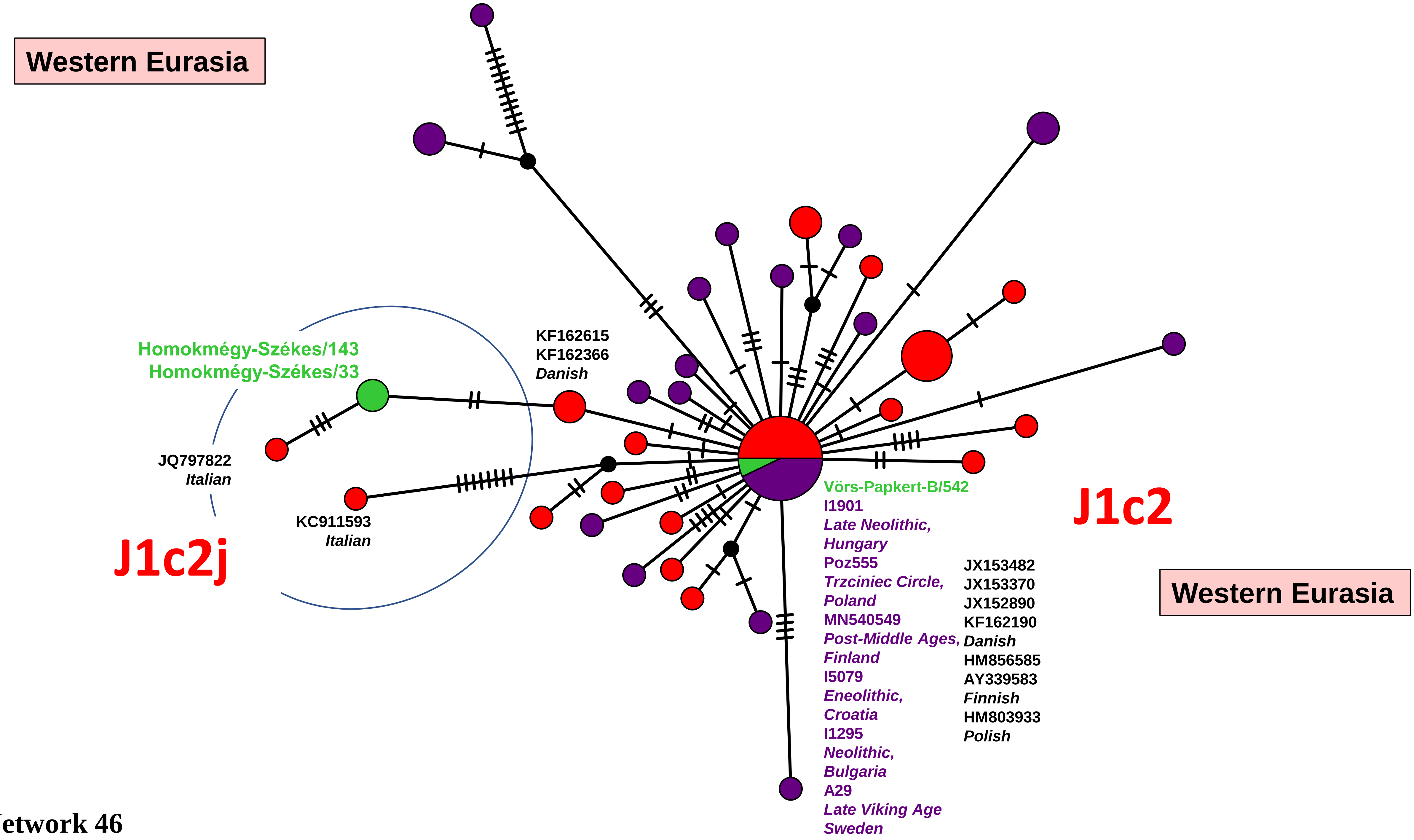

Western Eurasia
Homokmégy-Székes/143
Homokmégy-Székes/33
KF162615
KF162366
Danish
JQ797822
Italian
KC911593
Italian
J1c2j
Vörs-Papkert-B/542
I1901
Late Neolithic,
Hungary
Poz555
Trzciniec Circle,
Poland
MN540549
Post-Middle Ages,
Finland
I5079
Eneolithic,
Croatia
I1295
Neolithic,
Bulgaria
A29
Late Viking Age
Sweden
JX153482
JX153370
JX152890
KF162190
Danish
HM856585
AY339583
Finnish
HM803933
Polish
J1c2
Western Eurasia


Network 46

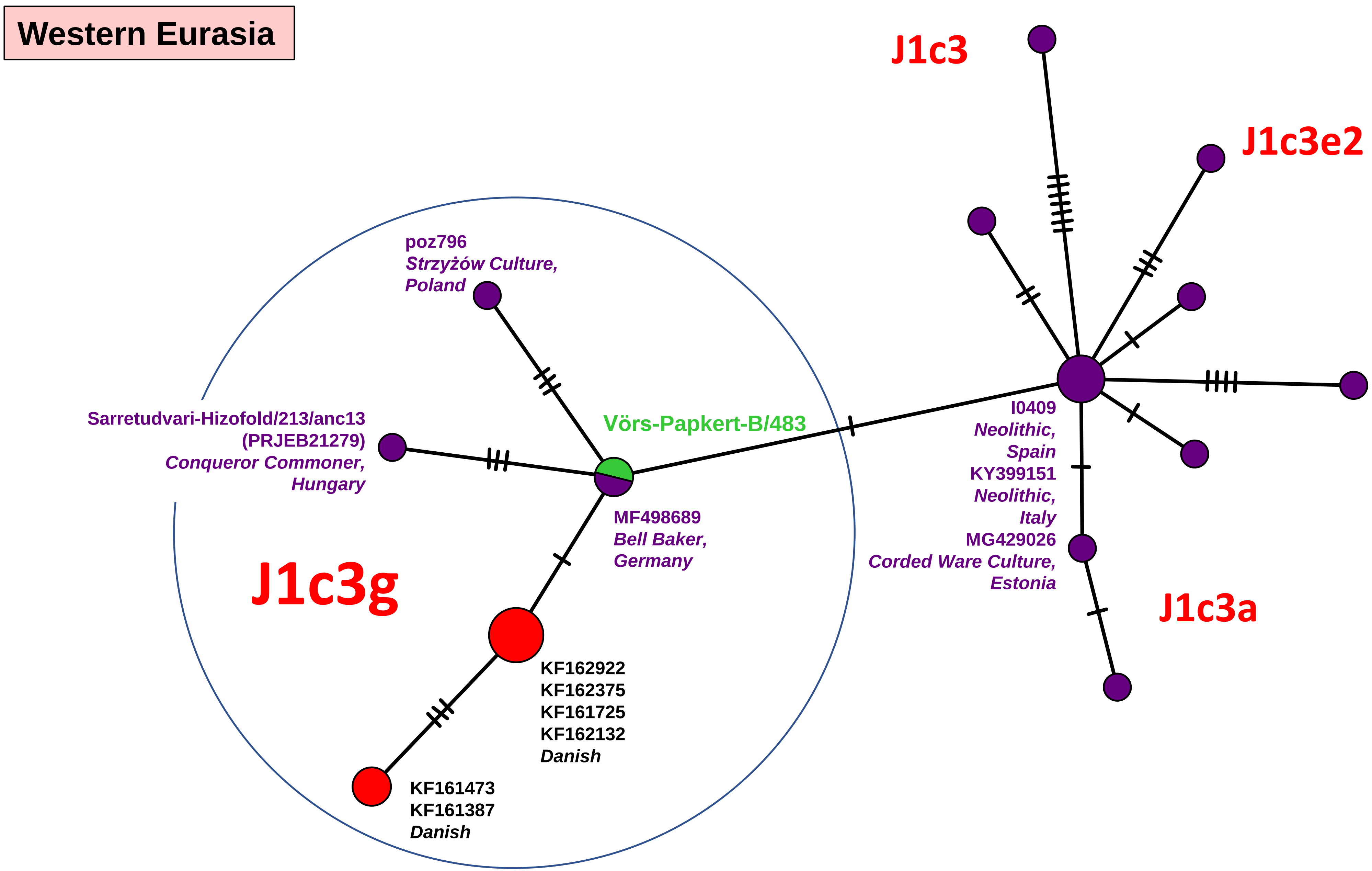
Western Eurasia
J1c3
J1c3e2
poz796
Strzyżów Culture,
Poland
Sarretudvari-Hizofold/213/anc13
(PRJEB21279)
Conqueror Commoner,
Hungary
Vörs-Papkert-B/483
I0409
Neolithic,
Spain
KY399151
Neolithic,
Italy
MG429026
Corded Ware Culture,
Estonia
MF498689
Bell Baker,
Germany
J1c3g
J1c3a
KF162922
KF162375
KF161725
KF162132
Danish
KF161473
KF161387
Danish

Network 47

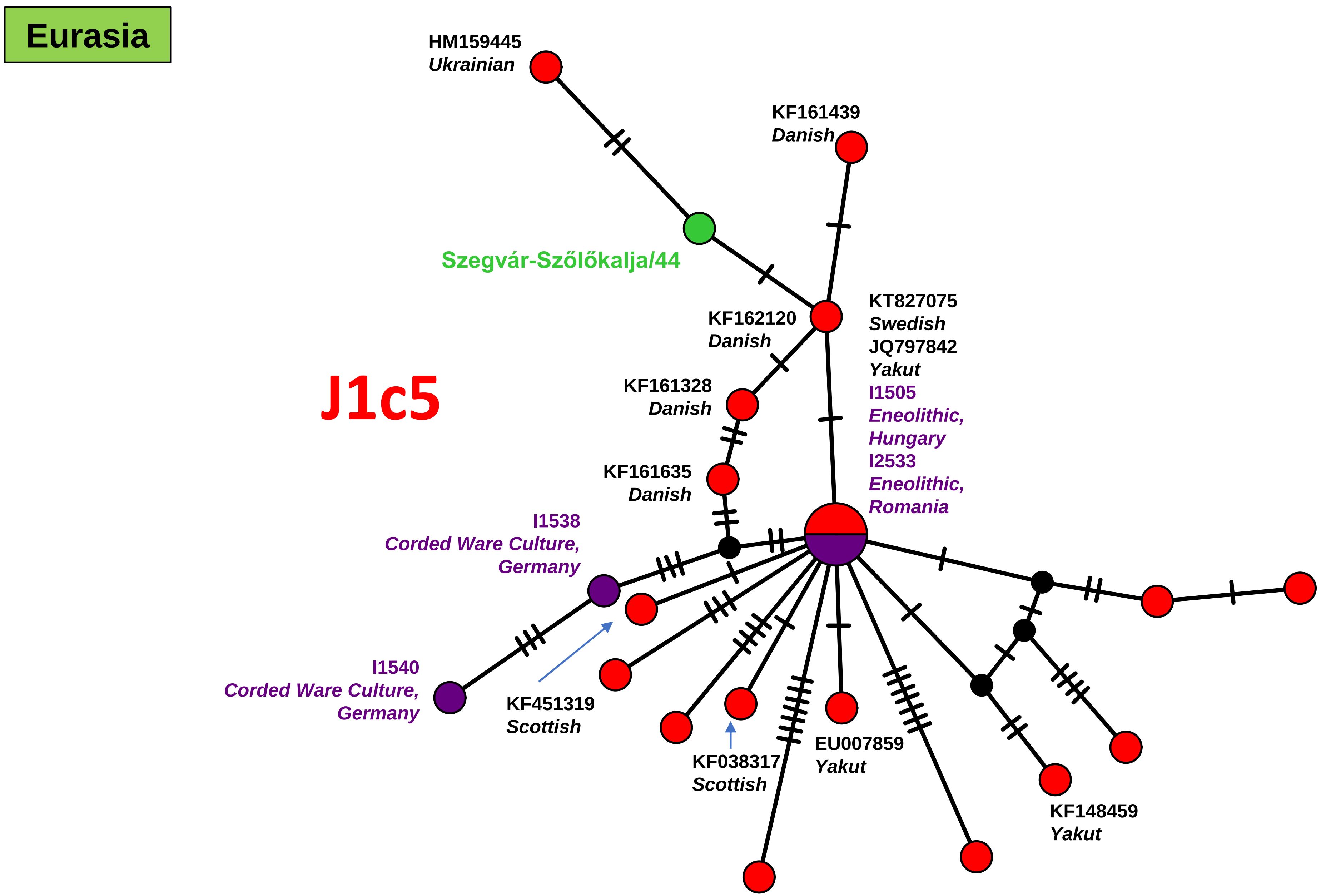
Eurasia
J1c5
HM159445
Ukrainian
KF161439
Danish
Szegvár-Szőlőkalja/44
KF162120
Danish
KT827075
Swedish
JQ797842
Yakut
I1505
Eneolithic,
Hungary
I2533
Eneolithic,
Romania
KF161328
Danish
KF161635
Danish
I1538
Corded Ware Culture,
Germany
I1540
Corded Ware Culture,
Germany
KF451319
Scottish
KF038317
Scottish
EU007859
Yakut
KF148459
Yakut

Network 48

Western Eurasia

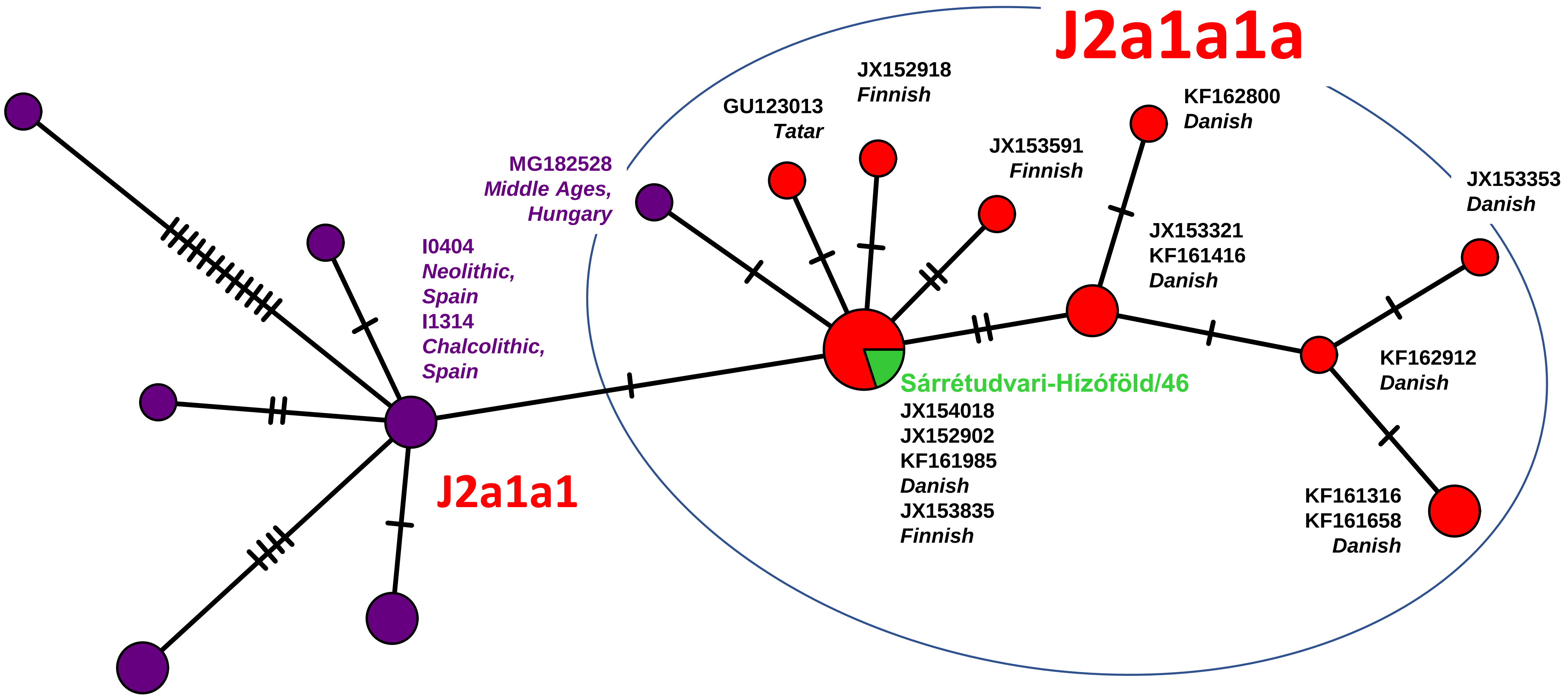


Network 49

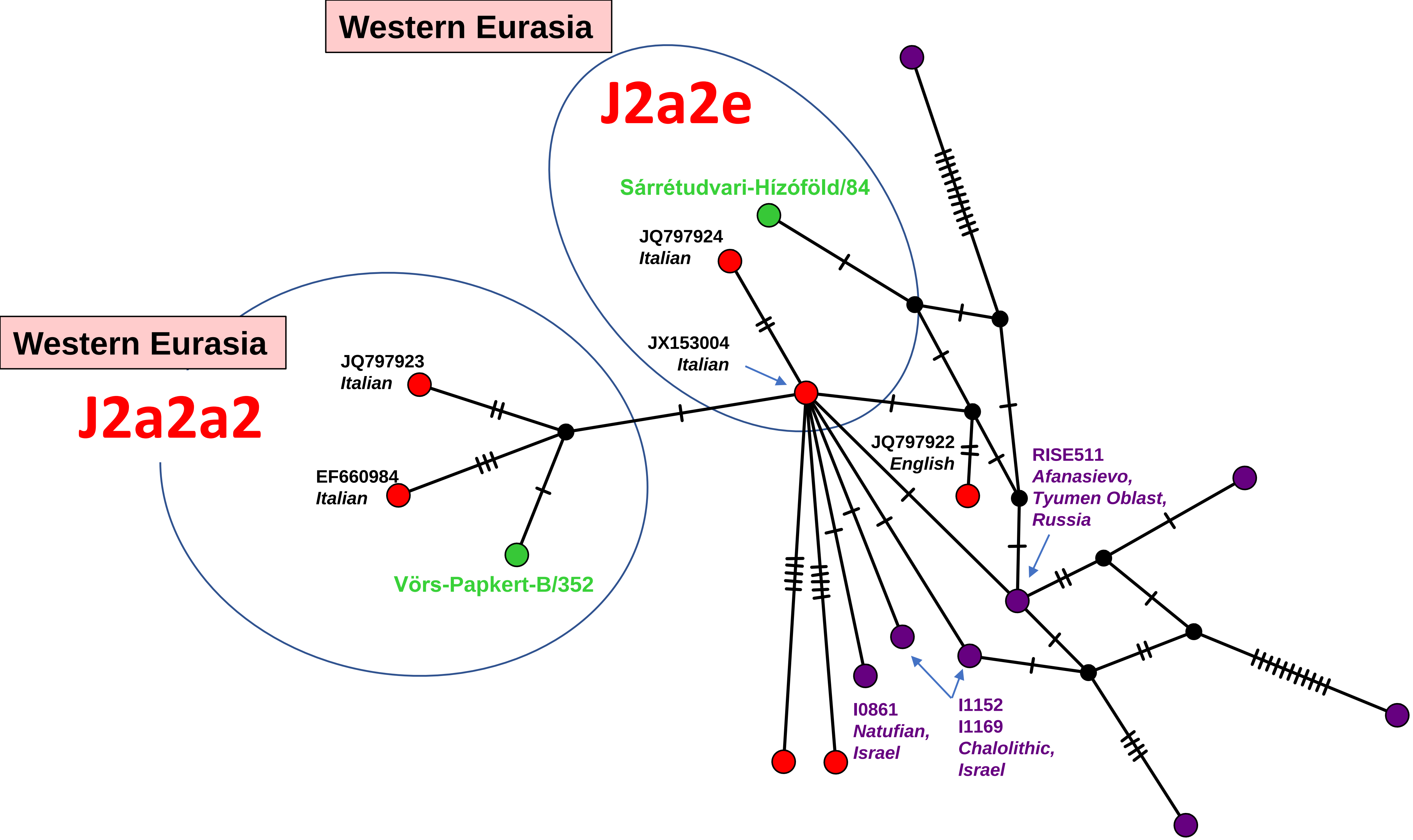
Western Eurasia
J2a2e
Sárrétudvari-Hízóföld/84
JQ797924
Italian
JX153004
Italian
Western Eurasia
J2a2a2
JQ797923
Italian
EF660984
Italian
Vörs-Papkert-B/352
JQ797922
English
RISE511
Afanasievo,
Tyumen Oblast,
Russia
I0861
Natufian,
Israel
I1152
I1169
Chalolithic,
Israel

Network 50

Eurasia

J2b1a

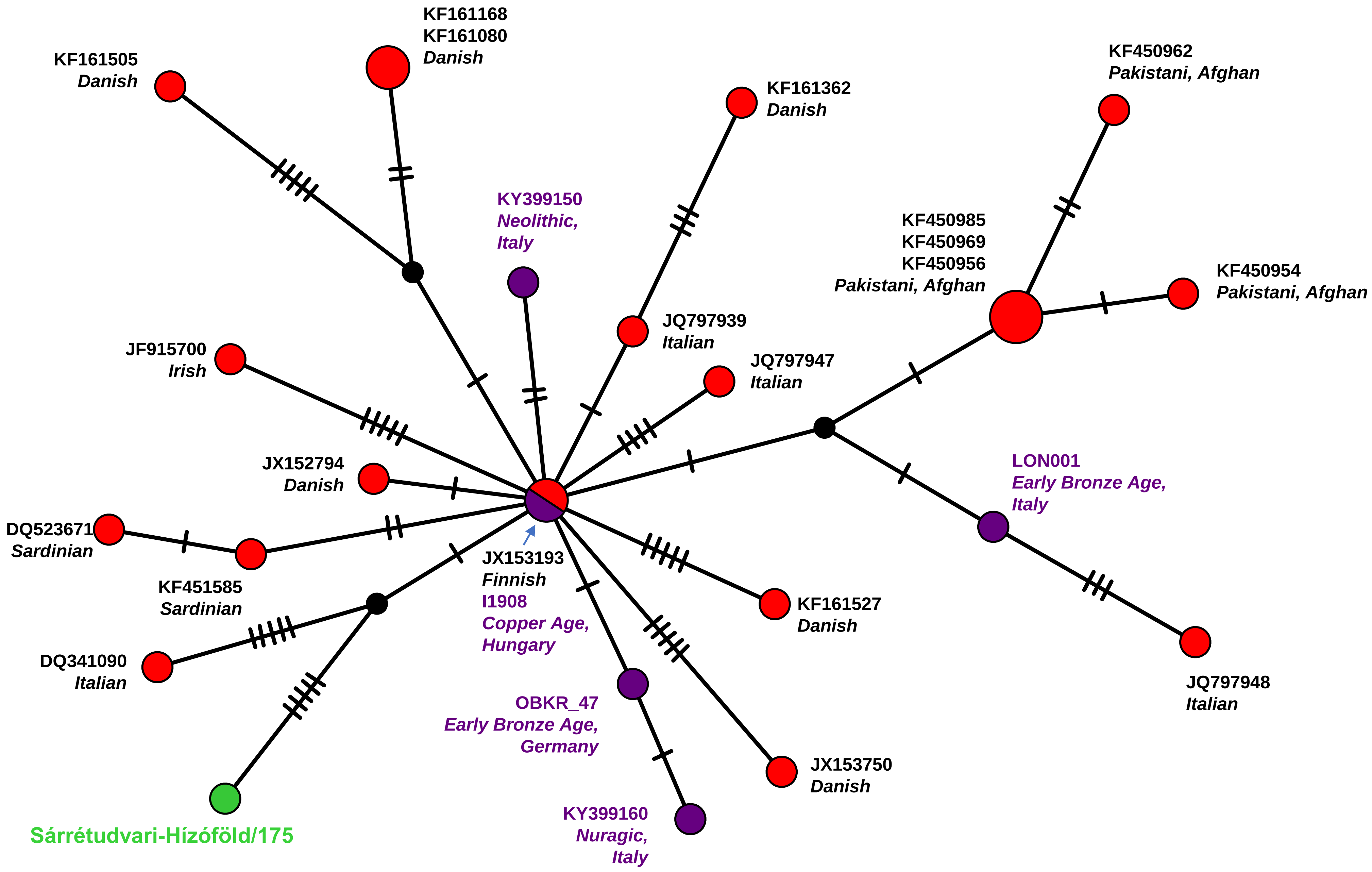


Network 51

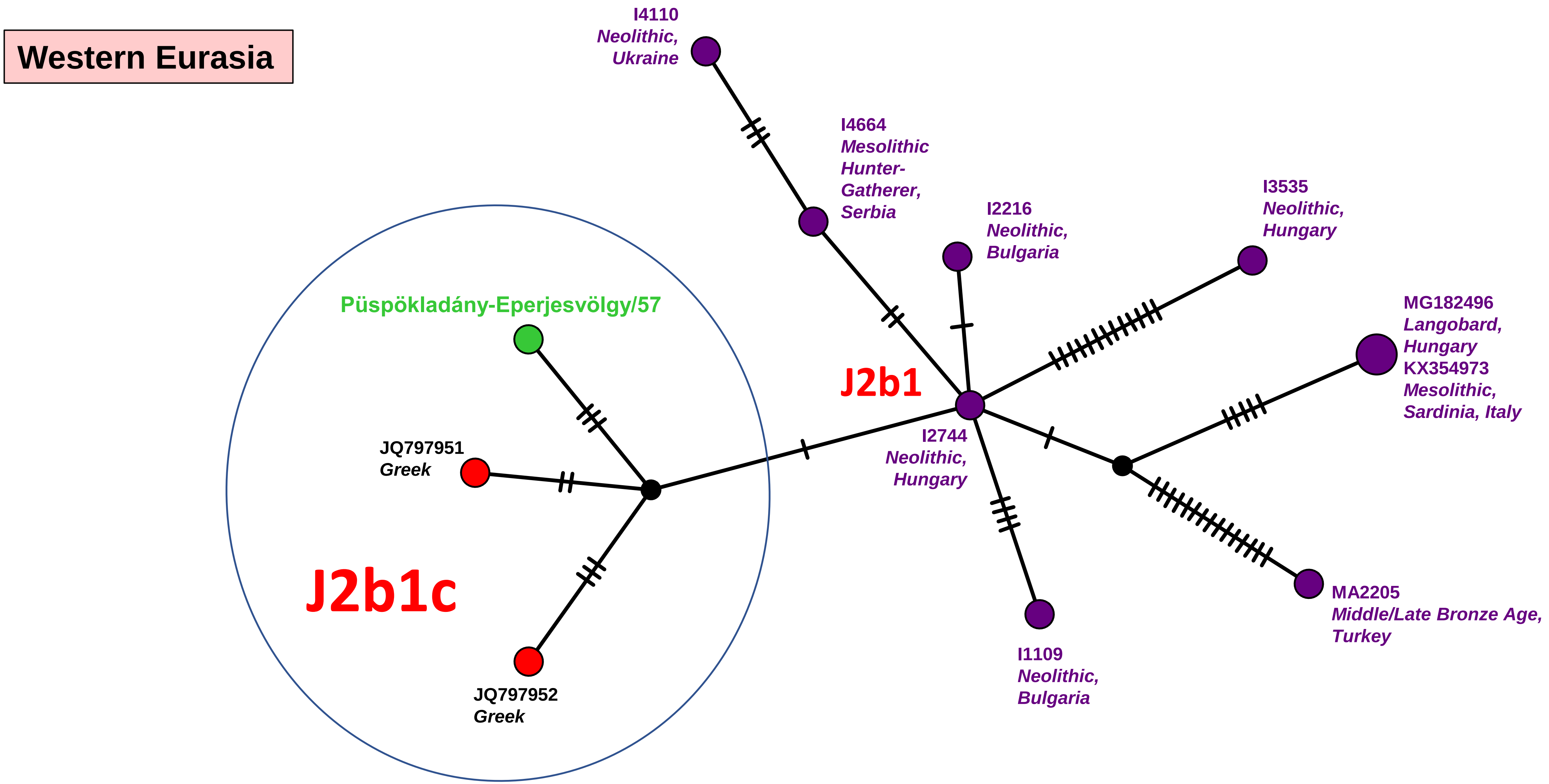

Western Eurasia
I4110
Neolithic,
Ukraine
I4664
Mesolithic
Hunter-
Gatherer,
Serbia
I2216
Neolithic,
Bulgaria
I3535
Neolithic,
Hungary
MG182496
Langobard,
Hungary
KX354973
Mesolithic,
Sardinia, Italy
Püspökladány-Eperjesvölgy/57
J2b1
JQ797951
Greek
I2744
Neolithic,
Hungary
J2b1c
JQ797952
Greek
I1109
Neolithic,
Bulgaria
MA2205
Middle/Late Bronze Age,
Turkey


Network 52

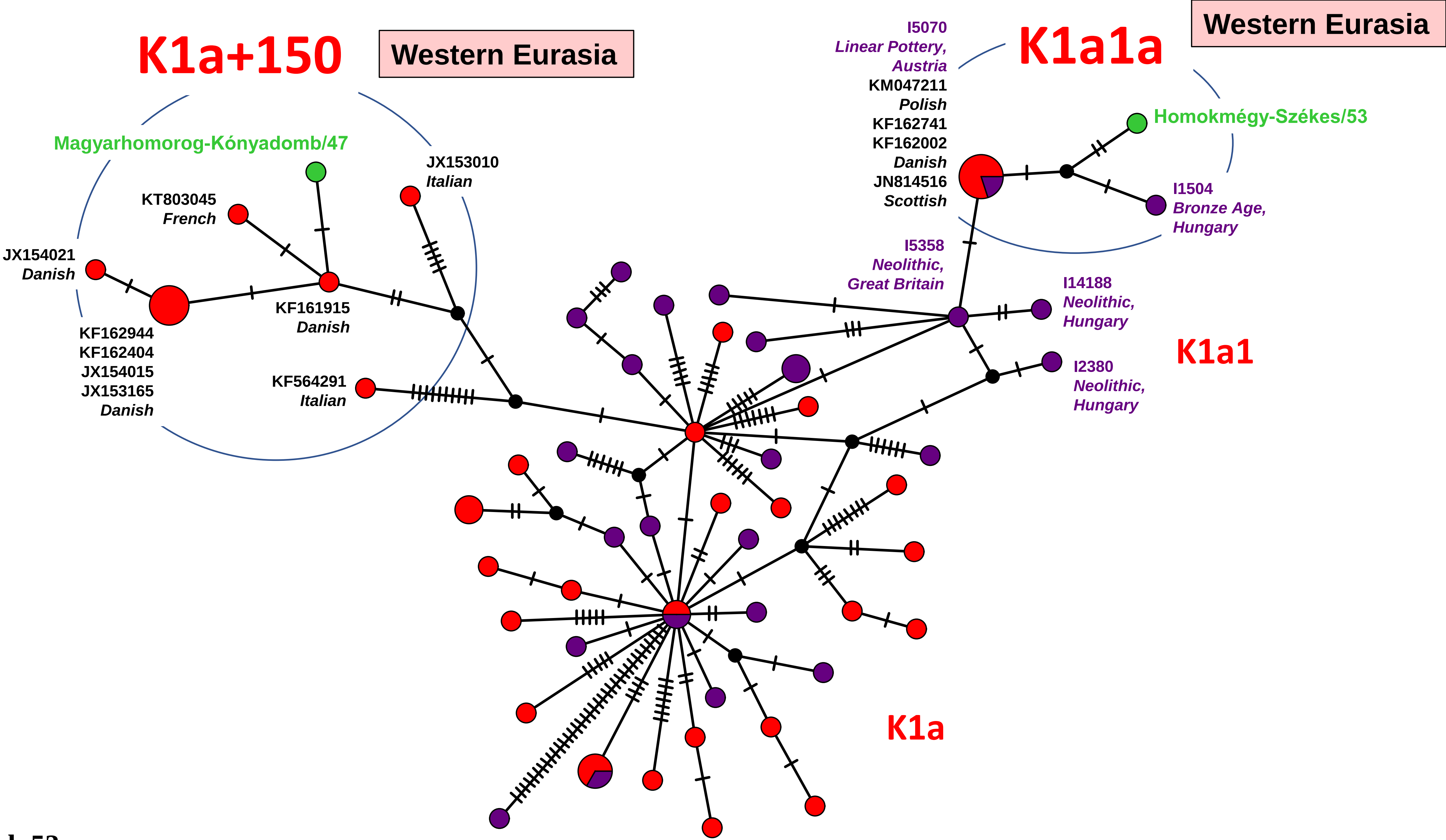
K1a+150
Western Eurasia
Magyarhomorog-Kónyadomb/47
JX153010
Italian
KT803045
French
JX154021
Danish
KF161915
Danish
KF162944
KF162404
JX154015
JX153165
Danish
KF564291
Italian
I5070
Linear Pottery,
Austria
KM047211
Polish
KF162741
KF162002
Danish
JN814516
Scottish
K1a1a
Western Eurasia
Homokmégy-Székes/53
I1504
Bronze Age,
Hungary
I5358
Neolithic,
Great Britain
I14188
Neolithic,
Hungary
K1a1
I2380
Neolithic,
Hungary
K1a

Network 53

Western Eurasia

# K1a1b1

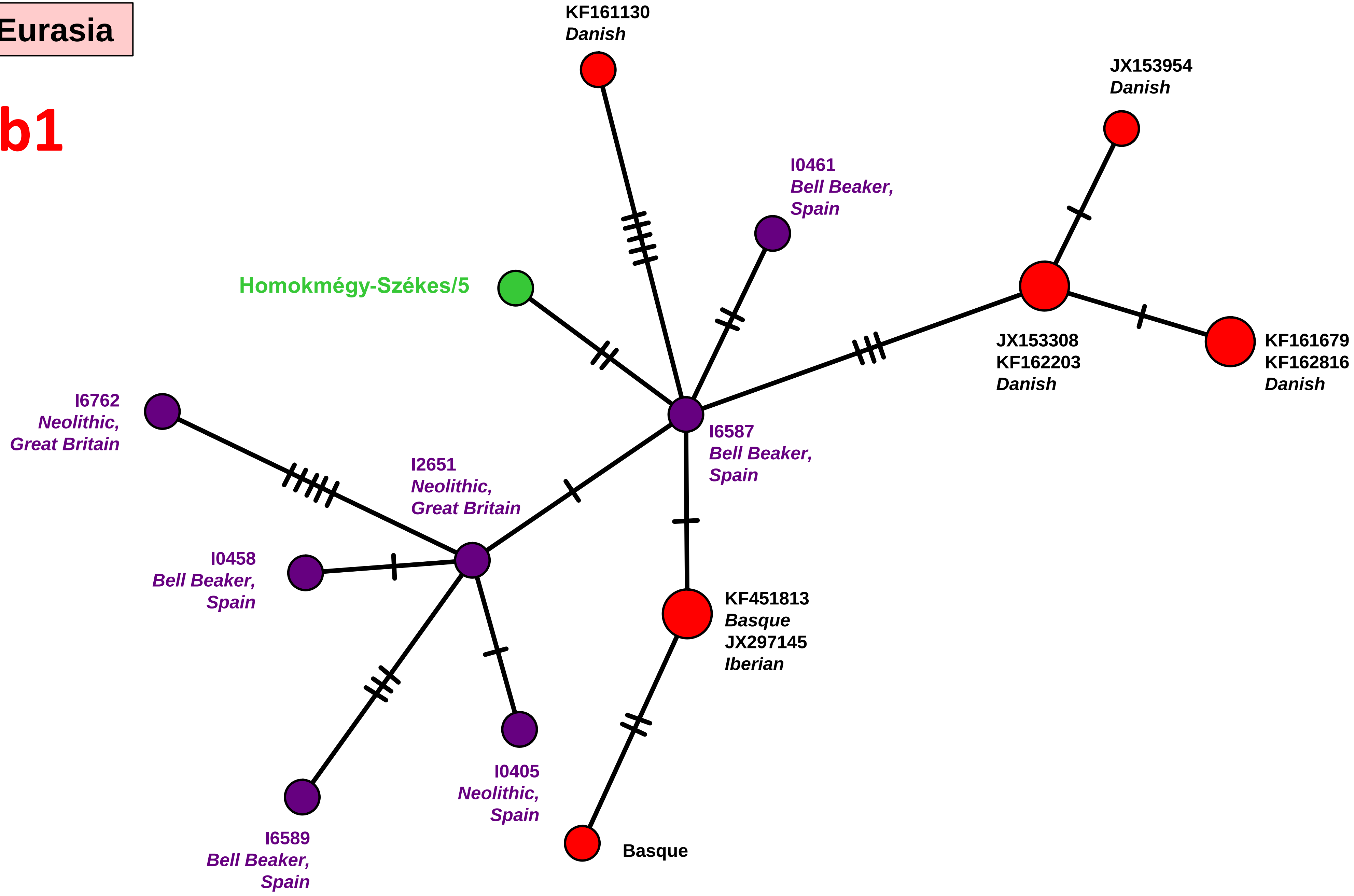


Network 54

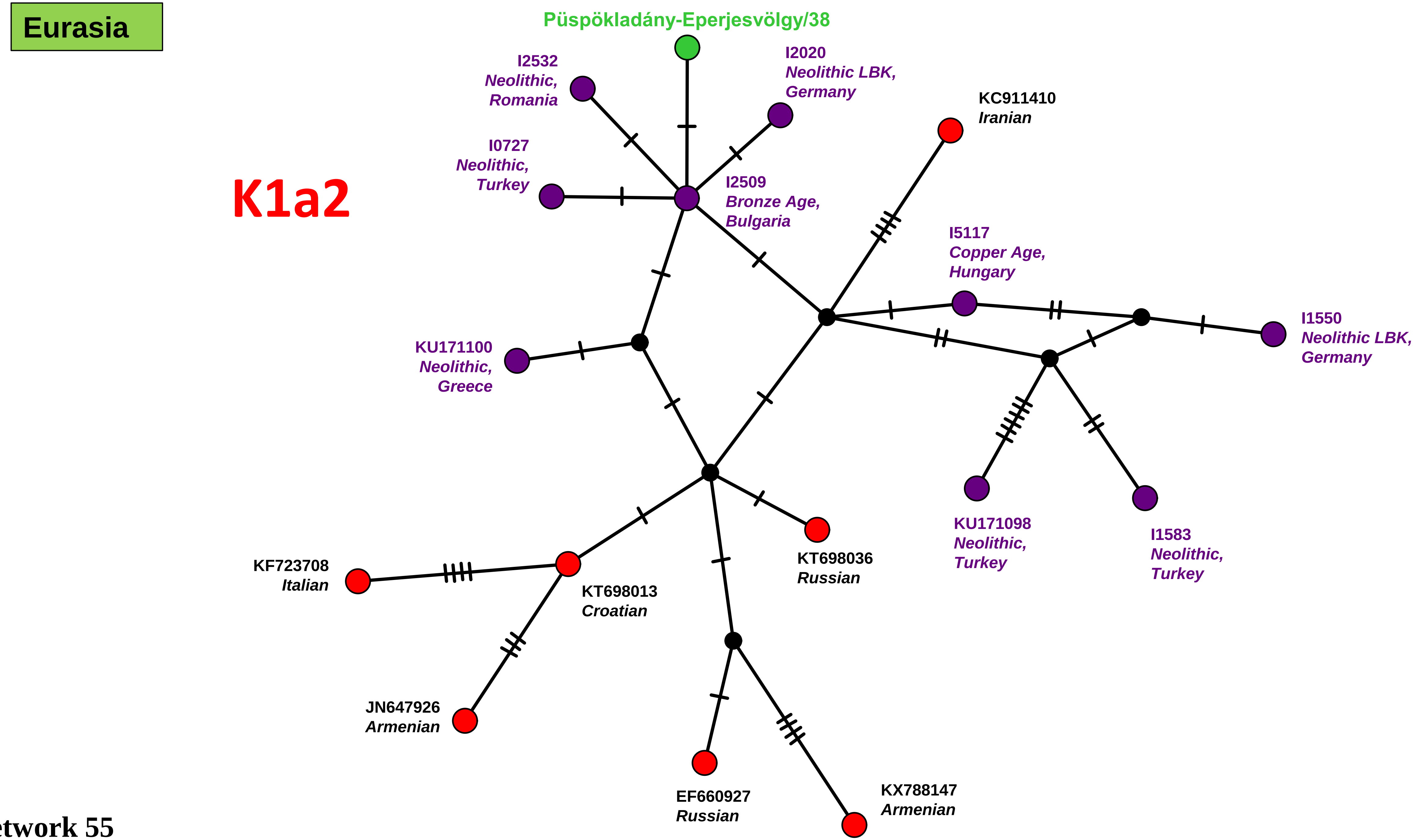
Eurasia
K1a2
Püspökladány-Eperjesvölgy/38
I2532
Neolithic,
Romania
I2020
Neolithic LBK,
Germany
KC911410
Iranian
I0727
Neolithic,
Turkey
I2509
Bronze Age,
Bulgaria
I5117
Copper Age,
Hungary
KU171100
Neolithic,
Greece
I1550
Neolithic LBK,
Germany
KU171098
Neolithic,
Turkey
I1583
Neolithic,
Turkey
KT698036
Russian
KF723708
Italian
KT698013
Croatian
JN647926
Armenian
EF660927
Russian
KX788147
Armenian

Network 55

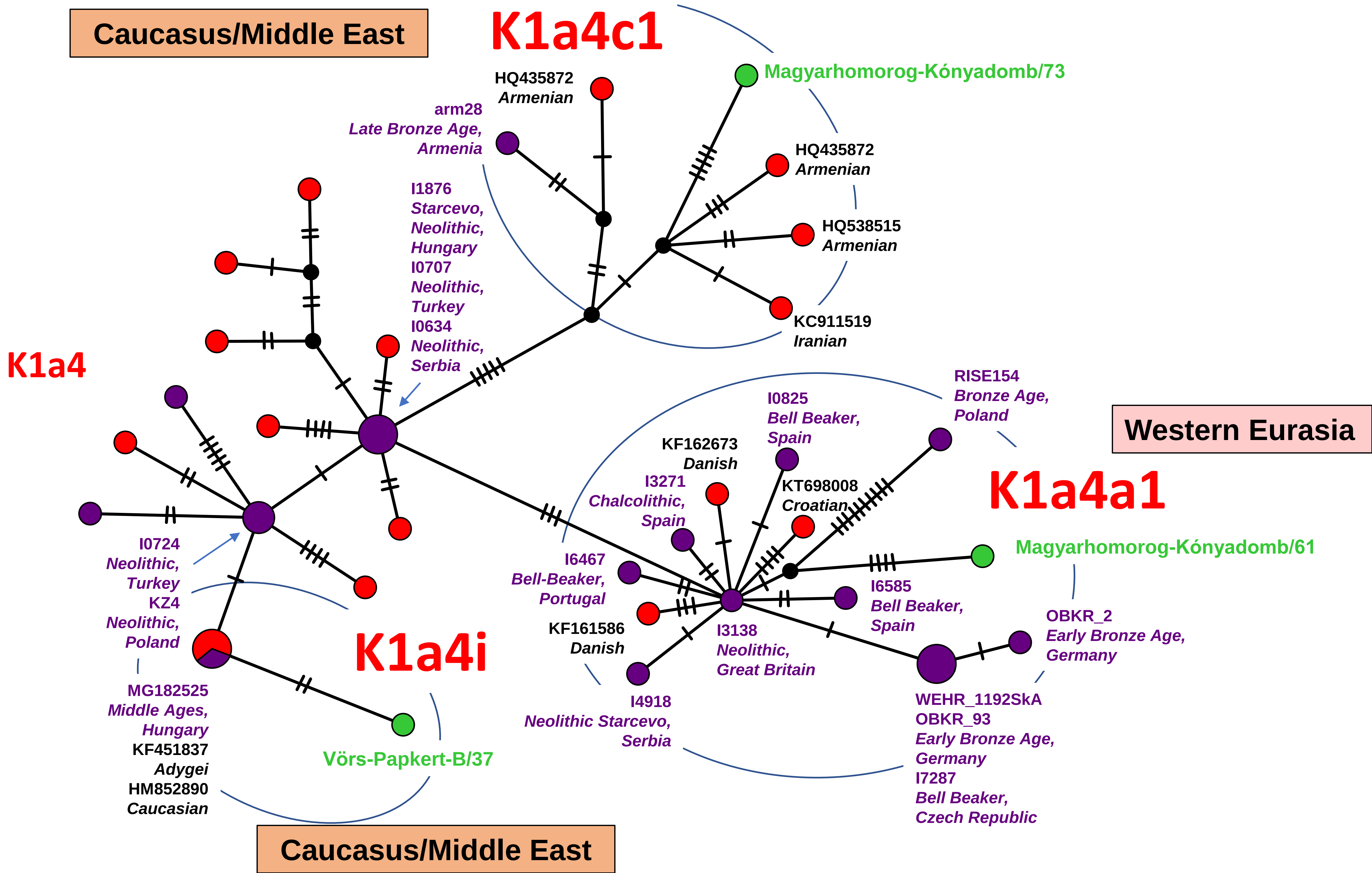
Caucasus/Middle East
K1a4c1
Magyarhomorog-Kónyadomb/73
HQ435872
Armenian
arm28
Late Bronze Age,
Armenia
HQ435872
Armenian
HQ538515
Armenian
KC911519
Iranian
I1876
Starcevo,
Neolithic,
Hungary
I0707
Neolithic,
Turkey
I0634
Neolithic,
Serbia
K1a4
RISE154
Bronze Age,
Poland
I0825
Bell Beaker,
Spain
Western Eurasia
KF162673
Danish
I3271
Chalcolithic,
Spain
KT698008
Croatian
K1a4a1
Magyarhomorog-Kónyadomb/61
I0724
Neolithic,
Turkey
KZ4
Neolithic,
Poland
I6467
Bell-Beaker,
Portugal
I6585
Bell Beaker,
Spain
OBKR_2
Early Bronze Age,
Germany
K1a4i
KF161586
Danish
I3138
Neolithic,
Great Britain
MG182525
Middle Ages,
Hungary
KF451837
Adygei
HM852890
Caucasian
Vörs-Papkert-B/37
I4918
Neolithic Starcevo,
Serbia
WEHR_1192SkA
OBKR_93
Early Bronze Age,
Germany
I7287
Bell Beaker,
Czech Republic
Caucasus/Middle East

Network 56

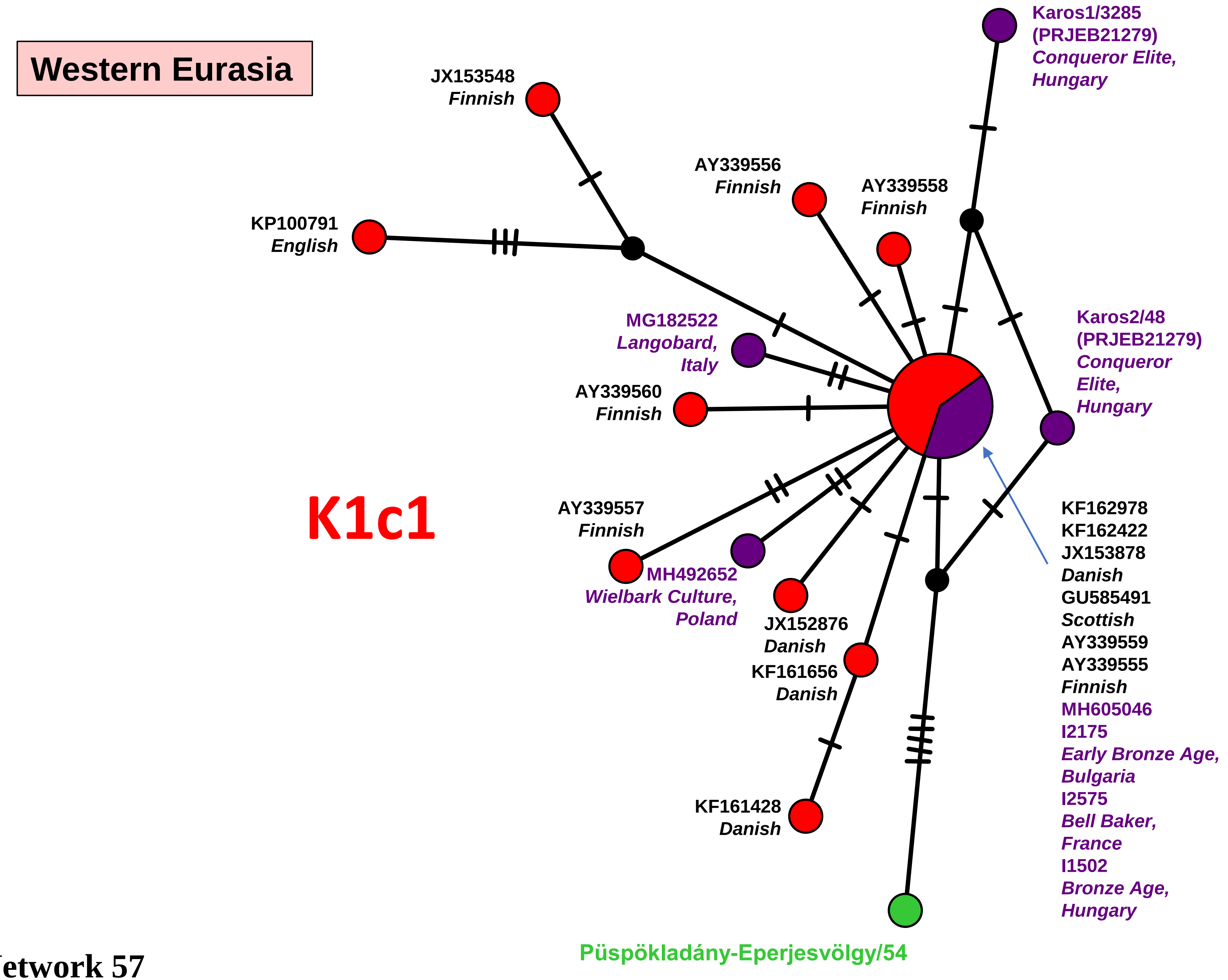
Western Eurasia
K1c1
JX153548
Finnish
KP100791
English
AY339556
Finnish
AY339558
Finnish
Karos1/3285
(PRJEB21279)
Conqueror Elite,
Hungary
MG182522
Langobard,
Italy
AY339560
Finnish
Karos2/48
(PRJEB21279)
Conqueror
Elite,
Hungary
AY339557
Finnish
MH492652
Wielbark Culture,
Poland
JX152876
Danish
KF161656
Danish
KF161428
Danish
KF162978
KF162422
JX153878
Danish
GU585491
Scottish
AY339559
AY339555
Finnish
MH605046
I2175
Early Bronze Age,
Bulgaria
I2575
Bell Baker,
France
I1502
Bronze Age,
Hungary
Püspökladány-Eperjesvölgy/54

Network 57

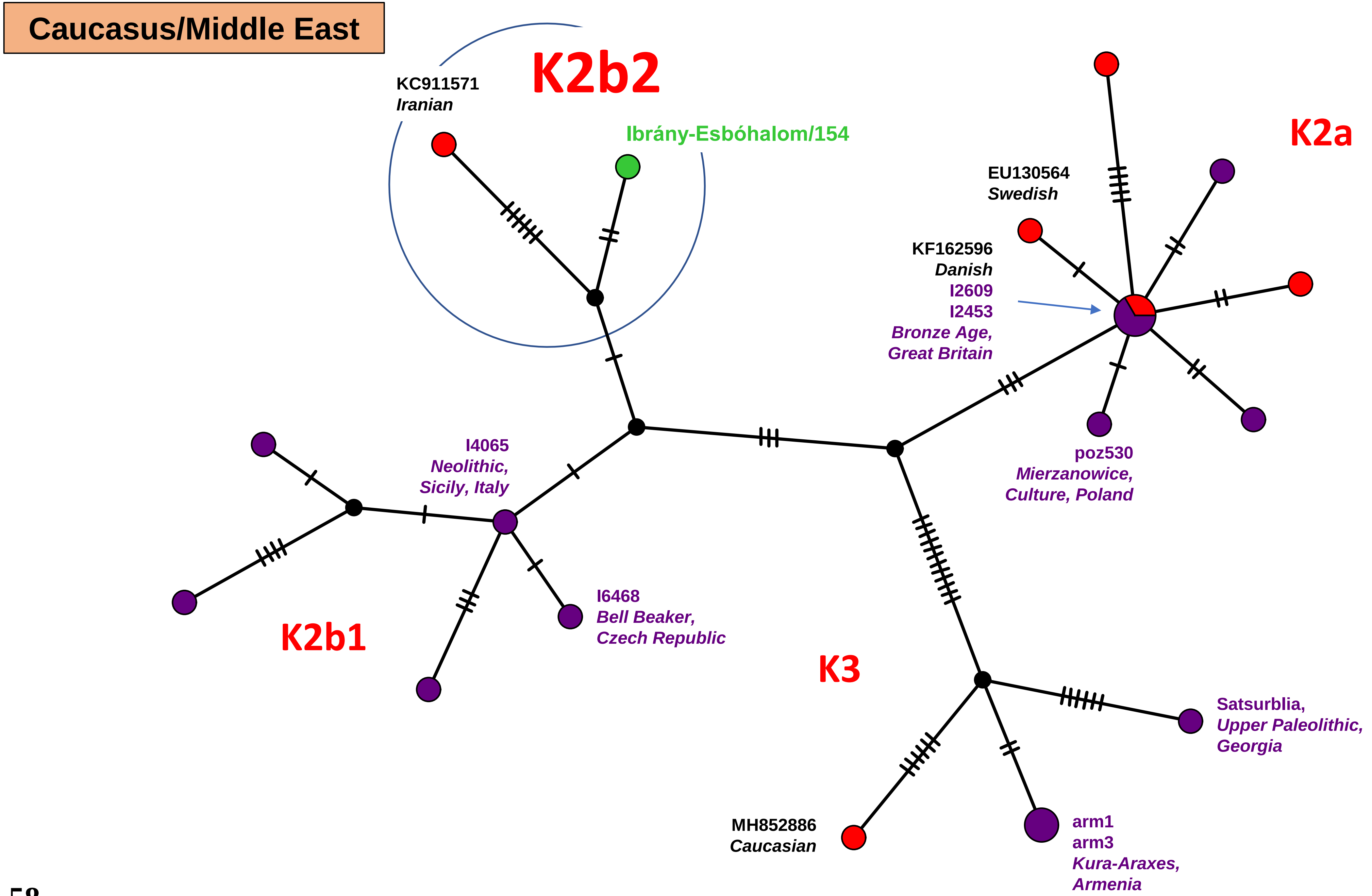

Caucasus/Middle East
K2b2
KC911571
Iranian
Ibrány-Esbóhalom/154
K2a
EU130564
Swedish
KF162596
Danish
I2609
I2453
Bronze Age,
Great Britain
I4065
Neolithic,
Sicily, Italy
poz530
Mierzanowice,
Culture, Poland
I6468
Bell Beaker,
Czech Republic
K2b1
K3
Satsurblia,
Upper Paleolithic,
Georgia
MH852886
Caucasian
arm1
arm3
Kura-Araxes,
Armenia


Network 58

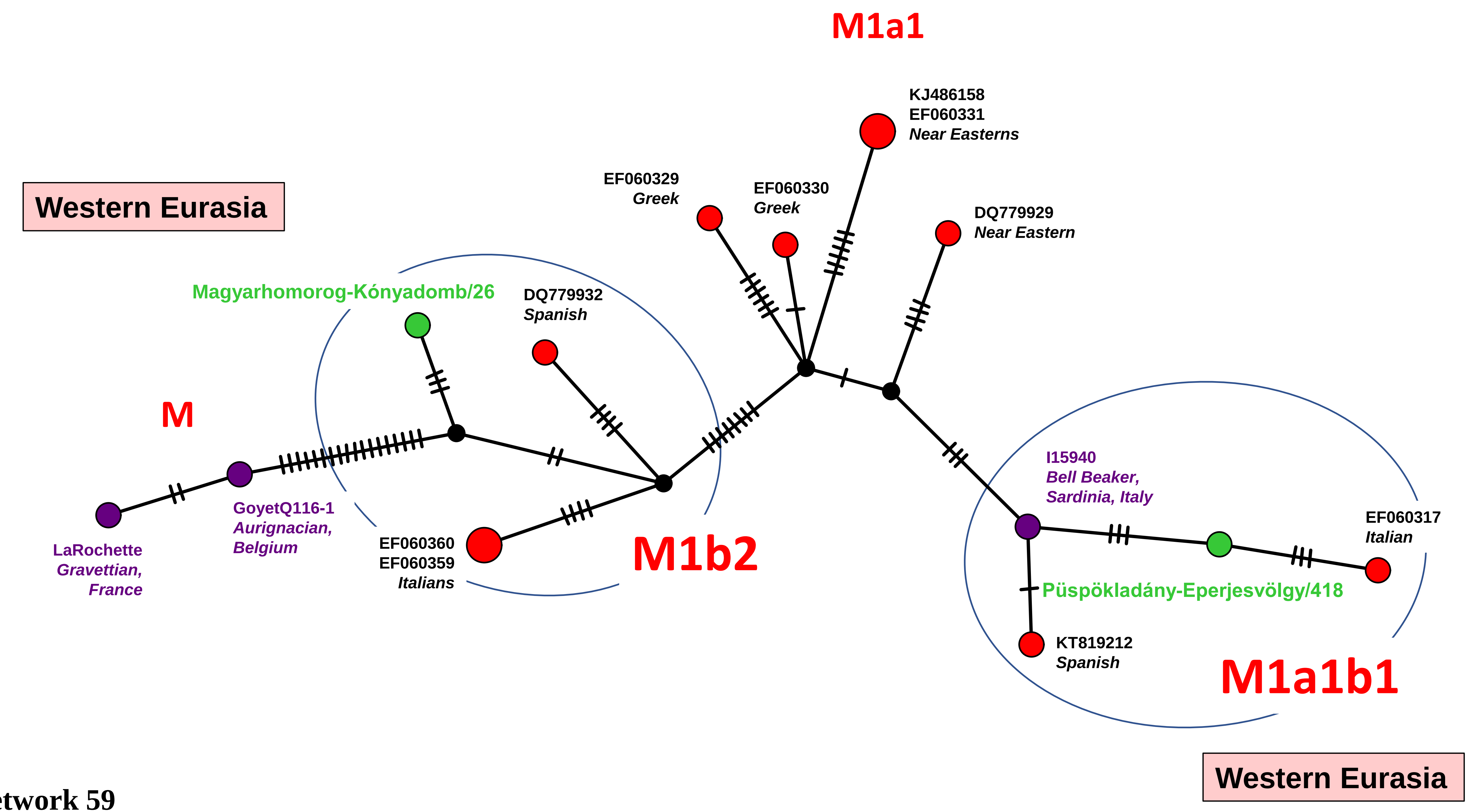
M1a1
KJ486158
EF060331
Near Easterns
Western Eurasia
EF060329
Greek
EF060330
Greek
DQ779929
Near Eastern
Magyarhomorog-Kónyadomb/26
DQ779932
Spanish
M
I15940
Bell Beaker,
Sardinia, Italy
GoyetQ116-1
Aurignacian,
Belgium
EF060317
Italian
LaRochette
Gravettian,
France
EF060360
EF060359
Italians
M1b2
Püspökladány-Eperjesvölgy/418
KT819212
Spanish
M1a1b1
Western Eurasia

Network 59

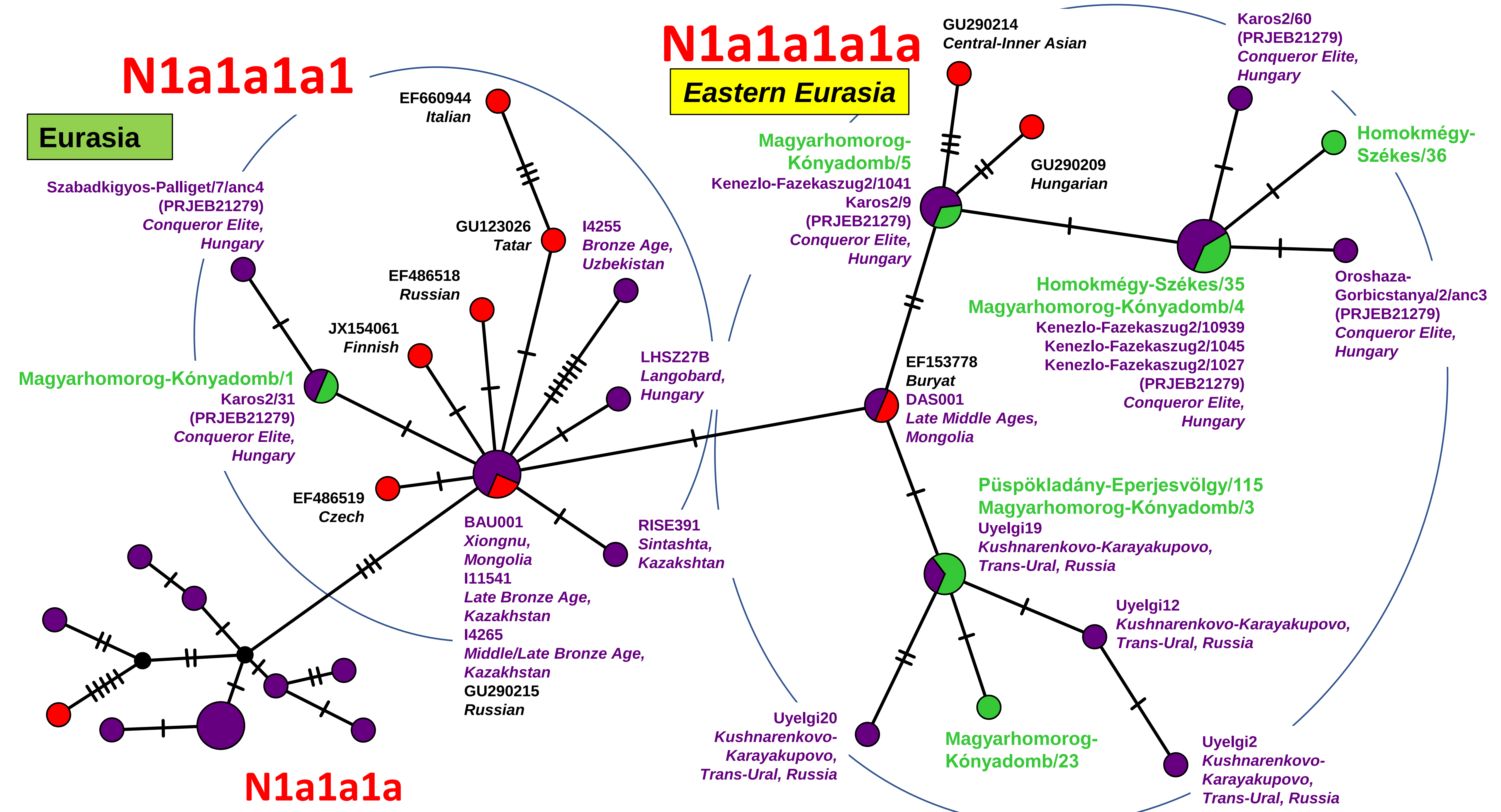
N1a1a1a1
Eurasia
N1a1a1a1a
Eastern Eurasia
N1a1a1a
EF660944
Italian
GU123026
Tatar
I4255
Bronze Age,
Uzbekistan
EF486518
Russian
JX154061
Finnish
Szabadkigyos-Palliget/7/anc4
(PRJEB21279)
Conqueror Elite,
Hungary
Magyarhomorog-Kónyadomb/1
Karos2/31
(PRJEB21279)
Conqueror Elite,
Hungary
LHSZ27B
Langobard,
Hungary
EF486519
Czech
BAU001
Xiongnu,
Mongolia
I11541
Late Bronze Age,
Kazakhstan
I4265
Middle/Late Bronze Age,
Kazakhstan
GU290215
Russian
RISE391
Sintashta,
Kazakhstan
GU290214
Central-Inner Asian
Karos2/60
(PRJEB21279)
Conqueror Elite,
Hungary
Magyarhomorog-
Kónyadomb/5
Kenezlo-Fazekaszug2/1041
Karos2/9
(PRJEB21279)
Conqueror Elite,
Hungary
GU290209
Hungarian
Homokmégy-
Székes/36
Homokmégy-Székes/35
Magyarhomorog-Kónyadomb/4
Kenezlo-Fazekaszug2/10939
Kenezlo-Fazekaszug2/1045
Kenezlo-Fazekaszug2/1027
(PRJEB21279)
Conqueror Elite,
Hungary
Oroshaza-
Gorbicstanya/2/anc3
(PRJEB21279)
Conqueror Elite,
Hungary
EF153778
Buryat
DAS001
Late Middle Ages,
Mongolia
Püspökladány-Eperjesvölgy/115
Magyarhomorog-Kónyadomb/3
Uyelgi19
Kushnarenkovo-Karayakupovo,
Trans-Ural, Russia
Uyelgi12
Kushnarenkovo-Karayakupovo,
Trans-Ural, Russia
Uyelgi20
Kushnarenkovo-
Karayakupovo,
Trans-Ural, Russia
Magyarhomorog-
Kónyadomb/23
Uyelgi2
Kushnarenkovo-
Karayakupovo,
Trans-Ural, Russia

Network 60

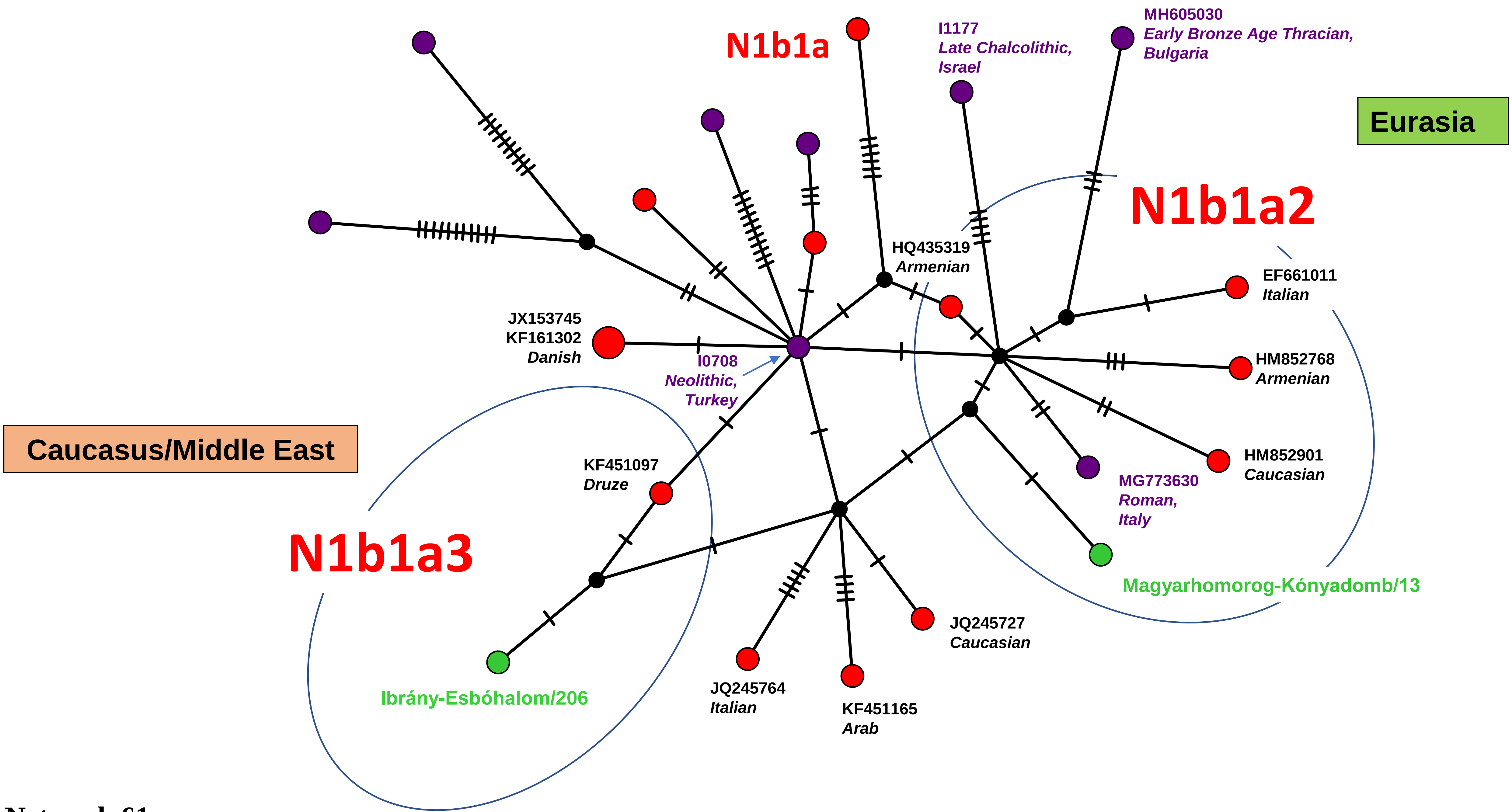

N1b1a
I1177
Late Chalcolithic,
Israel
MH605030
Early Bronze Age Thracian,
Bulgaria
Eurasia
N1b1a2
HQ435319
Armenian
EF661011
Italian
JX153745
KF161302
Danish
I0708
Neolithic,
Turkey
HM852768
Armenian
Caucasus/Middle East
KF451097
Druze
HM852901
Caucasian
MG773630
Roman,
Italy
N1b1a3
Magyarhomorog-Kónyadomb/13
JQ245727
Caucasian
Ibrány-Esbóhalom/206
JQ245764
Italian
KF451165
Arab


Network 61

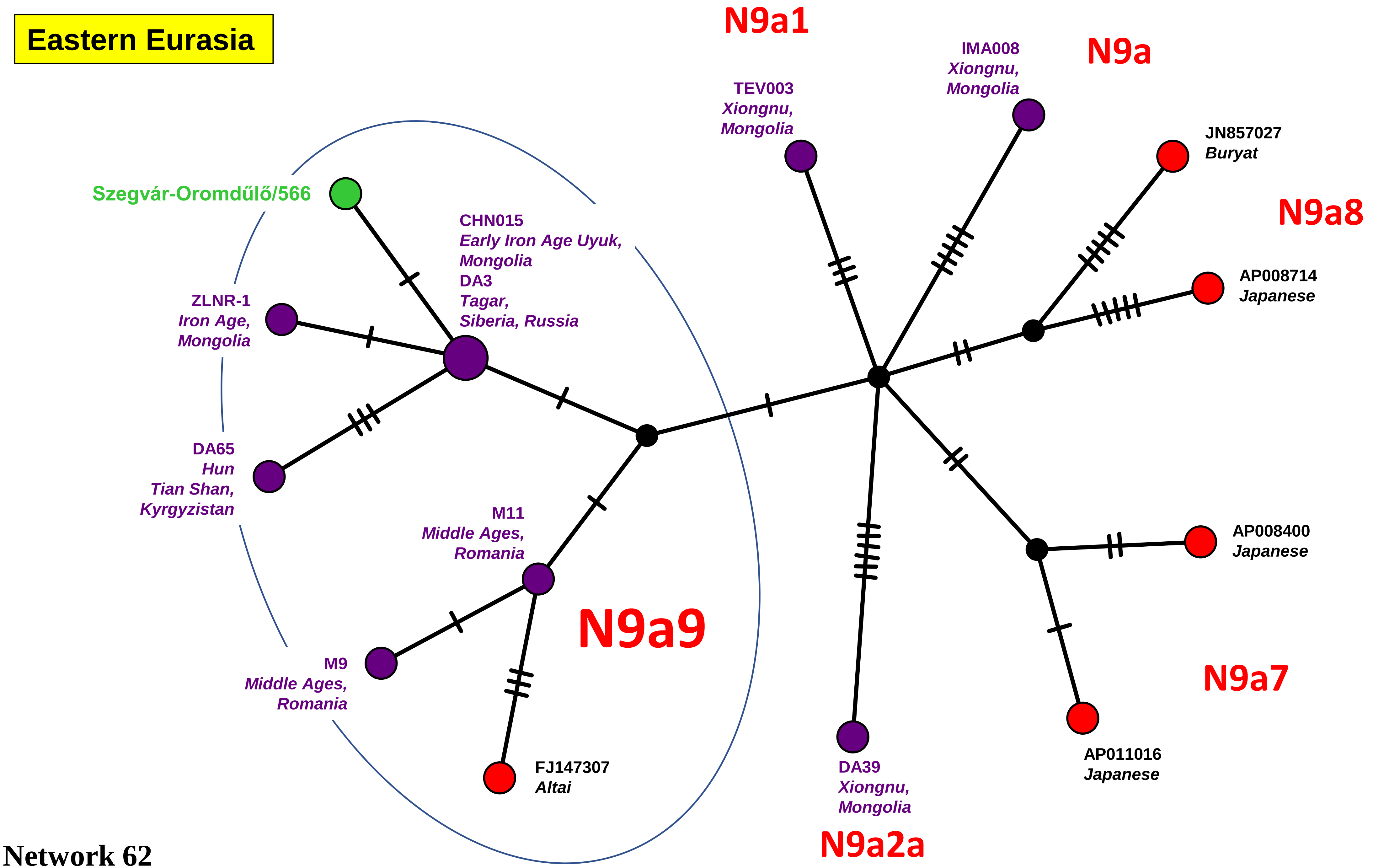

Eastern Eurasia
N9a1
N9a
N9a8
N9a9
N9a7
N9a2a
Szegvár-Oromdűlő/566
CHN015
Early Iron Age Uyuk,
Mongolia
DA3
Tagar,
Siberia, Russia
ZLNR-1
Iron Age,
Mongolia
DA65
Hun
Tian Shan,
Kyrgyzistan
M11
Middle Ages,
Romania
M9
Middle Ages,
Romania
FJ147307
Altai
TEV003
Xiongnu,
Mongolia
IMA008
Xiongnu,
Mongolia
JN857027
Buryat
AP008714
Japanese
AP008400
Japanese
AP011016
Japanese
DA39
Xiongnu,
Mongolia


Network 62

Caucasus/Middle East

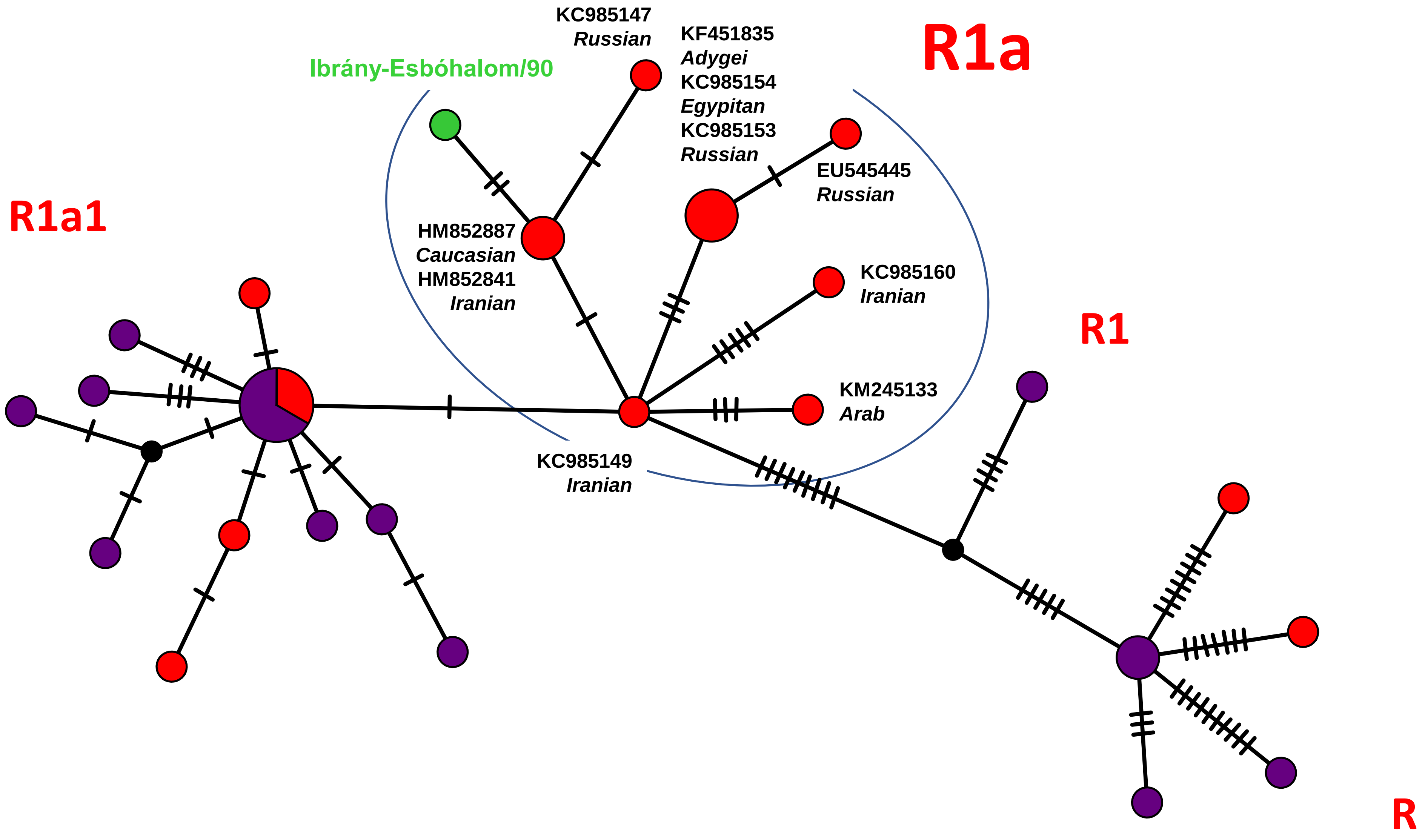


Network 63

**Eurasia**

# T1a1

Central Circle:

Magyarhomorog-Kónyadomb/151

KF162202
KF162064
KF161998
JX153684
JX153402
*Danish*
JN880467
*Irish*
HQ167734
*Ukrainian*
GU122980
*Tatar*
AF382006.
*Spanish*
KF057946
*Norwegian*
JQ797981
*Turkish*
JQ797980
*Caucasian*
JQ797979
*Swedish*
JQ797978
*Baltic*
I0550
*Bronze Age,*
*Germany*
I2105
*Yamnaya,*
*Ukraine*
I4892
*Bronze Age,*
*Czech Republic*
A25
*Late Viking Age,*
*Sweden*
chy002
*Late Sarmatian,*
*Orenburg Region,*
*Russia*
POST_50
*Early Bronze Age*
*Germany*
MG182461
*Langobard,*
*Hungary*
I6797
*Middle/Late Bronze*
*Age,*
*Kazakhstan*
I3769
*Late Bronze Age,*
*Kazakhstan*
I12979
*Iron Age,*
*Pakistan*

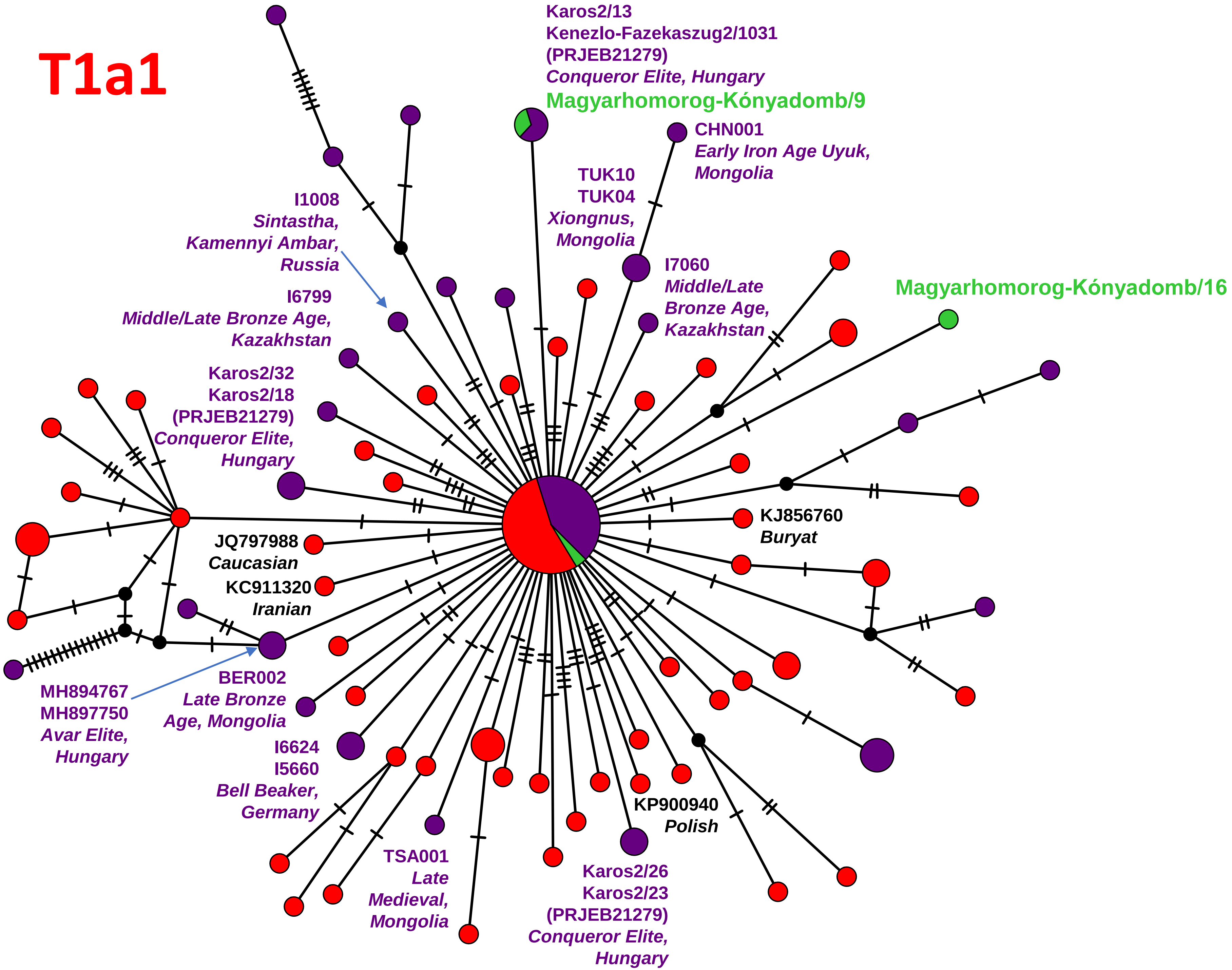


Network 64

Eurasia

# T1a1b

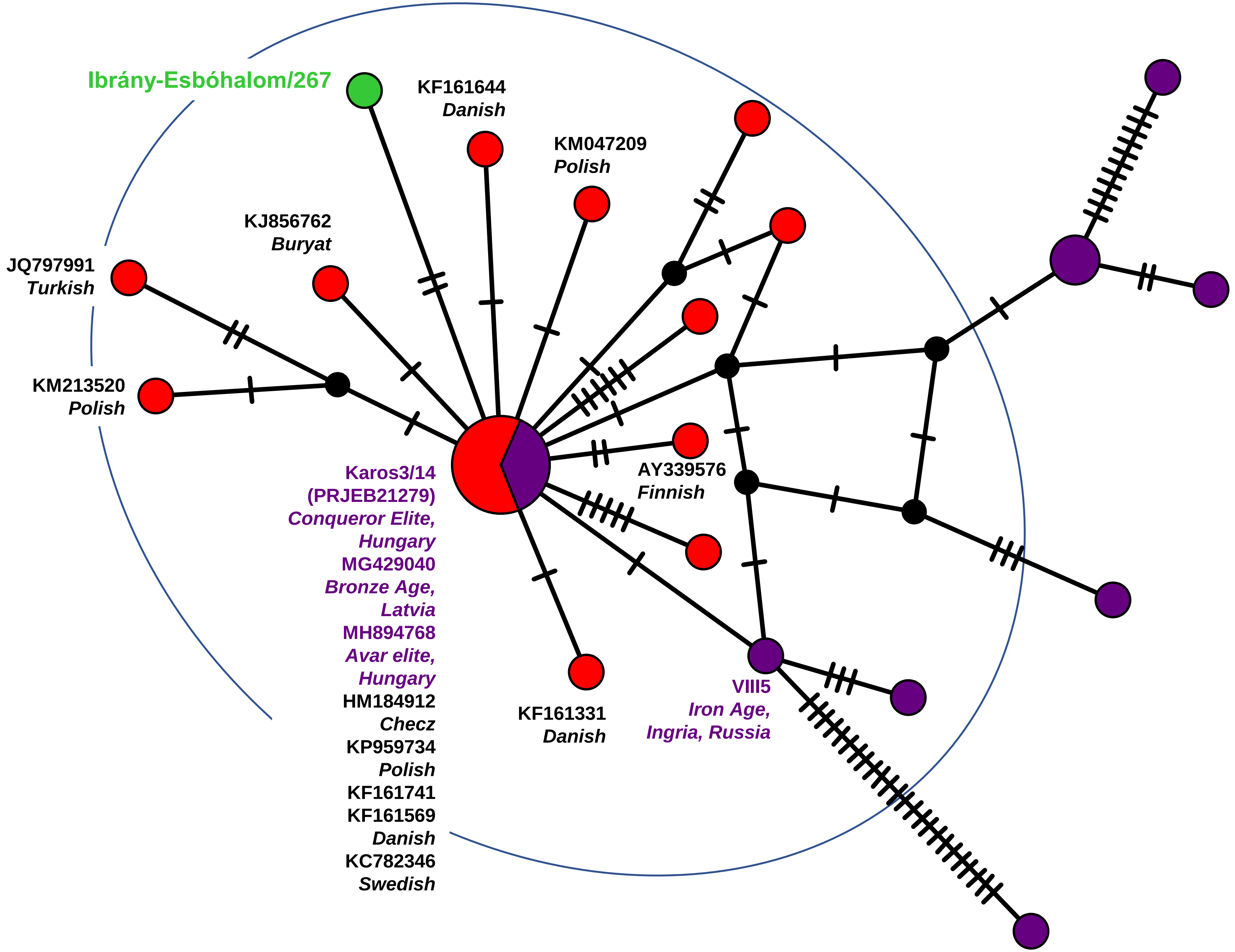


Network 65

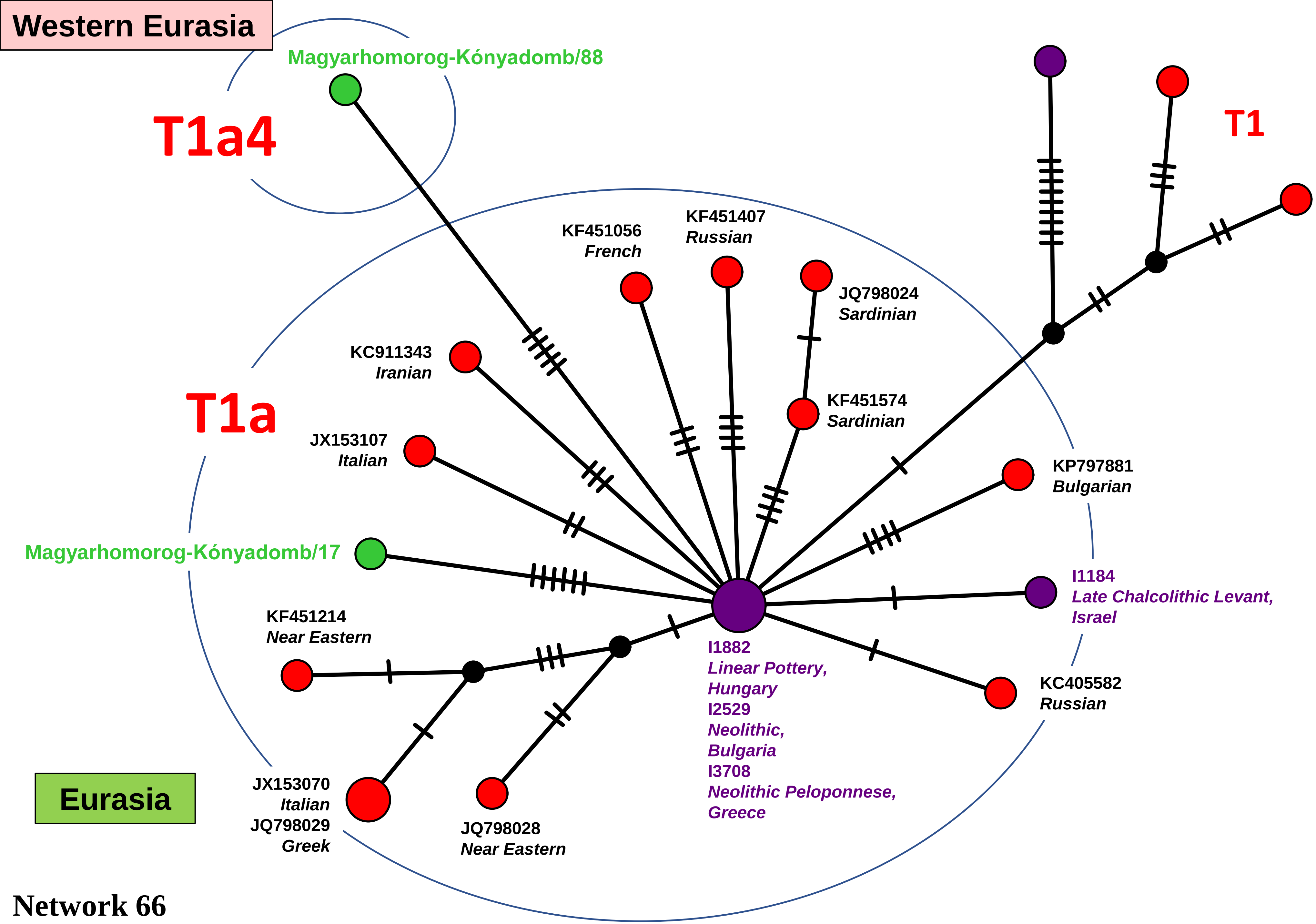
Western Eurasia
Magyarhomorog-Kónyadomb/88
T1a4
T1
KF451056
French
KF451407
Russian
JQ798024
Sardinian
KC911343
Iranian
T1a
KF451574
Sardinian
JX153107
Italian
KP797881
Bulgarian
Magyarhomorog-Kónyadomb/17
I1184
Late Chalcolithic Levant,
Israel
KF451214
Near Eastern
I1882
Linear Pottery,
Hungary
I2529
Neolithic,
Bulgaria
I3708
Neolithic Peloponnese,
Greece
KC405582
Russian
Eurasia
JX153070
Italian
JQ798029
Greek
JQ798028
Near Eastern

Network 66

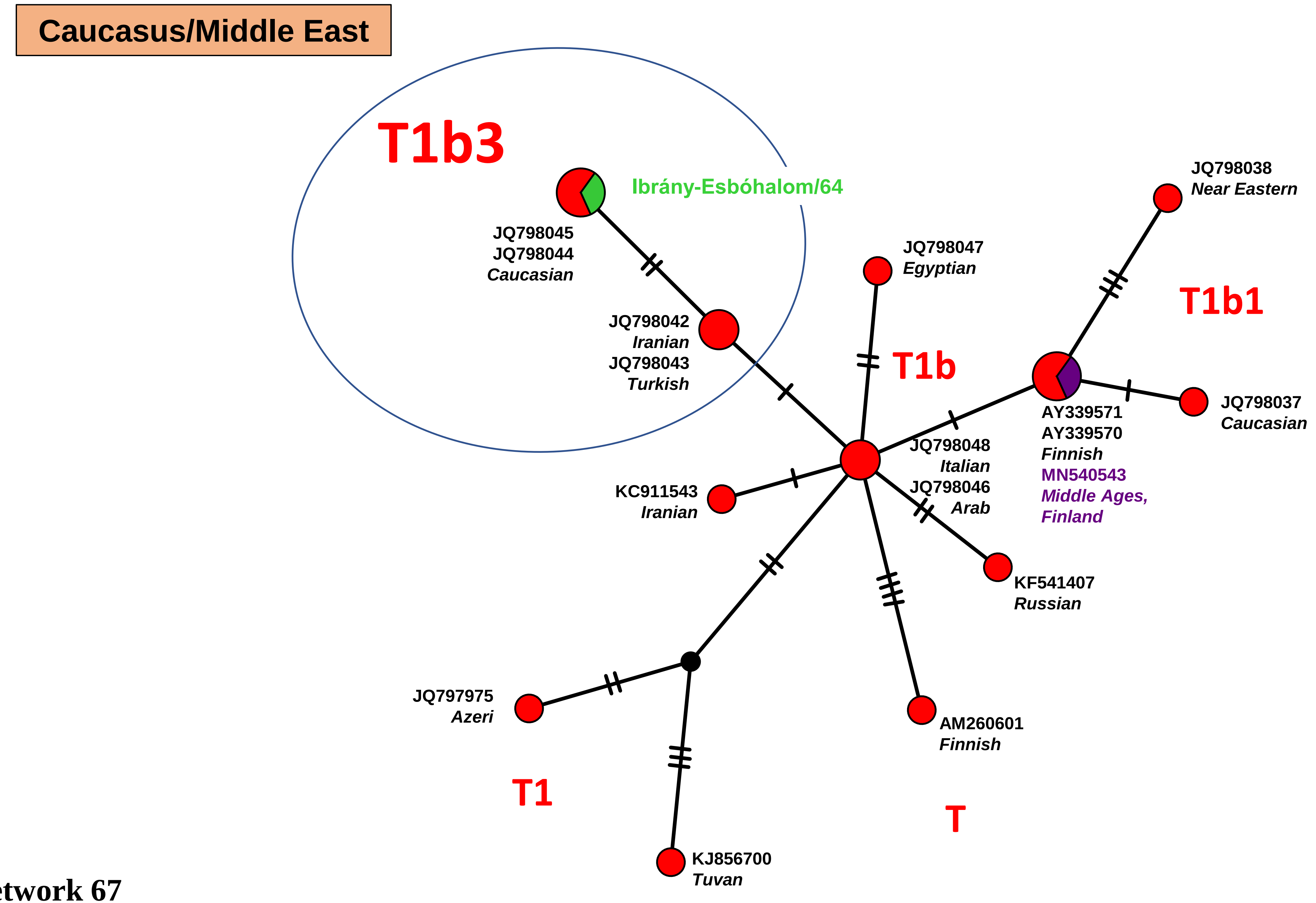
Caucasus/Middle East
T1b3
Ibrány-Esbóhalom/64
JQ798045
JQ798044
Caucasian
JQ798042
Iranian
JQ798043
Turkish
JQ798047
Egyptian
T1b
JQ798038
Near Eastern
T1b1
AY339571
AY339570
Finnish
MN540543
Middle Ages, Finland
JQ798037
Caucasian
JQ798048
Italian
JQ798046
Arab
KC911543
Iranian
KF541407
Russian
JQ797975
Azeri
AM260601
Finnish
T1
T
KJ856700
Tuvan

Network 67

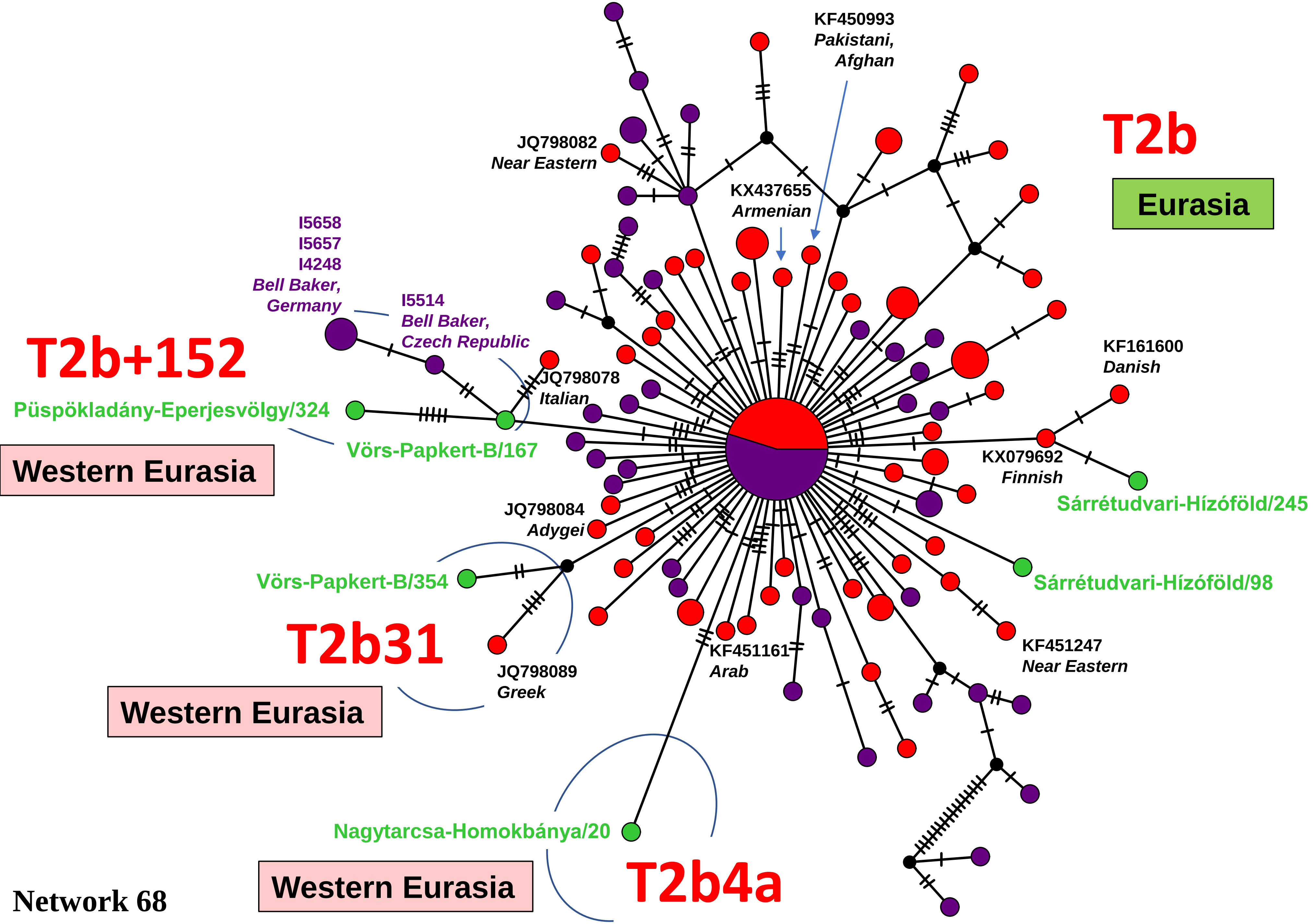


Middle circle:

STR_491
STR_355
Early Middle Ages,
Germany
poz678
Trzciniec Culture,
Poland
MN540533
Middle Ages
Crusade,
Finland
MH298073
Middle Ages,
Romania
MH043577
Punic,
Spain
MG429037
MG429032
Trzciniec Culture,
Lithuania
I5531
Bell Beaker,
Germany
I5382
Bell Beaker
Great Britain
I5377
Early Bronze Age,
Great Britain
I4189
Copper Age
Hungary

I1905
Late Neolithic,
Hungary
I2022
I0026
I0025
I0021
Linear Pottery,
Germany

KP635213
English
KM047204
Polish
KF451074
French
KF162406
KF162084
KF161799
KF161577
KF161348
KF161332
JX153338
JX153295
Danish
JQ664540
Crotian
JN032303
Finnish
KX017465
German

Network 68

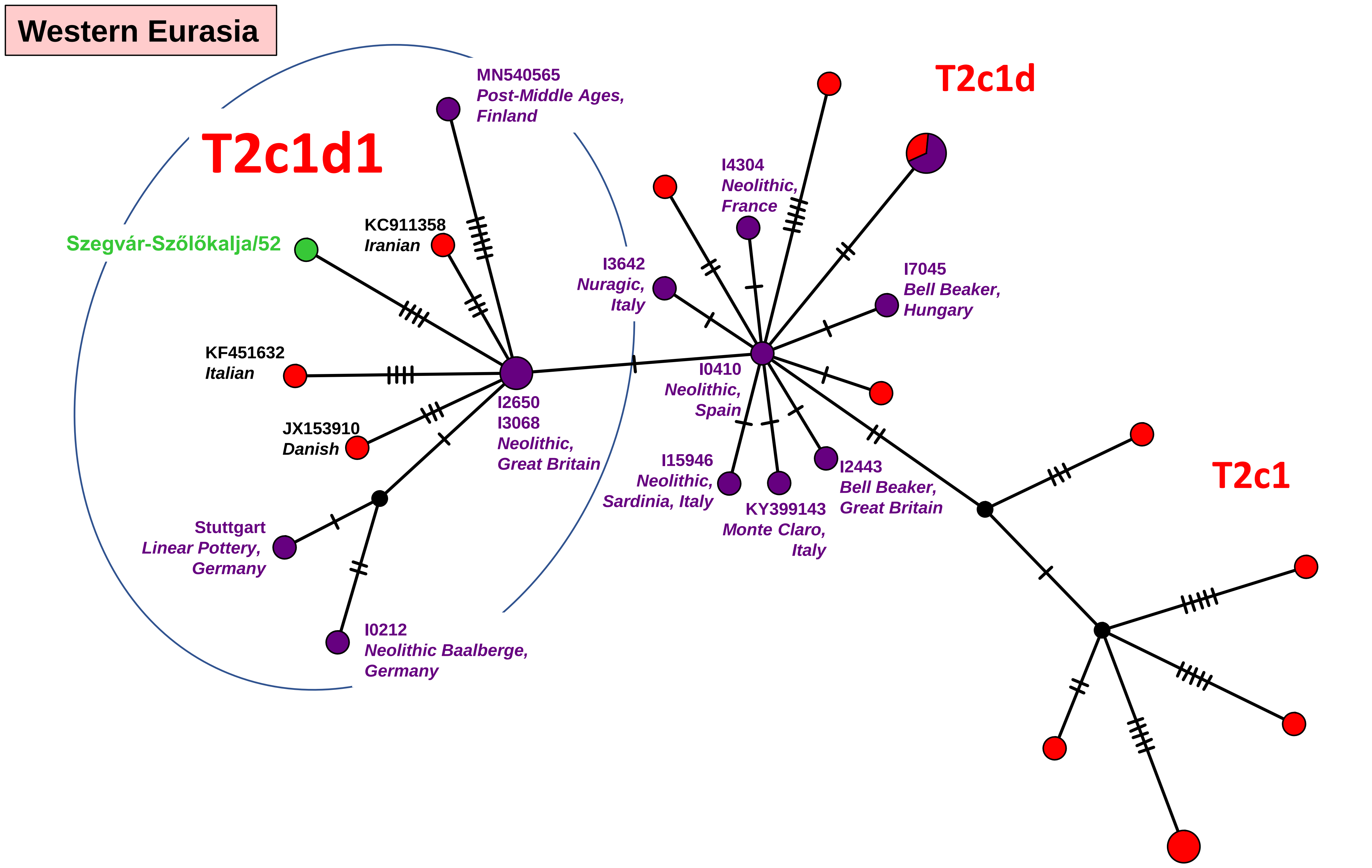

Western Eurasia
T2c1d1
T2c1d
T2c1
MN540565
Post-Middle Ages,
Finland
Szegvár-Szőlőkalja/52
KC911358
Iranian
KF451632
Italian
JX153910
Danish
Stuttgart
Linear Pottery,
Germany
I0212
Neolithic Baalberge,
Germany
I2650
I3068
Neolithic,
Great Britain
I3642
Nuragic,
Italy
I4304
Neolithic,
France
I7045
Bell Beaker,
Hungary
I0410
Neolithic,
Spain
I15946
Neolithic,
Sardinia, Italy
KY399143
Monte Claro,
Italy
I2443
Bell Beaker,
Great Britain


Network 69

### Eastern Eurasia

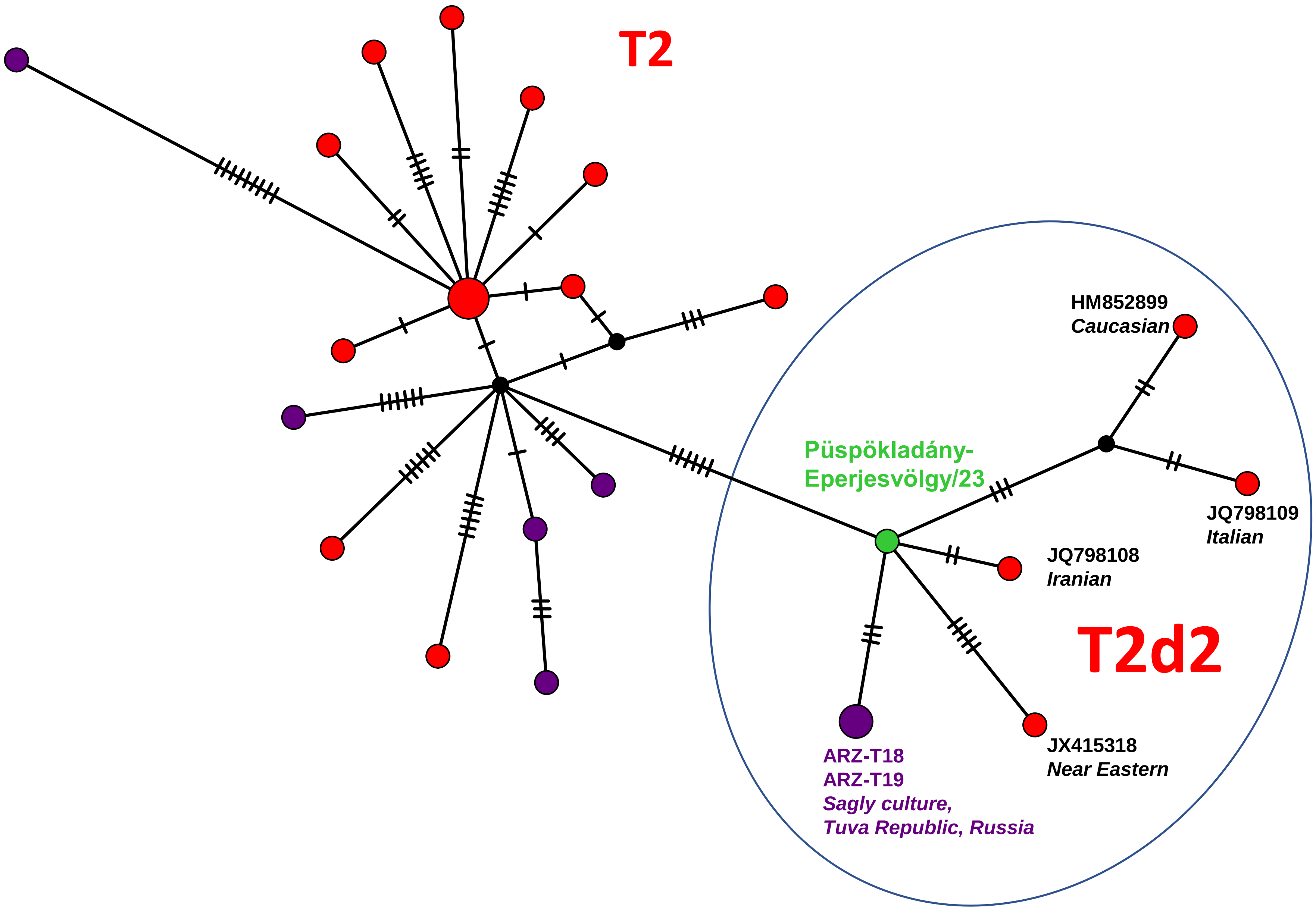


Network 70

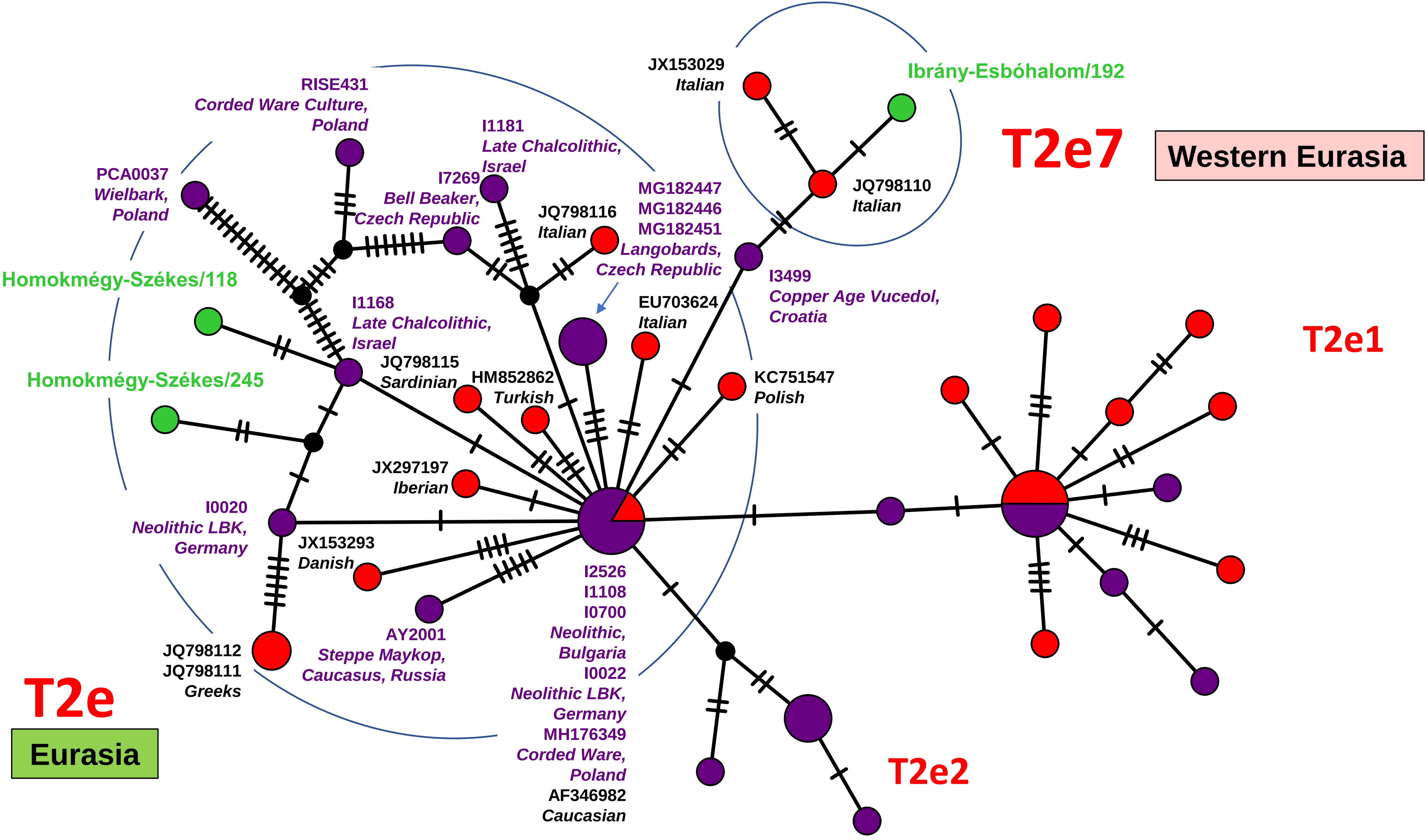

T2e
Eurasia
T2e7
Western Eurasia
T2e1
T2e2
JX153029
Italian
Ibrány-Esbóhalom/192
JQ798110
Italian
RISE431
Corded Ware Culture,
Poland
I1181
Late Chalcolithic,
Israel
PCA0037
Wielbark,
Poland
I7269
Bell Beaker,
Czech Republic
JQ798116
Italian
MG182447
MG182446
MG182451
Langobards,
Czech Republic
I3499
Copper Age Vucedol,
Croatia
Homokmégy-Székes/118
I1168
Late Chalcolithic,
Israel
EU703624
Italian
JQ798115
Sardinian
HM852862
Turkish
KC751547
Polish
Homokmégy-Székes/245
JX297197
Iberian
I0020
Neolithic LBK,
Germany
JX153293
Danish
I2526
I1108
I0700
Neolithic,
Bulgaria
I0022
Neolithic LBK,
Germany
MH176349
Corded Ware,
Poland
AF346982
Caucasian
AY2001
Steppe Maykop,
Caucasus, Russia
JQ798112
JQ798111
Greeks


Network 71

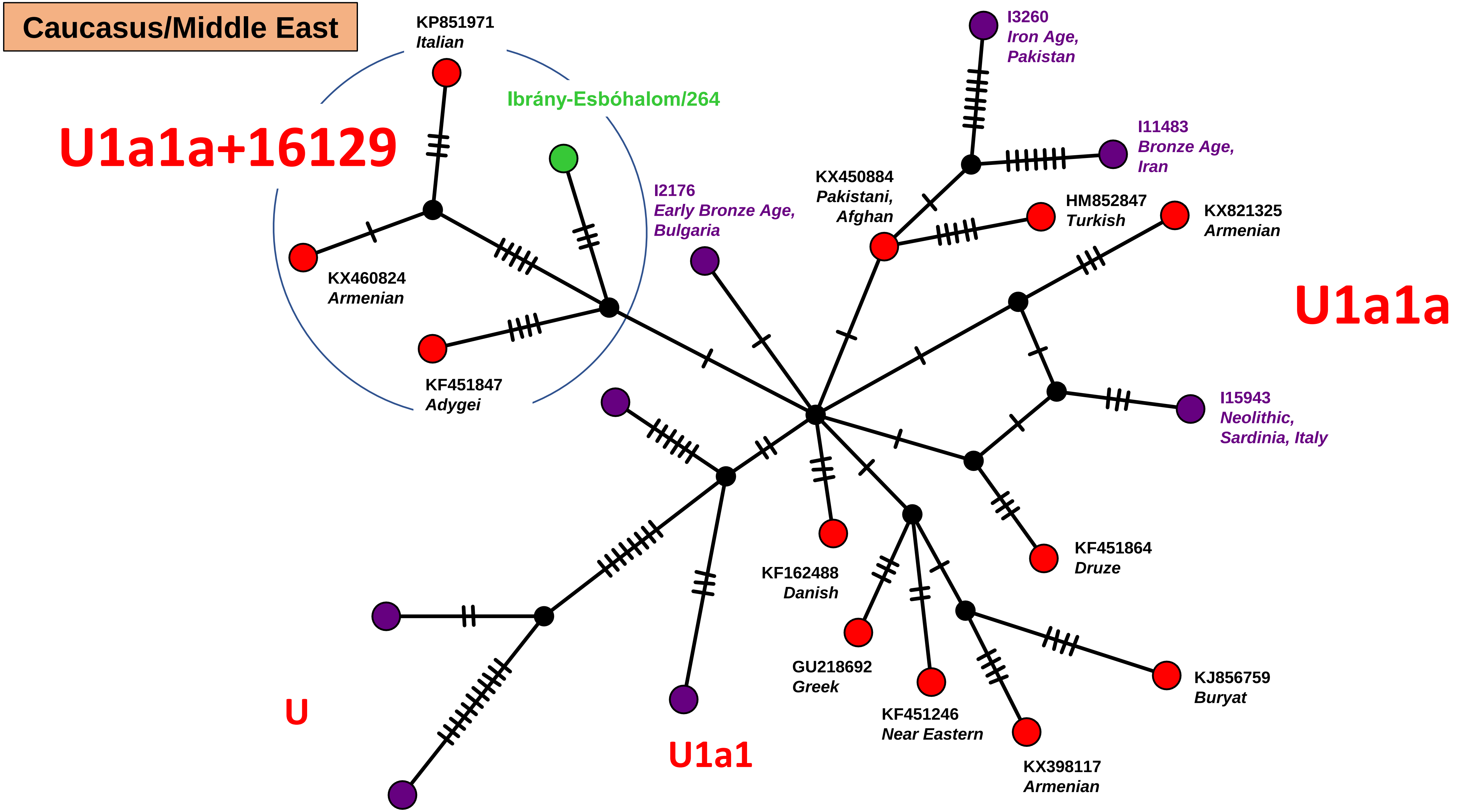
Caucasus/Middle East
KP851971
Italian
Ibrány-Esbóhalom/264
U1a1a+16129
KX460824
Armenian
KF451847
Adygei
I2176
Early Bronze Age,
Bulgaria
KX450884
Pakistani,
Afghan
I3260
Iron Age,
Pakistan
I11483
Bronze Age,
Iran
HM852847
Turkish
KX821325
Armenian
U1a1a
I15943
Neolithic,
Sardinia, Italy
KF451864
Druze
KF162488
Danish
GU218692
Greek
KF451246
Near Eastern
KX398117
Armenian
KJ856759
Buryat
U
U1a1

Network 72

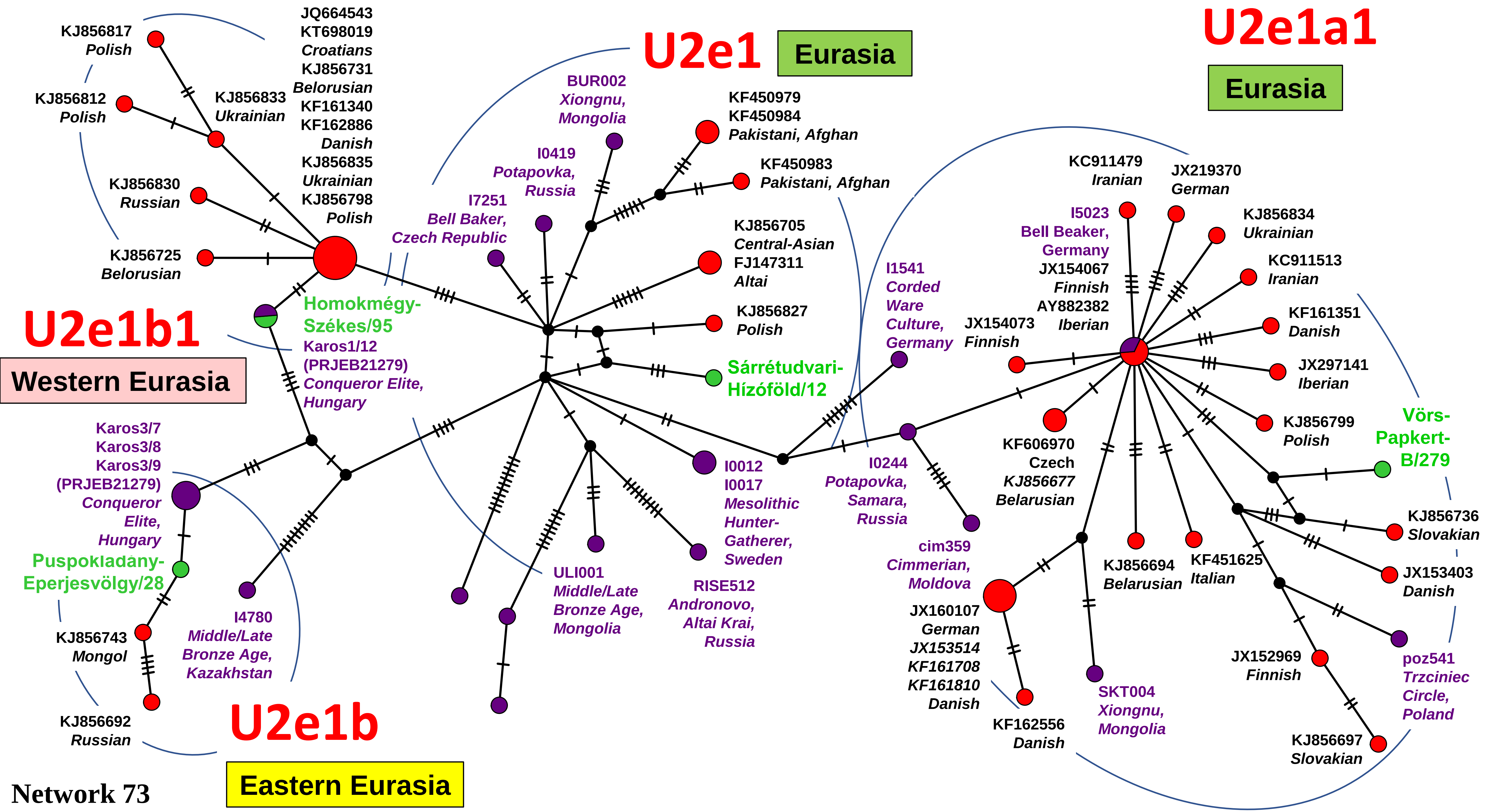

U2e1
Eurasia
U2e1a1
Eurasia
U2e1b1
Western Eurasia
U2e1b
Eastern Eurasia
KJ856817 Polish
KJ856812 Polish
KJ856833 Ukrainian
JQ664543 KT698019 Croatians KJ856731 Belorusian KF161340 KF162886 Danish KJ856835 Ukrainian KJ856798 Polish
KJ856830 Russian
KJ856725 Belorusian
Homokmégy-Székes/95
Karos1/12 (PRJEB21279) Conqueror Elite, Hungary
Karos3/7 Karos3/8 Karos3/9 (PRJEB21279) Conqueror Elite, Hungary
Puspokladány-Eperjesvölgy/28
KJ856743 Mongol
KJ856692 Russian
I4780 Middle/Late Bronze Age, Kazakhstan
BUR002 Xiongnu, Mongolia
I0419 Potapovka, Russia
I7251 Bell Baker, Czech Republic
KF450979 KF450984 Pakistani, Afghan
KF450983 Pakistani, Afghan
KJ856705 Central-Asian FJ147311 Altai
KJ856827 Polish
Sárrétudvari-Hízóföld/12
I0012 I0017 Mesolithic Hunter-Gatherer, Sweden
ULI001 Middle/Late Bronze Age, Mongolia
RISE512 Andronovo, Altai Krai, Russia
I1541 Corded Ware Culture, Germany
I0244 Potapovka, Samara, Russia
cim359 Cimmerian, Moldova
KC911479 Iranian
JX219370 German
I5023 Bell Beaker, Germany
JX154067 Finnish
AY882382 Iberian
JX154073 Finnish
KJ856834 Ukrainian
KC911513 Iranian
KF161351 Danish
JX297141 Iberian
KJ856799 Polish
Vörs-Papkert-B/279
KF606970 Czech KJ856677 Belarusian
KJ856736 Slovakian
JX153403 Danish
KJ856694 Belarusian
KF451625 Italian
JX160107 German JX153514 KF161708 KF161810 Danish
KF162556 Danish
SKT004 Xiongnu, Mongolia
JX152969 Finnish
poz541 Trzciniec Circle, Poland
KJ856697 Slovakian

Network 73

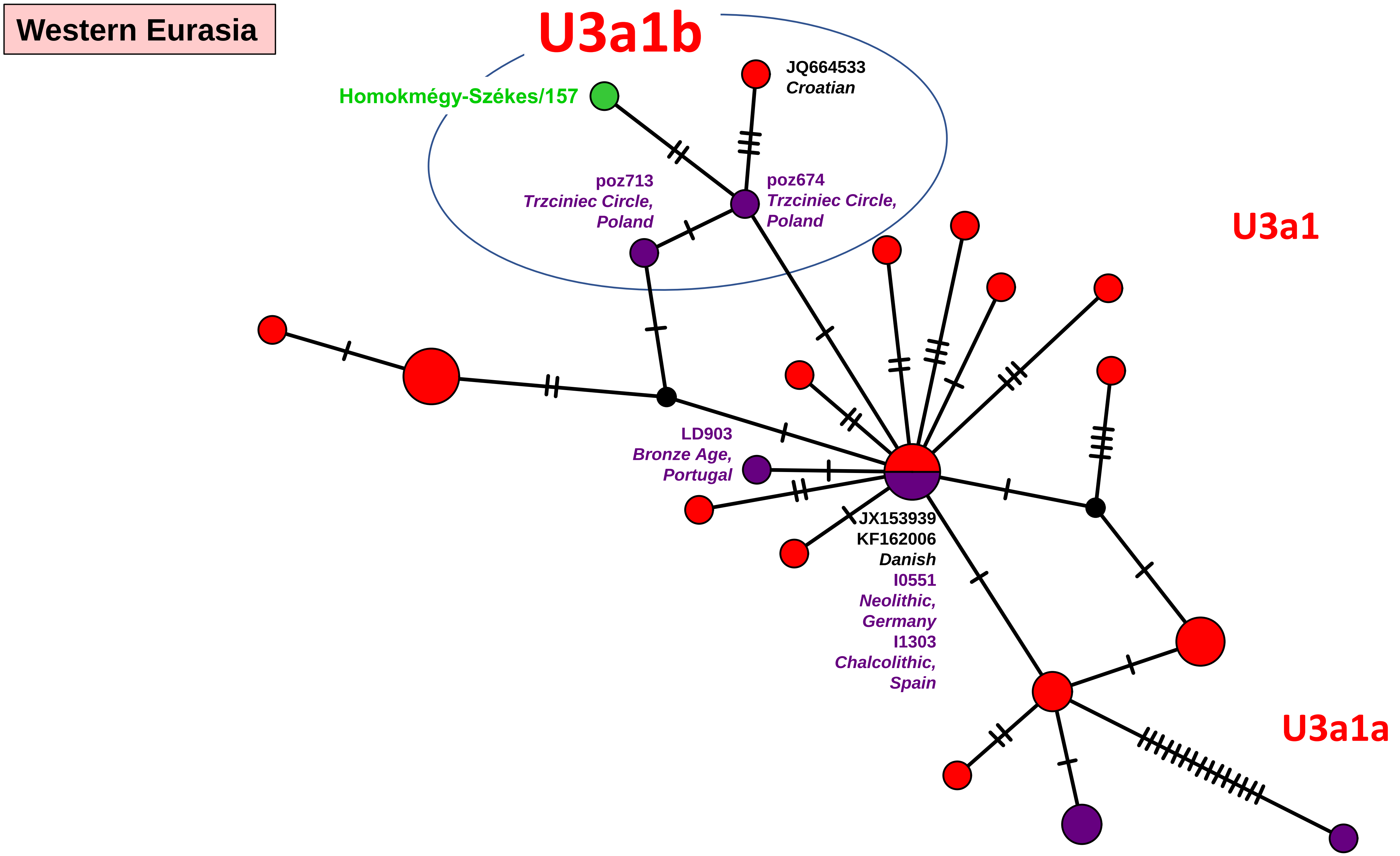
Western Eurasia
U3a1b
Homokmégy-Székes/157
JQ664533
Croatian
poz713
Trzciniec Circle,
Poland
poz674
Trzciniec Circle,
Poland
U3a1
LD903
Bronze Age,
Portugal
JX153939
KF162006
Danish
I0551
Neolithic,
Germany
I1303
Chalcolithic,
Spain
U3a1a

Network 74

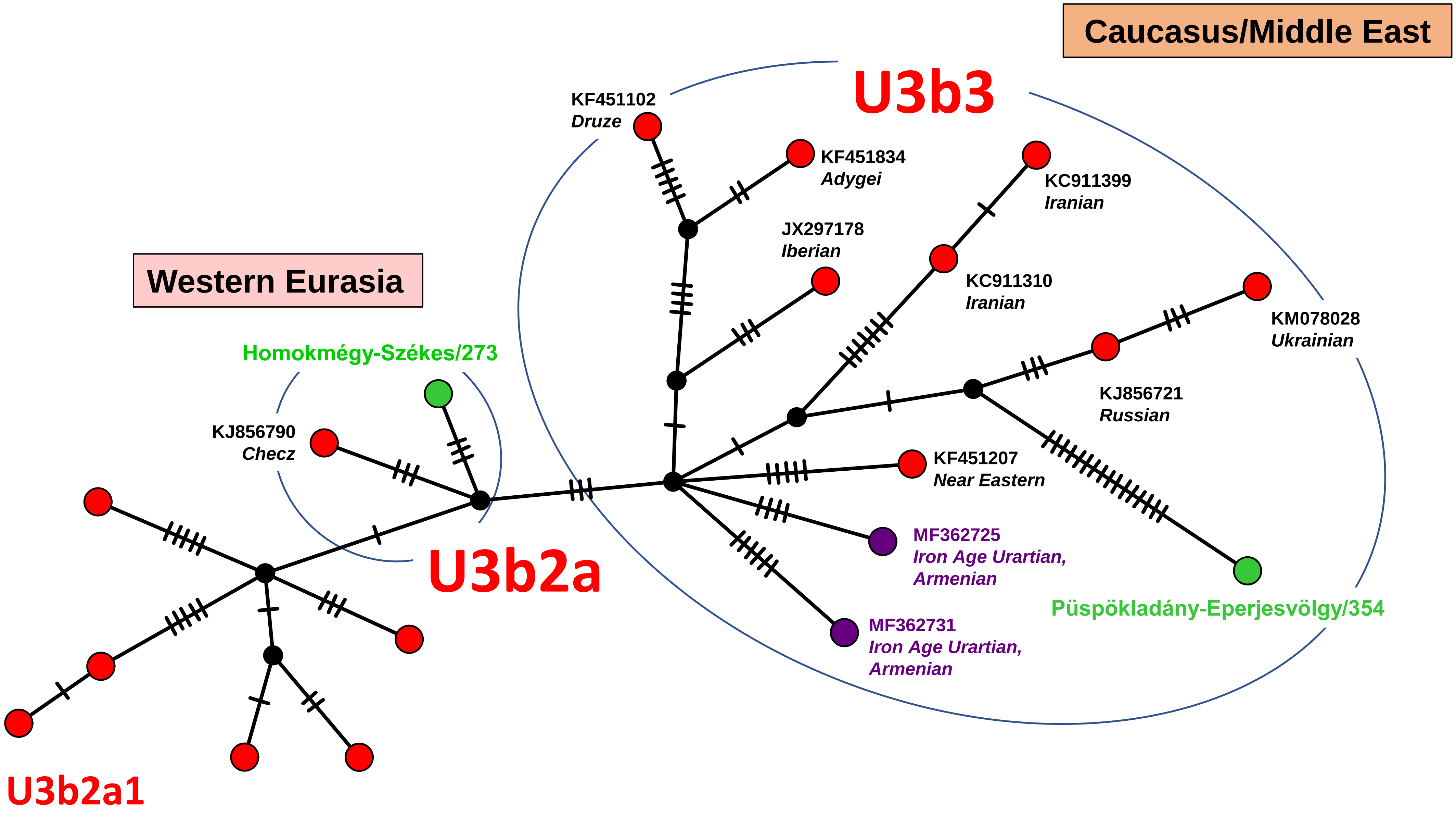
Caucasus/Middle East
U3b3
KF451102
Druze
KF451834
Adygei
KC911399
Iranian
JX297178
Iberian
KC911310
Iranian
KM078028
Ukrainian
KJ856721
Russian
KF451207
Near Eastern
MF362725
Iron Age Urartian,
Armenian
MF362731
Iron Age Urartian,
Armenian
Püspökladány-Eperjesvölgy/354
Western Eurasia
Homokmégy-Székes/273
KJ856790
Checz
U3b2a
U3b2a1

Network 75

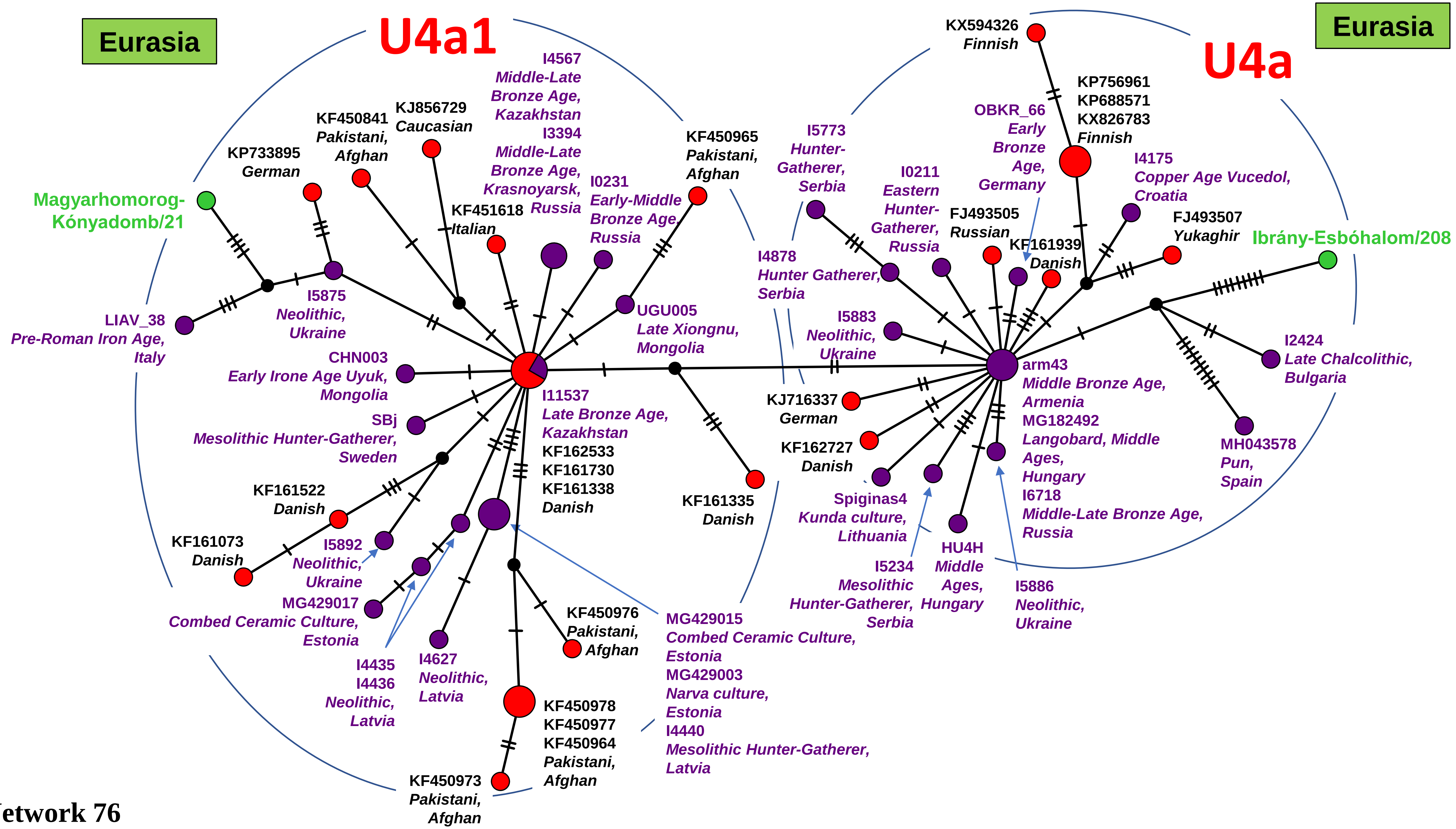

Eurasia
U4a1
Eurasia
U4a
KX594326
Finnish
KP756961
KP688571
KX826783
Finnish
OBKR_66
Early
Bronze
Age,
Germany
I4175
Copper Age Vucedol,
Croatia
FJ493507
Yukaghir
Ibrány-Esbóhalom/208
I5773
Hunter-
Gatherer,
Serbia
I0211
Eastern
Hunter-
Gatherer,
Russia
FJ493505
Russian
KF161939
Danish
I4878
Hunter Gatherer,
Serbia
I5883
Neolithic,
Ukraine
arm43
Middle Bronze Age,
Armenia
MG182492
Langobard, Middle
Ages,
Hungary
I6718
Middle-Late Bronze Age,
Russia
I2424
Late Chalcolithic,
Bulgaria
MH043578
Pun,
Spain
KJ716337
German
KF162727
Danish
Spiginas4
Kunda culture,
Lithuania
HU4H
Middle
Ages,
Hungary
I5234
Mesolithic
Hunter-Gatherer,
Serbia
I5886
Neolithic,
Ukraine
I4567
Middle-Late
Bronze Age,
Kazakhstan
I3394
Middle-Late
Bronze Age,
Krasnoyarsk, Russia
KF450965
Pakistani,
Afghan
I0231
Early-Middle
Bronze Age,
Russia
KJ856729
Caucasian
KF450841
Pakistani,
Afghan
KP733895
German
KF451618
Italian
Magyarhomorog-
Kónyadomb/21
I5875
Neolithic,
Ukraine
LIAV_38
Pre-Roman Iron Age,
Italy
UGU005
Late Xiongnu,
Mongolia
CHN003
Early Irone Age Uyuk,
Mongolia
I11537
Late Bronze Age,
Kazakhstan
KF162533
KF161730
KF161338
Danish
SBj
Mesolithic Hunter-Gatherer,
Sweden
KF161522
Danish
KF161073
Danish
I5892
Neolithic,
Ukraine
MG429017
Combed Ceramic Culture,
Estonia
I4435
I4436
Neolithic,
Latvia
I4627
Neolithic,
Latvia
KF450976
Pakistani,
Afghan
KF161335
Danish
MG429015
Combed Ceramic Culture,
Estonia
MG429003
Narva culture,
Estonia
I4440
Mesolithic Hunter-Gatherer,
Latvia
KF450978
KF450977
KF450964
Pakistani,
Afghan
KF450973
Pakistani,
Afghan


Network 76

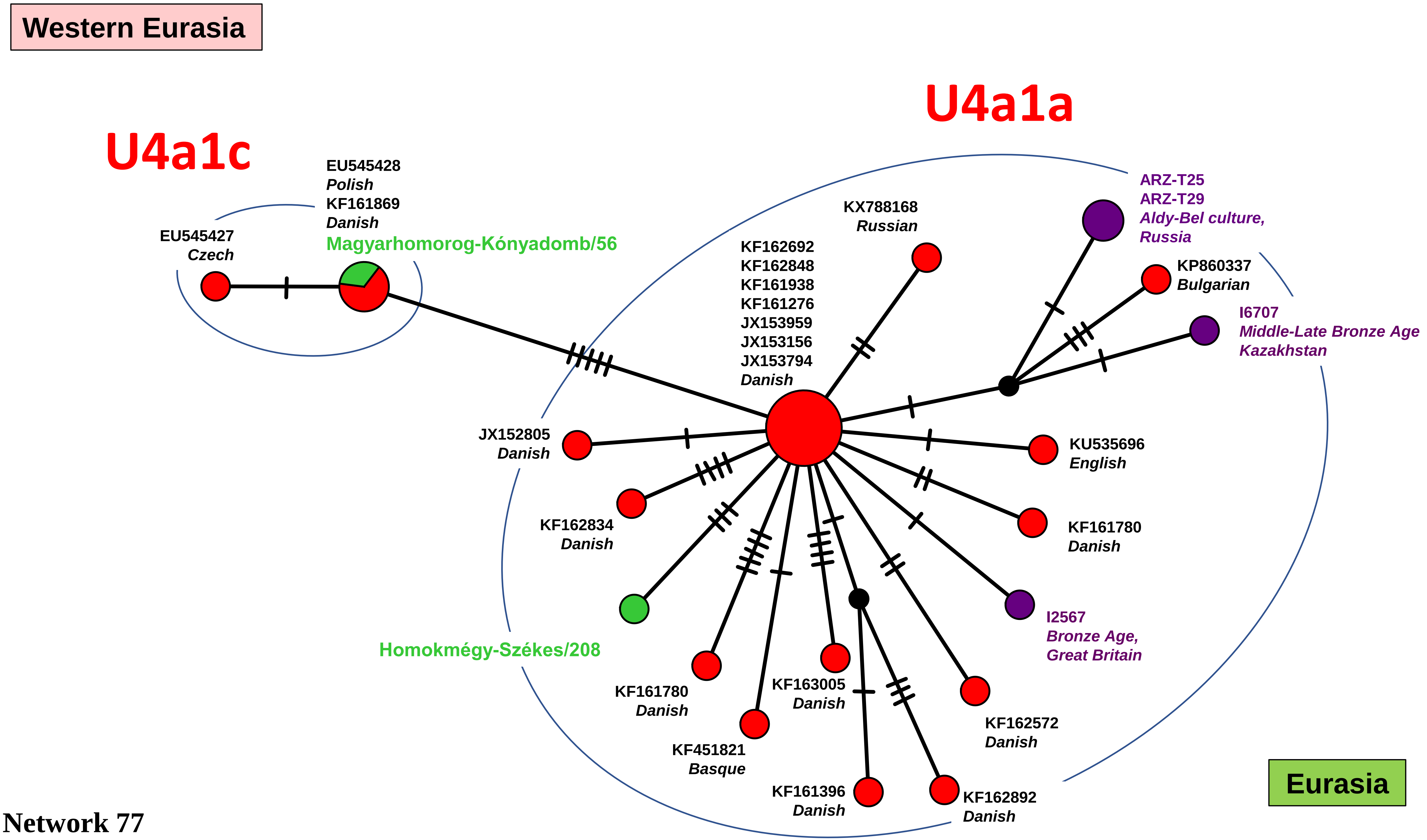
Western Eurasia
U4a1a
U4a1c
EU545428
Polish
KF161869
Danish
Magyarhomorog-Kónyadomb/56
EU545427
Czech
KX788168
Russian
KF162692
KF162848
KF161938
KF161276
JX153959
JX153156
JX153794
Danish
ARZ-T25
ARZ-T29
Aldy-Bel culture,
Russia
KP860337
Bulgarian
I6707
Middle-Late Bronze Age
Kazakhstan
JX152805
Danish
KU535696
English
KF162834
Danish
KF161780
Danish
I2567
Bronze Age,
Great Britain
Homokmégy-Székes/208
KF161780
Danish
KF163005
Danish
KF162572
Danish
KF451821
Basque
KF161396
Danish
KF162892
Danish
Eurasia

Network 77

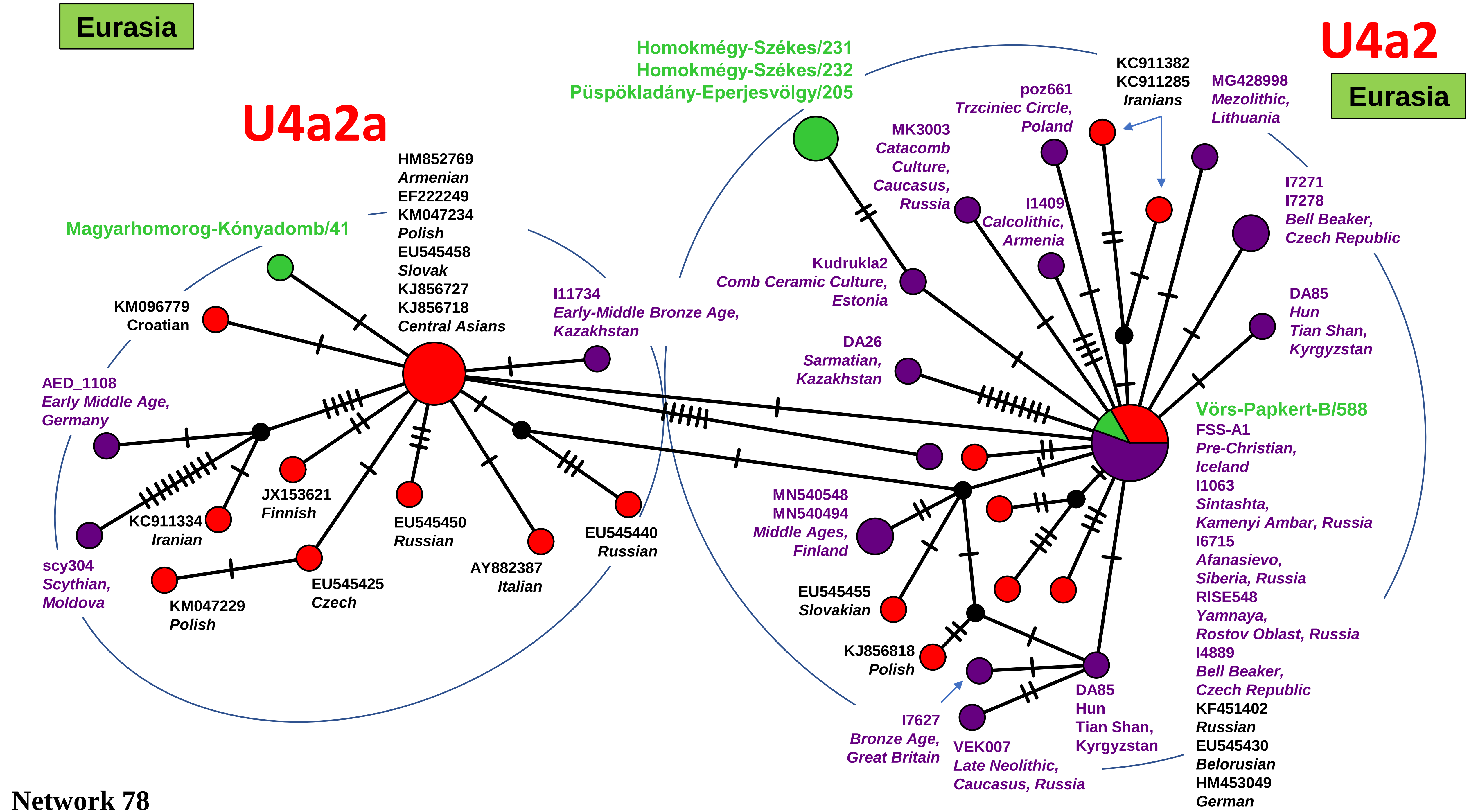
Eurasia
U4a2
Eurasia
U4a2a
Homokmégy-Székes/231
Homokmégy-Székes/232
Püspökladány-Eperjesvölgy/205
KC911382
KC911285
Iranians
MG428998
Mezolithic,
Lithuania
poz661
Trzciniec Circle,
Poland
MK3003
Catacomb
Culture,
Caucasus,
Russia
I1409
Calcolithic,
Armenia
I7271
I7278
Bell Beaker,
Czech Republic
Kudrukla2
Comb Ceramic Culture,
Estonia
DA85
Hun
Tian Shan,
Kyrgyzstan
DA26
Sarmatian,
Kazakhstan
Vörs-Papkert-B/588
FSS-A1
Pre-Christian,
Iceland
I1063
Sintashta,
Kamenyi Ambar, Russia
I6715
Afanasievo,
Siberia, Russia
RISE548
Yamnaya,
Rostov Oblast, Russia
I4889
Bell Beaker,
Czech Republic
KF451402
Russian
EU545430
Belorusian
HM453049
German
MN540548
MN540494
Middle Ages,
Finland
EU545455
Slovakian
KJ856818
Polish
I7627
Bronze Age,
Great Britain
VEK007
Late Neolithic,
Caucasus, Russia
DA85
Hun
Tian Shan,
Kyrgyzstan
HM852769
Armenian
EF222249
KM047234
Polish
EU545458
Slovak
KJ856727
KJ856718
Central Asians
Magyarhomorog-Kónyadomb/41
I11734
Early-Middle Bronze Age,
Kazakhstan
KM096779
Croatian
AED_1108
Early Middle Age,
Germany
JX153621
Finnish
KC911334
Iranian
EU545450
Russian
EU545440
Russian
AY882387
Italian
scy304
Scythian,
Moldova
EU545425
Czech
KM047229
Polish

Network 78

#### Western Eurasia

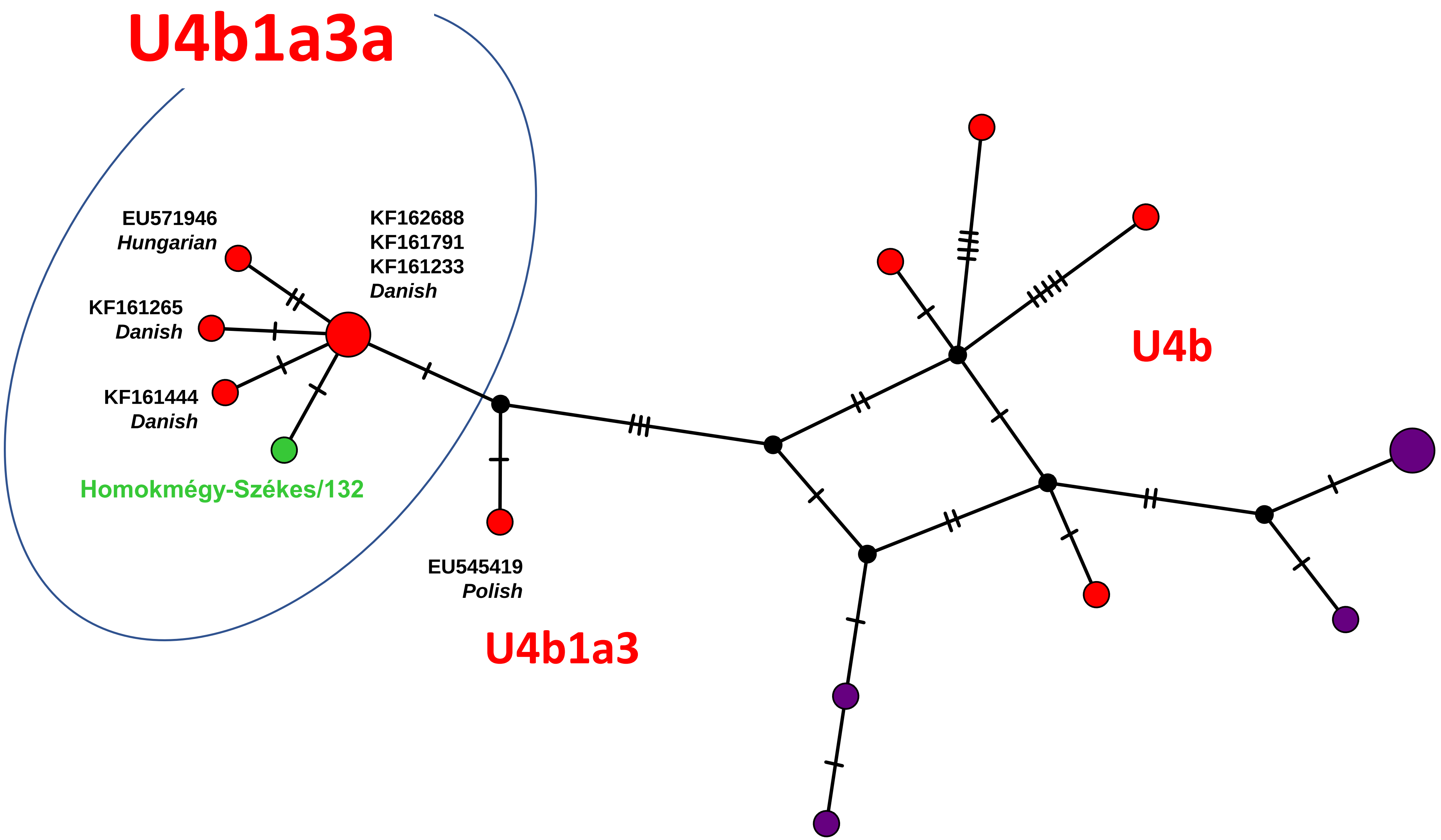


Network 79

**Eurasia**

# U4c1

**Central Circle**

KP406603
*Bulgarian*
HQ591466
*Polish*
EU545465
*Belarusian*
MH176340
*Yamnaya, Ukraine*
RISE412
*Bronze Age, Armenia*

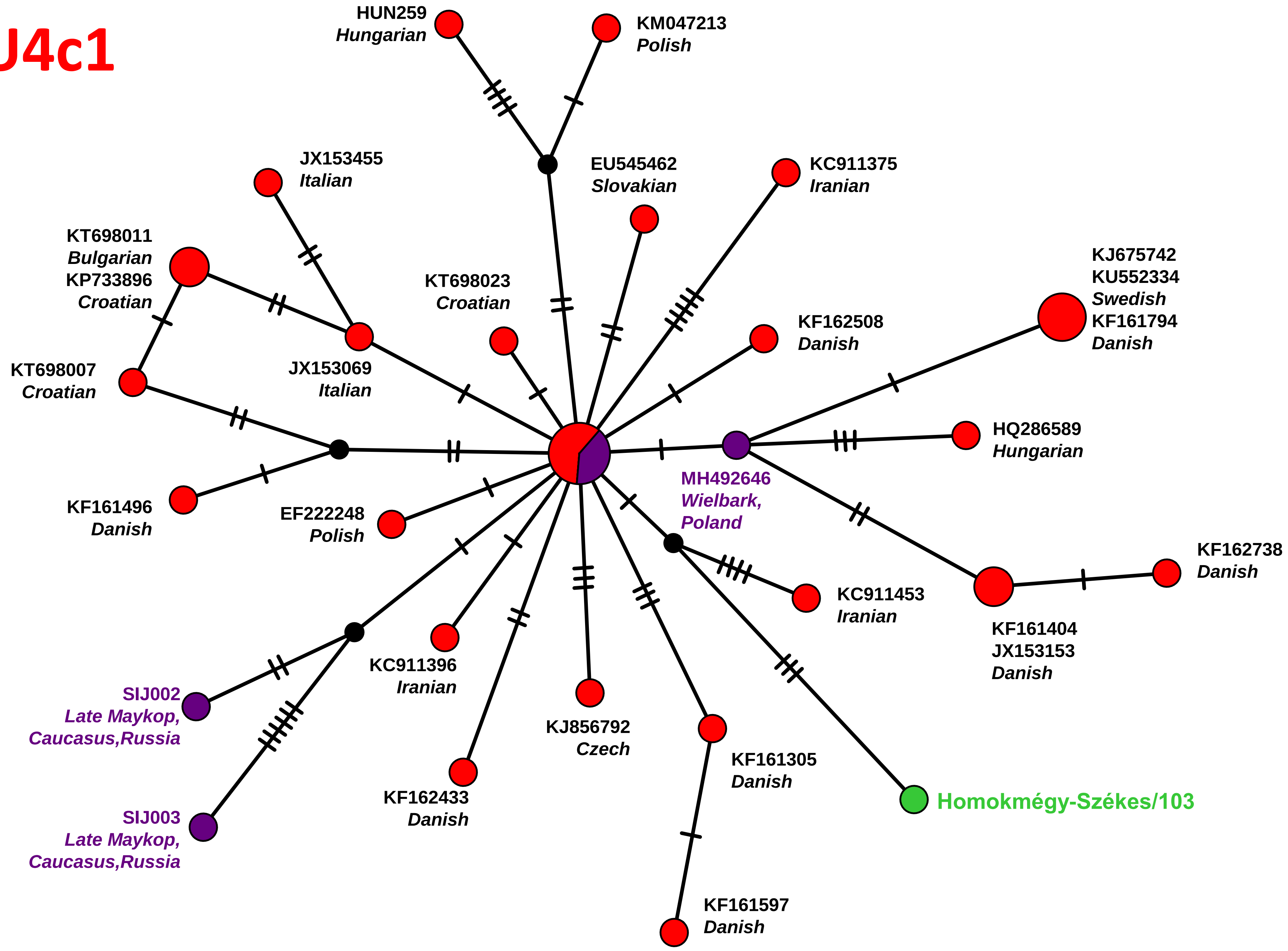


**Network 80**

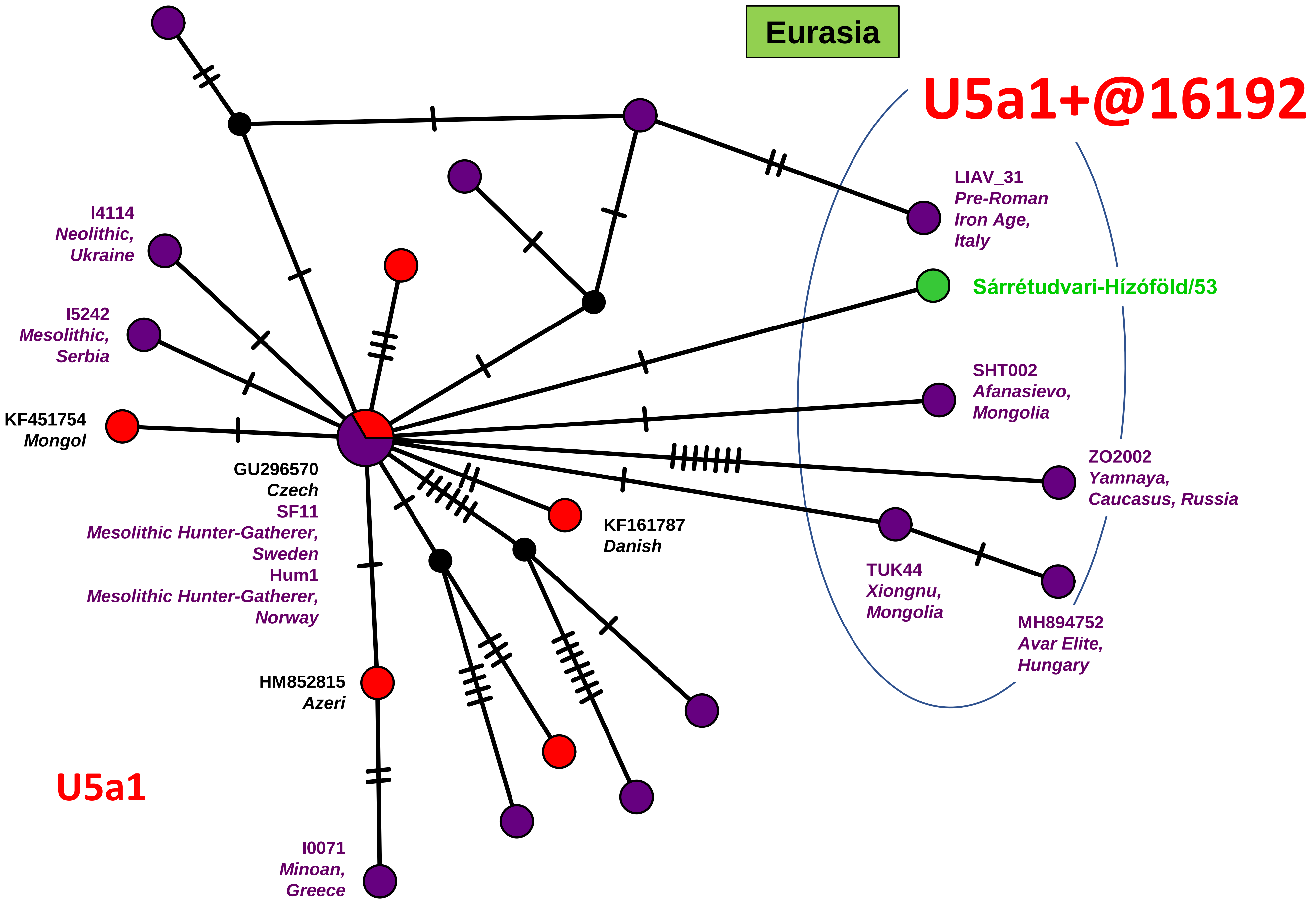
Eurasia
U5a1+@16192
LIAV_31
Pre-Roman
Iron Age,
Italy
Sárrétudvari-Hízóföld/53
SHT002
Afanasievo,
Mongolia
ZO2002
Yamnaya,
Caucasus, Russia
TUK44
Xiongnu,
Mongolia
MH894752
Avar Elite,
Hungary
I4114
Neolithic,
Ukraine
I5242
Mesolithic,
Serbia
KF451754
Mongol
GU296570
Czech
SF11
Mesolithic Hunter-Gatherer,
Sweden
Hum1
Mesolithic Hunter-Gatherer,
Norway
KF161787
Danish
HM852815
Azeri
I0071
Minoan,
Greece
U5a1

Network 81

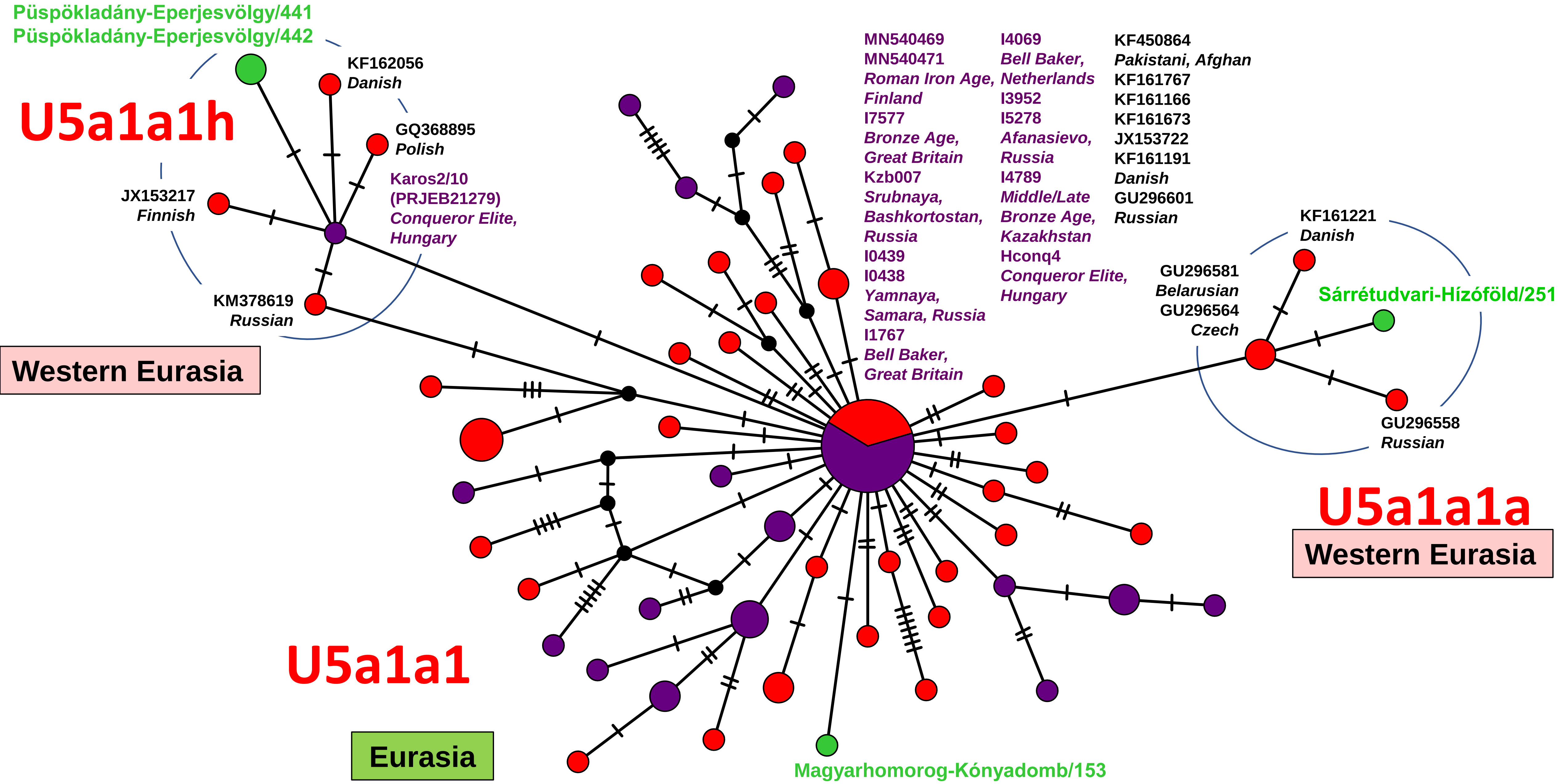
Püspökladány-Eperjesvölgy/441
Püspökladány-Eperjesvölgy/442
KF162056
Danish
U5a1a1h
GQ368895
Polish
Karos2/10
(PRJEB21279)
Conqueror Elite,
Hungary
JX153217
Finnish
KM378619
Russian
Western Eurasia
MN540469
MN540471
Roman Iron Age,
Finland
I7577
Bronze Age,
Great Britain
Kzb007
Srubnaya,
Bashkortostan,
Russia
I0439
I0438
Yamnaya,
Samara, Russia
I1767
Bell Baker,
Great Britain
I4069
Bell Baker,
Netherlands
I3952
I5278
Afanasievo,
Russia
I4789
Middle/Late
Bronze Age,
Kazakhstan
Hconq4
Conqueror Elite,
Hungary
KF450864
Pakistani, Afghan
KF161767
KF161166
KF161673
JX153722
KF161191
Danish
GU296601
Russian
KF161221
Danish
GU296581
Belarusian
GU296564
Czech
Sárrétudvari-Hízóföld/251
GU296558
Russian
U5a1a1a
Western Eurasia
U5a1a1
Eurasia
Magyarhomorog-Kónyadomb/153

Network 82

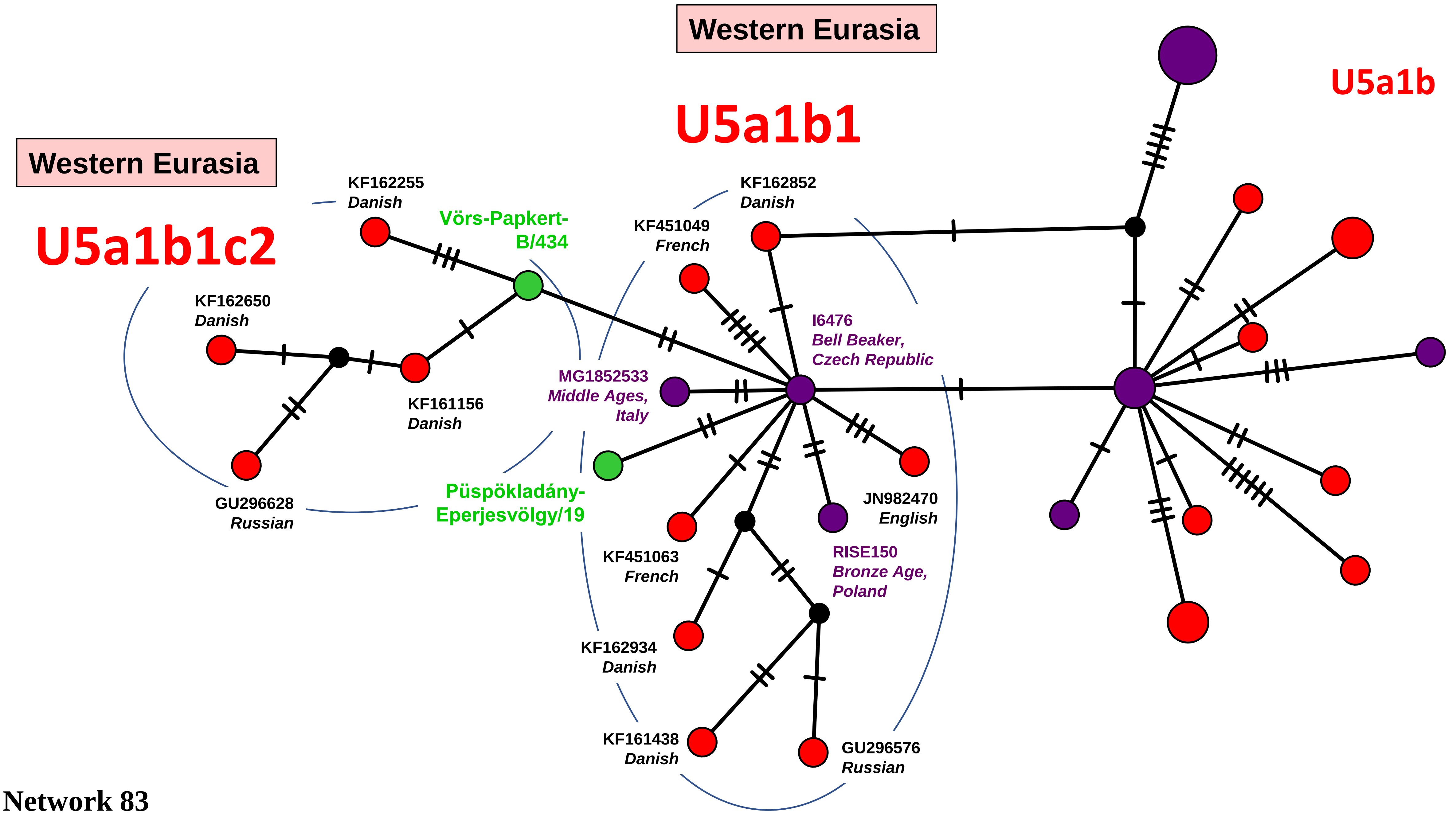
Western Eurasia
U5a1b1
U5a1b
Western Eurasia
U5a1b1c2
KF162255
Danish
Vörs-Papkert-
B/434
KF162852
Danish
KF451049
French
KF162650
Danish
I6476
Bell Beaker,
Czech Republic
MG1852533
Middle Ages,
Italy
KF161156
Danish
GU296628
Russian
Püspökladány-
Eperjesvölgy/19
JN982470
English
KF451063
French
RISE150
Bronze Age,
Poland
KF162934
Danish
KF161438
Danish
GU296576
Russian

Network 83

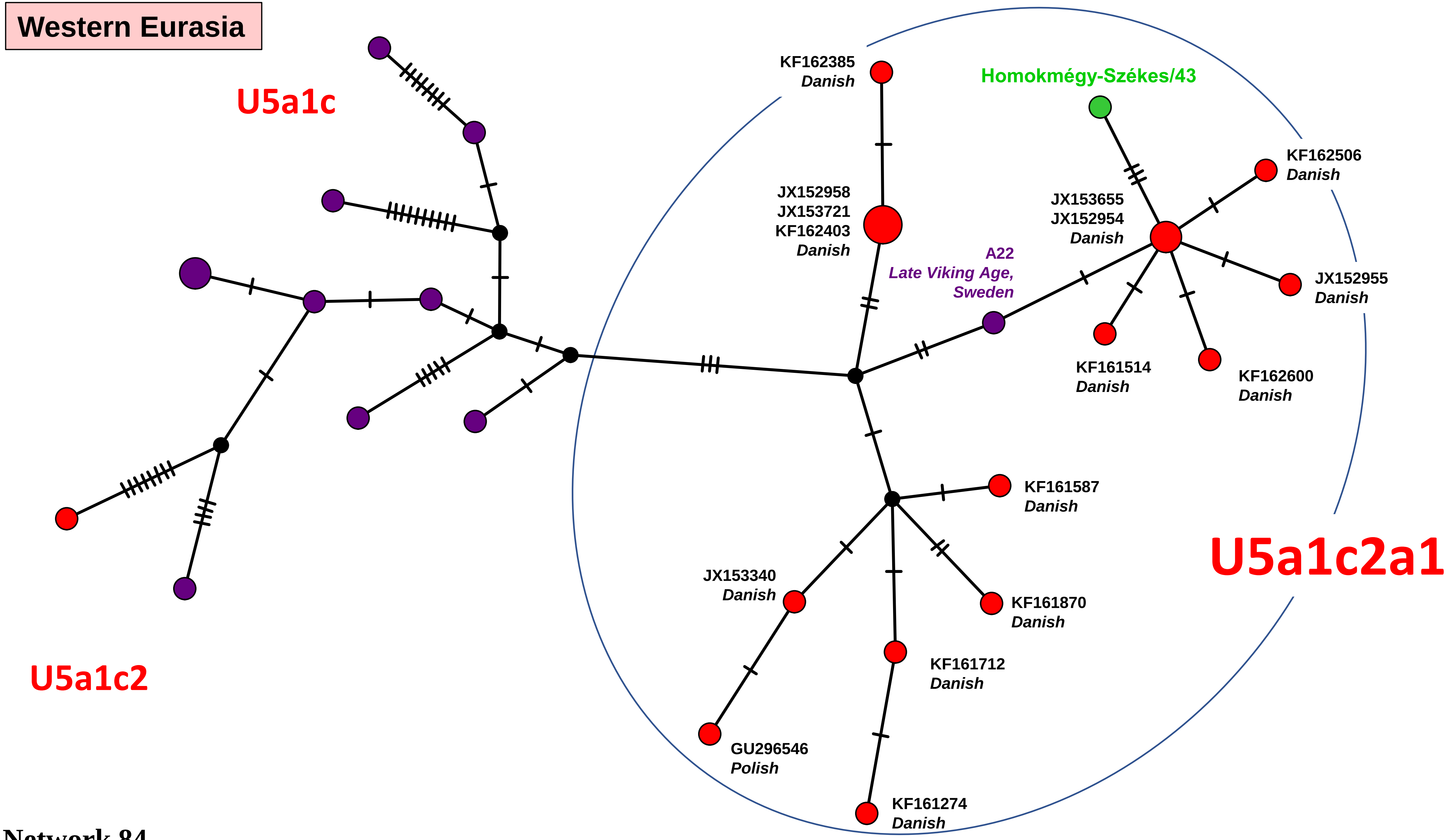

Western Eurasia
U5a1c
U5a1c2
U5a1c2a1
KF162385
Danish
Homokmégy-Székes/43
KF162506
Danish
JX152958
JX153721
KF162403
Danish
JX153655
JX152954
Danish
A22
Late Viking Age,
Sweden
JX152955
Danish
KF161514
Danish
KF162600
Danish
KF161587
Danish
JX153340
Danish
KF161870
Danish
KF161712
Danish
GU296546
Polish
KF161274
Danish


Network 84

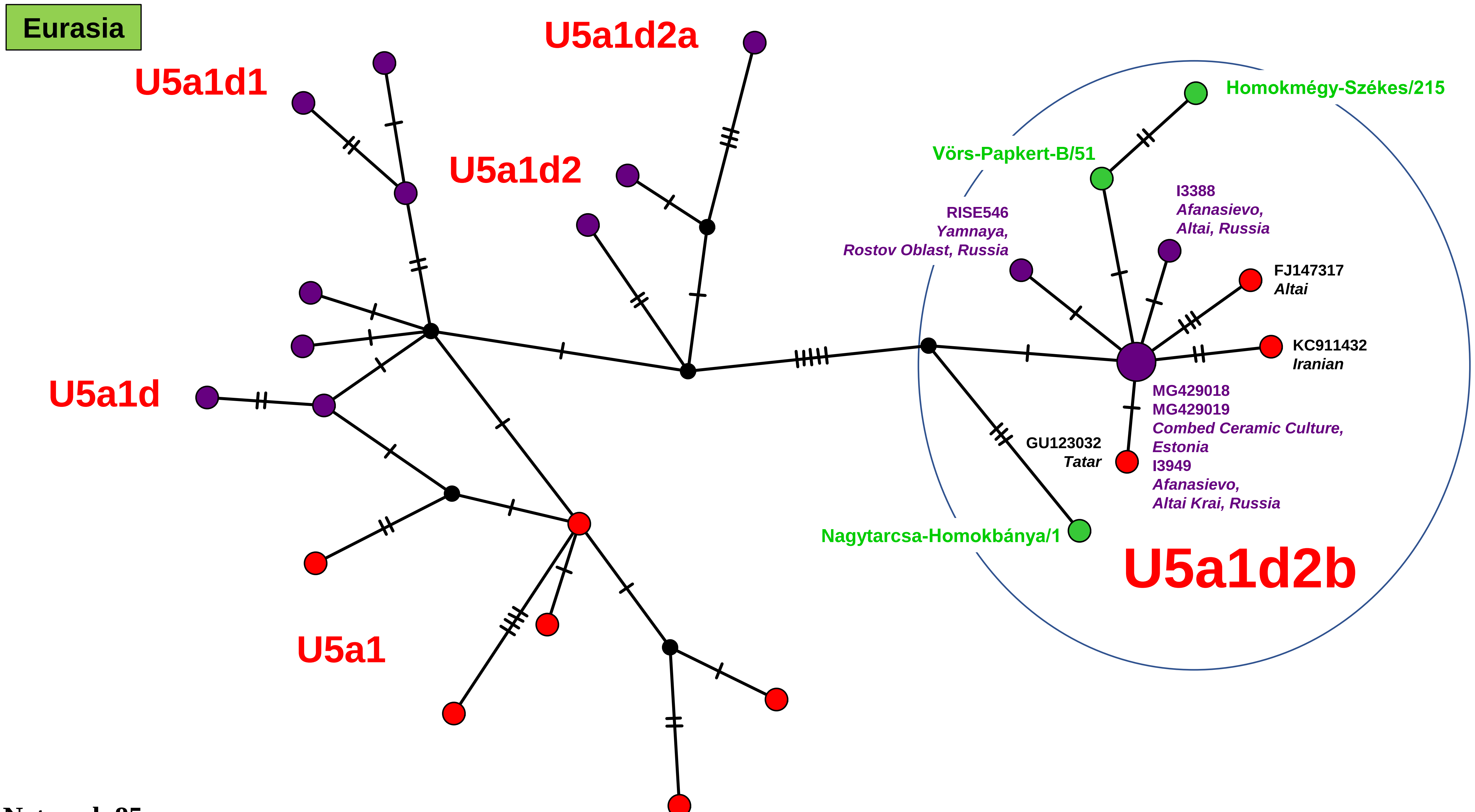
Eurasia
U5a1d2a
U5a1d1
U5a1d2
U5a1d
U5a1
Homokmégy-Székes/215
Vörs-Papkert-B/51
I3388
Afanasievo,
Altai, Russia
RISE546
Yamnaya,
Rostov Oblast, Russia
FJ147317
Altai
KC911432
Iranian
MG429018
MG429019
Combed Ceramic Culture,
Estonia
I3949
Afanasievo,
Altai Krai, Russia
GU123032
Tatar
Nagytarcsa-Homokbánya/1
U5a1d2b

Network 85

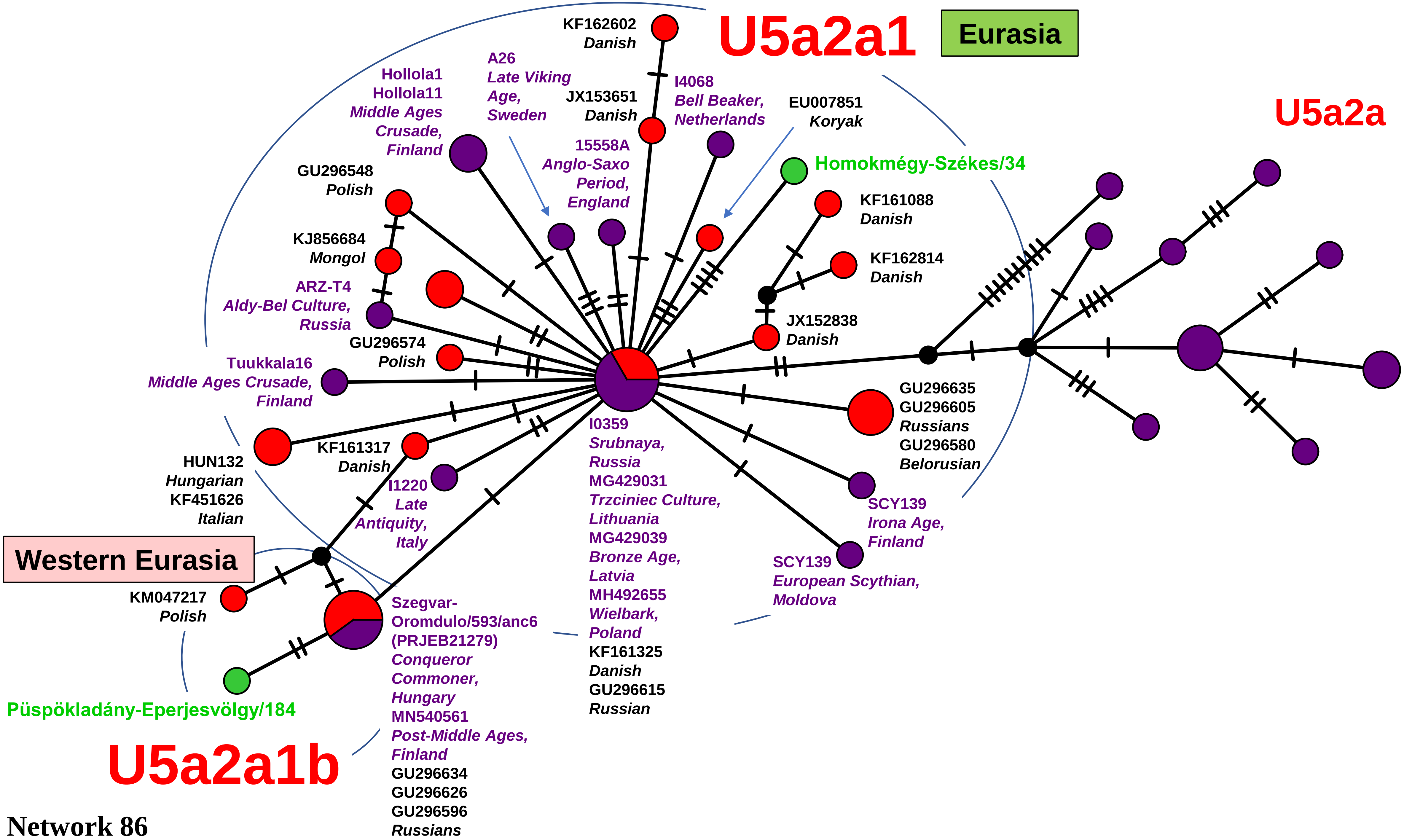

U5a2a1
Eurasia
U5a2a
KF162602
Danish
A26
Late Viking
Age,
Sweden
Hollola1
Hollola11
Middle Ages
Crusade,
Finland
JX153651
Danish
I4068
Bell Beaker,
Netherlands
EU007851
Koryak
15558A
Anglo-Saxo
Period,
England
Homokmégy-Székes/34
GU296548
Polish
KF161088
Danish
KJ856684
Mongol
KF162814
Danish
ARZ-T4
Aldy-Bel Culture,
Russia
JX152838
Danish
GU296574
Polish
Tuukkala16
Middle Ages Crusade,
Finland
GU296635
GU296605
Russians
GU296580
Belorusian
I0359
Srubnaya,
Russia
MG429031
Trzciniec Culture,
Lithuania
MG429039
Bronze Age,
Latvia
MH492655
Wielbark,
Poland
KF161325
Danish
GU296615
Russian
HUN132
Hungarian
KF451626
Italian
KF161317
Danish
I1220
Late
Antiquity,
Italy
SCY139
Irona Age,
Finland
SCY139
European Scythian,
Moldova
Western Eurasia
KM047217
Polish
Szegvar-
Oromdulo/593/anc6
(PRJEB21279)
Conqueror
Commoner,
Hungary
MN540561
Post-Middle Ages,
Finland
GU296634
GU296626
GU296596
Russians
Püspökladány-Eperjesvölgy/184
U5a2a1b


Network 86

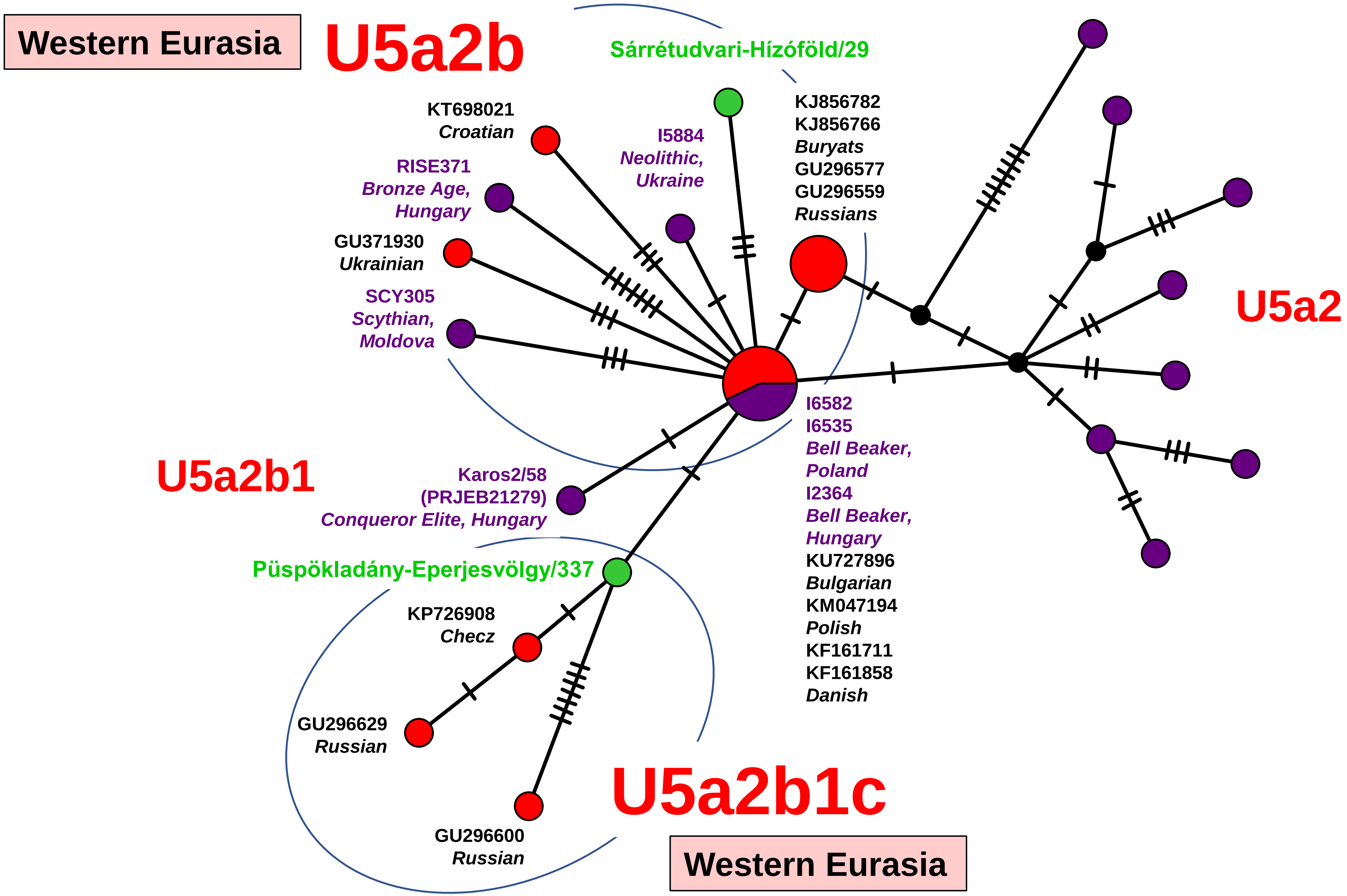

Western Eurasia
U5a2b
Sárrétudvari-Hízóföld/29
KT698021
Croatian
I5884
Neolithic,
Ukraine
KJ856782
KJ856766
Buryats
GU296577
GU296559
Russians
RISE371
Bronze Age,
Hungary
GU371930
Ukrainian
SCY305
Scythian,
Moldova
U5a2
I6582
I6535
Bell Beaker,
Poland
I2364
Bell Beaker,
Hungary
KU727896
Bulgarian
KM047194
Polish
KF161711
KF161858
Danish
U5a2b1
Karos2/58
(PRJEB21279)
Conqueror Elite, Hungary
Püspökladány-Eperjesvölgy/337
KP726908
Checz
GU296629
Russian
GU296600
Russian
U5a2b1c
Western Eurasia


Network 87

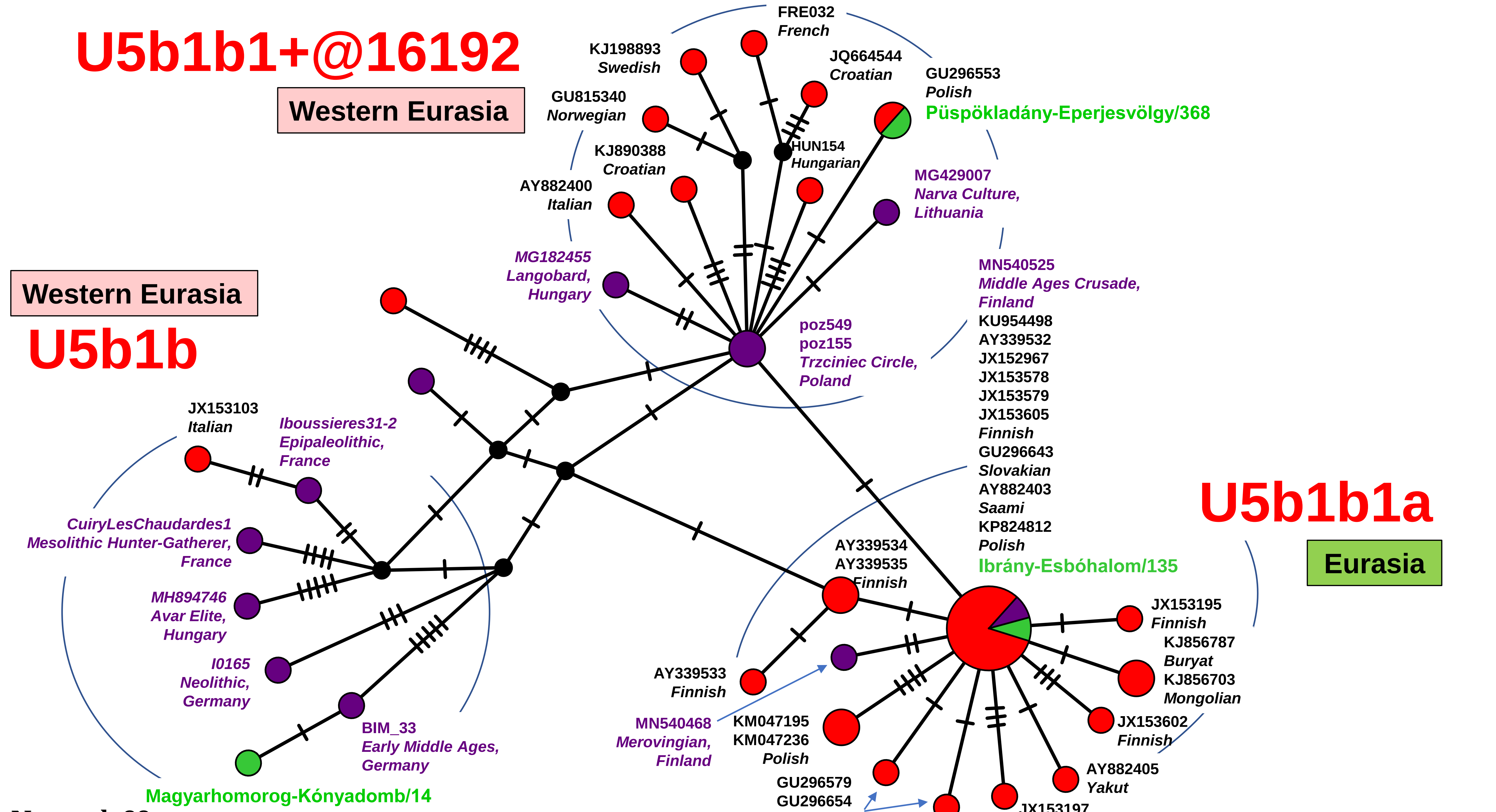

U5b1b1+@16192
Western Eurasia
U5b1b
Western Eurasia
U5b1b1a
Eurasia
FRE032
French
KJ198893
Swedish
JQ664544
Croatian
GU296553
Polish
Püspökladány-Eperjesvölgy/368
GU815340
Norwegian
HUN154
Hungarian
KJ890388
Croatian
AY882400
Italian
MG429007
Narva Culture,
Lithuania
MG182455
Langobard,
Hungary
MN540525
Middle Ages Crusade,
Finland
KU954498
AY339532
JX152967
JX153578
JX153579
JX153605
Finnish
GU296643
Slovakian
AY882403
Saami
KP824812
Polish
Ibrány-Esbóhalom/135
poz549
poz155
Trzciniec Circle,
Poland
JX153103
Italian
Iboussieres31-2
Epipaleolithic,
France
CuiryLesChaudardes1
Mesolithic Hunter-Gatherer,
France
MH894746
Avar Elite,
Hungary
I0165
Neolithic,
Germany
BIM_33
Early Middle Ages,
Germany
Magyarhomorog-Kónyadomb/14
AY339534
AY339535
Finnish
JX153195
Finnish
KJ856787
Buryat
KJ856703
Mongolian
AY339533
Finnish
MN540468
Merovingian,
Finland
KM047195
KM047236
Polish
JX153602
Finnish
GU296579
GU296654
Belarusians
AY882405
Yakut
JX153197
Finnish


Network 88

### Western Eurasia

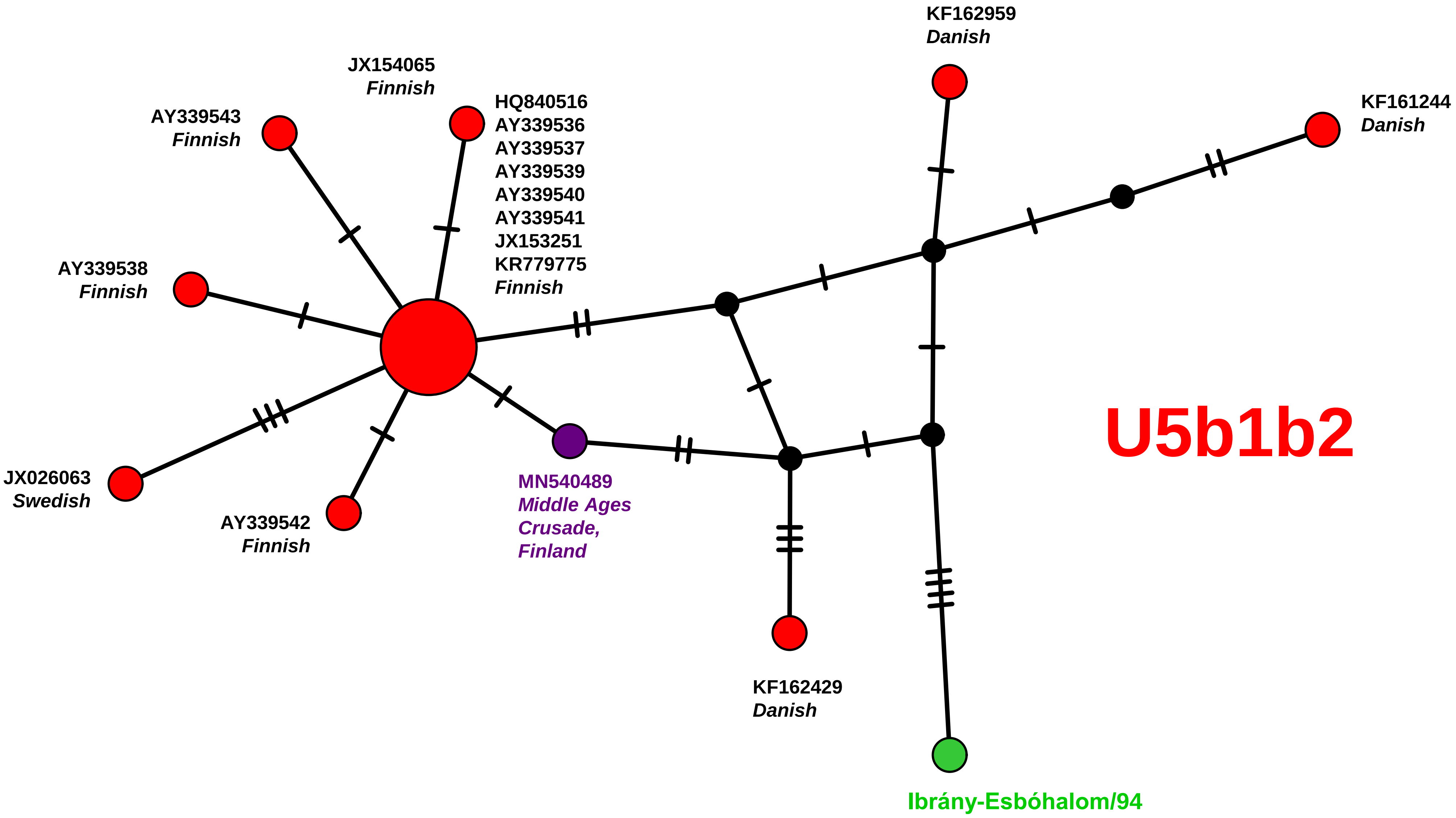


Network 89

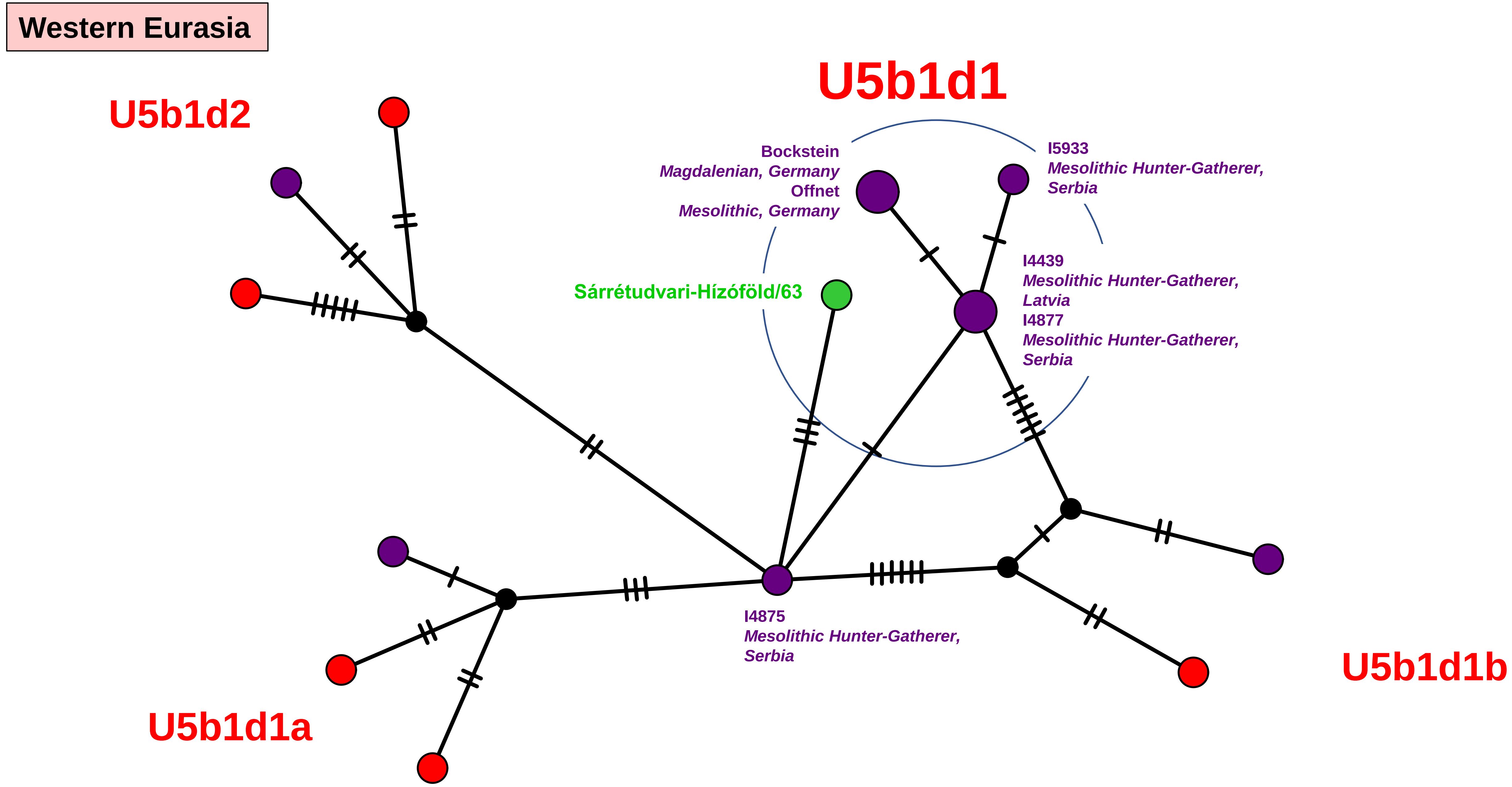
Western Eurasia
U5b1d2
U5b1d1
Bockstein
Magdalenian, Germany
Offnet
Mesolithic, Germany
I5933
Mesolithic Hunter-Gatherer,
Serbia
Sárrétudvari-Hízóföld/63
I4439
Mesolithic Hunter-Gatherer,
Latvia
I4877
Mesolithic Hunter-Gatherer,
Serbia
I4875
Mesolithic Hunter-Gatherer,
Serbia
U5b1d1a
U5b1d1b

Network 90

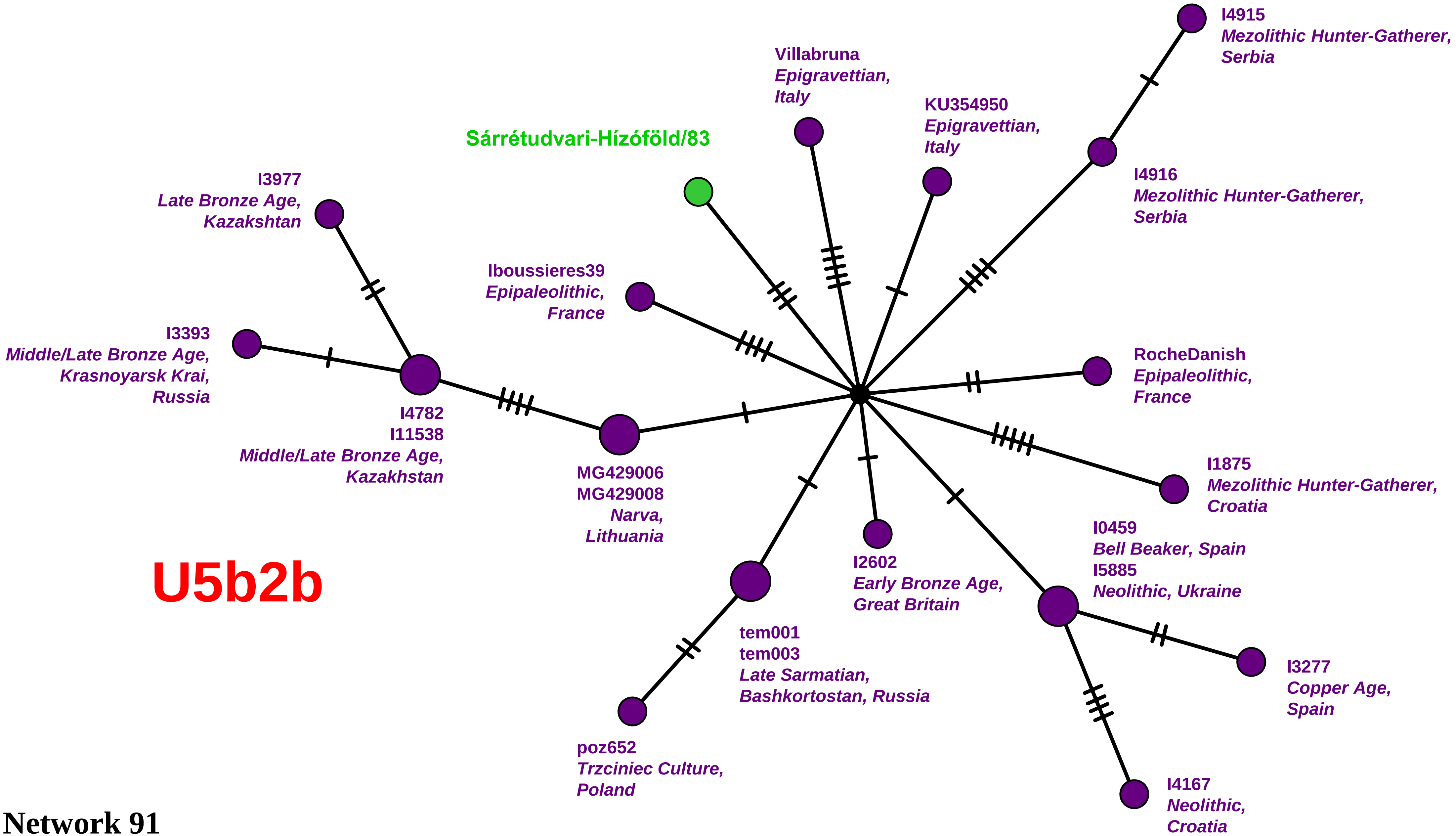


# U5b2b

Network 91

Western Eurasia

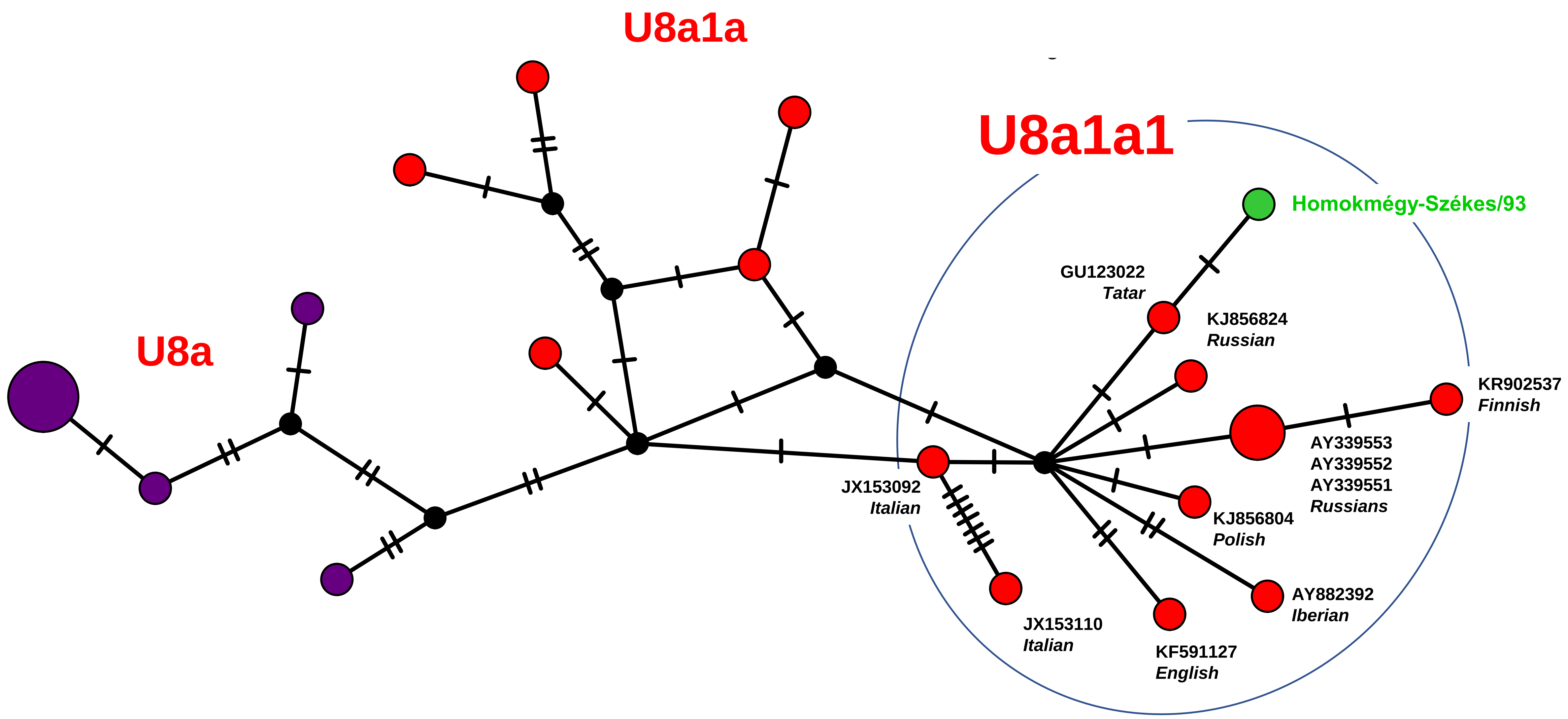


Network 92

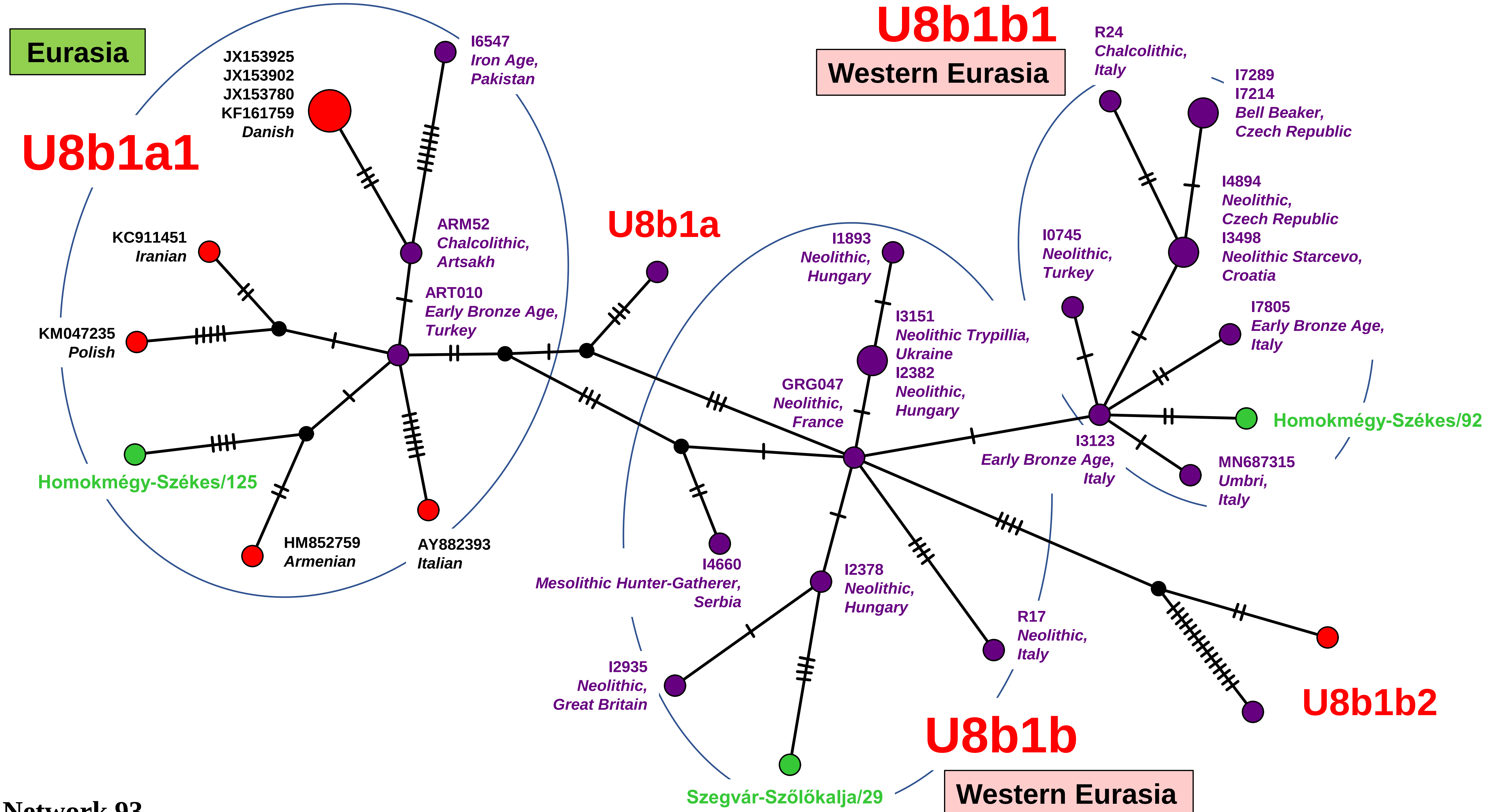
Eurasia
U8b1a1
JX153925
JX153902
JX153780
KF161759
Danish
I6547
Iron Age,
Pakistan
KC911451
Iranian
KM047235
Polish
ARM52
Chalcolithic,
Artsakh
ART010
Early Bronze Age,
Turkey
Homokmégy-Székes/125
HM852759
Armenian
AY882393
Italian
U8b1a
U8b1b1
Western Eurasia
R24
Chalcolithic,
Italy
I7289
I7214
Bell Beaker,
Czech Republic
I4894
Neolithic,
Czech Republic
I3498
Neolithic Starcevo,
Croatia
I0745
Neolithic,
Turkey
I1893
Neolithic,
Hungary
I3151
Neolithic Trypillia,
Ukraine
I2382
Neolithic,
Hungary
GRG047
Neolithic,
France
I7805
Early Bronze Age,
Italy
Homokmégy-Székes/92
I3123
Early Bronze Age,
Italy
MN687315
Umbri,
Italy
I4660
Mesolithic Hunter-Gatherer,
Serbia
I2378
Neolithic,
Hungary
R17
Neolithic,
Italy
I2935
Neolithic,
Great Britain
U8b1b2
U8b1b
Szegvár-Szőlőkalja/29
Western Eurasia

Network 93

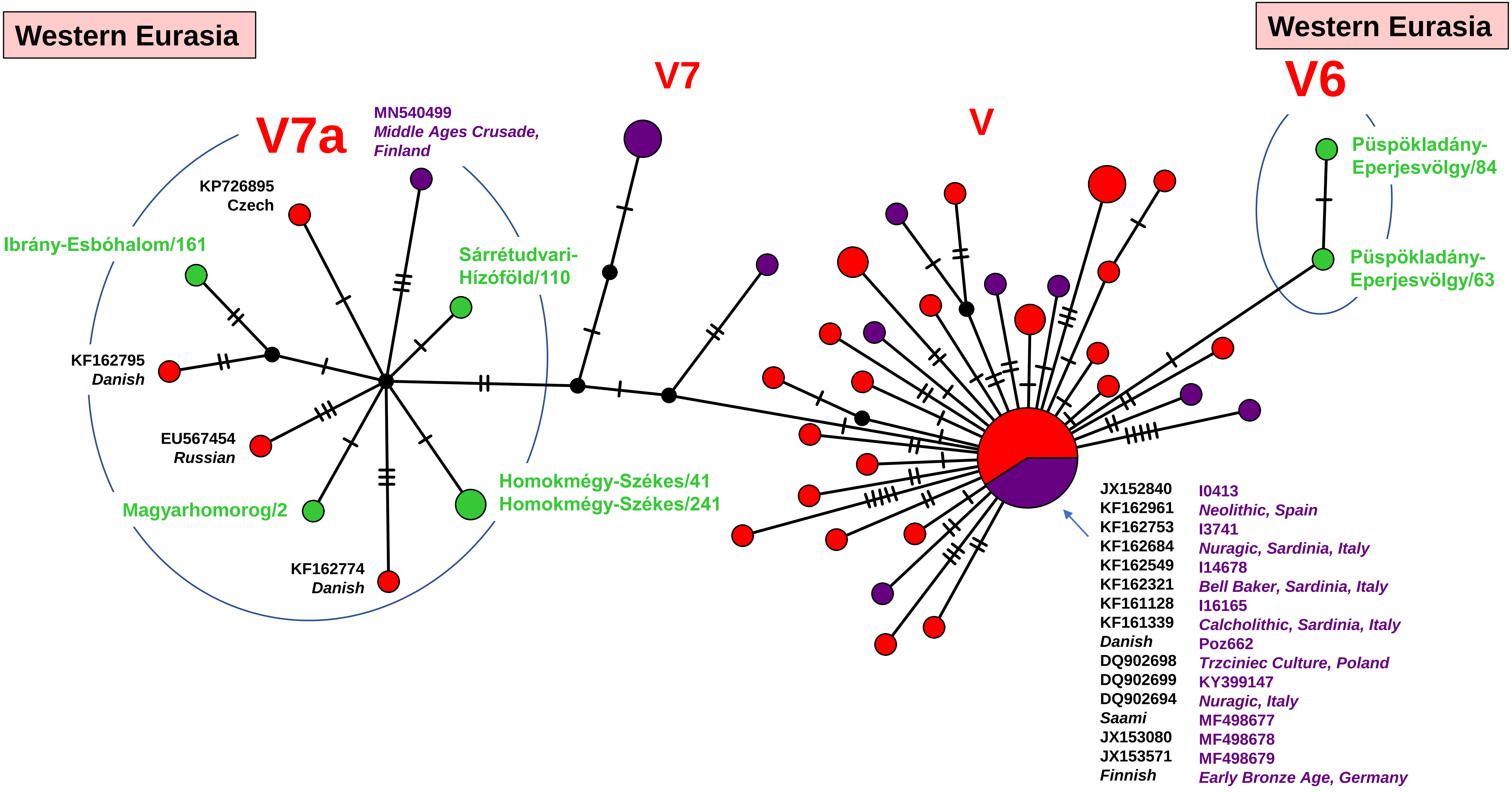
Western Eurasia
Western Eurasia
V7
V6
V7a
V
MN540499
Middle Ages Crusade, Finland
KP726895
Czech
Ibrány-Esbóhalom/161
Sárrétudvari-Hízóföld/110
KF162795
Danish
EU567454
Russian
Magyarhomorog/2
Homokmégy-Székes/41
Homokmégy-Székes/241
KF162774
Danish
Püspökladány-Eperjesvölgy/84
Püspökladány-Eperjesvölgy/63
JX152840
KF162961
KF162753
KF162684
KF162549
KF162321
KF161128
KF161339
Danish
DQ902698
DQ902699
DQ902694
Saami
JX153080
JX153571
Finnish
I0413
Neolithic, Spain
I3741
Nuragic, Sardinia, Italy
I14678
Bell Baker, Sardinia, Italy
I16165
Calcholithic, Sardinia, Italy
Poz662
Trzciniec Culture, Poland
KY399147
Nuragic, Italy
MF498677
MF498678
MF498679
Early Bronze Age, Germany

Network 94

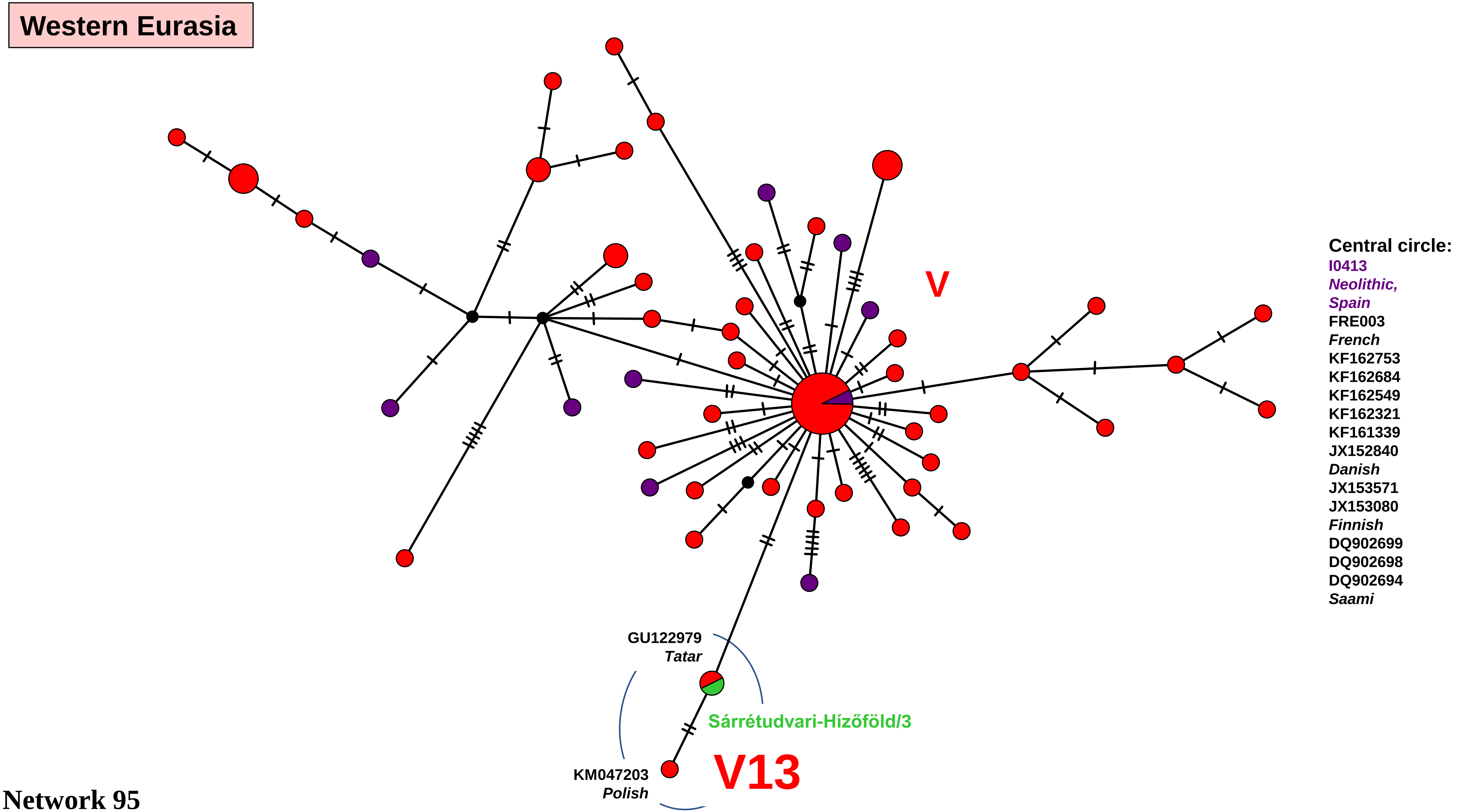

Western Eurasia
V
Central circle:
I0413
Neolithic,
Spain
FRE003
French
KF162753
KF162684
KF162549
KF162321
KF161339
JX152840
Danish
JX153571
JX153080
Finnish
DQ902699
DQ902698
DQ902694
Saami
GU122979
Tatar
Sárrétudvari-Hízóföld/3
KM047203
Polish
V13


Network 95

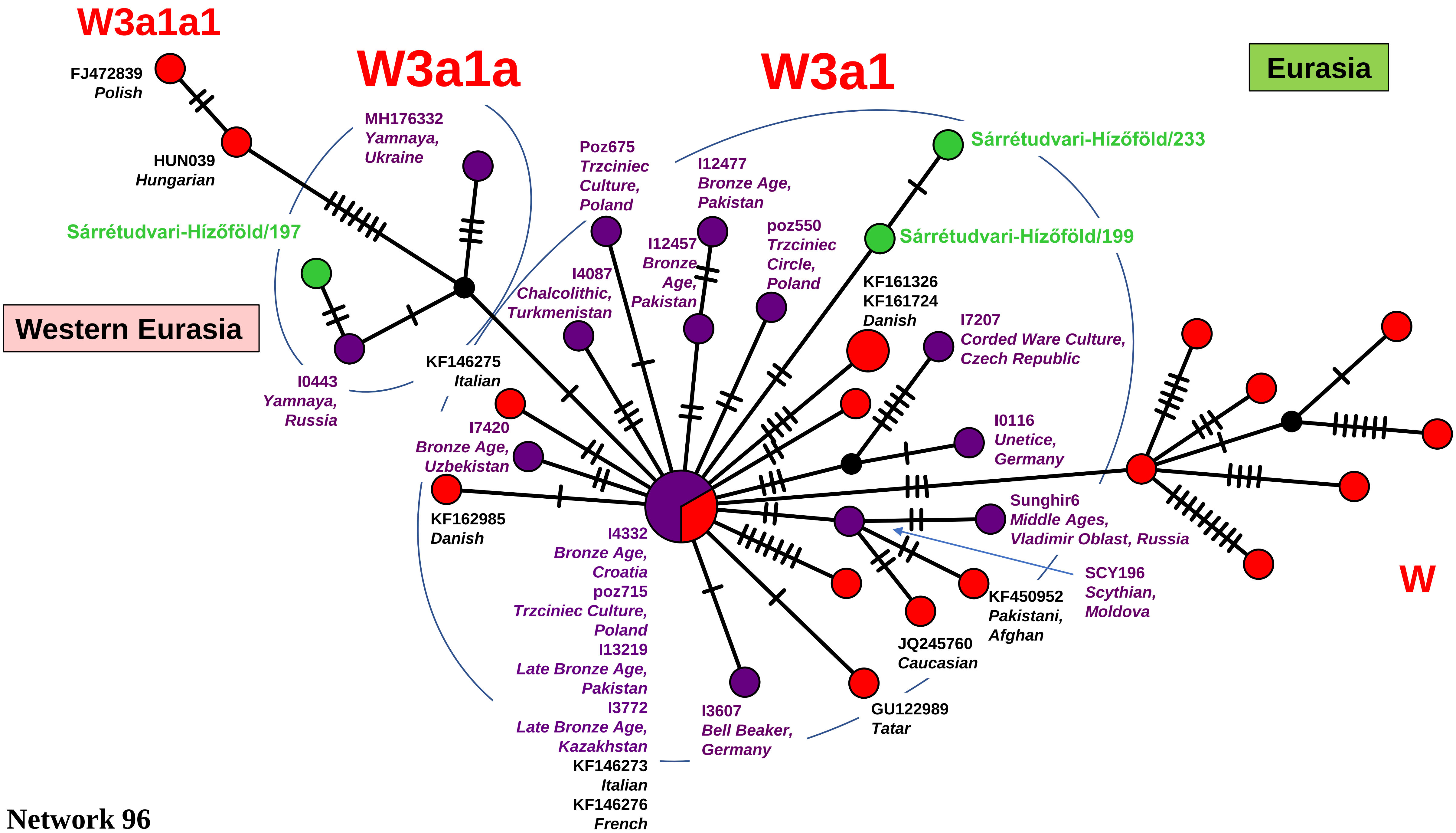

W3a1a1
FJ472839
Polish
HUN039
Hungarian
W3a1a
MH176332
Yamnaya,
Ukraine
W3a1
Eurasia
Sárrétudvari-Hízőföld/233
Poz675
Trzciniec
Culture,
Poland
I12477
Bronze Age,
Pakistan
Sárrétudvari-Hízőföld/197
Sárrétudvari-Hízőföld/199
I12457
Bronze
Age,
Pakistan
poz550
Trzciniec
Circle,
Poland
I4087
Chalcolithic,
Turkmenistan
KF161326
KF161724
Danish
Western Eurasia
I7207
Corded Ware Culture,
Czech Republic
KF146275
Italian
I0443
Yamnaya,
Russia
I7420
Bronze Age,
Uzbekistan
I0116
Unetice,
Germany
KF162985
Danish
Sunghir6
Middle Ages,
Vladimir Oblast, Russia
I4332
Bronze Age,
Croatia
poz715
Trzciniec Culture,
Poland
I13219
Late Bronze Age,
Pakistan
I3772
Late Bronze Age,
Kazakhstan
KF146273
Italian
KF146276
French
SCY196
Scythian,
Moldova
KF450952
Pakistani,
Afghan
W
JQ245760
Caucasian
I3607
Bell Beaker,
Germany
GU122989
Tatar


Network 96

Western Eurasia

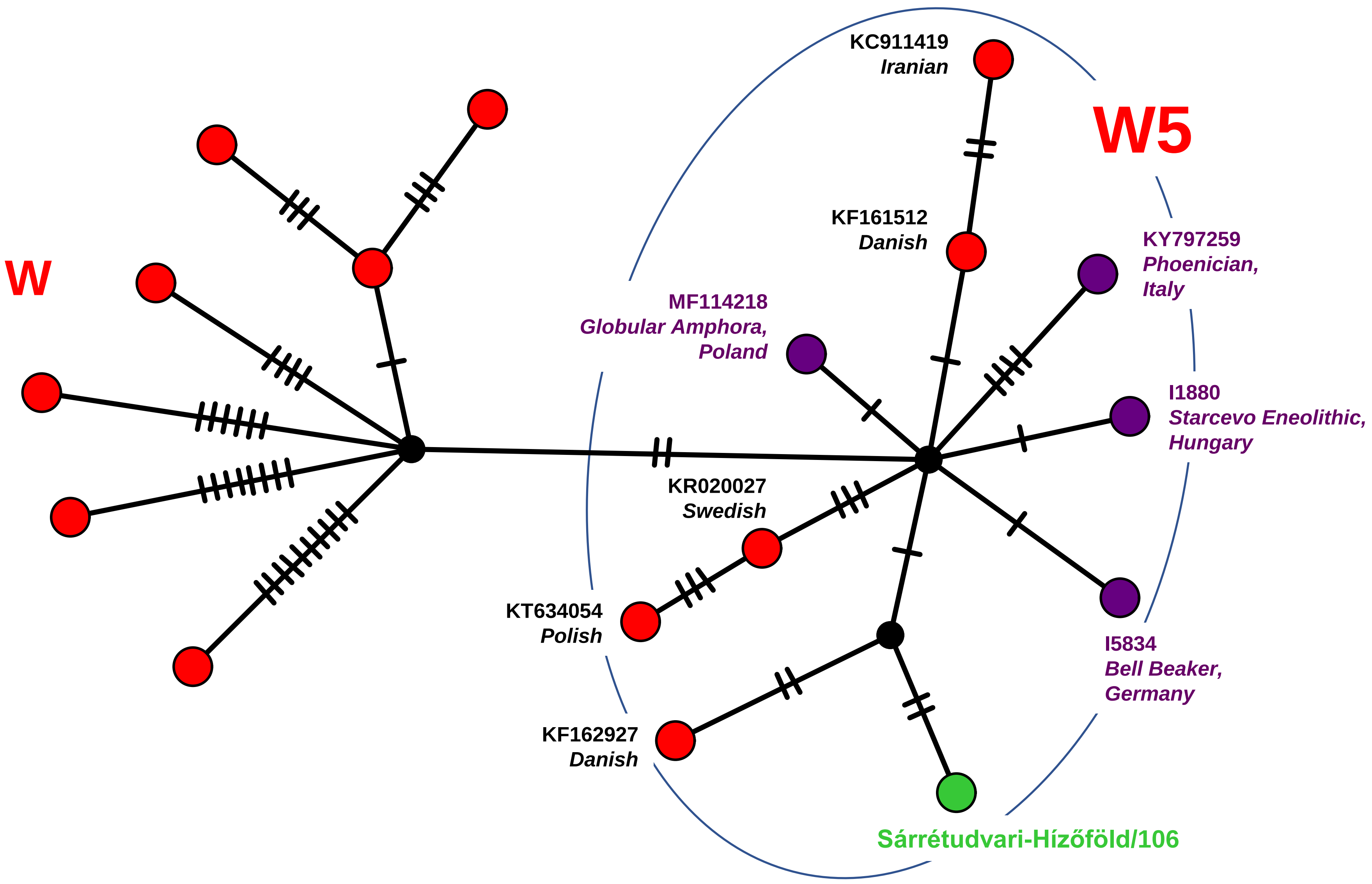


Network 97

**Caucasus/Middle East**

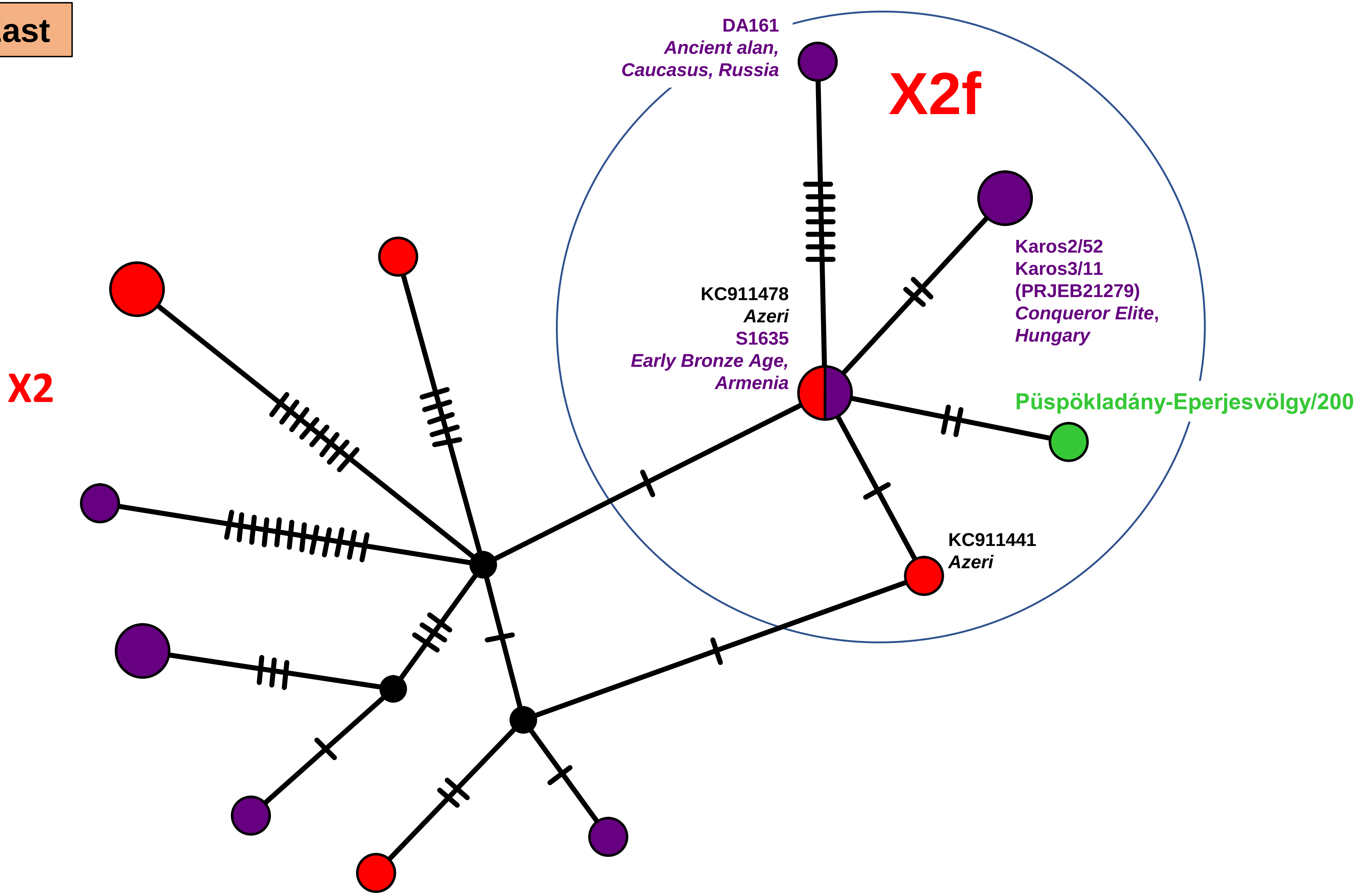


**Network 98**
