## Appendix A for "Maternal lineages from 10-11^th^ century commoner cemeteries of the Carpathian Basin"

**Basic description of the archeological categorization problems and the studied cemeteries.** (Detailed archaeological and anthropological description of each graves is provided in Table S1)

### 1. A brief summary of the problems associated with the archaeological categorization of 10-11th-century cemeteries in the Carpathian Basin

Assessment of the archaeological horizon, cemeteries and individual burials of the 10-11th-century Carpathian Basin which is the period of the Hungarian conquest and state formation, have undergone significant changes over the past 150 years. The first major summary and categorization of the finds was made by József Hampel [1] based on their dating patterns (tombs that can be dated with a coin, tombs that cannot be dated with a coin, and stray finds). However, this was soon replaced by his new classification based on ethnicity in which two groups were distinguished: the findings of newly arrived conquerors (the time covers ca.150 years, 895-1050), buried with horse riding- and weapon-related grave goods, often with coins of foreign origin (Western-European, Byzantian, Arabian) referred as Group A, and the local population resting in cemeteries composed of row of graves, buried with simpler jewels and grave goods, and often with coins of the kings of the House of Árpád referred as Group B. The latter group later spread in the public consciousness as the Bijelo Brdo culture and was defined as having Slavic ethnicity.

In the first half of the 20th century the number of archaeological findings of the 10-11th century period increased rapidly. During the collection and systematization of these finds, Béla Szőke noticed that Group A and Group B can not be sharply separated because the findings of the two groups often appear simultaneously in the cemeteries and the cemeteries composed of row of graves are also present in the central areas of the 10$^{th}$-century Hungarian territory. Based on his observations, he believed that the differences did not indicate different ethnicities, rather they reflect the social status. Thus he developed a new kind of systematization in which the ethnic-based classification was replaced by a division into social categories [2]. Hampel's group A was identified with the leading and middle class of the conquerors, and in connection with Hampel's group B, he assumed that those are the findings of the commoner people of the 11th century who were mostly of Hungarian origin, but often mixed with other ethnicities (e.g., in the peripheral areas). However, the analysis of the archaeological material related to the large-scale excavations of the second half of the 20th century already highlighted the problems of the method and its categories (e.g.,[3]), Szőke's system has been used until the last decades to classify individual burials and cemeteries. In addition to the fact that international research has shown that it is not possible or very difficult to distinguish legal and social categories using archeological methods alone [4,5], the social categories have not been sufficiently defined, so in the field of practice cemeteries were classified in a highly subjective way. On the other hand, the size of the cemetery significantly influenced the evaluation of the findings (e.g., the presence of 3-4 graves with horse riding- and weapon-related grave goods in a cemetery of 10-20 graves already elevated the given cemetery to the middle class, however, the same ratio in a cemetery composed of 100 graves resulted in the site being classified as a commoner cemetery). However based on the most recent investigations, only a low percentage of the 10th century cemeteries can be considered as fully excavated, which often resulted in erroneous conclusions [3]. In addition, frequently the cemeteries contained a mix of "rich" and "poor" grave goods or even burials without known grave goods, which drew attention to the fact that a cemetery may not necessarily correspond to a particular social category.

István Bóna attempted to reinterpret the 10-11th-century cemeteries, distinguishing two type of cemeteries [6]. The so-called military cemeteries are characterized by the rather low number of graves covering one or two generations and often showing a male surplus, which he considers were mostly used in the first half of the tenth century. While the larger village cemeteries with 30-100 graves were used later, during the 10-12th century respectively.

This criteria system was further developed by László Kovács, who, breaking with the ethnic and social classification, based the assessment of the sites on the chronological characteristics and quantity of the burials [7]. Within the archaeological horizon of the 10-12th centuries, he distinguished two main types in connection with the supposed form of the possibly related settlement: the so-called quarter and village cemeteries. The quarter group included the smaller cemeteries with 5/10-50/75 burials, which were used for a shorter time period and were abandoned between the second half of the 10th century and the turn of the millennium. These cemeteries characterized by mostly the types of objects and burial customs associated with the conquerors, and at the same time the coins of the kings of the Árpád Dynasty are completely missing. The other main group, the village cemeteries, is characterized by a larger number of graves of 50-200 and above, and their use can be traced back to the 10th, 10-11th, and 10-12th centuries, depending on the subtype. In connection with the grouping, it may be a problem to separate the 10th century quarter type and the 10th century village type, as there are overlaps based on the time of use and the number of graves, so sometimes the richness and quality of the finds may make the difference [3], but due to problems with the representational value and symbolic aspects of the objects [8,9] may mislead the conclusions. Questions arise relating to the evaluation of cemeteries dating from the 10th century which were in use through the 11-12th centuries, because based on the written sources concerning the period of the foundation of the Hungarian Kingdom and the early Árpádian age, as well as the results of micro-regional archeological research, significant internal population movements and organized settlements must be taken into account. With the help of archaeological methods alone it is not possible to determine beyond any doubt whether the $10^{th}$-century and later cemetery parts are contiguous with each other or whether a population change has taken place in the meantime [3]. However, this can also significantly affect the grouping of cemeteries (e.g., a site can be assessed as a larger 10-11th century village cemetery or as a separate 10th century quarter/village cemetery and a 11th century village cemetery too).

Several methods have been developed since the beginning of the research to categorize 10-11/12th century cemeteries and burials. Each method has its advantages and disadvantages depending on the issues raised by the particular research. Aware of all this, during the short archaeological summary of the cemeteries involved in our research, we tried to describe their characteristics objectively, but in each case, we indicated where the given site was classified in the Szőke and Kovács classifications.

### 2. Archaeological and anthropological description of the studied cemeteries

#### Homokmégy-Székes (Bács-Kiskun County)

The excavation of the site was carried out by Zsolt Gallina and Sándor Varga between 1996-2002 [10]. The cemetery which consisted of 206 graves is divided into northern and southern parts based on the type and the orientation of the graves, the grave goods, and the position of the arms. The northern part is further splitted into east and west sides based on the density of the graves. Due to the characteristics of the soil, the shape of the graves has been preserved well: several grave pits with sidewall niche have been found, as well as traces of other Avar age burial customs, which are quite rare in the $10^{th}$ century (e.g., patterns of post-holes in the grave pit). The archeological findings include jewelry and clothing ornaments for example hoops around the head (e.g., S-terminalled and penannular hair rings, earrings), neck jewelry (e.g., neckrings, beads), and arm jewelry (e.g., bracelets, rings), dress fittings, as well as weapons (archery equipment, ax) and implements (e.g., fire-lightning equipment, knife).

Based on the composition of the findings, the cemetery dates back to the period from the second third of the 10$^{th}$ century to the first third of the 11$^{th}$ century. It is believed that the first generation may still have belonged to the conquest period, and the last generation may have been buried there during the reign of king (St.) Stephen I. According to a former classification based on the size of the cemetery and the composition of the findings the cemetery was described as „commoner cemetery”. In a more recent classification, the cemetery belongs to the group of the 10$^{th}$-century village cemeteries.

The state of preservation of the anthropological material is generally good or medium, excluding the sub-adult skeletons. During the anthropological examinations [11], 136 adult and 50 sub-adult remains were distinguished of which 63 was described as male and 83 as female. Based on the distribution of taxonomic features, in addition to the predominance of Europids (88%), a smaller proportion of components belonging to the Mongoloid type also appears (9.3%). Based on biological distance calculations, the population of the cemetery shows similarities to other 10$^{th}$ century and Avar-era series.

**Ibrány-Esbóhalom (Szabolcs-Szatmár-Bereg County)**

The excavation of the cemetery was carried out between 1985 and 1990 under the leadership of Eszter Istvánovits [12]. The remains of 274 individuals were found in 269 graves. Based on the finds, the 10$^{th}$-century part of the cemetery can be well characterized by the low number of burials with weapon- and horse riding-related grave-goods and dress fittings (e.g., pendent mounts), as well as simple wire jewelries (e.g., bracelets) and implements (e.g., knives, fire-lightning equipment). Within the 10$^{th}$-century part, a group with different ethnicities is also distinguished, based on the grave orientation, burial customs, and findings. In addition to the coins related to the reign of the kings of the Árpád dynasty, the burials of the 11$^{th}$ century are characterized by S-terminalled hair rings, beads and rings. In virtue of the archaeological material, continuity was assumed between the two parts of the site. The use of the cemetery dates back to 940-1075. Based on the number of the graves and the composition of the archaeological material, it was characterized as a commoner cemetery. The state of preservation of the anthropological material is medium, often poor. In anthropological studies [13] 98 men, 82 women, 74 children (inf I-II) and 20 young (juvenile) individuals have been distinguished, thus there is a male surplus for adults. Taxonomic analysis revealed a predominance of Europeans, Mongoloid traits were observed in case of 4 individuals. Craniometric analysis showed a discrepancy between the 10$^{th}$ and 11$^{th}$ century parts, raising the question that the two cemetery parts may hide different populations.

**Magyarhomorog-Kónyadomb (Hajdú-Bihar County)**

The systematic excavation of the cemetery was started by István Dienes between 1961-1971, then it was finished by László Kovács between 1985-1988 [14]. The cemetery has a „pagan" segment with 17 graves from the 10th century characterized by a high number of burials with weapon- and horse riding-related grave-goods, and a significant male surplus. The larger part consists of 523 „Christian" graves of the 11-12th century in which the gender rate is more balanced. The cemetery is one of the few sites where the burial custom of giving tomb furnishing can be observed after the turn of the millennium and the adoption of Christianity, (e.g., weapon- and horse riding-related grave-goods in the tombs dated with coins related to the reign of the Árpád dynasty kings), which was forbidden by the Christian liturgy. The different types of jewelry were the most common archaeological findings: hoops around the head (e.g., S-terminalled hair rings), neck jewelry (e.g., beads), bracelets and various types of rings. Based on the location of the tombs containing coins, the cemetery started from a central core and expanded evenly towards its edges. During the archaeological analysis, the possibility of discontinuity between the two parts of the cemetery arose. Therefore, the archaeologists suggest that the cemetery is separated to a 10$^{th}$-century so-called quarter cemetery and a 11-12$^{th}$-century village cemetery part. During the anthropological analysis [15], 11 men, 3 women, and 3 children of unknown sex were identified in the 10$^{th}$-century part. The state of preservation was generally of medium quality. Based on the taxonomic analysis, the europid groups dominated (about 60%), but overall, the proportion of those showing Europo-Mongoloid traits is significant and one individual was classified as Mongoloid type. In case of paleopathological alterations, developmental abnormalities predominated, which address several further questions about kin relationships within the group. Concerning the 11-12$^{th}$-century village cemetery, the state of preservation of the skeletons was generally low. 126 males, 174 females, 187 adolescents (infantia I-II), and 36 juvenile individuals were distinguished. According to the craniometric analysis, the dolichocran skulls were dominant, and the taxonomic analysis revealed that skulls with europid characteristics were in the highest number, but also europo-mongolid and in 6 cases mongolid types were also present (the state of preservation highly limited the classification). In our analysis, we examined whether the "Christian" part could have been contiguous with the "pagan" part, i.e., it hides descendants of the same population or its population originates from elsewhere. Special attention should also be paid to the possible connections of the previously described cemeteries of Karos and Kenézlő of a similar age.

**Nagytarcsa-Homokbánya (Pest County)**

In 1967, under the leadership of László Kovács 21 graves were excavated and there are data on another 7 disturbed graves, but the cemetery can only be considered as partially excavated (estimated at about 40-50 graves) [16]. The poor archeological findings consisted of penannular hair rings, twisted neckrings, wire bracelets, rings, and ball buttons. Two burials with weapon related grave-goods (archery equipment, an ax) and four burials with horse riding-related deposit (e.g., pear-shaped stirrups) were excavated in the cemetery. On the grounds of the findings the cemetery was dated to the second half of the 10th century. The state of preservation of the anthropological material is moderate, often low. The anthropological analysis [17] identified the skeletons of 8 male, 15 female, and 4 unspecified children. Five skulls belonging to the europid type proved to be suitable for taxonomic analysis.

**Püspökladány-Eperjesvölgy (Hajdú-Bihar County)**

The excavation was carried out under the leadership of Ibolya M. Nepper and Márta Sz. Máthé between 1977-1982 [18]. 637 graves were found in the cemetery, but due to double burials the remains of 641 individuals were identified. Based on the findings, the cemetery can be divided into 2 parts. The „pagan" western part (about one-third of the cemetery) dating back to the 10$^{th}$ century is characterized by burials with weapon- (e.g., archery equipment, sabre, sword) and horse riding-related (pear-shaped stirrups) grave-goods, as well as jewelry - hoops around the head (e.g., penannular hair rings), necklaces (e.g., beads), arm/hand jewelry (e.g., bracelets, rings), dress fittings and implements (e.g., knife, fire-lightning equipment). The other part was likely used after the adoption of Christianity in the 11th century, based on the more common occurrence of coin-dated burials and relatively late grave-good types (e.g., twisted and braided rings, foil beads, S-terminalled hair rings). On the strength of the size of the cemetery and the composition of the archaeological material, this cemetery is belonging to the commoner cemeteries in the former classification and to the 10-11$^{th}$-century village cemeteries in the more recent classification. The anthropological material is of medium or poor preservation. During the anthropological analysis of the cemetery [19,20], 191 male, 163 female, and 256 sub-adult (unspecified sex) individuals were described. According to studies on craniometric and body height data, continuity was assumed between the 10th-century and 11th-century parts of the cemetery.

**Sárrétudvari-Hízóföld (Hajdú-Bihar County)**

The site was excavated between 1980 and 1985 under the leadership of Ibolya M. Nepper [18]. The site with 262 graves is considered the largest 10th-century cemetery in Hungary. The cemetery contains very high proportion of burials with weapon- (archery equipment, sabers, ax) and horse riding-related (eg: pear-shaped stirrups) grave-goods, and the archeological findings consist of jewelry - hoops around the head (penannular hair rings), neck jewelry (neckrings, beads), arm jewelry (e.g., bracelets, beads), - dress fittings, and implements (e.g., knives, fire-lightning equipment). Based on the composition of the findings and the lack of coins and grave-goods dated to the reign of the kings of the Árpád dynasty, the cemetery can be dated to the 10th century. Formerly it was classified as a commoner cemetery, and in the new classification it belongs to the 10th-century village cemeteries.

The skeletal remains are of good / medium preservation. During the extensive anthropological analysis (e.g., [21]), 265 individuals were determined, of whom 98 belonged to sub-adult- and 162 skeletons to adult categories. Based on the skulls suitable for taxonomic studies, the series shows European characteristics with the presence of cromagnoid and nordoid elements.

**Szegvár-Oromdűlő (Csongrád County)**

The site contained 372 graves which were excavated under the leadership of Gábor Lőrinczy between 1980/1983 and 1996 [22], but many burials (estimated about 75-85) were destroyed due to previous disturbances. 5 additional graves were excavated 30-40 m away from the tight array of the cemetery. The archaeological material is characterized by jewelry - hoop jewelry around the head (penannular hair rings, coiled hair rings, S-terminalled hair rings, hoop with spiral pendant), neckrings, bracelets, rings, less frequently implements (knives, fire-lightning equipment), and in one grave dress fittings were excavated. The use of the cemetery dates back to the period between the second third of the 10th century and the middle of the 11th century. Based on its size and the composition of the grave goods, the cemetery was classified formerly as a commoner cemetery of the 10-11th century and as a 10-11th-century village cemetery recently.

In anthropological studies [23], skeletons of 110 male, 114 female, and a total of 148 sub-adults of indeterminate sex were described. During the taxonomic analysis, the predominance of europid type skulls (cromagnoid, nordoid) was detected in both the 10th- and 11th-century groups, but a small proportion of Mongoloid features were also described. Based on the craniometric data, it is assumed that a population change occurred at the turn of the 10-11$^{th}$ centuries.

**Szegvár-Szőlőkalja (Csongrád County)**

The site of 62 graves containing the burials of 63 individuals was excavated in 1979 by Katalin Hegedűs [24]. A burial of opposite orientation (E-W) was found at a distance of 30-35 m from the cemetery array. The poor archaeological findings consist of penannular hair rings, beads, wire bracelets, ball buttons, and knives. Horse riding-related equipment (a fragment of a bridle) and a weapon (arrowheads) were unearthed in one burial. Based on the composition of the archaeological material, the site dates back to the 10$^{th}$ century and it was classified as a commoner cemetery formerly and 10$^{th}$-century villager cemetery recently.

During the analysis of anthropological findings [24], 25 male and 25 female skeletons were described. Based on taxonomic studies, the population composition is heterogenous, the individuals belonged to the europid, mongoloid and europo-mongoloid types.

**Vörs-Papkert (Somogy County)**

The cemetery was excavated between 1983 and 1993 under the leadership of László Költő, Szilvia Honti and József Szentpéteri [25]. The cemetery as a whole is still unpublished, but it is dated back from the turn of the 8-9th centuries to the turn of the 10-11th centuries. The 716 excavated burials are mostly from the Late Avar and Carolingian periods, but sporadic burials of 33 people can be dated to the time of the Hungarian conquest. Therefore, the study of the cemetery could help investigate the supposed survival of the Transdanubian late Avar population through the 9-10th centuries. Extensive anthropological and serological investigations were carried out at the cemetery [26,27], but contradictory results were obtained (e.g., concerning the sex determination).
